# Nucleus size and its effect on the chromatin structure in living cells

**DOI:** 10.1101/2021.07.27.453925

**Authors:** Artem K. Efremov, Ladislav Hovan, Jie Yan

**Affiliations:** Institute of Systems and Physical Biology, Shenzhen Bay Laboratory, Shenzhen, China 518132; Mechanobiology Institute, National University of Singapore, Singapore 117411; School of Pharmaceutical Sciences, University of Geneva, Geneva, Switzerland 1211

**Keywords:** DNA, DNA-protein interaction, DNA-binding protein, DNA packaging, polymer field theory

## Abstract

DNA-architectural proteins play a major role in organization of chromosomal DNA in living cells by packaging it into chromatin, whose spatial conformation is determined by an intricate interplay between the DNA-binding properties of architectural proteins and physical constraints applied to the DNA by a tight nuclear space. Yet, the exact effects of the cell nucleus size on DNA-protein interactions and chromatin structure currently remain obscure. Furthermore, there is even no clear understanding of molecular mechanisms responsible for the nucleus size regulation in living cells. To find answers to these questions, we developed a general theoretical framework based on a combination of polymer field theory and transfer-matrix calculations, which showed that the nucleus size is mainly determined by the difference between the surface tensions of the nuclear envelope and the endoplasmic reticulum membrane as well as the osmotic pressure exerted by cytosolic macromolecules on the nucleus. In addition, the model demonstrated that the cell nucleus functions as a piezoelectric element, changing its electrostatic potential in a size-dependent manner. This effect has been found to have a profound impact on stability of nucleosomes, revealing a previously unknown link between the nucleus size and chromatin structure. Overall, our study provides new insights into the molecular mechanisms responsible for regulation of the cell nucleus size, as well as the potential role of nuclear organization in shaping the cell response to environmental cues.

**SIGNIFICANCE STATEMENT:** The cell nucleus plays a central role in the life of eukaryotic cells, providing the highest level of control of intracellular processes. Depending on the stage of the cell cycle and / or surrounding environment, the size of the cell nucleus may undergo changes that are believed to cause chromatin reorganization, affecting gene transcription. At present, however, there is no clear understanding of the molecular mechanisms that may be responsible for such regulation, whose exact effect on chromatin structure remains unclear. In this study, by developing an advanced computational approach, we explore these issues from a physical perspective, revealing previously unknown mechanisms contributing to organization of the cell nucleus and chromatin.

## INTRODUCTION

Homeostasis and biological functioning of living cells rely on sophisticated synergistic cooperation between multiple molecular subsystems that must coexist with each other in a tight and highly crowded cellular space. This makes the problem of space allocation to each of the cell components one of the most important for intracellular organization, especially taking into account that many of the subcellular systems have very different requirements to the surrounding microenvironment needed for their proper operation. However, at the present time, there is still no clear understanding of molecular mechanisms responsible for size regulation of the most of cellular organelles, including even the major ones, such as the cell nucleus [1–4].

Indeed, while many models with different levels of detail have been developed in the past to describe regulation of the cell volume [5–7], no similar analogue has been created so far to address the problem of the nucleus size control. As a result, such a very well-known phenomenon as correlation between the cell and nucleus volumes under a broad range of experimental conditions still remains poorly understood [8–11]. Furthermore, lack of the knowledge of the main molecular mechanisms responsible for the nucleus size regulation makes it hard to fully comprehend potential effects of various environmental factors in shaping nuclear organization.

Specifically, based on recent experimental studies it has been suggested that by modulating DNA-binding properties of architectural proteins, cells may be able to actively respond to a wide range of environmental cues via changing the condensation level of various parts of DNA and switching between different gene expression patterns [12–18]. Furthermore, it has been found that such a mechanism not only plays the central role in cell response to chemical signals [12–14], but also to extracellular mechanical forces as well [16–25]. In the latter case, rearrangement of the chromatin structure and changes in gene transcription have been found to be tightly associated with alternations in the nucleus size and shape [18, 21, 25]. Yet, despite a profound role of mechanical forces in regulation of the nuclear organization, the exact molecular mechanisms underlying it still remain unclear.

The main difficulty in understanding the role of mechanical forces in shaping the chromatin structure comes from the fact that the latter is predominantly determined by a tight interplay between several key factors, whose exact contributions, however, cannot be easily quantified in experimental studies. Namely, physical constraints applied by the nuclear envelope (NE) to DNA results in appearance of highly crowded microenvironment inside the cell nucleus, whose behaviour in a large part is driven by electrostatic and volume-exclusion interactions between DNA and proteins [26–32]. The strength of these interactions is determined by nucleoplasmic concentrations of DNA and proteins, which are dictated by the nucleus size. As it is hard to reproduce similar unique microenvironment in *in vitro* studies with the same DNA and protein composition, the main understanding of the role of such physical interactions as well as their interplay with mechanical forces in chromatin organization currently can be gained only by means of theoretical modelling.

Unfortunately, extreme complexity of this task and strong limitations of the existing computational methods do not allow to run molecular dynamics simulations for long DNA molecules comparable in size to the cell genome. Even the most advanced modern mathematical models, which heavily rely on coarse-grained description of chromatin, can currently predict an approximate conformation of DNA molecules no longer than ~ 100 Mbp in length [33–46], – about two orders of magnitude smaller than the typical size of chromosomal DNA in human cells (~ 6.2 Gbp [47]). And while these models have provided many interesting insights into molecular mechanisms underlying large-scale chromatin organization, it should be noted that due to their coarse-grained nature, they have a number of limitations.

First, they do not take in consideration one of the major physical forces being involved in chromatin organization – electrostatic DNA-DNA and protein-DNA interactions, which make a significant contribution into the total free energy of chromatin, dominating over all other energy contributions made by the rest of physical and chemical factors. As a result, the role of intracellular ionic environment in chromatin organization still remains poorly understood despite experimental studies showing that it plays an important role in shaping the chromatin structure [26–30, 32, 48].

Second, use of extend segments of the chromatin fiber as elementary modelled units makes it nearly impossible to take into consideration formation and dissociation of individual nucleoprotein complexes from DNA as well as their transition between various states, – processes which play the central role in governing genome-wide changes in chromatin organization in response to environmental cues [12–14, 17, 18, 49]. Thus, capacity of the existing models to address in detail the core property of chromatin – its ability to dynamically reorganize in response to signals received from surrounding microenvironment, is compromised by application of the coarse-grained approach that lacks insights into behaviour of individual nucleoprotein complexes.

Finally, it should be noted that the possible number of ways nucleoprotein complexes, such as nucleosomes, can be positioned on the chromosomal DNA is extremely large (≫ 10^1000000^). Since none of the existing models describing chromatin organization can deal with so many possible DNA-protein conformations, it seems that the above problem cannot be solved on the basis of currently used theoretical methods, and an alternative approach needs to be developed.

In this study, by utilizing elements of polymer field theory and statistical physics, we have constructed a general theoretical framework that allows one to address these issues and predict the size of the cell nucleus and the conformation of chromosomal DNA, taking into account DNA-protein interactions and the main physical forces responsible for nuclear organization. As a result, it has been found that the nucleus size in the general case is predominantly determined by a tug-of-war between the osmotic pressure exerted by cytosolic macromolecules on the NE and the difference between the surface tensions of the NE and endoplasmic reticulum (ER) membrane. In addition, the model showed the existence of a previously unknown link between the chromatin structure and the nucleus size, revealing a new potential molecular pathway, which may contribute to cell mechanosensing.

## METHODS

In this work, we used the previously developed theoretical framework based on transfer-matrix calculations that allows one to compute the grand partition function of DNA in the presence of DNA-binding proteins [50–52], extending it to include in consideration physical forces discussed in *Introduction.* While all mathematical details can be found in Appendices A-M, here we will only outline the central idea of the study and the main model assumptions.

Packaging of long chromosomal DNA into a tiny nuclear space in eukaryotic cells is mainly done with the help of special DNA-architectural proteins, histones. These proteins assemble into positively charged octamer complexes that wrap negatively charged DNA around themselves, leading to formation of compact nucleosomes. Nucleosome assembly is performed via help of histone-binding chaperones, such as Asf1 or Nap1, which are involved in transportation of H2A·H2B and H3·H4 histone dimers into the cell nucleus [53, 54]. Correspondingly, in the model we considered two types of chaperones: 1) c_1_ chaperones that can either be in empty / unloaded state (c_1_u) or histone-bound state (c_1_b), in which they form a complex with a H2A·H2B dimer, and 2) c_2_ chaperones that also can be either in unloaded state (c_2_u) or histone-bound state (c_2_b), but this time forming a complex with a H3·H4 dimer, see schematic Figure 1(a). Since experimental data suggest that some histone-binding chaperones, such as Nap1, shuttle between the nucleus and cytosol in eukaryotic cells [55], in our model it was assumed for simplicity that both histone-loaded and unloaded chaperones c_1_ and c_2_ can move through nuclear pore complexes (NPCs) without any restriction, see also Appendix M for more details.

**FIG. 1.**
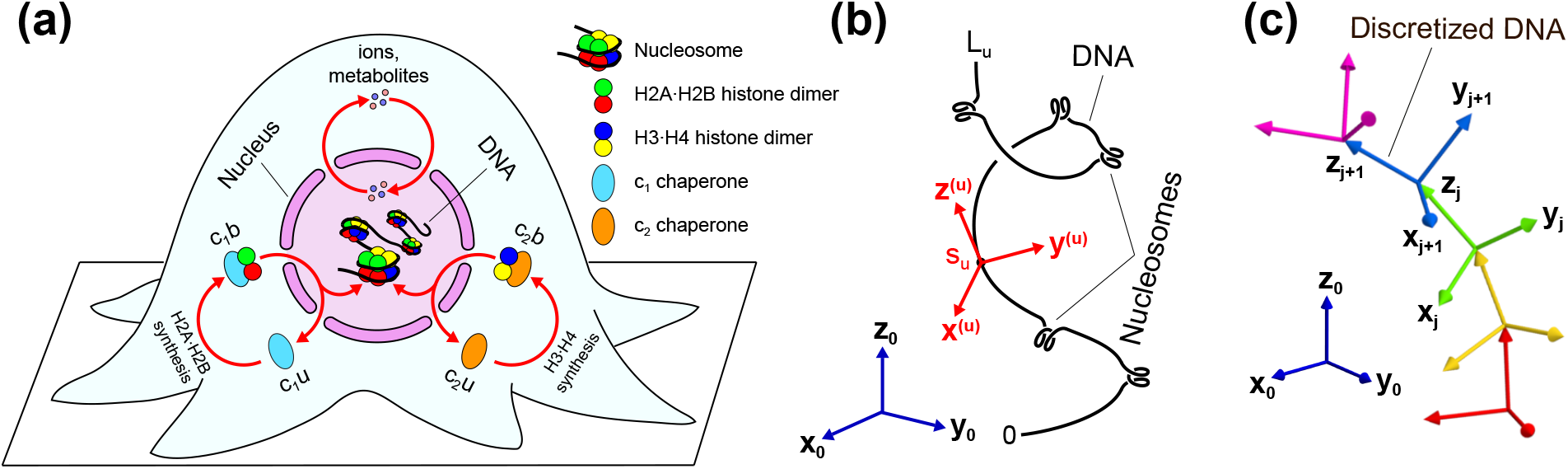
Semiflexible polymer model of chromosomal DNA packaging in the nucleus of a eukaryotic cell. **(a)** Main protein complexes involved in histone transportation and nucleosome formation. In the model, nucleosomes assemble on DNA from H2A·H2B and H3·H4 histone dimers, which are transported by histone-binding chaperones schematically indicated as c_1_ and c_2_ in the figure. Specifically, by picking up newly synthesised histone dimers in the cell cytoplasm, chaperones switch from an unloaded state (c_1_u or c_2_u) to a histone-bound state (c_1_b or c_2_b, respectively). Then by moving through NPCs, chaperones enter the the cell nucleus, where they can deposit histone dimers onto the chromosomal DNA via nucleosome assembly. In addition to histone-binding chaperones, NPCs also allow free diffusion of ions and small metabolites between the cell nucleus and cytosol, helping to maintain stability of the ionic microenvironment in the nucleoplasm. **(b)** Chromosome conformations. In the model, conformations of chromosomes were described by triplets of unit vectors, (**x**^(*u*)^ (s_*u*_), **y**^(*u*)^ (*s_u_*), **z**^(*u*)^ (*s_u_*)), *u* = 1,…, *Q*, with each triplet indicating the orientation of the respective (u^th^) DNA molecule with reference to the fixed coordinate system, (**x**_0_,**y**_0_,**z**_0_), at a point corresponding to the arc length *s_u_* ∈ [0,*L_u_*] along the DNA. Here *L_u_* is the length of the *u*^th^ DNA molecule. **(c)** Discretized polymer model of DNA. To calculate the partition function, each DNA polymer was represented by a polygonal chain comprised of small straight segments, each of which was treated as a rigid body with an attached local Cartesian coordinate frame, (**x**_*j*_, **y**_*j*_, **Z**_*j*_), representing the DNA segment orientation in space with respect to the fixed global coordinate system, (**x**_0_, **y**_0_, **z**_0_). More details regarding description of DNA and nucleoprotein complexes can be found in Appendices A-B.

In the model, the cell nucleus is represented by a spherically shaped NE enclosing nucleoplasm of *V*_nucl_ volume. The nucleus itself is submerged into the cell cytosol of *V*_cyto_ volume, which may have an arbitrary shape. Cytosol in the model serves as a buffer solution cushioning changes in the nucleoplasmic concentrations of unloaded and histone-bound chaperones that may result from rearrangement of the chromatin structure.

Besides chaperones, the cytosol also buffers any changes in the nucleoplasmic levels of Na^+^, K^+^ and Cl^-^ ions, which have a major effect on the strength of DNA-histone interactions and chromatin organization [28, 30, 32]. Microelectrode measurements as well as radioautographic and extractive analysis suggest that these monovalent ions can rather freely move between the cell nucleus and cytosol [56, 57]. Furthermore, their cytosolic concentrations are usually kept at nearly constant levels by transmembrane ion pumps and ion channels to maintain electroneutrality of intracellular environment and at the same time to counterbalance osmotic pressure created by the cell metabolites and proteins onto the cell membrane [6, 7, 58]. Following previously published experimental and theoretical studies [5, 58, 59], the total cytosolic concentrations of monovalent Na^+^ and K^+^ ions on one hand, and Cl^-^ ions and negatively charged cell metabolites on the other hand were both put equal to *c*_ions_ = 150 mM in all our calculations.

As for the chromosomal DNA, it was modelled as a set of polymers constrained inside the cell nucleus, whose total number (*Q*) matches the number of chromosomes in the studied eukaryotic cells (in human cells *Q* = 46), with the total length of DNA pieces 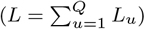 being equal to the total size of the cell genome. Here *L_u_* is the length of the *u*^th^ DNA polymer comprising the *u*^th^ chromosome. Thus, to describe chromatin organization in human cells the total length of all DNA polymers was set equal to *L* = 6.2 Gbp in all calculations [47].

Three-dimensional conformations of the DNA polymers were represented in the model by **R**^(*u*)^(*s_u_*), *u* = 1,…,*Q*, functions, where *s_u_* ∈ [0, *L_u_*] is the arc length along the *u*^th^ DNA polymer and **R**^(*u*)^ is a three-dimensional Euler rotation matrix assigned to each point residing on the contour of the *u*^th^ chromosome, which is a function of the arc length, *s_u_*. This matrix indicates orientation of the DNA backbone at the corresponding point with respect to the global coordinate system, (**x**_0_, **y**_0_, **z**_0_), such that the unit vector **z**^(*u*)^(su) = **R**^(*u*)^(*s_u_*) **z**_0_ resulting from the rotation of **z**_0_-axis of the global coordinate system via Euler matrix **R**^(*u*)^(*s_u_*) is tangential to the DNA backbone; whereas, the other two unit vectors, **x**^(*u*)^(*s_u_*) = **R**^(*u*)^(*s_u_*) **x**_0_ and **y**^(*u*)^(*s_u_*) = **R**^(*u*)^(*s_u_*) **y**_0_, are normal to the DNA backbone, keeping track of the DNA twist angle, see schematic Figure 1(b).

Nucleosomes formed on the chromosomal DNA were represented in the model by solenoid-like structures schematically shown in Figure 1(b), whose geometry matches that of DNA wrapped around histone octamers in nucleosome complexes [60, 61].

To find out how confinement of DNA inside the cell nucleus as well as electrostatic interactions between the system components affect the chromatin conformation, we used statistical physics approach by calculating the partition function of the system. Indeed, while living cells usually operate far from thermodynamic equilibrium, previous studies suggest that DNA interaction with proteins still can be accurately described by equilibrium thermodynamics since assembly of nucleoprotein complexes on DNA is frequently governed by DNA-binding affinities and positioning entropies of the proteins [62–66]. Moreover, it has been shown in previous theoretical studies that methods of statistical physics seem to correctly predict positions of nucleosomes on the chromosomal DNA near transcription start sites as well as on the global genomic scale [67–71], suggesting that they can be used to provide accurate insights into organization of chromosomal DNA in living cells. This approach is further supported by multiple theoretical studies of a large-scale chromatin organization based on thermodynamic equilibrium calculations / simulations, which demonstrated good agreement with numerous experimental observations, see, for example, ref. [33, 34, 36, 38, 40–42, 44–46]. Furthermore, simple estimations based on existing experimental data indicate that, under physiological conditions, active processes underlying nuclear transport of proteins through NPCs should have a negligible effect on histone-binding chaperones shuttling between the cell cytosol and nucleus, whose nucleoplasmic and cytosolic concentrations appear to closely follow the Boltzmann distribution (see Appendix M). Altogether, these results suggest that the statistical physics / equilibrium thermodynamics approach provides an accurate description of the chromosomal DNA organization in nuclei of living cells, which for this reason was adopted in our study.

To calculate the partition function (*Z*) of DNA polymers described above, we utilized an implicit solvent method, which leads to the following formula [see ref. [72] and Appendix A for more details]:

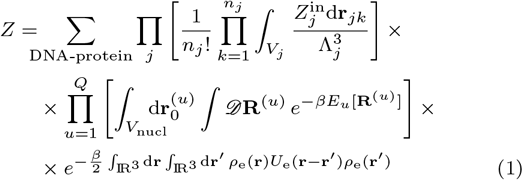

Here ∑_DNA-protein_ is the sum over all possible ways, in which nucleosomes can be positioned on chromosomal DNA. Π_*j*_ is the product over all types of particles (ions, proteins, cell metabolites, etc.) diffusing inside the cell, which are enumerated by index *j. n_j_* is the total number of particles of type *j* inside the cell. *p* is the product over all particles of the same type, *j*, whose positions in space are described by the radius vectors **r**_*jk*_. Integration *∫_V_j__* d**r**_*jk*_ in the above formula is performed over the volume *V_j_* accessible to particles of type *j*. In the case of ions, small proteins and cell metabolites, *V_j_* corresponds to the union of the cell nucleus and cytosol volumes: *V_j_* = *V*_nucl_ ∪ *V*_cyto_; whereas, in the case of macromolecular complexes that cannot move through NPCs into the nucleoplasm, we have *V_j_* = *V*_cyto_. Λ_*j*_ and 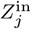 are thermal de Broglie wavelength and the partition function describing inner degrees of freedom of the *j*^th^ type of particles, respectively. 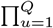, is the product over all chromosomes confined inside the nucleus, whose 3D conformations are represented by **R**^(*u*)^ functions. 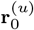 is the radius vector describing position of the starting end of the DNA polymer comprising the *u*^th^ chromosome. 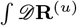 are Feynman-like path integrals along the contours of the respective DNA polymers (*u* = 1, …,*Q*), which are calculated over all possible chromosome conformations. *E_u_*[**R**^(*u*)^] is the energy of the DNA polymer comprising the *u*^th^ chromosome, which incorporates elastic deformations of the DNA as well as its interaction with DNA-binding proteins (histones), see details in Appendices A-B. In the general case, *E_u_* energy is determined by the 3D conformation of the *u*^th^ chromosome, which is described by **R**^(*u*)^(*s_u_*) function. *β* = 1/*k*_B_T is the inverse thermodynamic temperature, where *k*_B_ is the Boltzmann constant and *T* is temperature of the surrounding environment. *U*_e_ (**r**) = 1/4*πεr* is the core part of the electrostatic potential, where *r* = ||**r**|| is the length of the distance vector, **r**, between the electrically charged particles. *ε* is permittivity of intracellular media, which in the model was put equal to water permittivity. Finally, *ρ*_e_(**r**) is the distribution of electrical charges inside the cell, which is described by Eq. (A4) (Appendix A). More details regarding the above mathematical expression as well as treatment of the volume-exclusion effect can be found in Appendices A and L.

To calculate the partition function given by Eq. (1), in this study we employed the mean field approach by considering randomly fluctuating electrostatic field, *ψ*, created by electrically charged ions, metabolites and DNA, based on which it can be shown that Eq. (1) can be reduced to the following simple formula (see Appendix A for details):

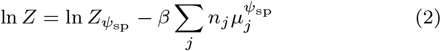

Where *Z*_*ψ*sp_ and 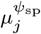 are the partition functions of DNA and electrochemical potential of the *j*^th^ type of particles in the presence of the stationary phase electrostatic field, *ψ*_sp_, which solves the following functional equation (see Appendix A):

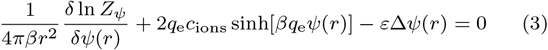

Here *r* is the radial distance measured from the center of the cell nucleus. *q*_e_ = 1.6 · 10^-19^ C is the elementary charge. *Z_ψ_* is the partition function of DNA interacting with histone octamers in the presence of electrostatic field *ψ*, see Appendices B-D. 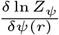 is the functional derivative of the DNA partition function, *Z_ψ_*, with respect to the electrostatic field, *ψ*, see Appendix E.

To solve Eq. (3), electrostatic field *ψ*(*r*) was represented in our study in a form of the following Fourier-Bessel expansion series:

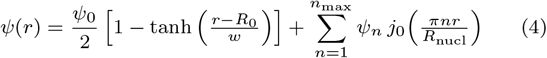

Where *R*_nucl_ is the radius of the cell nucleus. *n*_max_ is a number of Fourier-Bessel modes used in calculations (in this study, nmax = 18). As for *R*_0_,*w,ψ*_0_,*ψ*_1_,…,*ψn*_max_, these are the model fitting parameters, whose values were determined by Nelder-Mead simplex algorithm [73], which was used to minimize deviation of the left side of Eq. (3) from zero, see Appendix A for details.

Finally, to calculate the partition function of chromosomal DNA, *Z_ψ_*, in an arbitrary electrostatic potential field, *ψ*, we used a discretized semiflexible polymer chain model in which DNA is partitioned into short straight segments, whose size (*b* = 3.4 nm) is much smaller than the bending persistence length of DNA (*A* = 50 nm [74, 75]), see schematic Figure 1(c). Since the elastic bending and twisting energies of DNA can be described based on relative orientations of neighbouring DNA segments, it is then possible to utilize transfer-matrix calculations to obtain the partition function of DNA, see Appendices B-E. As a result, it can be shown that up to a non-essential prefactor, *Z_ψ_* can be represented in the following form:

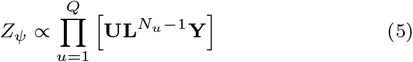

Where *N_u_* is the total number of segments in a discretized polygonal chain representing the DNA polymer comprising the *u*^th^ chromosome. **U** and **Y** are boundary condition vectors that depict physical states of the DNA end segments in each chromosome. **L** is a transfer-matrix, which characterizes DNA segments’ interactions with DNA-binding proteins and surrounding electrostatic potential field, *ψ*, as well as describes local bending and twisting elasticities of DNA.

The main advantage of Eq. (5) is that the value of each matrix product, **UL**^*N_u_*-1^**Y**, in this equation is mainly determined by the dominant eigenvalue (λ_max_) of the transfer-matrix **L** when *N_u_* ≫ 1, which is the case for long DNA molecules found in eukaryotic cells. Specifically, it can be shown that 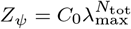, where 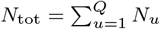 is the total number of DNA segments in all of the chromosomes, and *C*_0_ is a constant independent of *N*_tot_. Since both Co and the dominant eigenvalue *λ*_max_ can be found with the help of standard techniques, such as the power iteration method [76] (also see Appendices E and I), this makes it possible to calculate the partition function of an arbitrarily long DNA in the presence of DNA-binding proteins in solution and electrostatic interactions between all of the system components – a task, which cannot be handled by any of the existing theoretical methods employed in previous theoretical studies of DNA organization in nuclei of living cells. Furthermore, the above approach provides direct connection between the mesoscale structure of the entire cell genome and properties of individual protein complexes, which could not be achieved in prior studies.

In addition, it should be noted that, in contrast to previous theoretical works [33–46], in our model the chromatin structure is not fixed and nucleosomes are allowed to form / disassemble anywhere on DNA, enabling evaluation of the grand partition function of DNA over all possible DNA-protein conformations, which so far has been considered as a problem that cannot be solved for genomesize DNA by using modern computational systems.

By taking derivatives of the partition function logarithm defined by Eq. (2) with respect to various model parameters, such as the volume of the cell nucleus or histones binding energy to DNA, etc., it is possible to find the mean pressure created by DNA on the NE, the average DNA conformation and occupancy by nucleosomes, DNA distribution inside the nucleus, and many other quantities characterizing the physical state of chromosomal DNA and the cell nucleus. The full list of these quantities and formulas that were applied to calculate them can be found at the end of Appendix I. The rest of the model parameters as well as their values used in the calculations are listed in supplementary Table I.

The source code of the computer programs can be downloaded from http://www.artem-efremov.org website.

Finally, it should be noted that while here we have mainly focused on the model approach to study DNA organization in nuclei of living cells, the same method, with minor modifications, can be also employed to gain insights into DNA packaging in viral particles, see *Results* and Appendix A.

## RESULTS

### Viral DNA packaging

To test the developed theoretical approach, we first applied it to the description of DNA packaging in a viral particle in the presence of electrostatic DNA-DNA and ion-DNA interactions. By varying the size of the viral capsid, it has been found that the free energy of a DNA-loaded viral particle, *G*, changes almost as an inverse cubic function of the capsid radius (i.e., 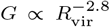), deviating from this trend only when the particle size reaches the value comparable to the DNA bending persistence length (~ 50 nm [74, 75]). At this point the slope of the free energy curve becomes flatter, which may seem to be counter-intuitive as one expects that at this point DNA will start to apply more pressure on the capsid wall due to accumulation of the DNA elastic bending energy, see Figure 2(a). However, additional calculations performed for a polymer with zero bending and twisting elasticities showed nearly the same results [see dashed curves in Figure 2(a)], suggesting that the elastic bending and twisting energies of DNA make rather negligible contribution into the total free energy of a viral particle.

**FIG. 2.**
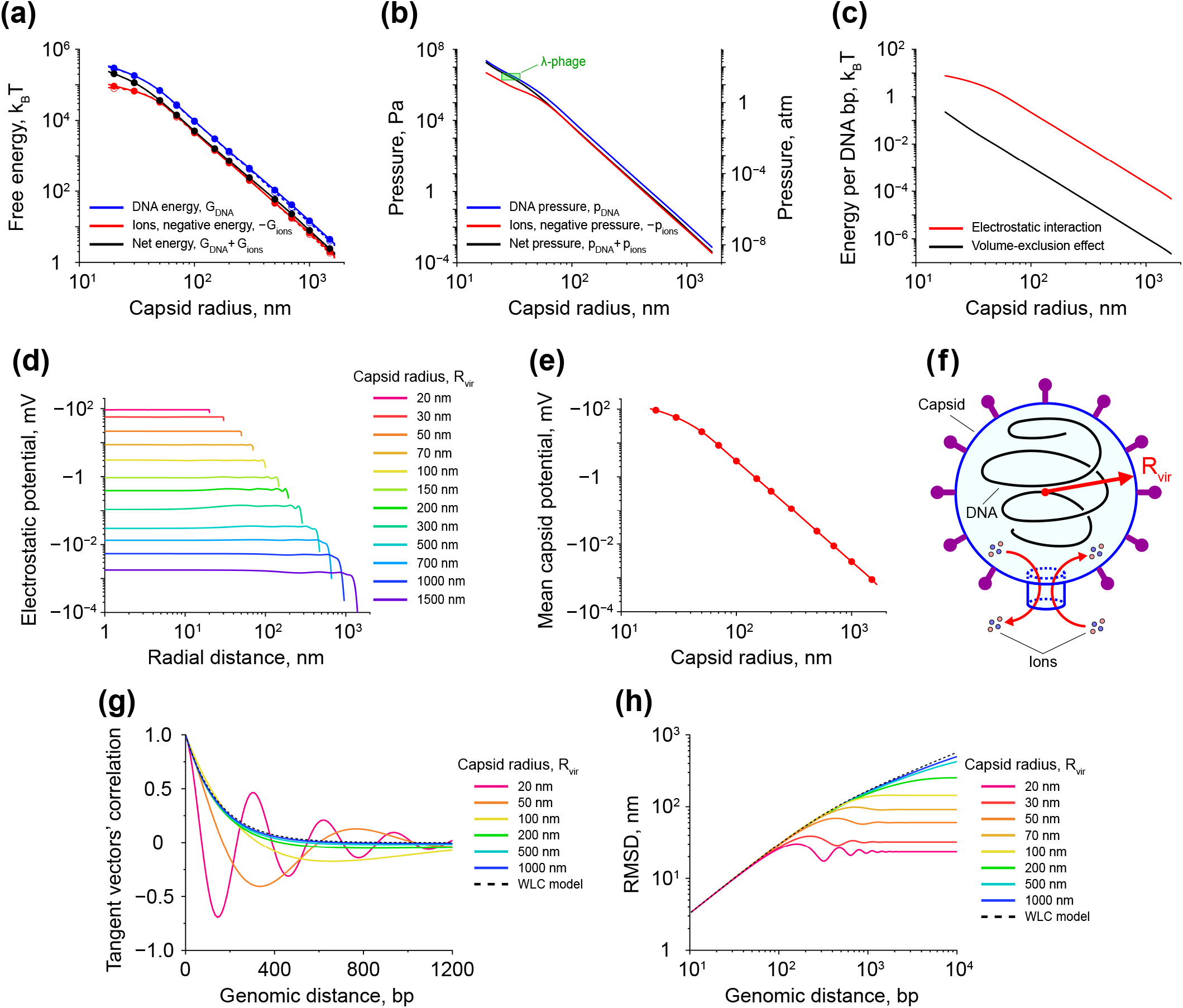
DNA packaging in viral particles. **(a)** Contributions made by DNA (blue curve) and ions (red curve) into the total free energy of a viral particle (black curve) as functions of the capsid size. Data points shown in the graph are the results of transfer-matrix calculations; whereas, lines indicate smoothing spline interpolation. Solid curves correspond to the case of a semiflexible DNA with the bending and twisting persistence lengths of *A* = 50 nm and *C* = 95 nm, respectively; whereas, dashed curves demonstrate results obtained for a freely-joint polymer chain of the same length but with zero bending and twisting elasticities between the polymer segments. All energy values in the graph are shown up to a non-essential additive constant. **(b)** Pressures generated by DNA and ions on the walls of a viral particle as functions of the capsid size. Green rectangle indicates the area corresponding to experimentally measured values for λ-phage particles [78, 79]. **(c)** Electrostatic and volume-exclusion DNA interaction energies per single DNA base-pair as functions of the capsid size. **(d)** Distribution of the electrostatic potential in viral particles of different sizes. **(e)** Volume-averaged value of the electrostatic potential in viral particles of different sizes. Each data point shown in the panel represents the corresponding curve displayed in panel (d). Solid line demonstrates interpolation of the data points by a smoothing spline. **(f)** Schematic picture of DNA packaging in a viral particle. Results shown in panels (g-h) suggest that DNA assumes a coil-like conformation inside the capsid. **(g)** and **(h)** The average correlation function between a couple unit vectors tangent to the DNA backbone and the root mean squared distance (RMSD) between two points residing on DNA as functions of the genomic distance between the vectors and points, respectively. Mathematical details regarding the formulas used in the above calculations can be found in Appendices I-J, L. In all computations, the DNA length was 14 *μ*m, corresponding to the size of EMBL3 λ-phage genome of 41.5 kbp [78], with the size of DNA segments in the discretized polymer model being equal to *b* = 3.4 nm.

Indeed, by estimating other energy terms, such as electrostatic DNA-DNA and ion-DNA interactions as well as DNA-DNA volume exclusion effect, it has been found that the vast majority of the viral particle free energy comes from electrostatic interactions between the system components, which are by ~ 1 — 2 orders of magnitude stronger than the DNA-DNA volume-exclusion effect, see Figure 2(c) and Appendix L. This result is in good agreement with previous experimental studies showing that DNA-DNA electrostatic interactions make the dominant contribution to the viral particle energy [77].

Calculations further demonstrated that the mean electrostatic field inside viral particles is rather uniform. At the same time the absolute value of the electrostatic field appeared to rapidly increase with decreasing size of the viral capsid, *R*_vir_, see Figures 2(d-e). At *R*_vir_ = 50 nm the electrostatic potential was found to reach the value of ~ −20 mV, at which the energy associated with movement of monovalent ions between the inner space of the capsid and outside environment becomes comparable to the thermal energy (1 *k*_B_*T* ≈ 27 e·mV). Beyond this point positively charged ions start to accumulate inside the capsid, enhancing electrostatic screening effect of DNA-DNA interactions. As a result, the system free energy grows slower at smaller values of the capsid size, explaining unusual flattening of the energy curves at *R*_vir_ ≤ 50 nm shown in Figure 2(a).

To compare the above results to experimental data, we next calculated derivatives of the ions and DNA free energy terms with respect to the capsid volume to estimate pressures generated by these components on the wall of the viral capsid, see Figure 2(b). From the figure it can be seen that overall behaviour of the pressure curves seems to be similar to that of energy curves displayed in Figure 2(a), with the only difference being much steeper slopes of the pressure curves.

Previous experimental studies indicate that the amount of pressure applied by viral DNA to the capsid wall can potentially reach a very high value of ~ 10 — 40 atm [78, 79] – an interval, which is marked by the green rectangle in Figure 2(b), that in addition shows the range of λ-phage particle sizes. By looking at Figure 2(b), it can be seen that the theoretically predicted DNA and net pressure curves pass exactly through the center of the experimentally measured range, suggesting that our model provides accurate description of the physical properties of packed viral DNA.

To further test the model, we calculated the correlation between unit vectors tangent to the DNA backbone as well as the root mean squared distance (RMSD) between two points residing on DNA as functions of the genomic distance along the DNA, see Figures 2(g-h). It has been found that the DNA conformation becomes more and more ordered with decreasing size of the viral capsid, and starting from *R*_vir_ ≈ 50 nm the correlation function takes the form of damped oscillations [see Figure 2(g)], indicating that DNA becomes folded into a coil-like structure, as illustrated in schematic Figure 2(f). This result is in good agreement with existing experimental data [80, 81] and theoretical simulations [82–84]. RMSD plot shown in Figure 2(h) further confirms this observation, suggesting that our model provides rather accurate description of DNA packaging in viral particles.

Interestingly, RMSD graphs shown in Figure 2(h) look very similar to those reported in a recent study of polymer molecules confined in 2D spaces [85], indicating strong similarity between polymers’ behaviour in 2D and 3D cases.

Altogether, the above results demonstrate that the developed model correctly describes behaviour of DNA confined in a tight space, making it suitable for study of DNA packaging in viral particles and nuclei of living cells. In this work, we used this approach to gain understanding of molecular mechanisms responsible for regulation of the cell nucleus size and its potential downstream effects on the chromatin structure, such as changes in the strength of DNA-protein interactions.

### Mechanical equilibrium of the cell nucleus

In the general case, the nucleus size is determined by mechanical equilibrium corresponding to a net zero pressure acting on the NE. This includes the pressure created by chromosomal DNA (*p*_DNA_) as well as the osmotic pressure resulting from the gradients of electrically charged ions and metabolites across the NE (*p*_ions_), which develop due to the mean negative electrostatic potential of the cell nucleus with respect to the cytoplasm, see schematic Figure 3(e). Cytosolic macromolecular compounds that cannot move through NPCs into the nucleus, have also been previously shown to be involved in generation of outside osmotic pressure on the NE, *p*_macro_ [86, 87]. In addition, it has been suggested that lamin proteins, which polymerize into a dense lamina network underlying the NE [88, 89], may be responsible for the nucleus growth during the cell cycle in metazoan cells [90, 91]. On the other hand, proteins binding to the ER membrane have been found to slow down this process [92]. As a result, it has been hypothesized that the NE and ER may be involved in a tug-of-war for the shared lipid membrane [1], which from a physical point of view can be described in terms of opposite pressures, *p*_NE_ and *p*_ER_, acting on the NE created by the surface tensions of the NE and ER membrane, *σ*_NE_ and *σ*_ER_, see Appendix K for details.

**FIG. 3.**
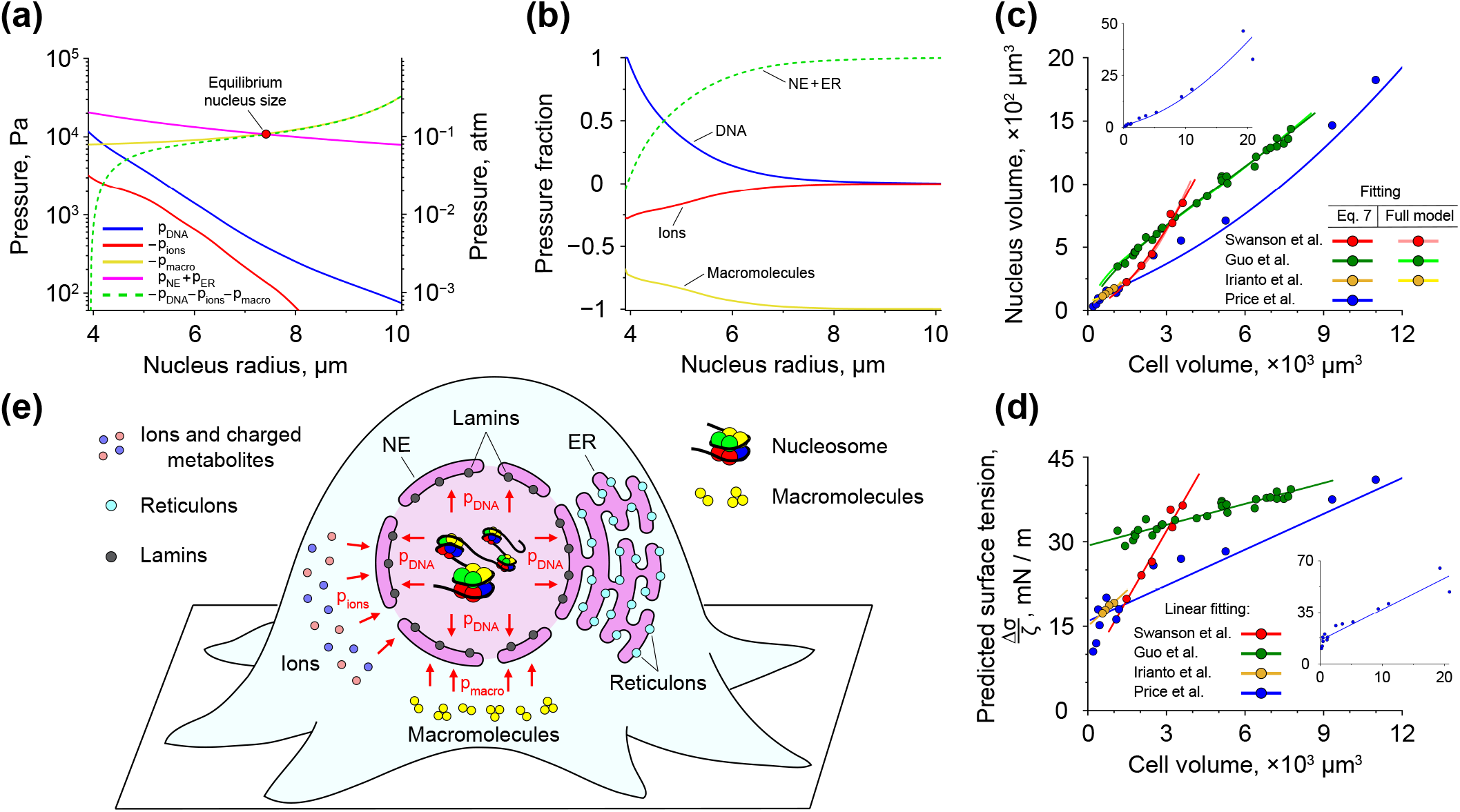
Cell nucleus size regulation. **(a)** Pressures exerted on the NE by DNA, ions, macromolecules, and the surface tensions of the NE and ER membrane. The yellow curve representing the osmotic pressure created by macromolecules is plotted for the case of *n*_macro_ = 10^10^ molecules. The magenta curve demonstrates the net pressure *p*_NE_ + *p*_ER_ generated by the difference in the NE and ER membrane surface tension, which corresponds to Δ*σ* = 20 mN/m; whereas, the green dashed curve indicates the equilibrium value of such a pressure, *p*_NE_ + *p*_ER_ = – *p*_DNA_ – *p*_ions_ – *p*_macro_ [see Eq. (6)], as a function of the cell nucleus radius. In transfer-matrix calculations, the total length of DNA was set equal to 2.1 m, corresponding to the size of human genome of *~* 6.2 Gbp [47]. The length of bare DNA segments in the discretized polymer model was *b* = 3.4 nm. **(b)** Relative contributions of different cell components to the total positive / negative pressure acting on the NE. Results shown in panels (a) and (b) were obtained for a cell with the total volume of *V*_cell_ = 8000 *μ*m^3^. Smaller size cells (*V*_cell_ = 4000 *μ*m^3^) show very similar behaviour, see Figure S1. **(c)** Correlation between the nucleus and cell volumes. Solid curves indicate fitting of experimental data from ref. [8, 9, 11, 15] either to the full model described by Eq. (6) or to the reduced model described by Eq. (7). In the former case, the fitting was done by varying a single model parameter – the difference between the surface tensions of the NE and ER membrane, Δ*σ*; whereas in the latter case, the linear fitting of Δ*σ/ζ* ratio shown in panel (d) was used as an input in Eq. (8). Inset displays a large scale view of the fitting of experimental data from ref. [8]. **(d)** *Δσ/ζ* ratio as a function of the nucleus size. Data points show values calculated by using Eq. (7) based on experimental data displayed in panel (c). Solid lines indicate data points fitting to a linear function. Inset demonstrates a large scale view of the fitting of data points obtained based on experimental data from ref. [8]. **(e)** Schematic picture of cellular components exerting pressure on the NE. The size of the cell nucleus in the general case is determined by mechanical equilibrium described by Eq. (6), which involves a tight interplay between the pressures generated by DNA (*p*_DNA_), ions (*p*_ions_) and macromolecules (*p*_macro_). In addition, it has been previously hypothesised that NE and ER, which share the same bilayer lipid membrane, may be involved in a tug-of-war with each other via indirect membrane-binding competition between lamins and reticulons [1] that can be described in terms of *p*_NE_ and *p*_ER_ pressures associated with the surface tensions of the NE and ER membrane.

Summing all the above pressures, it is easy to obtain the following formula for mechanical equilibrium of the NE:

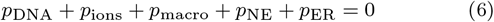

It should be noted that previously it has been proposed that the cell cytoskeleton may also exert mechanical forces on the nucleus, affecting its size and shape [93]. However, experimental measurements demonstrate that a typical mechanical stress developed by the actin network is generally small (~ 20 – 1000 Pa [11, 94]) in comparison to other pressures acting on the NE (see results below). Furthermore, it has been shown that disruption of the actin filaments by cytochalasin D does not influence osmolarity-induced change in the nucleus size [87]. Altogether, these results indicate that while the actin cytoskeleton may develop strong enough forces to affect the nucleus shape, it is unlikely to be involved in the nucleus size / volume regulation via directly generating pressure on the NE. Though, under certain conditions, it may have an indirect effect on the nucleus size by modulating the average level of lamin A/C and nucleocytoplasmic shuttling of histone-modifying enzymes [18, 93, 95, 96] as well as by causing changes in the cell shape / volume [11, 97], thus influencing *p*_NE_, *p*_DNA_, *p*_ions_ and *p*_macro_ terms in Eq. (6).

Anyway, by using formulas shown in Appendix K, we plotted the pressure terms from Eq. (6) as functions of the nucleus radius, *R*_nucl_, see Figure 3(a) and Figure S1(a). From the graphs it can be seen that the equilibrium size of the nucleus is predominantly determined by the difference in the surface tensions of the NE and ER as well as the osmotic pressure generated by cytosolic macromolecules. Indeed, from direct comparison of relative contributions of various cell components into the total positive and negative pressures acting on the NE [Figure 3(b) and Figure S1(b)] it can be concluded that neither DNA, nor ions make a substantial contribution in a wide range of the model parameters. Only in the case when the nucleus is compressed strong enough, the model predicts that DNA starts to produce sufficiently large repulsing force counteracting the applied pressure, which is in good agreement with existing experimental studies showing that DNA does not directly influence the nucleus size, rather setting its minimum possible value [10, 86, 98].

Hence, in the most cases, an approximate equilibrium radius of the nucleus, *R*_nucl_, can be found as a unique real-valued solution of the following cubic equation that can be derived from Eq. (6) (see Appendix K for details):

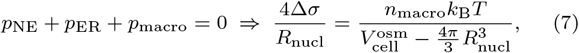

and thus

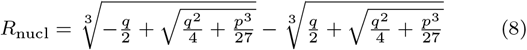

Where Δ*σ* = *σ*_ER_ – *σ*_NE_ is the difference between the surface tensions of the NE and ER membrane. 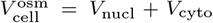 is the osmotically active volume of a cell, which typically occupies ~ 70% of the total cell volume 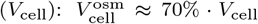 [11, 99]. nmacro is the total number of macromolecules in the cell cytosol. Taking into account that most of the macromolecules are either sufficiently large proteins or protein complexes that cannot move through NPCs, it is natural to expect that their number is approximately proportional to the total number of proteins in a living cell: *n*_macro_ ≈ *ζn*_pr_ = *ζc*_pr_ *V*_cell_. Here *ζ* is a proportionality coefficient (0 < *ζ* < 1), and *c*_pr_ is the average protein concentration in living cells: *c*_pr_ = 2.7 · 10^6^ proteins/*μ*m^3^ [100]. Finally, *q* and *p* coefficients in Eq. (8) are: 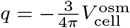 and 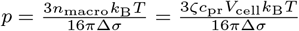.

We then used Eq. (7) to estimate Δ*σ/ζ* ratio based on experimentally measured volumes of different types of cells and their nuclei [8, 9, 11], see Figure 3(d). From the figure it can be seen that Δ*σ/ζ* ratio predicted by the model changes almost linearly with the cell volume in all considered cases. As a result, by fitting each of the data set shown in Figure 3(d) to a linear function and then substituting this function into Eq. (8), a nearly perfect fit of the experimentally measured correlations between the cell and nucleus volumes could be achieved, see Figure 3(c). Fitting of the experimental data to the full model described by Eq. (6) led to very similar results [Figure 3(c)], indicating robustness of Eq. (7)-Eq. (8).

Interestingly, the calculated values of Δ*σ/ζ* ratio appeared to be of the order of ~ 20 −40 mN/m [Figure 3(d)], which is close to the elastic modulus of the NE measured in isolated nuclei of *Xenopus laevis* oocytes (28 ± 8 mN/m [86]). This result suggests that the nuclear lamina network likely plays a major role in governing the nucleus size regulation in *Xenopus* oocytes, – a model prediction, which is in good agreement with previous experimental studies [90, 91].

By using Eq. (6) and 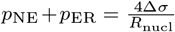 formula (Appendix K), we have also estimated the difference between the surface tensions of the NE and ER membrane, Δ*σ*, that has to be maintained by a living cell in order to retain the nucleus size at a specific value in the case of *ζ* = 0.4 - 0.5, see Figure S1(c). It can be seen from the figure that bigger nuclei require considerably larger Δ*σ*, which corresponds to a high surface free energy density of the order of ~ 1 – 10 *k*_B_*T*/nm^2^ (~ 4 – 40 mN/m). This observation further stresses importance of the nuclear lamina network as a potential power source driving the cell nucleus growth both throughout the cell cycle [90, 91] and during the cell differentiation process [101].

Interestingly, the value of Δ*σ* predicted by the model seems to be slightly higher than the rupture surface tension of the plasma membrane in living cells, which was previously found to be of the order of several mN/m [102, 103]. However, it should be noted that since numeric estimations based on *in vitro* observations of force-induced membrane tether formation suggest that the surface tension of ER is likely to be very small (|*σ*_ER_| < 0.1 mN/m [104]), the large positive value of Δ*σ* implies a high negative membrane tension of the NE. I.e., the model predicts that the nuclear envelope is strongly compressed in living cells rather than stretched as in membrane rupture experiments [102, 103]. In this case, most of the external pressure exerted by cytosolic macromolecules on the NE will be borne by the lamina network underlying the nuclear envelope, which may explain how the nuclear envelope maintains its integrity despite a strong pressure applied to it.

### Effect of the nucleus size on chromatin

Since existing experimental studies suggest that electrostatic interactions play a major role in DNA organization, we next checked how the size of the cell nucleus affects its mean electrostatic potential with respect to the cytoplasm. From the results shown in Figures 4(a-b) it can be seen that the absolute magnitude of this potential experiences a rapid increase with the decreasing size of the nucleus, reaching the value of several millivolts, which is within the experimentally measured range of the nuclear potential in *Xenopus* oocytes, MDCK cells and isolated murine pronuclei (~ from –10 mV to 0 mV [57, 105, 106]).

**FIG. 4.**
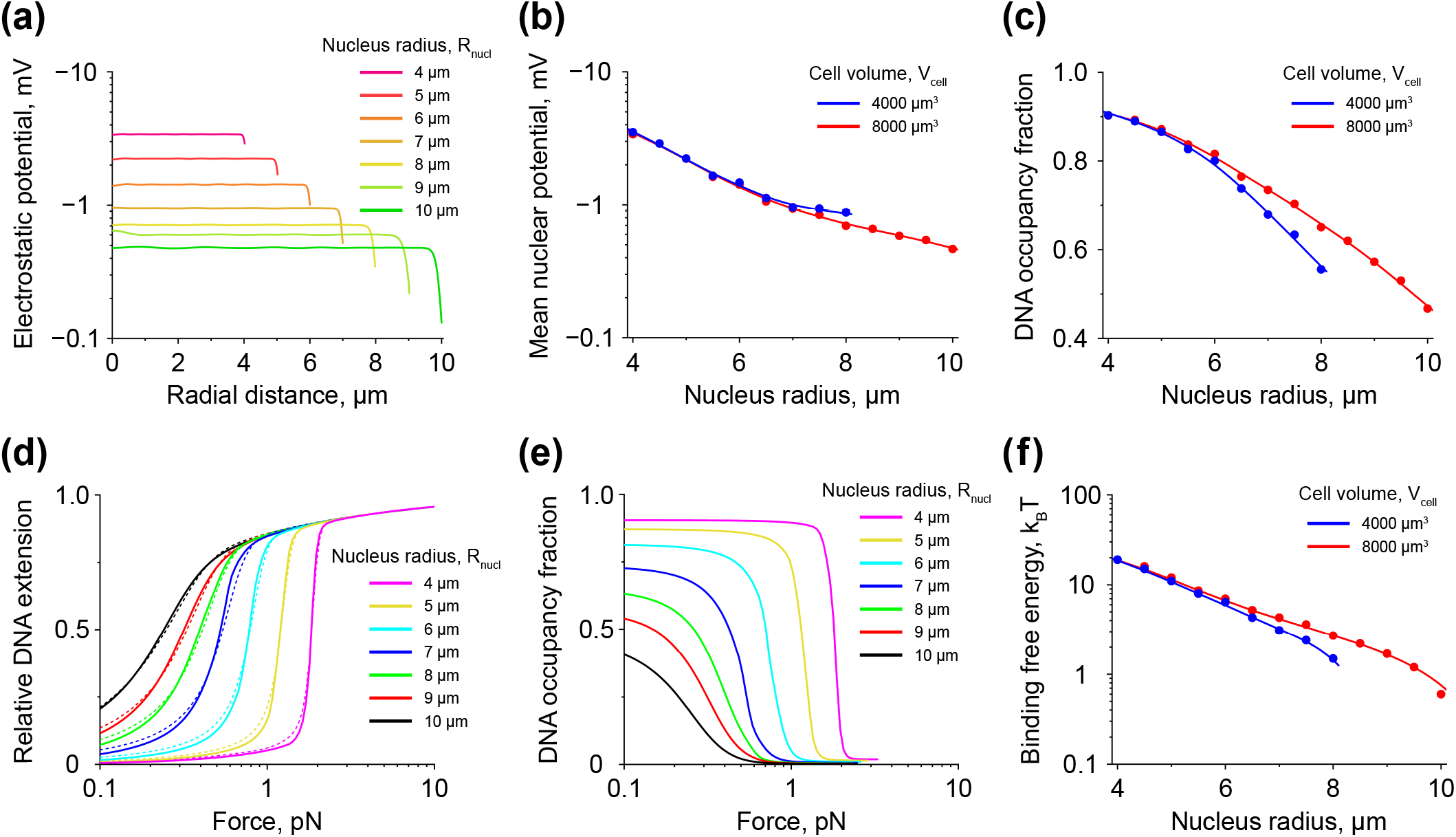
Nuclear electrostatic potential, and correlation between the nucleus size and nucleosome stability. **(a)** Distribution of the electrostatic potential in nuclei of different sizes. The plot shows results obtained for a cell with the total volume of *V*_cell_ = 8000 *μ*m^3^. Smaller size cells (*V*_cell_ = 4000 *μ*m^3^) demonstrate a very similar nuclear electrostatic potential distribution, see Figure S2. **(b)** Volume-averaged value of the electrostatic potential in nuclei of different sizes. Data points indicate values calculated based on the curves shown in panel (a) and Figure S2; whereas, solid lines demonstrate smoothing spline interpolation. **(c)** Occupancy fraction of chromosomal DNA by nucleosomes as a function of the cell nucleus size. Data points represent results of transfer-matrix calculations; whereas, solid lines show smoothing spline interpolation. **(d)** and **(e)** Force-extension and force-DNA occupancy fraction curves obtained by mechanical stretching of a small part of chromosomal DNA (2 *μ*m in length) in nuclei of different sizes. Solid curves on panels (d) and (e) demonstrate results of transfer-matrix calculations performed for a cell with the total volume of *V*_cell_ = 8000 *μ*m^3^ (see also Figure S3 for *V*_cell_ = 4000 *μ*m^3^ case). Dashed curves in panel (d) display results of the force-extension curves’ fitting to the previously published model of a mechanically stretched DNA [52]. **(f)** Binding free energy of histone octamers to DNA as a function of the cell nucleus size. Data points shown in the graph are calculated based on the force-extension curves’ fitting displayed in panel (d) and Figure S3(a). Solid curves indicate prediction of the binding free energy of histone octamers by Eq. (9).

As histone dimers possess a strong electrical charge (~ +37.2 *q*_e_, where *q*_e_ = 1.6·10^-19^ C), it is clear that such a nuclear electrostatic potential may have a profound effect on stability of nucleosomes formed on the chromosomal DNA. Indeed, the model calculations demonstrate that the DNA occupancy by nucleosomes can experience considerable variation in response to changes in the nucleus size [Figure 4(c)], suggesting strong dependence of the histone binding free energy to DNA on the nuclear electrostatic potential, – a result, which is supported by *in vitro* experimental studies demonstrating importance of electrostatic DNA-protein interactions and surrounding ionic environment in regulation of nucleosome spacing on DNA [27].

To investigate the role of the nuclear electrostatic potential in stabilization of nucleosomes, we performed calculations of the average pulling force needed to be applied to the chromosomal DNA in order to induce mechanical unfolding of nucleosomes, – a quantitative indicator of the stability of nucleoprotein complexes, which is frequently used in single-molecule studies [107, 108]. From the results shown in Figures 4(d-e) it can be seen that in the case of small nuclei one needs to apply a pulling force, which is several times larger than in the case of bigger nuclei to initiate mechanical unfolding of nucleosomes, indicating nucleosome stabilization effect by the nuclear electrostatic potential.

By fitting the force-extension curves shown in Figure 4(d) to our previously model of a mechanically stretched DNA [52], we estimated the average binding free energy of histone octamers to DNA in cell nuclei of different sizes, see Figure 4(f). It has been found that the nucleus size indeed has a strong effect on the binding free energy of histone octamers to DNA, causing changes of the order of ~ 1 – 20 *k*_B_*T* in response to the nucleus size variations.

In fact, it is not hard to obtain the following approximate formula for the total binding free energy of histone octamers to DNA, 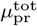, which is equal to the difference between the electrochemical potentials of the initial and final states of the protein complexes contributing to nucleosome assembly (for more details see Appendix B):

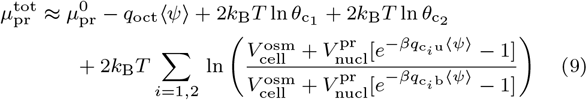

Here 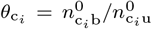 (*i* = 1, 2) are occupancy ratios of histone-binding chaperones, where 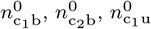 and 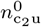 are the total average numbers of c1 and c_2_ chaperones in histone-bound and unloaded states, respectively, see Appendix A. *q*_oct_ is the electrical charge of a histone octamer. *q*_c_*i*_b_ and *q*_c_*i*_u_ are the total electrical charges of the *i*^th^ chaperone, *c_i_*, in histone-bound and unloaded states, respectively. 〈*ψ*〉 is the mean value of the nuclear electrostatic potential obtained by averaging over the nucleus volume. 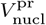 is a free nucleus volume accessible to proteins, such as histone-binding chaperones, which, for the sake of simplicity, was set equal in this study to the total nucleus volume, 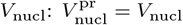, see notes at the end of Appendix M. Finally,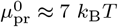 is the standard Gibbs free energy of nucleosome formation in the presence of histone-binding chaperones under the standard experimental conditions: *θ*_*c*_1__ = *θ*_*c*_2__ = 1 and *ψ* = 0 [62, 63].

As can be seen from comparison of the solid curves in Figure 4(f) plotted by using Eq. (9) and the data points obtained by fitting of the force-extension curves shown in Figure 4(d), the above formula accurately predicts behaviour of the binding free energy of histone octamers to DNA as a function of the nucleus size.

Interestingly, from Eq. (9) it can be seen that not only the electrical charges of histones are important for stabilization of nucleosomes by the nuclear electrostatic potential, but also electrical charges of histone-binding chaperones responsible for their transportation. Indeed, simple model calculations based on Eq. (9) show that depending on the electrical charge of histone-binding chaperones, a living cell can switch between two regimes: 1) nucleus size-sensing, in which the binding free energy of histone octamers to DNA depends on the nucleus size, and 2) nucleus-size insensitive, in which the binding free energy of histone octamers to DNA becomes nearly independent from the nucleus size, indicating existence of a previously unknown potential regulatory pathway of nucleosome stability.

Altogether, the above results suggest that contingent on physical properties of histone-binding chaperones, living cells may have very diverse response to changes in the nucleus size in terms of chromatin structure, whose reorganization can be driven by variation in the average strength of the nuclear electrostatic potential.

## DISCUSSION

In this study, we have developed a general theoretical framework aimed at description of DNA packaging in viral particles and nuclei of living cells, which allowed us to address the question of nuclear organization by taking into account main physical forces contributing to it. As a result, it has been found that the nucleus size in eukaryotic cells is predominantly determined by a tug-of-war between the osmotic pressure exerted by cytosolic macromolecules on the NE and the difference between the surface tensions of the NE and ER membrane, see Figure 3 and Figure S1. This finding is in good agreement with previous experimental studies showing that cytosolic macromolecules and lamins play a central role in the nucleus size regulation in metazoan cells [3, 86, 87, 90, 91, 109, 110].

It should be noted that although there may exist other molecular mechanisms which might contribute to the nucleus size regulation, it seems that the tug-of-war between cytosolic macromolecules, NE and ER will likely play the central role in this process. For example, it is known that yeasts do not have lamin analogues, and thus their nuclei do not possess a lamina network. Because of that, these cells have to rely on alternative mechanisms to regulate volumes of their nuclei, for instance, through accumulation of nuclear proteins, which may be used by cells to build up internal osmotic pressure on the NE counterbalancing the outside pressure created by cytosolic macromolecules [4]. In addition, despite the lack of lamins, yeasts may create a difference in the surface tensions of the NE and ER membrane, Δ*σ*, by maintaining a slightly distinct lipid composition of the NE and ER membrane [111, 112]. In both cases, the equilibrium nucleus size still will be described by the same Eq. (6)-Eq. (7), in which an additional term corresponding to the osmotic pressure created by a mobile fraction of macromolecules confined inside the cell nucleus, *p*_nucl_ = –*n*_nucl_*k*B*T*/*V*_nucl_, has to be introduced:

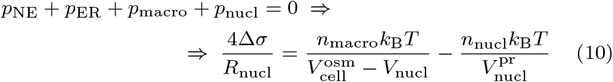

Here *n*_nucl_ is the total number of mobile macromolecules confined in the cell nucleus. 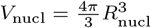 is the nucleus volume and 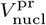 is a free nucleus volume accessible to proteins 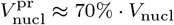 [31]).

From the above formula, it can be seen that in the absence of the surface tension (Δ*σ* = 0), the ratio between the nucleus and cell volumes is almost equal to *n*_nucl_/*n*_macro_ ratio of macromolecular content of the nucleoplasm and cytoplasm, which, as has been suggested in a recent theoretical study [113], plays an important role in determining the size of the nucleus in yeast cells that lack lamina network. This prediction is in good agreement with experimental measurements, which show that in yeast the nucleus volume changes in proportion to the cell volume. However, in the case of higher eukaryotes, the nucleus volume does not seem to be strictly proportional to the cell volume [Figure 3(c)], indicating a potential role of the membrane tension of the NE in regulation of the nucleus size.

Previously, it has been proposed that the actin cytoskeleton may also contribute to regulation of the nucleus size in mammalian cells via application of mechanical forces to the NE [97]. However, it should be noted that independent experimental studies show that the actin network typically generates rather small mechanical stress (~ 20 — 1000 Pa [11, 94]) in comparison to pressures developed by other cell components on the NE (see Figure 3). Thus, while mechanical forces created by the actin cytoskeleton may be strong enough to perturb the nucleus shape, it seems unlikely that they may cause considerable changes in the nucleus volume, leading instead only to small fluctuations of the nucleus size of the order of tens to hundreds of nanometers, which indeed have been observed in recent experimental studies [114, 115].

On the other hand, the role of the actin cytoskeleton in regulation of the cell shape / volume as well as the average level of lamin A/C and nucleocytoplasmic shuttling of histone-modifying enzymes has been documented in several previous studies [11, 18, 93, 95–97], suggesting that it may have rather an indirect effect on the nucleus size by modulating *p*_NE_, *p*_DNA_, *p*_ions_ and *p*_macro_ terms in Eq. (6).

Interestingly, existing experimental data indicate that abnormal expression of lamins as well as their defective localization frequently result in aberrant nuclear sizes and anomalies in the nuclear-to-cell volume ratio (i.e., N/C ratio), both of which strongly correlate with development of human-related diseases, such as certain types of cancers [116–118]. Based on this and other observations, it has been previously suggested that there may exist a unique connection between the nucleus size, the condensation state of chromatin and the transcription level of various genes, leading to hypothesis of the nucleus size-dependent regulation of gene transcription [15, 18, 26, 29, 98, 119]. However, possible molecular mechanisms that might be involved in such regulation remained unclear.

To gain insights into the above problem, we used the developed theoretical approach to predict a potential effect of the nucleus size on chromatin structure. As a result, it has been found that by influencing the average density of negatively charged DNA, the nucleus size determines the mean electrostatic potential of the cell nucleus with respect to the cytoplasm, which in turn defines stability of nucleosomes as well as the average DNA occupancy by them, see Figures 4, S2 and S3. Since in a recent experimental study it has been shown that stability of nucleosomes bound to promoter regions has a strong impact on the transcriptional level of the downstream genes [120], these results point to a possibility that model-predicted changes in nucleosome stability and the average DNA occupancy level, which are caused by the nucleus size variations, may have a profound effect on the global gene expression profile.

Indeed, in ref. [98, 119], it has been found that the nucleus size and N/C ratio strongly correlate with the total amount of RNA molecules in living cells as well as their synthesis rate by RNA polymerase II. This observation is further supported by a recent experimental study in which it has been demonstrated that application of external mechanical pressure on NIH 3T3 cells leads to decrease in the nucleus volume accompanied by condensation of chromatin and subsequent decrease in transcriptional activity of RNA polymerase II [18]. And while it has been suggested that such downregulation of RNA polymerase II takes place due to shuttling of MRTF-A transcription cofactor to the cell cytoplasm [18], simple estimations based on our model indicate that this phenomenon may actually have a deeper physical origin.

Namely, based on the results presented in Figure 4(f), it can be shown that pressure-induced changes in the nucleus volume measured in ref. [18] lead to ~ 4 – 5 *k*_B_*T* increase in the binding free energy of histone octamers to DNA. As suggested by our recent theoretical study [66], this amount of free energy may be enough to tip the balance in DNA-binding competition between transcription factors and histone octamers in favour of nucleosome formation, resulting in release of transcription factors from the chromosomal DNA and global downregulation of gene transcription. Thus, our model predicts that the observed correlation between N/C ratio and RNA synthesis rate in living cells may be caused by physical mechanisms related to stabilization of nucleosomes in a nucleus size-dependent manner, providing new insights into this interesting and important phenomenon.

A very similar mechanism may also underlie recently reported nucleus size-dependent formation of paraspeckles, – nucleoprotein condensates in nuclei of living cells responsible for gene expression regulation [121]. Specifically, in ref. [121], it has been found that the average number of paraspeckles increases with the nucleus size in numerous types of human cells. Moreover, experimental data indicate that accessibility of DNA for binding of paraspeckle components is one of the main factors required for assembly and maintenance of paraspeckles and other nuclear long-noncoding RNA condensates in living cells [121]. Thus, the nucleus size-dependent stabilization of nucleosomes discovered in our study may have a drastic effect on paraspeckle assembly by modulating DNA-binding competition between the paraspeckle components and histone octamers, which could explain the observed correlation between the number of paraspeckles and the size of the cell nucleus.

It should be noted that the above increase of ~ 4 – 5 *k*_B_*T* in the binding free energy of histone octamers to DNA in response to pressure application to living cells is caused by a rather mild reduction of the cell nucleus volume by ~ 20 – 30% [18]. At the same time, it has been previously reported that nuclei of living cells experience significantly larger volume increase by ~ 100 – 200% (i.e., by 2 – 3 times) during G1 phase of the cell cycle [91, 122, 123], which, according to our model, leads to a profound drop in the binding free energy of histone octamers to DNA by ~ 8 – 11 *k*_B_*T*. Therefore, expansion of nuclei observed during G1 phase may have a considerable impact on the condensation state of chromatin and the average synthesis rate of RNA, which may help cells to faster produce and double all their cytoplasmic content in preparation for cell division.

Likewise, the nucleus size-dependent regulation of nucleosome stability may also play an important role during cell differentiation process. Indeed, as cells undergo differentiation, they need to synthesize a large amount of proteins to fulfil their new biological functions. Thus, one might expect that differentiated cells on average require more intensive production of multiple RNAs in comparison to stem cells, which, according to ref. [98, 119] and our study, can be achieved by increase in the nucleus size and/or N/C ratio. This hypothesis is supported by experimental findings showing that differentiation of human embryonic stem cells is indeed accompanied by expansion of their nuclei [101], which may be potentially caused by nuclear accumulation of lamins A/C, – a phenomenon that has been found to take place not only in differentiating human cells, but also in cells from other animal species [101, 124, 125].

Interestingly, among different proteins that constitute the nuclear lamina network, lamin A has the strongest effect on the nuclear lamina architecture and its elastic properties [48, 89]. Furthermore, it has been revealed that lamin A is expressed in living cells in a mechanosensitive way, enhancing matrix-directed stem cell differentiation [126]. This observation together with our findings rises an interesting possibility of a previously unknown pathway of gene transcription regulation. Namely, from the results published in ref. [25, 126], it follows that the cell shape and/or mechanical properties of the surrounding environment have a profound effect on the expression level of lamin A, which, in turn, has a strong influence on the elastic properties of the nuclear lamina network and therefore the surface tension of the NE [48]. The latter, according to our study, determines the nucleus size and, as a result, nucleosome stability, which may have a global impact on gene transcription profiles.

Existence of such a molecular link between expression of lamin A and the global gene transcription profile is supported by a recent study [17]. In this work, it has been demonstrated that incubation of mouse fibroblasts on micropatterned substrates causes nuclear reprogramming in the absence of any exogenous reprogramming factors, which is accompanied by downregulation of lamin A and changes in 3D chromatin organization, with a systematic progression of cells from mesenchymal to stem cell transcriptome.

Remarkably, our model suggests that such a nucleus size-dependent regulation may in fact be quite selective, involving only a certain set of genes instead of the whole cell genome. Specifically, the model shows that stabilization of nucleosomes in a nucleus size-dependent manner is strongly affected by electrical charges of histone-binding chaperones, see Figure 5. Thus, by using histone-binding chaperones or chaperone-related cofactors with different electrical charges, living cells may be able to have several alternative regulatory pathways, some of which might be nucleus size-sensing, while others not, making it possible for cells to precisely fine-tune stability of nucleosomes at specific locations on the chromosomal DNA in response to nucleus size variations. Such a biological function of histone-binding chaperones and their cofactors can be performed in cooperation with histone-modifying epigenetic factors [127], and/or importins and exportins, which regulate nucleocytoplasmic trafficking of proteins [128], including histone-binding chaperones [54, 129, 130] and lamins [109], see also Appendix M.

**FIG. 5.**
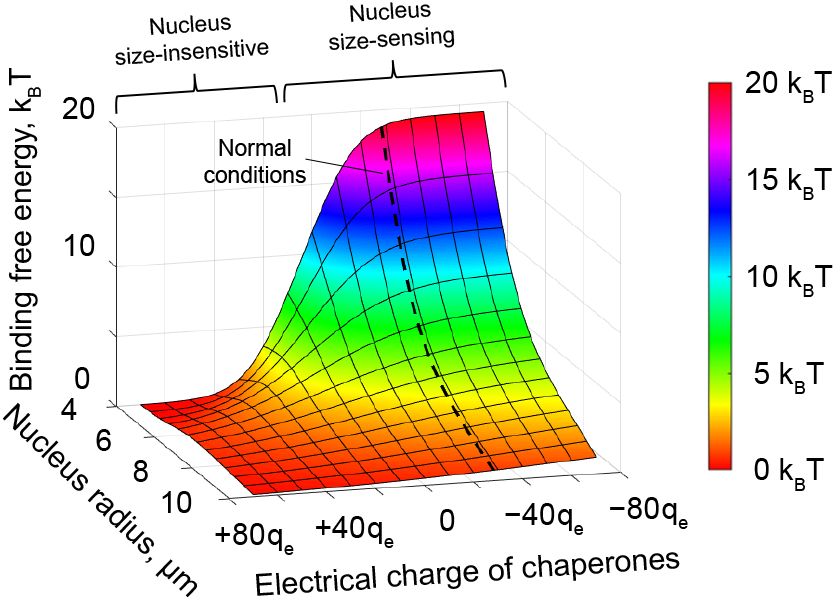
Role of the electrical charge of chaperones in regulation of the binding free energy of histone octamers to DNA. The plot demonstrates results of calculations based on Eq. (9) performed for different electrical charges of histone-binding chaperones and cell nucleus sizes. For the sake of simplicity, it was assumed in the calculations that H2A·H2B-binding chaperones as well as H3·H4-binding chaperones, both have similar electrical charges: *q*_c_1_u_ = *q*_c_2_u_. Dashed curve shown in the graph indicates model predictions based on estimation of the chaperone electrical charge (*q*_c_1_u_ = *q*_c_2_u_ = −28.2 *q*_e_, where *q*_e_ = 1.6 · 10^-19^ C is the elementary charge), which was obtained from experimental measurements of relative histone concentrations in the cytosol and nucleoplasm of HeLa cells [135], see Appendix A for details. It can be seen from the plot that the binding free energy of histone octamers to DNA may either change with the size of the cell nucleus, or remain insensitive to nucleus size variations, depending on the value of the electrical charge of histone-binding chaperones.

Furthermore, from Eq. (9) it follows that not only the nuclear electrostatic potential and electrical charges of histones and chaperones contribute to the binding free energy of histone octamers to DNA, but also the loading ratios of histone-binding chaperones, indicating that molecular pathways responsible for histone synthesis and degradation may as well affect nucleosome stability.

Presence of nuclear ion channels adds an extra level of regulation to the aforementioned pathways. Indeed, while some experimental studies suggest that NPCs and nuclear ion channels serve as passive elements allowing free diffusion of electrically charged molecules point to a possibility of a more specialized role of K^+^-selective channels in modulation of the nuclear electrostatic potential in certain types of living cells [106]. In the latter case, experimental measurements show that K^+^-selective channels contribute to the formation of a greater nuclear electrostatic potential (~ −10 mV) [106] in comparison to cells in which ions can freely move between the cell cytoplasm and nucleoplasm (~ −4 mV) [57]. As a result, stabilization of nucleosomes by the nuclear electrostatic potential will be stronger in cells expressing K^+^-selective NE channels, leading to more significant changes in chromatin organization.

Taken together, the above results indicate that living cells can utilize several possible pathways to fine-tune chromatin structure in response to environmental cues. However, it should be noted that all these pathways operate on top of a more general physical background created by the molecular processes discussed in our study, which make a major contribution into chromatin organization.

Finally, it should be noted that versatility of the theoretical approach developed in our work allows one to incorporate into the model practically all of the aforementioned mechanisms for a better understanding of their effect on chromatin, warranting future study. Furthermore, with a few minor modifications, the model can be also used to investigate the role of other types of interactions in shaping the chromatin landscape, such as HP1-dependent crosslinking of nucleosomes [131, 132] and formation of contacts between the nuclear lamina network and chromosomal DNA [133, 134]. This may provide valuable information about phase-separation processes taking place inside nuclei of living cells, such as self-organization of hetero- and euchromatin domains, making it possible to better comprehend their role in regulation of the cell response to various environmental cues.

## AUTHOR CONTRIBUTIONS

L.H. and A.K.E. made the computer program used for model calculations. Y.J. supported the study, providing useful suggestions. A.K.E. designed the research, derived formulas, carried out computations, analyzed the data and wrote the paper.

## ACKNOWLEDGEMENTS

We would like to thank Yuqi Guo from Shenzhen Bay Laboratory for providing images of living cells stained with ANG-2 sodium indicator. This work was supported by the National Research Foundation, Prime Minister’s Office, Singapore and the Ministry of Education under the Research Centres of Excellence programme (Y.J.) and the start-up funds from Shenzhen Bay Laboratory (A.K.E.).

## Notes

### Competing Interest Statement

The authors have declared no competing interest.

### Summary of Updates

New references, results, supplementary figures (S5, S6, S7, S15, S16) and appendix section (Appendix M) have been added to the manuscript and SI.

