## Supplement Information for "Nucleus size and its effect on the chromatin structure in living cells"

### SUPPLEMENTARY INFORMATION

#### Appendix A: General idea. Polymer field theory background.

In this study, we are interested in understanding the role of physical conditions, which are typically met inside viral particles and nuclei of eukaryotic cells, in regulation of DNA organization and DNA-protein interactions (in the case of living cells).

To this aim, we utilize previously developed transfer-matrix framework [1–3], further extending it by including interactions between the system components mentioned in *Introduction* section to describe crowded microenvironment existing inside a viral capsid or cell nucleus. Namely, it can be shown that electrostatic (or volume-exclusion) interactions between DNA segments and surrounding molecules can be accounted for in partition function calculations via a randomly fluctuating field, $\psi$, that describes the net electrostatic potential (or local average excluded volume) acting on each element of the system, which is generated by the rest of the molecules and DNA [4, 5].

To demonstrate this, let's recall that in the general case the conformation of a DNA polymer can be completely specified by a function, $\mathbf{R}(s)$, which describes 3D orientation of the DNA backbone versus the arc length, $s$, in terms of the Euler rotation matrix, $\mathbf{R}$ (see *Methods*). The latter shows how much the DNA backbone at the point corresponding to the arc length $s$ is rotated with respect to the global coordinate system, $(\mathbf{x}_0, \mathbf{y}_0, \mathbf{z}_0)$, see schematic Figure 1(b) and Figure S4(a). Specifically, the unit vector $\mathbf{z}(s) = \mathbf{R}(s)\mathbf{z}_0$ resulting from rotation of $\mathbf{z}_0$-axis of the global coordinate system via Euler matrix $\mathbf{R}(s)$ represents tangential direction of the DNA backbone at the point corresponding to the arc length, $s$. The remaining two unit vectors, $\mathbf{x}(s) = \mathbf{R}(s)\mathbf{x}_0$ and $\mathbf{y}(s) = \mathbf{R}(s)\mathbf{y}_0$, are normal to the DNA backbone and keep track of the DNA twist angle, $\varphi(s)$.

Having at hands $\mathbf{R}(s)$ function describing the DNA conformation, it is then straightforward to calculate the total energy of DNA, $E[\mathbf{R}]$, which is confined inside a spherical shell of radius $R_0$, such as a viral capsid or cell nucleus. In the simplest case, when electrostatic and volume-exclusion interactions between the system components are neglected, and DNA-binding proteins are absent in surrounding solution, $E[\mathbf{R}]$ comprises only the elastic bending and twisting energies of DNA as well as the DNA confinement term:

$$E[\mathbf{R}] = \frac{k_\mathrm{B}TA}{2}\int_0^L \left(\frac{\mathrm{d}\mathbf{z}(s)}{\mathrm{d}s}\right)^2 \mathrm{d}s + \frac{k_\mathrm{B}TC}{2}\int_0^L \left(\frac{\mathrm{d}\varphi(s)}{\mathrm{d}s}\right)^2 \mathrm{d}s + \frac{1}{L}\int_0^L U_{R_0}(\mathbf{r}_\mathrm{d}(s))\,\mathrm{d}s \tag{A1}$$

Where $k_\mathrm{B}$ is Boltzmann constant, $T$ is temperature of surrounding environment. $A$ and $C$ are bending and twisting persistence lengths of DNA. $\mathbf{r}_\mathrm{d}(s)$ is the position vector of a point on the DNA polymer corresponding to the arc length $s \in [0, L]$, where $L$ is the contour length of DNA. $U_{R_0}(\mathbf{r})$ is a spherical potential well describing DNA confinement inside the shell: $U_{R_0}(\mathbf{r}) = 0$ if $\mathbf{r}$ is inside the ball $B_{R_0}$ of radius $R_0$ (i.e., $\mathbf{r} \in B_{R_0}$), and $U_{R_0}(\mathbf{r}) = \infty$ otherwise (i.e., if $\mathbf{r} \notin B_{R_0}$). Dimensionality of $U_{R_0}$ is: $[U_{R_0}] = \mathrm{J}$.

By knowing the total energy of DNA, $E[\mathbf{R}]$, one then can find its partition function, $Z$, as:

$$Z = \int_{B_{R_0}} \mathrm{d}\mathbf{r}_0 \int e^{-\beta E[\mathbf{R}]} \mathscr{D}\mathbf{R} \tag{A2}$$

Where $\mathbf{r}_0$ is the position vector of the starting end of DNA, and $\int_{B_{R_0}} \mathrm{d}\mathbf{r}_0$ is the integral over all possible values of the position vector: $\mathbf{r}_0 \in B_{R_0}$. $\int \mathscr{D}\mathbf{R}$ is a Feynman-like path integral over all possible DNA conformations described by $\mathbf{R}(s)$ function. $\beta$ is the inverse of thermodynamic temperature: $\beta = 1/k_\mathrm{B}T$.

In the case of DNA interaction with DNA-binding proteins, the above Eq. (A1) has to be supplemented with several additional energy terms describing the binding energy of proteins to DNA and elastic deformations of DNA caused by formation of nucleoprotein complexes. Subsequently, a sum, $\sum_{\text{DNA-protein}}$, over all possible ways by which nucleoprotein complexes, such as nucleosomes, can be positioned on the chromosomal DNA has to be introduced into Eq. (A2) in addition to the path integral, $\int \mathscr{D}\mathbf{R}$. While all these points will be discussed in detail in Appendices B and C, here we would like only to note that the final mathematical form of Eq. (A2) does not change much in the presence of DNA-binding proteins:

$$Z = \sum_{\text{DNA-protein}} \int_{B_{R_0}} d\mathbf{r}_0 \int e^{-\beta E[\mathbf{R}]} \mathscr{D}\mathbf{R} \tag{A3}$$

So far, we have discussed the partition function of DNA in the absence of any long-range interactions between the system components, such as electrostatic interactions between DNA segments, ions and other molecules freely diffusing in surrounding solution. To include these interactions into the partition function calculations, we first need to make a few comments about the studied biological systems.

In the model, viral particles are represented by spherical shells containing viral DNA, which are submerged in surrounding buffer solution, whose volume, $V_{\text{buf}}$, is much larger than the volume of individual viral particles, $V_{\text{vir}}$ ($V_{\text{buf}} \ggg V_{\text{vir}}$). Furthermore, following previous experimental findings [6, 7], it is assumed that ions can freely move between the inner space of the capsid and outer environment, with the total concentrations of monovalent positive ($Na^+$ and $K^+$) and negative ($Cl^-$) ions in the buffering solution both being equal to the same physiological level of $c_{\text{ions}} = 150$ mM [8], see schematic Figure 2(f).

Analogously, nuclei of living cells are represented in the model by spherically shaped nuclear envelopes (NE), which enclose the cell nucleoplasm of $V_{\text{nucl}}$ volume containing the chromosomal DNA. Since experimental studies suggest that ions can freely move between the cell cytosol and nucleoplasm [9, 10], it is assumed that as in the case of viral particles, the cytosol serves the role of buffering solution, in which the total concentrations of monovalent positive ions ($Na^+$ and $K^+$), on one hand, and monovalent negative ions ($Cl^-$) and electrically charged metabolites, on the other, are maintained by the cell at the same physiological level of $c_{\text{ions}} = 150$ mM [11–14]. Furthermore, staining of living cells with ANG-2 sodium indicator demonstrates lack of pronounced exclusion regions for $Na^+$ ions in nuclei of living cells [Figure S5], suggesting that ions can access the entire nucleus volume rather freely. Thus, the cytosol and the nucleus comprise so-called osmotically active volume of a cell, $V_{\text{cell}}^{\text{osm}} = V_{\text{nucl}} + V_{\text{cyto}}$, which has been previously found to occupy $\sim 70\%$ of the total cell volume: $V_{\text{cell}}^{\text{osm}} = 70\% \cdot V_{\text{cell}}$ [15, 16]. Here $V_{\text{cyto}}$ is the cytosol volume.

While there seems to be not much difference between living cells and viral particles in terms of their physical description, what really distinguishes these two cases is the presence of a large amount of DNA-binding proteins in living cells, which are absent in viral particles.

Indeed, packaging of long chromosomal DNA into a tiny nuclear space in eukaryotic cells is mainly done with the help of special DNA-architectural proteins, histones. These proteins assemble into positively charged octamer complexes that wrap negatively charged DNA around themselves, leading to formation of compact nucleosomes. Nucleosome assembly is performed with the help of histone-binding chaperones, such as Asf1 or Nap1, which are involved in transportation of H2A·H2B and H3·H4 histone dimers into the cell nucleus [17, 18]. Experimental studies show that histones almost never can be found freely diffusing inside living cells on their own since as soon as they are synthesized or dissociate from DNA they immediately become picked up by histone-binding chaperons [17, 18]. Thus, as mentioned in *Methods* section, to account for this fact, we introduced two types of histone-binding chaperones into the model: 1) $c_1$ chaperones that can either be in unloaded ($c_1$u) or histone-bound ($c_1$b) state, in which they form a complex with H2A·H2B dimer, and 2) $c_2$ chaperones that also can be either in unloaded ($c_2$u) or histone-bound ($c_2$b) state, but this time forming a complex with H3·H4 dimer, see schematic Figure 1(a).

Since experimental data suggest that several main histone-binding chaperones, such as Nap1, carry both nuclear localization and nuclear export signals, making it possible for them to shuttle between the two cell compartments [19, 20], in our model it is assumed for simplicity that both histone-loaded and unloaded chaperones $c_1$ and $c_2$ can move through nuclear pores without any restriction, see Figure 1(a).

From the primary sequence of histone-binding chaperones it can be found that typically they carry a strong negative electrical charge. It is easy to estimate the average charge of histone-binding chaperones under physiological conditions based on the measurements of the nuclear electrostatic potential ($\sim -10$ mV to $-4$ mV [10, 21]) and the distribution of the mobile histone fraction between the cell nucleus and cytosol [22]. By using the Boltzmann law, it can be shown that the average net electrical charge of histone-loaded chaperones lies somewhere in the range from $+5q_e$ to

$+13q_\mathrm{e}$. Here $q_\mathrm{e} = 1.6 \cdot 10^{-19}$ C is the elementary charge. Since the average electrical charge of H2A·H2B and H3·H4 histone dimers is $\sim +37.2\,q_\mathrm{e}$, it can be seen that binding of chaperones leads to electrostatic screening of a large part of the positive histone charge. Based on these estimations, the electrical charge of histone-bound chaperones in the model was put equal to $q_{\mathrm{c_1 b}} = q_{\mathrm{c_2 b}} = +9\,q_\mathrm{e}$; whereas, the electrical charge of unloaded chaperones was set to $q_{\mathrm{c_1 u}} = q_{\mathrm{c_2 u}} = -28.2\,q_\mathrm{e}$.

As viral particles and cell nuclei are very similar to each other in terms of physical description (apart from the DNA-binding proteins), we will mainly focus below on derivation of formulas for the more complex case of chromosomal DNA organization in a cell nucleus. Yet, for experienced readers it will be rather straightforward to obtain similar mathematical expressions for the viral particle scenario following the same logic, which for this reason will not be considered in detail in Appendix sections below.

Finally, it can be shown that electrostatic interactions between electrically charged molecules, such as DNA, ion, metabolites and proteins, make much more larger contribution into the system free energy than the volume-exclusion effect under physiological conditions, see the main text and Appendix L). Thus, we mainly focus on electrostatic interactions between the system components in this study, neglecting for the most part of the work the volume-exclusion effect.

To incorporate electrostatic forces into the partition function calculations, the first thing we need to do is to define an electrical charge density function, $\rho_\mathrm{e}(\mathbf{r})$:

$$\rho_\mathrm{e}(\mathbf{r}) = \sum_j \sum_{k=1}^{n_j} q_j \delta(\mathbf{r} - \mathbf{r}_{jk}) + \sum_{u=1}^{Q} \int_0^{L_u} \rho_\mathrm{d}^u(s_u)\, \delta(\mathbf{r} - \mathbf{r}_\mathrm{d}^u(s_u))\, \mathrm{d}s_u \tag{A4}$$

Here the first sum, $\sum_j$, runs over all of types of charged particles, except the DNA, that can be found diffusing in the cell cytosol and nucleoplasm (i.e., charged ions, metabolites, proteins, etc.). The second sum, $\sum_{k=1}^{n_j}$, runs over all charged particles of the same type, $j$, where $n_j$ is their total number. $q_j$ is the electrical charge of the $j^\mathrm{th}$ type of particles, and $\mathbf{r}_{jk}$ is the position vector of the $k^\mathrm{th}$ particle of type $j$. Sum $\sum_{u=1}^{Q}$ runs over all chromosomes confined inside the cell nucleus, which are enumerated by index $u$. $Q$ is the total number of chromosomes in the cell. $L_u$ is the contour length of the $u^\mathrm{th}$ chromosome. $\mathbf{r}_\mathrm{d}^u(s_u)$ is the position vector of a point situating on the $u^\mathrm{th}$ chromosome, which corresponds to the arc length, $s_u$, measured along the contour of the respective DNA polymer ($s_u \in [0, L_u]$). $\rho_\mathrm{d}^u(s_u)$ is the net linear charge density of DNA in the $u^\mathrm{th}$ chromosome, which in the general case may depend on the arc length, $s_u$. Indeed, while bare DNA has a uniform linear charge density of $\rho_\mathrm{d} = 2e^- = -2q_\mathrm{e}$ per single base-pair, the net charge of protein-covered DNA may be of a smaller value due to the electrostatic screening effect created by positively charged DNA-bound proteins, such as histones. Dimensionality of $\rho_\mathrm{d}^u$ is: $[\rho_\mathrm{d}^u] = \mathrm{C/m}$.

We would like to stress that the double sum in the left part of Eq. (A4) corresponds to freely diffusing molecules; whereas, the integral on the right side describes the linear charge density of protein-covered DNA. Thus, in the case of DNA-binding proteins, the total number of mobile molecules in the cytosol and nucleoplasm, $n_j$, is not fixed as these proteins can be sequestered from solution via binding to DNA.

Anyway, by having the electrical charge density function, $\rho_\mathrm{e}(\mathbf{r})$, one then can calculate the partition function of the system of DNA polymers comprising the cell chromosomes by using the following formula:

$$Z = \sum_{\text{DNA-protein}} \prod_j \left[ \frac{1}{n_j!} \prod_{k=1}^{n_j} \int_{V_j} \frac{Z_j^\mathrm{in} \mathrm{d}\mathbf{r}_{jk}}{\Lambda_j^3} \right] \times \prod_{u=1}^{Q} \left[ \int_{V_\mathrm{nucl}} \mathrm{d}\mathbf{r}_0^{(u)} \int e^{-\beta E_u[\mathbf{R}^{(u)}]} \mathscr{D}\mathbf{R}^{(u)} \right] e^{-\frac{\beta}{2} \int_{\mathbb{R}^3} \mathrm{d}\mathbf{r} \int_{\mathbb{R}^3} \mathrm{d}\mathbf{r}'\, \rho_\mathrm{e}(\mathbf{r}) U_\mathrm{e}(\mathbf{r}-\mathbf{r}') \rho_\mathrm{e}(\mathbf{r}')} \tag{A5}$$

Here $\Lambda_j$ and $Z_j^\mathrm{in}$ are the thermal de Broglie wavelength and the partition function describing inner degrees of freedom of the $j^\mathrm{th}$ type of particles, respectively. $U_\mathrm{e}(\mathbf{r}) = \frac{1}{4\pi\varepsilon r}$ is the core part of the electrostatic potential, where $r = \|\mathbf{r}\|$ is the length of the distance vector, $\mathbf{r}$, between particles. Dimensionality of $U_\mathrm{e}$ is: $[U_\mathrm{e}] = 1/\mathrm{F}$. $\varepsilon = \varepsilon_r \varepsilon_0$ is the water permittivity, which equals to the product of the water relative permittivity, $\varepsilon_r$ ($\varepsilon_r \sim 75$ at 37°C), and the vacuum permittivity, $\varepsilon_0 = 8.85 \cdot 10^{-12}$ F/m. $E_u[\mathbf{R}^{(u)}]$ is the total energy of DNA comprising the $u^\mathrm{th}$ chromosome, which includes all of the energy terms from Eq. (A1) and those related to DNA interaction with DNA-binding proteins, see

Appendix B for details. $\mathbf{R}^{(u)}$ is a shorthand notation of $\mathbf{R}^{(u)}(s)$ function representing the conformation of DNA in the $u^{\text{th}}$ chromosome.

In the above formula, each integral $\int_{V_{\text{nucl}}}\mathbf{r}_0^{(u)}$, $u = 1, ..., Q$ is calculated over all possible values of the position vector, $\mathbf{r}_0^{(u)}$, describing the starting point of DNA in the $u^{\text{th}}$ chromosome. Similarly, each functional integral, $\int \mathscr{D}\mathbf{R}^{(u)}$, is carried over all possible conformations of the DNA comprising the $u^{\text{th}}$ chromosome. $\int_{\mathbb{R}^3}\mathrm{d}\mathbf{r}$ and $\int_{\mathbb{R}^3}\mathrm{d}\mathbf{r}'$ integrations are performed over all 3D space, $\mathbb{R}^3$. As for $\int_{V_j}\mathrm{d}\mathbf{r}_{jk}$ integrals, they are calculated over the volume $V_j$ accessible by diffusing particles of type $j$. For example, in the case of particles that can freely shuttle between the cell cytosol and nucleus, $V_j$ simply equals to the osmotically active volume of a cell: $V_j = V_{\text{cell}}^{\text{osm}}$. On the other hand, for cytosolic macromolecules, which cannot move through nuclear pore complexes into the cell nucleus, we have: $V_j = V_{\text{cyto}}$. As we will see in Appendix K, this results in generation of osmotic pressure on the NE by cytosolic macromolecules, which plays an important role in regulation of the cell nucleus volume.

It should be noted that while all of the terms in Eq. (A5) have clear physical interpretations, this formula has a limited use as the energy term in the right-most exponential function, $\rho_{\text{e}}(\mathbf{r})U_{\text{e}}(\mathbf{r}-\mathbf{r}')\rho_{\text{e}}(\mathbf{r}')$, requires calculation of the interaction energy between each pair of electrically charged particles present in a cell, which is beyond capabilities of modern computers.

To circumvent this drawback of Eq. (A5), it is convenient instead to represent electrostatic interaction between charged particles by using an alternative way – via an action, $S_\psi[\mathbf{R}^{(u)}, \mathbf{r}_{jk}]$, of a randomly fluctuating electrostatic potential, $\psi$, which depends on the conformation of each DNA polymer as well as the position of each electrically charged particle:

$$S_\psi[\mathbf{R}^{(u)}, \mathbf{r}_{jk}] = \int_{\mathbb{R}^3}\mathrm{d}\mathbf{r}\,\rho_{\text{e}}(\mathbf{r})\psi(\mathbf{r}) - \frac{1}{2}\int_{\mathbb{R}^3}\mathrm{d}\mathbf{r}\int_{\mathbb{R}^3}\mathrm{d}\mathbf{r}'\,\psi(\mathbf{r})\,U_{\text{e}}^{-1}(\mathbf{r}-\mathbf{r}')\,\psi(\mathbf{r}') \tag{A6}$$

Where $\psi$ has dimensionality of: $[\psi] = \text{V}$. In the above equation, $U_{\text{e}}^{-1}(\mathbf{r}-\mathbf{r}')$ is the functional inverse of $U_{\text{e}}(\mathbf{r}-\mathbf{r}')$, which is defined as a generalized function satisfying the following mathematical condition:

$$\int_{\mathbb{R}^3}\mathrm{d}\mathbf{r}'\,U_{\text{e}}^{-1}(\mathbf{r}-\mathbf{r}')\,U_{\text{e}}(\mathbf{r}'-\mathbf{r}'') = \int_{\mathbb{R}^3}\mathrm{d}\mathbf{r}'\,U_{\text{e}}(\mathbf{r}-\mathbf{r}')\,U_{\text{e}}^{-1}(\mathbf{r}'-\mathbf{r}'') = \delta(\mathbf{r}-\mathbf{r}'') \tag{A7}$$

Thus, dimensionality of $U_{\text{e}}^{-1}$ function is: $[U_{\text{e}}^{-1}] = \text{F/m}^6$.

By using Eq. (A6) and Eq. (A7), it is then not hard to see that:

$$\frac{1}{2}\int_{\mathbb{R}^3}\mathrm{d}\mathbf{r}\int_{\mathbb{R}^3}\mathrm{d}\mathbf{r}'\,\rho_{\text{e}}(\mathbf{r})U_{\text{e}}(\mathbf{r}-\mathbf{r}')\rho_{\text{e}}(\mathbf{r}') = S_\psi[\mathbf{R}^{(u)}, \mathbf{r}_{jk}] + \frac{1}{2}\int_{\mathbb{R}^3}\mathrm{d}\mathbf{r}\int_{\mathbb{R}^3}\mathrm{d}\mathbf{r}'\,\widetilde{\psi}(\mathbf{r})\,U_{\text{e}}^{-1}(\mathbf{r}-\mathbf{r}')\,\widetilde{\psi}(\mathbf{r}') \tag{A8}$$

Where $\widetilde{\psi}(\mathbf{r})$ is a shifted electrostatic potential defined as:

$$\widetilde{\psi}(\mathbf{r}) = \psi(\mathbf{r}) - \int_{\mathbb{R}^3}\mathrm{d}\mathbf{r}''\,U_{\text{e}}(\mathbf{r}-\mathbf{r}'')\,\rho_{\text{e}}(\mathbf{r}'') \tag{A9}$$

As a result, it immediately follows that:

$$\int e^{-\beta S_\psi[\mathbf{R}^{(u)},\mathbf{r}_{jk}]}\mathscr{D}\psi = e^{-\frac{\beta}{2}\int_{\mathbb{R}^3}\mathrm{d}\mathbf{r}\int_{\mathbb{R}^3}\mathrm{d}\mathbf{r}'\,\rho_{\text{e}}(\mathbf{r})U_{\text{e}}(\mathbf{r}-\mathbf{r}')\rho_{\text{e}}(\mathbf{r}')}\int e^{\frac{\beta}{2}\int_{\mathbb{R}^3}\mathrm{d}\mathbf{r}\int_{\mathbb{R}^3}\mathrm{d}\mathbf{r}'\,\widetilde{\psi}(\mathbf{r})\,U_{\text{e}}^{-1}(\mathbf{r}-\mathbf{r}')\,\widetilde{\psi}(\mathbf{r}')}\mathscr{D}\widetilde{\psi} =$$
$$= \text{const}\times e^{-\frac{\beta}{2}\int_{\mathbb{R}^3}\mathrm{d}\mathbf{r}\int_{\mathbb{R}^3}\mathrm{d}\mathbf{r}'\,\rho_{\text{e}}(\mathbf{r})U_{\text{e}}(\mathbf{r}-\mathbf{r}')\rho_{\text{e}}(\mathbf{r}')} \tag{A10}$$

Here we have taken into account that $\int\mathscr{D}\widetilde{\psi} = \int\mathscr{D}\psi$, where $\int\mathscr{D}\psi$ and $\int\mathscr{D}\widetilde{\psi}$ are field integrals over all possible configurations of the electrostatic potentials $\psi$ and $\widetilde{\psi}$, respectively, see ref. [5].

Substituting the above formula into Eq. (A5), it can be seen that:

$$Z = \frac{\sum_{\text{DNA-protein}} \prod_j \left[\frac{1}{n_j!} \prod_{k=1}^{n_j} \int_{V_j} \frac{Z_j^{\text{in}} \mathrm{d}\mathbf{r}_{jk}}{\Lambda_j^3}\right] \times \prod_{u=1}^{Q} \left[\int_{V_{\text{nucl}}} \mathrm{d}\mathbf{r}_0^{(u)} \int e^{-\beta E_u[\mathbf{R}^{(u)}]} \mathscr{D}\mathbf{R}^{(u)}\right] \int e^{-\beta S_\psi[\mathbf{R}^{(u)},\mathbf{r}_{jk}]} \mathscr{D}\psi}{\int e^{\frac{\beta}{2} \int_{\mathrm{I\!R}^3} \mathrm{d}\mathbf{r} \int_{\mathrm{I\!R}^3} \mathrm{d}\mathbf{r}' \, \psi(\mathbf{r}) \, U_{\mathrm{e}}^{-1}(\mathbf{r}-\mathbf{r}') \, \psi(\mathbf{r}')} \mathscr{D}\psi} \tag{A11}$$

Since multiplication of the partition function by an arbitrary constant simply leads to a fixed offset of the system free energy, which does not have any effect on behaviour of the studied system, the denominator in Eq. (A11) can be omitted. As a result, the partition function can be rewritten in the following simpler form:

$$Z = \sum_{\text{DNA-protein}} \prod_j \left[\frac{1}{n_j!} \prod_{k=1}^{n_j} \int_{V_j} \frac{Z_j^{\text{in}} \mathrm{d}\mathbf{r}_{jk}}{\Lambda_j^3}\right] \times \prod_{u=1}^{Q} \left[\int_{V_{\text{nucl}}} \mathrm{d}\mathbf{r}_0^{(u)} \int e^{-\beta E_u[\mathbf{R}^{(u)}]} \mathscr{D}\mathbf{R}^{(u)}\right] \int e^{-\beta S_\psi[\mathbf{R}^{(u)},\mathbf{r}_{jk}]} \mathscr{D}\psi \tag{A12}$$

To further streamline the above formula, let's find an explicit mathematical expression for $\int_{\mathrm{I\!R}^3} \mathrm{d}\mathbf{r} \, \rho_{\mathrm{e}}(\mathbf{r})\psi(\mathbf{r})$ term from Eq. (A6). By using Eq. (A4), we get:

$$\int_{\mathrm{I\!R}^3} \mathrm{d}\mathbf{r} \, \rho_{\mathrm{e}}(\mathbf{r})\psi(\mathbf{r}) = \sum_j \sum_{k=1}^{n_j} q_j \psi(\mathbf{r}_{jk}) + \sum_{u=1}^{Q} \int_0^{L_u} \rho_{\mathrm{d}}^u(s_u) \, \psi(\mathbf{r}_{\mathrm{d}}^u(s_u)) \, \mathrm{d}s_u \tag{A13}$$

Combining together Eq. (A6), Eq. (A12) and Eq. (A13), it is not hard to see that:

$$Z = \sum_{\text{DNA-protein}} \int \mathscr{D}\psi \prod_j \frac{(Z_j^\psi)^{n_j}}{n_j!} \times e^{\frac{\beta}{2} \int_{\mathrm{I\!R}^3} \mathrm{d}\mathbf{r} \int_{\mathrm{I\!R}^3} \mathrm{d}\mathbf{r}' \, \psi(\mathbf{r}) \, U_{\mathrm{e}}^{-1}(\mathbf{r}-\mathbf{r}') \, \psi(\mathbf{r}')} \times$$
$$\times \prod_{u=1}^{Q} \left[\int_{V_{\text{nucl}}} \mathrm{d}\mathbf{r}_0^{(u)} \int e^{-\beta\left\{E_u[\mathbf{R}^{(u)}] + \int_0^{L_u} \rho_{\mathrm{d}}^u(s_u) \, \psi(\mathbf{r}_{\mathrm{d}}^u(s_u)) \, \mathrm{d}s_u\right\}} \mathscr{D}\mathbf{R}^{(u)}\right] \tag{A14}$$

Here $Z_j^\psi$ is the partition function of an individual particle of type $j$ in the field $\psi$:

$$Z_j^\psi = \frac{Z_j^{\text{in}}}{\Lambda_j^3} \int_{V_j} e^{-\beta q_j \psi(\mathbf{r})} \, \mathrm{d}\mathbf{r} = \frac{Z_j^{\text{in}} I_j^\psi}{\Lambda_j^3}, \quad \text{where} \quad I_j^\psi = \int_{V_j} e^{-\beta q_j \psi(\mathbf{r})} \, \mathrm{d}\mathbf{r} \tag{A15}$$

By applying the Stirling's approximation ($\ln n_j! \approx n_j \ln n_j - n_j$), Eq. (A14) can be reduced to:

$$Z \approx \sum_{\text{DNA-protein}} \int \mathscr{D}\psi \prod_j \left(\frac{e Z_j^\psi}{n_j}\right)^{n_j} \times e^{\frac{\beta}{2} \int_{\mathrm{I\!R}^3} \mathrm{d}\mathbf{r} \int_{\mathrm{I\!R}^3} \mathrm{d}\mathbf{r}' \, \psi(\mathbf{r}) \, U_{\mathrm{e}}^{-1}(\mathbf{r}-\mathbf{r}') \, \psi(\mathbf{r}')} \times$$
$$\times \prod_{u=1}^{Q} \left[\int_{V_{\text{nucl}}} \mathrm{d}\mathbf{r}_0^{(u)} \int e^{-\beta\left\{E_u[\mathbf{R}^{(u)}] + \int_0^{L_u} \rho_{\mathrm{d}}^u(s_u) \, \psi(\mathbf{r}_{\mathrm{d}}^u(s_u)) \, \mathrm{d}s_u\right\}} \mathscr{D}\mathbf{R}^{(u)}\right] \tag{A16}$$

To further simplify the above formula, we need to recall that $\prod_j$ product runs over all types of diffusing ions and molecules, $j$, that can be found inside a living cell. This includes histone-bound and histone-free chaperones, whose total number can vary depending on how many histones are bound to DNA. However, it should be noted that experimental measurements suggest that these numbers experience only very small fluctuations under physiological conditions. Indeed, it has been previously shown that the mobile fraction of histones constitutes $\sim 1-9\%$ of all histones found in mammalian cells [22–24], which seems to be enough to cover basic needs of cells as fluctuations in the cytoplasmic histone concentration were found to be of the order of only $\sim 8-10\%$ [22]. Thus, it can be concluded that deviations of the total number of histone-bound and histone-unloaded chaperones as well as the total number of nucleosomes formed on the chromosomal DNA from their average values are very small at any given moment of time.

Specifically, let $n^0_{\mathrm{c_1 b}}$, $n^0_{\mathrm{c_2 b}}$, $n^0_{\mathrm{c_1 u}}$ and $n^0_{\mathrm{c_2 u}}$ be the average total numbers of $\mathrm{c_1}$ and $\mathrm{c_2}$ chaperones in histone-bound and unloaded states, and $n^0_{\rm nucl}$ be the average total number of nucleosomes on the chromosomal DNA in a living cell. Then the above notes regarding fluctuations of the histone-bound and unloaded chaperone numbers, $n_{\rm c_1b}$, $n_{\rm c_2b}$, $n_{\rm c_1u}$ and $n_{\rm c_2u}$, as well as the nucleosome number, $n_{\rm nucl}$, are equivalent to the following inequalities:

$$\begin{aligned} &|n_{\rm c_1b} - n^0_{\rm c_1b}| \ll n^0_{\rm c_1b} \quad \text{and} \quad |n_{\rm c_2b} - n^0_{\rm c_2b}| \ll n^0_{\rm c_2b} \quad \text{and} \quad |n_{\rm nucl} - n^0_{\rm nucl}| \ll n^0_{\rm nucl} \\ &|n_{\rm c_1u} - n^0_{\rm c_1u}| \ll n^0_{\rm c_1u} \quad \text{and} \quad |n_{\rm c_2u} - n^0_{\rm c_2u}| \ll n^0_{\rm c_2u} \end{aligned} \tag{A17}$$

In other words, histone-binding chaperones buffer potential changes in the intracellular concentration of the mobile fraction of histones, helping to maintain cell homeostasis.

Using the introduced notations, it is straightforward to see that the total numbers of histone-bound and unloaded chaperones in the case when there are $n_{\rm nucl}$ nucleosomes on the chromosomal DNA equal to $n^0_{\rm c_1b} - 2n_{\rm nucl} + 2n^0_{\rm nucl}$ and $n^0_{\rm c_1u} + 2n_{\rm nucl} - 2n^0_{\rm nucl}$, respectively, with similar equations holding for $\rm c_2$ chaperones as well. Hence, by taking into account the above notes, it is not hard to see that Eq. (A16) can be rewritten as:

$$Z \approx \int \mathscr{D}\psi \prod_j{}' \left(\frac{eZ^\psi_j}{n_j}\right)^{n_j} \times \left(\frac{eZ^\psi_{\rm c_1b}}{n^0_{\rm c_1b}}\right)^{n^0_{\rm c_1b}+2n^0_{\rm nucl}} \left(\frac{eZ^\psi_{\rm c_2b}}{n^0_{\rm c_2b}}\right)^{n^0_{\rm c_2b}+2n^0_{\rm nucl}} \left(\frac{eZ^\psi_{\rm c_1u}}{n^0_{\rm c_1u}}\right)^{n^0_{\rm c_1u}-2n^0_{\rm nucl}} \left(\frac{eZ^\psi_{\rm c_2u}}{n^0_{\rm c_2u}}\right)^{n^0_{\rm c_2u}-2n^0_{\rm nucl}} \times$$

$$\times\, Z_\psi\, e^{\frac{\beta}{2}\int_{\rm I\!R^3}d\mathbf{r}\int_{\rm I\!R^3}d\mathbf{r}'\,\psi(\mathbf{r})\,U_{\rm e}^{-1}(\mathbf{r}-\mathbf{r}')\,\psi(\mathbf{r}')} \tag{A18}$$

Where $\prod'_j$ is the product which runs over all molecule types, $j$, except for histone-bound and unloaded chaperones. $Z^\psi_{\rm c_1b}$, $Z^\psi_{\rm c_2b}$, $Z^\psi_{\rm c_1u}$ and $Z^\psi_{\rm c_2u}$ are the partition functions of histone-bound and unloaded chaperones, which are defined by Eq. (A15). Finally, $Z_\psi$ is the partition function of DNA covered with nucleosomes in the electrostatic potential $\psi$:

$$Z_\psi = \sum_{\text{DNA-protein}} \left(\frac{Z^\psi_{\rm c_1b}}{n^0_{\rm c_1b}}\right)^{-2n_{\rm nucl}} \left(\frac{Z^\psi_{\rm c_2b}}{n^0_{\rm c_2b}}\right)^{-2n_{\rm nucl}} \left(\frac{Z^\psi_{\rm c_1u}}{n^0_{\rm c_1u}}\right)^{2n_{\rm nucl}} \left(\frac{Z^\psi_{\rm c_2u}}{n^0_{\rm c_2u}}\right)^{2n_{\rm nucl}} \times$$

$$\times \prod_{u=1}^{Q}\left[\int_{V_{\rm nucl}} d\mathbf{r}_0^{(u)} \int e^{-\beta\left\{E_u[\mathbf{R}^{(u)}]+\int_0^{L_u}\rho^u_{\rm d}(s_u)\,\psi(\mathbf{r}^u_{\rm d}(s_u))\,ds_u\right\}}\mathscr{D}\mathbf{R}^{(u)}\right] \tag{A19}$$

For the sake of formulas simplicity, it makes sense to include $Z^\psi_{\rm c_1b}$, $Z^\psi_{\rm c_2b}$, $Z^\psi_{\rm c_1u}$ and $Z^\psi_{\rm c_2u}$ terms back into the product $\prod'_j$ in Eq. (A18), and as a result, one can easily obtain the following formula for the partition function:

$$Z = \int \mathscr{D}\psi \prod_j \left(\frac{eZ^\psi_j}{n_j}\right)^{\tilde{n}_j} \times Z_\psi\, e^{\frac{\beta}{2}\int_{\rm I\!R^3}d\mathbf{r}\int_{\rm I\!R^3}d\mathbf{r}'\,\psi(\mathbf{r})\,U_{\rm e}^{-1}(\mathbf{r}-\mathbf{r}')\,\psi(\mathbf{r}')} \tag{A20}$$

Where for ions and non-chaperone molecules $\tilde{n}_j = n_j$; whereas in the case of histone-binding chaperones, $n_j$ equals to $n^0_{\rm c_1b}$, $n^0_{\rm c_2b}$, $n^0_{\rm c_1u}$ or $n^0_{\rm c_2u}$ and $\tilde{n}_j$ equals to $n^0_{\rm c_1b} + 2n^0_{\rm nucl}$, $n^0_{\rm c_2b} + 2n^0_{\rm nucl}$, $n^0_{\rm c_1u} - 2n^0_{\rm nucl}$ or $n^0_{\rm c_2u} - 2n^0_{\rm nucl}$ for histone-bound and unloaded chaperones $c_1$ and $c_2$, respectively.

To calculate the field integral in Eq. (A20), in this study we use the stationary phase approximation. Namely, it can be shown that the value of this functional integral is approximately proportional to [5]:

$$Z \propto \prod_j \left(\frac{eZ^{\psi_{\rm sp}}_j}{n_j}\right)^{\tilde{n}_j} \times Z_{\psi_{\rm sp}} e^{\frac{\beta}{2}\int_{\rm I\!R^3}d\mathbf{r}\int_{\rm I\!R^3}d\mathbf{r}'\,\psi_{\rm sp}(\mathbf{r})U_{\rm e}^{-1}(\mathbf{r}-\mathbf{r}')\psi_{\rm sp}(\mathbf{r}')} \tag{A21}$$

Where $\psi_{\rm sp}$ is the potential field that extremizes the integrand function in Eq. (A20). This formula defines the partition function up to a constant prefactor, which is not important for this study since it results only in a small constant offset of the total free energy of the system that does not affect its behaviour [5].

To find the stationary phase electrostatic potential, $\psi_{\rm sp}$, one needs to solve the following functional equation:

$$\frac{\delta}{\delta\psi(\mathbf{r})}\left[\sum_j \tilde{n}_j \ln Z_j^{\psi} + \ln Z_{\psi} + \frac{\beta}{2}\int_{\rm I\!R^3} \mathrm{d}\mathbf{r} \int_{\rm I\!R^3} \mathrm{d}\mathbf{r}'\, \psi(\mathbf{r}) U_{\rm e}^{-1}(\mathbf{r}-\mathbf{r}')\psi(\mathbf{r}')\right] = 0 \tag{A22}$$

Here $\sum_j$ is the sum over all particle types, $j$, diffusing in a living cell, and $\frac{\delta}{\delta\psi(\mathbf{r})}$ is a functional derivative with respect to the field $\psi(\mathbf{r})$.

Since in this study the cell nucleus is represented by a sphere, it is clear that its electrostatic potential, $\psi$, in the general case will be spherically symmetric, depending only on the radial distance, $r = \|\mathbf{r}\|$, measured from the center of the nucleus. On the other hand, outside of the nucleus the electrostatic field drops within a distance of a few Debye lengths ($\lambda_D \approx 0.8$ nm) from NE to a constant level corresponding to the cytoplasmic electrostatic potential, $\psi_{\rm cyto}$, which we will use as a reference point in our calculations. As a result, it can be shown that to solve Eq. (A22), all we need to do is to find a nuclear electrostatic potential, $\psi(r)$ ($r \leq R_{\rm nucl}$), which satisfies a spherically symmetric version of Eq. (A22):

$$\frac{\delta}{\delta\psi(r)}\left[\sum_j \tilde{n}_j \ln Z_j^{\psi} + \ln Z_{\psi} + \frac{\beta}{2}\int_{\rm I\!R^3} \mathrm{d}\mathbf{r} \int_{\rm I\!R^3} \mathrm{d}\mathbf{r}'\, \psi(\mathbf{r}) U_{\rm e}^{-1}(\mathbf{r}-\mathbf{r}')\psi(\mathbf{r}')\right] = 0 \tag{A23}$$

Where this time the functional derivative, $\frac{\delta}{\delta\psi(r)}$, is taken with respect to a spherically symmetric field, $\psi(r)$.

From Eq. (A23) it can be seen that to obtain the potential $\psi_{\rm sp}$, we need in the first place to know the exact mathematical forms of the following three functions: 1) $U_{\rm e}^{-1}(\mathbf{r})$, 2) $\frac{\delta}{\delta\psi(r)} \ln Z_j^{\psi}$ and 3) $\frac{\delta}{\delta\psi(r)} \ln Z_{\psi}$. Let's find them one by one.

##### *A. Functional inverse of potential $U_e$.*

To obtain the functional inverse of $U_{\rm e}(\mathbf{r})$, all we need to do is to recall the convolution theorem which states that the Fourier transform of a convolution of two functions equals to the pointwise product of their Fourier transforms. In application to Eq. (A7), this means that:

$$\mathscr{F}[U_{\rm e} * U_{\rm e}^{-1}] = \mathscr{F}[U_{\rm e}] \cdot \mathscr{F}[U_{\rm e}^{-1}] = \mathscr{F}[\delta] = 1 \tag{A24}$$

Where $\mathscr{F}[U_{\rm e}]$, $\mathscr{F}[U_{\rm e}^{-1}]$, $\mathscr{F}[U_{\rm e} * U_{\rm e}^{-1}]$ and $\mathscr{F}[\delta]$ are Fourier transforms of the potential $U_{\rm e}$, its functional inverse, $U_{\rm e}^{-1}$, their convolution, $U_{\rm e} * U_{\rm e}^{-1}$, and the Dirac $\delta$-function, respectively.

It is straightforward to find the Fourier transform of the potential $U_{\rm e}$ by performing integration in spherical coordinates, in which $\mathbf{z}$-axis is collinear to the wave-vector $\mathbf{k}$:

$$\mathscr{F}[U_{\rm e}] = \frac{1}{4\pi\varepsilon}\lim_{m\to+0}\int_{\rm I\!R^3} \frac{1}{r} e^{-mr} e^{-i(\mathbf{k}\cdot\mathbf{r})}\,\mathrm{d}\mathbf{r} = \frac{1}{4\pi\varepsilon}\lim_{m\to+0}\int_0^{2\pi}\mathrm{d}\varphi\int_0^{\pi}\mathrm{d}\theta\int_0^{\infty}\mathrm{d}r\, r e^{-mr} e^{-ikr\cos\theta}\sin\theta = \frac{1}{\varepsilon k^2} \tag{A25}$$

Where $k = \|\mathbf{k}\|$.

From Eq. (A24) and Eq. (A25) it then immediately follows that:

$$\mathscr{F}[U_{\rm e}^{-1}] = \varepsilon k^2 \tag{A26}$$

And as a result, we have:

$$U_{\rm e}^{-1}(\mathbf{r}) = -\varepsilon\Delta\delta(\mathbf{r}) \tag{A27}$$

Where $\Delta$ is the Laplace operator. To see this, one just needs to substitute the Fourier form of the Dirac $\delta$-function: $\delta(\mathbf{r}) = \frac{1}{(2\pi)^3}\int_{\rm I\!R^3} e^{-i(\mathbf{k}\cdot\mathbf{r})}\mathrm{d}\mathbf{r}$ into Eq. (A27) and then compare the obtained result to Eq. (A26).

By using Eq. (A27) for the functional inverse of the potential $U_\mathrm{e}$, it is easy to see that:

$$\frac{\beta}{2}\frac{\delta}{\delta\psi(r)}\int_{\mathrm{I\!R}^3}\mathrm{d}\mathbf{r}\int_{\mathrm{I\!R}^3}\mathrm{d}\mathbf{r}'\,\psi(\mathbf{r})U_\mathrm{e}^{-1}(\mathbf{r}-\mathbf{r}')\psi(\mathbf{r}')=-2\pi\beta\varepsilon\frac{\delta}{\delta\psi(r)}\int_0^\infty \mathrm{d}r\,r^2\psi(r)\Delta\psi(r)=-4\pi\beta\varepsilon r^2\Delta\psi(r) \tag{A28}$$

*B. Functional derivative of the partition function of freely diffusing ions and molecules.*

As for the functional derivative of $Z_j^\psi$ partition function, from Eq. (A15) it is easy to see that for $r\leq R_\mathrm{nucl}$ we have:

$$\frac{\delta\ln Z_j^\psi}{\delta\psi(r)}=\frac{\delta}{\delta\psi(r)}\ln\left(\frac{Z_j^\mathrm{in}I_j^\psi}{\Lambda_j^3}\right)=-\frac{4\pi\beta q_j}{I_j^\psi}\,\chi_j r^2 e^{-\beta q_j\psi(r)} \tag{A29}$$

Where $\chi_j=1$ if particles of type $j$ can freely move between the cell nucleus and cytosol and $\chi_j=0$, otherwise.

Substituting Eq. (A28) and Eq. (A29) into Eq. (A23), it is easy to see that the functional equation for the stationary phase electrostatic potential, $\psi_\mathrm{sp}$, turns into:

$$\frac{1}{4\pi\beta r^2}\frac{\delta\ln Z_\psi}{\delta\psi(r)}-\sum_j\frac{\tilde{n}_j q_j}{I_j^\psi}\,\chi_j e^{-\beta q_j\psi(r)}-\varepsilon\Delta\psi(r)=0 \tag{A30}$$

While so far we have considered DNA organization in nuclei of living cells, it should be noted that in the case of viral particles, one can arrive to the same Eq. (A30) by using a very similar line of reasoning. Furthermore, by taking into account that the volume of buffering solution is much larger than the volume of the viral capsid (i.e., $V_\mathrm{buf}\ggg V_\mathrm{vir}$), for viral particles we have:

$$I_j^\psi=\int_{V_j}e^{-\beta q_j\psi(\mathbf{r})}\,\mathrm{d}\mathbf{r}\approx V_\mathrm{buf} \tag{A31}$$

Thus, in the case of viral particles, it is easy to see that Eq. (A30) simplifies to:

$$\frac{1}{4\pi\beta r^2}\frac{\delta\ln Z_\psi}{\delta\psi(r)}-\sum_j c_j q_j e^{-\beta q_j\psi(r)}-\varepsilon\Delta\psi(r)=0 \tag{A32}$$

Where we have taken into account that ions can freely exchange between interior and exterior of the viral capsid, putting $\chi_j=1$. In the above equation, $c_j=n_j/V_\mathrm{buf}$ is the average concentration of the $j^\mathrm{th}$ type of ions in the buffer solution.

It is not hard to see that in the absence of DNA the above formula transforms into a very well-known Poisson-Boltzmann equation, which forms the core of the Debye-Hückel theory [25], indicating consistency of the derived formulas with previously published studies:

$$\sum_j c_j q_j e^{-\beta q_j\psi(r)}+\varepsilon\Delta\psi(r)=0 \tag{A33}$$

It is possible to further simplify Eq. (A30) by noting that in the case of living cells the value of $\sum_j$ sum is predominantly determined by contribution of monovalent ions, such as $K^+$, $Na^+$, $Cl^-$, as well as by small cell metabolites, which on average have a negative charge of $\sim -1q_\mathrm{e}$ [11–14]. Indeed, as has been noted after Eq. (A3), previous studies suggest that the total cytosolic concentrations of positive and negative monovalent ions as well as small metabolites are maintained by living cells at a relatively stable level of $c_\mathrm{ions}=150$ mM [11–14]. In contrast, the average protein concentration in living cells has been found to be of the order of only $\sim 0.3-2.5$ mM [26], i.e., two orders of magnitude smaller than the total concentrations of positive and negative monovalent ions and small cell metabolites. Thus, to the first order of approximation, protein contribution to $\sum_j$ sum in Eq. (A30) can be neglected.

Furthermore, under quasi-equilibrium conditions, which are used in this study, spatial distributions of electrically charged ions and small metabolites inside the cytosol and nucleus of a living cell follow the Boltzmann law:

$$c_j(\mathbf{r}) = \frac{\tilde{n}_j}{I_j^{\psi}}\, e^{-\beta q_j \psi(\mathbf{r})} \tag{A34}$$

Thus, for the cytosolic ion concentration, $c_{\rm ions}$, we have:

$$c_{\rm ions} = \frac{1}{I_+^{\psi}} \sum_{\substack{\text{positive} \\ \text{monovalent}}} \tilde{n}_j = \frac{1}{I_-^{\psi}} \sum_{\substack{\text{negative} \\ \text{monovalent}}} \tilde{n}_j \tag{A35}$$

Where the first sum is calculated over $j$ indexes corresponding to positive monovalent ions; whereas, the second sum is performed over $j$ indexes corresponding to negative monovalent ions and small cell metabolites. Here we have taken into account that the electrostatic potential of the cell cytoplasm, $\psi_{\rm cyto}$, is used in our study as a reference point (thus, $\psi_{\rm cyto} = 0$ mV). As for $I_+^{\psi}$ and $I_-^{\psi}$, these are simply $I_j^{\psi}$ integrals of positive and negative monovalent ions, which are equal to:

$$\begin{cases} I_+^{\psi} = \displaystyle\int_{V_{\rm cell}^{\rm osm}} e^{-\beta q_{\rm e} \psi(\mathbf{r})} d\mathbf{r} = V_{\rm cell}^{\rm osm} + 4\pi \int_0^{R_{\rm nucl}} r^2 \left[e^{-\beta q_{\rm e}\psi(r)} - 1\right] dr \\ I_-^{\psi} = \displaystyle\int_{V_{\rm cell}^{\rm osm}} e^{\beta q_{\rm e} \psi(\mathbf{r})} d\mathbf{r} = V_{\rm cell}^{\rm osm} + 4\pi \int_0^{R_{\rm nucl}} r^2 \left[e^{\beta q_{\rm e}\psi(r)} - 1\right] dr \end{cases} \tag{A36}$$

Substituting Eq. (A35) into Eq. (A30), we finally obtain the following equation for the stationary field, $\psi_{\rm sp}$:

$$\frac{1}{4\pi\beta r^2}\frac{\delta \ln Z_{\psi}}{\delta\psi(r)} + 2 q_{\rm e} c_{\rm ions} \sinh[\beta q_{\rm e}\psi(r)] - \varepsilon\Delta\psi(r) = 0 \tag{A37}$$

Where $\sinh(x)$ is the hyperbolic sine function.

In a very similar way, by taking into consideration the dominant contribution of monovalent ions into $\sum_j$ sum of Eq. (A32), it can be shown that in the case of viral particles the stationary phase electrostatic potential is determined by the same functional Eq. (A37), which has been used in our study for both scenarios – viral particles and living cells.

To complete Eq. (A37), all that remains is to find a mathematical expression for $\frac{\delta \ln Z_{\psi}}{\delta\psi(r)}$. In Appendicies B-E, it will be shown how to calculate this functional derivative. But before proceeding to derivation of formulas for $\ln Z_{\psi}$ and $\frac{\delta \ln Z_{\psi}}{\delta\psi(r)}$, let's imaging for a moment that we have found a way to compute them. Then what will be the next step?

In this study, to solve Eq. (A37) in the cell nucleus, we approximate electrostatic field $\psi(r)$ by the first $n_{\rm max}$ terms of the Fourier-Bessel expansion series:

$$\psi(r) = \frac{\psi_0}{2}\left[1 - \tanh\left(\tfrac{r-R_0}{w}\right)\right] + \sum_{n=1}^{n_{\rm max}} \psi_n\, j_0\!\left(\tfrac{\pi n r}{R_{\rm nucl}}\right) \tag{A38}$$

Where the left term represents the average level of the nuclear electrostatic potential, with deviations from it being described by the sum $\sum_n$. In the above formula, $\tanh(x)$ is the hyperbolic tangent function. $R_0$ ($R_0 \approx R_{\rm nucl}$), $w$ and $\psi_0$ are the model parameters characterizing the average shape of the nuclear electrostatic potential. $j_0(x)$ is the spherical Bessel function of the first kind, and $\psi_n$ are expansion coefficients. $n_{\rm max}$, is the total number of the Fourier-Bessel expansion series terms used in the model computations, which is determined by the required accuracy of the final result. In all our calculations, $n_{\rm max}$ was set equal to: $n_{\rm max} = 18$.

By substituting the above expansion formula into the left part of Eq. (A37), squaring it and minimizing the integral of the obtained function over the radial distance, $r$, with respect to $R_0$ and $w$ model parameters as well as the expansion coefficients, $(\psi_0, \psi_1, \psi_2, ...)$, it is possible to find the shape of the field $\psi_{\rm sp}(r)$ at the stationary phase with a desired accuracy level:

$$\min_{[R_0,w,\psi_0,\psi_1,...]} \int_0^{R_{\rm nucl}} \left[Q(r,R_0,w,\psi_0,\psi_1,...)\right]^2 {\rm d}r \to 0 \ \ \text{as} \ \ \psi \to \psi_{\rm sp} \tag{A39}$$

Here $Q(r,R_0,w,\psi_0,\psi_1,...)$ is the left part of Eq. (A37), where the electrostatic potential $\psi$ is defined by Eq. (A38):

$$Q(r,R_0,w,\psi_0,\psi_1,...) = \frac{1}{4\pi\beta r^2}\frac{\delta \ln Z_\psi}{\delta\psi(r)} + 2q_{\rm e}c_{\rm ions}\sinh[\beta q_{\rm e}\psi(r)] - \varepsilon\Delta\psi(r) \tag{A40}$$

For the above minimization procedure we used Nelder-Mead simplex algorithm [27].

By knowing the stationary phase electrostatic potential, $\psi_{\rm sp}$, it is then straightforward to calculate the partition function defined by Eq. (A21). Specifically, by taking logarithm of Eq. (A21) and substituting into it Eq. (A27), it can be found that up to a non-essential constant term, we have:

$$\ln Z = \ln Z_{\psi_{\rm sp}} - \beta\sum_j \tilde{n}_j\mu_j^{\psi_{\rm sp}} - 2\pi\beta\varepsilon\int_0^{R_{\rm nucl}} r^2\psi_{\rm sp}(r)\Delta\psi_{\rm sp}(r)\,{\rm d}r \tag{A41}$$

Here $\mu_j^{\psi_{\rm sp}}$ denotes the electrochemical potential of the $j^{\rm th}$ particle type in field $\psi_{\rm sp}$:

$$\mu_j^{\psi_{\rm sp}} = k_{\rm B}T\ln\left(\frac{n_j}{eZ_j^{\psi_{\rm sp}}}\right) = \mu_j^0 - k_{\rm B}T\ln\left(\frac{I_j^{\psi_{\rm sp}}}{V_j}\right) + k_{\rm B}T\ln\langle c_j\rangle \tag{A42}$$

Where $\langle c_j\rangle = n_j/V_j$ is the average concentration of the $j^{\rm th}$ type of particles in a living cell, and $\mu_j^0$ is their standard chemical potential in zero $\psi$ field ($\psi = 0$): $\mu_j^0 = k_{\rm B}T\ln(\Lambda_j^3/eZ_j^{\rm in})$.

By taking into account that the value of the sum $\sum_j$ in Eq. (A41) is mainly determined by the contribution of monovalent ions, small cell metabolites and cytosolic macromolecules, for which $\tilde{n}_j = n_j$, we can rewrite it as:

$$\ln Z = \ln Z_{\psi_{\rm sp}} - \beta\sum_{\substack{\text{ions,}\\ \text{metabolites,}\\ \text{macromolecules}}} n_j\mu_j^{\psi_{\rm sp}} - 2\pi\beta\varepsilon\int_0^{R_{\rm nucl}} r^2\psi_{\rm sp}(r)\Delta\psi_{\rm sp}(r)\,{\rm d}r \tag{A43}$$

It should be noted that under physiological conditions, the integral on the right side of Eq. (A43) makes a negligible contribution into the partition function, $Z$. Thus, it can be omitted in calculations.

To obtain the final mathematical expression for the total free energy of the system, which includes the main molecular mechanisms involved in the nucleus size regulation, we need to add to Eq. (A43) two additional free energy terms, $G_{\rm NE}$ and $G_{\rm ER}$, associated with the surface tensions of the NE and ER membrane, $\sigma_{\rm NE}$ and $\sigma_{\rm ER}$, see Appendix K. Indeed, as discussed in *Results* section and Appendix K, surface tensions of the NE and ER membrane lead to appearance of $p_{\rm NE}$ and $p_{\rm ER}$ pressures acting on the NE, which play the major role in the nucleus size regulation. As a result, we get the following formula for the total free energy of the system, $G$:

$$G = -k_{\rm B}T\ln Z + G_{\rm NE} + G_{\rm ER} = -k_{\rm B}T\ln Z_{\psi_{\rm sp}} + \sum_{\substack{\text{ions,}\\ \text{metabolites,}\\ \text{macromolecules}}} n_j\mu_j^{\psi_{\rm sp}} + G_{\rm NE} + G_{\rm ER} \tag{A44}$$

By taking derivatives of the total free energy, $G$, or, which is almost the same thing, the logarithm of the partition function, $\ln Z$, with respect to various model parameters, such as the nucleus volume or the binding energy of histones to DNA, etc., it is then possible to find the mean pressure created by DNA and other elements onto the NE, the average DNA occupancy fraction by nucleosomes, and many other quantities characterizing the physical state of the system. The full list of formulas, which were used in this study to calculate all these observables, can be found in

Appendices I-K.

In order to assess stability of the stationary phase potential, $\psi_{\rm sp}$, obtained by solving Eq. (A37), we have calculated the matrix of second derivatives (Hessian) of the logarithm of the partition function, $\ln Z$, defined by Eq. (A43) with respect to the expansion coefficients shown in Eq. (A38) for different sizes of viral particles ($R_{\rm vir} = 20 - 1500$ nm) and nuclei of living cells ($R_{\rm nucl} = 4 - 10$ $\mu$m). The final results of the calculations are shown in Figure S6. As can be seen from the figure, all of the obtained Hessian matrices have a nearly diagonal form. Since the diagonal elements in these matrices are all negative, it follows that they are negative-definite in the entire tested range of viral particles' and cell nuclei sizes. Thus, it can be concluded that $\psi_{\rm sp}$ electrostatic potential is a stable stationary point corresponding to the maximum of the partition function or, which is the same, to the minimum of the free energy, $G$.

It should be emphasized that all of the results presented in this study were obtained by using the mean field approach described above. Unfortunately, at the present stage of the development of polymer field theory, there is no simple way to evaluate the full contribution of fluctuations of the electrostatic potential, $\psi$, to the free energy of a system consisting of confined self-interacting semiflexible polymers, such as DNA. Yet, it is possible to make rough estimations of the correction to the free energy of the system due to fluctuations in the electrostatic potential by using the Gaussian approximation in order to perform field integration in Eq. (A20) over the amplitudes of spherically-symmetric modes, $(\psi_1, \psi_2, ..., \psi_{n_{\rm max}})$ [Eq. (A38)], in the vicinity of the stationary phase potential, $\psi_{\rm sp}$. The final results of such calculations are displayed in Figure S7.

As can be seen from Figure S7(a), spherically-symmetric fluctuations of the electrostatic potential have a negligible effect on the free energy of viral particles in the physiologically relevant range of $R_{\rm vir} = 20 - 100$ nm. In the case of larger capsid radii ($R_{\rm vir} > 300$ nm) that correspond to dilute / semidilute DNA conditions, such fluctuations make a more substantial contribution to the free energy of the system, which, however, does not lead to any significant changes in other parameters characterizing the physical state of viral particles, such as the pressure exerted by DNA on the capsid wall, see Figure S7(b). As for nuclei of living cells, spherically-symmetric fluctuations of the nuclear electrostatic potential seem to play an even more minor role in this case, as can be seen from Figures S7(c-d).

Based on the above results, it can be concluded that the mean field approach provides rather accurate description of the physical state of DNA in viral particles and nuclei of living cells at physiologically relevant values of the capsid radius ($R_{\rm vir} = 20 - 100$ nm) and the cell nucleus size ($R_{\rm nucl} = 4 - 10$ $\mu$m), especially taking into account the fact that microenvironment inside nuclei of living cells seem to be far from any critical point under physiological conditions [28]. Thus, the theoretical framework developed in our work can be used to gain useful insights into the molecular processes underlying DNA packaging in viral particles and nuclei of living cells.

Finally, it should be noted that previous theoretical and experimental studies indicate that chemical and biological systems may be a subject to so-called charge inversion phenomenon, which plays an important role in determining their physicochemical behaviour, see references in [29]. However, it should be noted that a vast majority of the conclusions in support of the charge inversion phenomenon in biological systems have been drawn so far mainly based on *in vitro* studies, in which highly artificial experimental conditions were typically applied [29]. As a result, such observations may not be relevant to understanding molecular processes taking place in living cells and their organelles.

In particular, experimental measurements show that physicochemical conditions, which are commonly found in nuclei of living cells, appear to be unfavourable for the dominant contribution of the charge inversion mechanism into chromatin organization at the global nucleus scale. Indeed, multiply negatively charged molecules in nuclei of living cells are mainly represented by nucleosomes and bare DNA linkers connecting them. On the other hand, mobile histones, which diffuse in the nucleoplasm in a form of chaperone-bound protein complexes, form one of the most abundant group of multiply positively charged particles. Yet, from photobleaching experiments it is known that the mobile fraction of histones constitutes only $1\% - 9\%$ of all histones present in the cell nucleus, see ref. [22–24]. In other words, there are much more negatively charged nucleosomes than positively charged mobile histones, – i.e., multiply negatively charged particles are in great excess (by $1 - 2$ orders of magnitude) in comparison to mobile multiply positively charged particles. Furthermore, the absolute value of the net electrical charge carried by each nucleosome is more than an order of magnitude larger than that of chaperone-bound histone dimers due to a partial electrostatic screening of positive histone charges by negatively charged chaperones. As a result, it can be concluded that it is nearly impossible for chromatin to undergo charge inversion on a sufficiently large scale under typical physiological conditions, unless other types of strongly positively charged particles are present in a very large amount in nuclei of living cells.

It is known that behaviour of such highly off-stoichiometric mixtures of polyelectrolytes to a large extent is determined by interaction between the unmatched polyelectrolytes (DNA and nucleosomes in the case of living cells) and monovalent counterions of the corresponding type (i.e., $Na^+$ and $K^+$ ions), which can be very well described by mean field theories based on the Poisson-Boltzmann equation such as Eq. (A37), see comments in ref. [29]. Indeed, by labelling HEK-293 cells with ANG-2 fluorescent sodium indicator, we did not find any exclusion regions of $Na^+$ ions in nuclei of living cells, suggesting that there is no significant charge inversion of chromatin, see Figure S5. On the other hand, the average level of $Na^+$ ions was substantially elevated in nuclei of living cells in comparison to the cell cytoplasm, which was likely a result of $Na^+$ ions interaction with the negative electrostatic potential of the cell nucleus and chromatin. Thus, while the charge inversion phenomenon may potentially play an important role in many chemical and biological systems, it does not seem to be the main factor determining the large-scale organization of chromatin.

In the rest of Appendix sections (Appendices B-H), we derive formulas for the DNA partition function, $Z_\psi$, and its functional derivative, $\frac{\delta \ln Z_\psi}{\delta \psi(r)}$, in an arbitrary potential field, $\psi$, which are needed to complete Eq. (A37).

#### Appendix B: Energy of a mechanically stretched DNA in a potential field, $\psi$

In order to show how to calculate the partition function of chromosomal DNA in an arbitrary potential field, $\psi$, we need first to introduce a discretized semiflexible polymer model of DNA. To this aim, we will start with a simpler scenario of a single DNA molecule (instead of $Q$ chromosomes) interacting with DNA-binding proteins and a potential field, $\psi$.

Following our previous studies [1–3], DNA in the model is represented by a discretized polygonal chain consisting of short segments, which are treated as rigid bodies with a local coordinate system $(\mathbf{x}_j, \mathbf{y}_j, \mathbf{z}_j)$ attached to each of them, see schematic Figure 1(c). Here $j$ is the index enumerating all DNA segments from 1 to $N$, where $N$ is the total number of segments in the polygonal chain representing the DNA molecule. 3D orientation of each of the coordinate systems, and thus each DNA segment, is described by the Euler rotation matrix $\mathbf{R}_j = \mathbf{R}_j(\alpha_j, \beta_j, \gamma_j) = \mathbf{R}_{\alpha_j}\mathbf{R}_{\beta_j}\mathbf{R}_{\gamma_j}$ resulting from the composition of three successive revolutions through Euler angles $\alpha_j$, $\beta_j$ and $\gamma_j$ about the fixed global coordinate frame $(\mathbf{x}_0, \mathbf{y}_0, \mathbf{z}_0)$, see Figure S4(a).

Besides the 3D orientation, DNA segments in the model are additionally characterized by their physical state. Namely, by attaching or dissociating from proteins, DNA segments can switch between protein-bound and bare DNA states. Furthermore, upon formation of nucleoprotein complexes, such as nucleosomes, DNA typically binds to several different parts of a protein. As a result, each nucleoprotein complex usually comprises a number of DNA segments, each bound to a specific site on the protein surface. Thus, to accurately describe nucleoprotein complexes, we need to have a mean to label DNA segments in the model according to their binding sites on the protein surface. This can be done by introducing integer indexes, $k_j$ ($j = 1, ..., N$), to indicate whether a given DNA segment, $j$, is bound to a nucleoprotein complex, and if yes, then to specify the index of the binding site on the protein surface with which this DNA segment interacts. Specifically, assuming that the protein of interest occupies $K$ segments upon binding to DNA, one can assign $K$ DNA binding sites on the protein surface – from 1 (the first DNA binding site on the protein surface) to $K$ (the last DNA binding site on the protein surface). Correspondingly, for each DNA segment bound to a protein we put the value of $k_j$ equal to the index of the respective binding site on the surface of the protein – from $k_j = 1$ (if the DNA segment is bound to the first binding site on the protein surface) to $k_j = K$ (if the DNA segment is bound to the last binding site on the protein surface). For DNA segments, which are not bound to any protein, we simply put $k_j = 0$. Thus, in the presence of DNA-protein interactions, indexes $k_j$ ($j = 1, ..., N$) take integer values in the range from 0 to $K$, with $k_j = 0$ representing bare DNA segments, and $k_j = 1, ..., K$ corresponding to protein-bound DNA segments. In the latter case, for a given DNA segment, $j$, parameter $k_j$ equals to the index of the DNA binding site on the protein surface to which this DNA segment is bound. As an example, see schematic

Figure S4(b) for the case of $K = 12$.

Knowing the orientations of DNA segments, $(\mathbf{R}_1, ..., \mathbf{R}_N)$, their physical states, $(k_1, ..., k_N)$, and the position vector of the starting end of DNA, $\mathbf{r}_0$, one then can find the total conformational energy of DNA, $E_{\text{tot}}$, which in the general case can be written as a sum of the following energy terms:

$$E_{\text{tot}}(k_1, ..., k_N, \mathbf{R}_1, ..., \mathbf{R}_N, \mathbf{r}_0) = E_{\text{DNA}} + E_{\text{protein}} + E_{\text{conf}} + E_\psi + E_f \tag{B1}$$

Here $E_{\text{DNA}}$ term includes the bending and twisting elastic deformation energies of bare DNA segments; whereas, $E_{\text{protein}}$ is the energy term associated with formation of nucleoprotein complexes on DNA. $E_{\text{conf}}$ is the energy of DNA confinement in the cell nucleus. $E_\psi$ is the potential energy describing DNA interaction with field $\psi$. Finally, as in the main text we consider several scenarios in which a small part of chromosomal DNA is stretched by mechanical force, we added into the above equation the last energy term, $E_f$, which represents the potential energy of DNA associated with the stretching force, $f$. Without loss of generality, it can be assumed that this force point in the direction of $\mathbf{z}_0$-axis of the global coordinate system.

To provide more detailed insights into the above energy terms, we will consider first a simple abstract case of DNA interacting with proteins that occupy $K = 3$ DNA segments upon binding to DNA. After finding formulas for the total energy and the partition function of DNA in such a hypothetical case, we will then generalize the obtained results to proteins that have an arbitrary large binding site size on DNA, aiming at description of DNA interaction with histone octamers, which are involved in formation of nucleosomes. Thus, starting from here to the end of Appendix C, indexes $k_j$ will be assumed to take integer values from 0 to 3, with indexes $k_j = 0$ corresponding to bare DNA segments, and indexes $k_j = 1$, 2 and 3 – to DNA segments bound to proteins.

Let's now consider one by one energy terms from Eq. (B1).

The first term from Eq. (B1) describes elastic deformations of bare DNA segments, and, as has been discussed in ref. [1–3], it simply equals to:

$$E_{\text{DNA}} = \sum_{j=1}^{N-1} \delta_{k_j 0} \delta_{k_{j+1} 0} E_{\text{bare}}(\mathbf{R}_j, \mathbf{R}_{j+1}) \tag{B2}$$

Here $\delta_{k_j n}$ is the Kronecker delta ($\delta_{k_j n} = 1$ if $k_j = n$ and $\delta_{k_j n} = 0$, otherwise). $E_{\text{bare}}(\mathbf{R}_j, \mathbf{R}_{j+1})$ is the local elastic deformation energy of DNA corresponding to the vertex joining the $j^{\text{th}}$ and $(j+1)^{\text{th}}$ segments of the polygonal chain representing the DNA, which comprises the bending and twisting energies of bare DNA:

$$E_{\text{bare}}(\mathbf{R}_j, \mathbf{R}_{j+1}) = \frac{a}{2\beta}(\mathbf{R}_j \mathbf{z}_0 - \mathbf{R}_{j+1} \mathbf{z}_0)^2 + \frac{c}{2\beta}[\Delta\varphi_j(\mathbf{R}_j, \mathbf{R}_{j+1})]^2 \tag{B3}$$

Where $\Delta\varphi_j(\mathbf{R}_j, \mathbf{R}_{j+1})$ is the local DNA twist angle between the $j^{\text{th}}$ and $(j+1)^{\text{th}}$ DNA segments. $a = A/b$ and $c = C/b$ are dimensionless parameters designating the bending and twisting elasticities of bare DNA segments in the semiflexible polymer chain model of DNA, where $A$ and $C$ are the bending and twisting persistence lengths of B-DNA (Table I). Here $b$ is the length of bare DNA segments in the polygonal chain representing the DNA molecule. Calculations show that the exact value of $b$ does not have any significant effect on the final results as soon as $b$ is selected to be much smaller than the bending persistence length of DNA (i.e., $b \ll A$), see Figure S8(a). For this reason, in all our calculations, we used the value of $b$ corresponding to a single B-DNA turn: $b = 3.4$ nm.

Direct comparison of Eq. (B3) to Eq. (A1) shows that the above formula is just a discretized version of the first two integrals in Eq. (A1), which describe elastic bending and twisting deformation energies of bare DNA.

Let's now have a look at the next energy term from Eq. (B1), $E_{\text{protein}}$.

To derive a mathematical expression for it, we need to provide several additional details regarding the treatment of nucleoprotein complexes in our study. Specifically, in this study, we utilize a coarse-grained representation of nucleoprotein complexes, which has been originally introduced for DNA-wrapping proteins in ref. [3]. According to this approximation, all DNA segments constrained inside each of the nucleoprotein complexes are replaced by a straight line connecting the entry and exit points of DNA in the corresponding complex. This line is then subdivided into smaller intervals, whose total number, $K$, equals to the total number of DNA segments bound to a protein in a single nucleoprotein complex, see schematic Figure S4(c) and ref. [3] for more details. Thus, the length of each subinterval is $b_1 = ... = b_K = r_{\rm pr}/K$, where $r_{\rm pr}$ is the average distance between the entry and exit points of DNA in nucleoprotein complexes. As will be shown in Appendices G-K, by replacing protein-bound DNA segments with such subintervals, it is possible to considerably simplify calculations of the DNA partition function as all of the new segments will have the same 3D orientation. For example, in the case of $j$, $j+1$, ..., $j+K-1$ DNA segments being bound to a nucleoprotein complex, the above replacement leads to the following simple constraint on orientations of the protein-bound DNA segments: $\mathbf{R}_j = \mathbf{R}_{j+1} = ... = \mathbf{R}_{j+K-1} = \mathbf{R}_{\rm pr,j}$, see Figure S4(c). Here $\mathbf{R}_{\rm pr,j}$ is a Euler rotation matrix describing the orientation of the line connecting the entry and exit points of DNA in the considered nucleoprotein complex.

As for DNA segments immediately upstream and downstream of the entry and exit points of DNA, it is clear that they must have specific orientations with respect to the main body of a nucleoprotein complex, corresponding to a relaxed conformation of DNA. Namely, if $\mathbf{R}_{j-1}$ and $\mathbf{R}_{j+K}$ are Euler rotation matrices describing the orientations of the DNA segments sitting next to the entry and exit points of a nucleoprotein complex, which occupies $j$, $j+1$, ..., $j+K-1$ DNA segments, then in the relaxed conformation their orientations must satisfy the following two general equations: $\mathbf{R}^{\rm (rel)}_{j-1}\mathbf{A}_{\rm in} = \mathbf{R}_j = \mathbf{R}_{\rm pr,j}$ and $\mathbf{R}^{\rm (rel)}_{j+K} = \mathbf{R}_{j+K-1}\mathbf{A}_{\rm out} = \mathbf{R}_{\rm pr,j}\mathbf{A}_{\rm out}$, where $\mathbf{A}_{\rm in}$ and $\mathbf{A}_{\rm out}$ are some rotation matrices that define the relaxed orientations of the entry and exit DNA segments relative to the core part of the nucleoprotein complex.

To impose the above constraints in the partition function calculations, we utilize in this study previously developed approach based on Dirac $\delta$-functions defined on SO(3) group of Euler rotation matrices [3]. For example, in the case of a nucleoprotein complex occupying $j$, $j+1$, ..., $j+K-1$ DNA segments, the constraint $\mathbf{R}_j = \mathbf{R}_{j+1} = ... = \mathbf{R}_{j+K-1}$ applied to the DNA segments bound to the protein can be enforced in DNA partition function calculations by using the following Dirac $\delta$-functions: $\delta(\mathbf{R}_j - \mathbf{R}_{j+1})$, $\delta(\mathbf{R}_{j+1} - \mathbf{R}_{j+2})$, ..., $\delta(\mathbf{R}_{j+K-2} - \mathbf{R}_{j+K-1})$. However, for the sake of formulas simplicity, instead of inserting such $\delta$-functions directly under the integral sign of Eq. (C1) defining the DNA partition function, it is more convenient to add them to $E_{\rm protein}$ energy term in a form of Dirac $\delta$-function logarithms, such as $\ln[\delta(\mathbf{R} - \mathbf{R}')]$. The latter is defined as a generalized function, which after exponentiation results in the Dirac $\delta$-function: $\exp[\ln\delta(\mathbf{R} - \mathbf{R}')] = \delta(\mathbf{R} - \mathbf{R}')$. Here $\mathbf{R}$ and $\mathbf{R}'$ are Euler rotation matrices describing orientations of neighbouring DNA segments in a nucleoprotein complex.

As such generalized functions have to be added to $E_{\rm protein}$ for each of the DNA segment bound to a protein, it is clear that $E_{\rm protein}$ energy term then must contain the following sum: $-\frac{1}{\beta}\sum_{j=1}^{N-1}\sum_{n=1}^{K}\delta_{k_j n}\ln\left[\delta(\mathbf{R}_j - \mathbf{R}_{j+1})\right]$.

It should be noted that such representation of nucleoprotein complexes has a small drawback – by using Dirac $\delta$-functions to impose predefined relative orientations on protein-bound DNA segments, we implicitly offset the free energies of the corresponding nucleoprotein complexes by a constant term, $\mu_{\rm off}$, resulting from the change in the entropy contribution by protein-bound DNA segments. Nevertheless, the exact value of $\mu_{\rm off}$ can be easily found using transfer-matrix calculations and subtracted from the energy of each nucleoprotein complex as described in Appendix F of ref. [3].

In addition to the above sum and $\mu_{\rm off}$ correction, $E_{\rm protein}$ energy term must also contain elastic deformation energies, $E_{\rm in}$ and $E_{\rm out}$, of the DNA segments sitting next to the entry ("in") and exit ("out") points of nucleoprotein complexes described in this study by the following equations:

$$
\begin{aligned}
E_{\rm in}(\mathbf{R}_{\rm in}, \mathbf{R}_{\rm first}) &= \frac{a_{\rm pr}}{2\beta}\left(\mathbf{R}_{\rm in}\mathbf{A}_{\rm in}\mathbf{z}_0 - \mathbf{R}_{\rm first}\mathbf{z}_0\right)^2 + \frac{c_{\rm pr}}{2\beta}\left[\Delta\varphi(\mathbf{R}_{\rm in}\mathbf{A}_{\rm in}, \mathbf{R}_{\rm first})\right]^2 \\
E_{\rm out}(\mathbf{R}_{\rm last}, \mathbf{R}_{\rm out}) &= \frac{a_{\rm pr}}{2\beta}\left(\mathbf{R}_{\rm last}\mathbf{A}_{\rm out}\mathbf{z}_0 - \mathbf{R}_{\rm out}\mathbf{z}_0\right)^2 + \frac{c_{\rm pr}}{2\beta}\left[\Delta\varphi(\mathbf{R}_{\rm last}\mathbf{A}_{\rm out}, \mathbf{R}_{\rm out})\right]^2
\end{aligned}
\tag{B4}
$$

Where $a_{\rm pr}$ and $c_{\rm pr}$ are dimensionless bending and twisting elasticities of the entry and exit DNA segments in nucleoprotein complexes. $\mathbf{R}_{\rm in}$ and $\mathbf{R}_{\rm out}$ are Euler rotation matrices denoting orientations of the entry and exit DNA segments; whereas, $\mathbf{R}_{\rm first}$ and $\mathbf{R}_{\rm last}$ matrices designate the orientations of the first and the last DNA segments inside a nucleoprotein complex, see Figures S4(b-c) for details. $\mathbf{A}_{\rm in}$ and $\mathbf{A}_{\rm out}$ are Euler rotation matrices that determine the relaxed orientations of the entry and exit DNA segments relative to the core part of a nucleoprotein complex. Finally,

$\Delta\varphi(\mathbf{R}_{\rm in}\mathbf{A}_{\rm in}, \mathbf{R}_{\rm first})$ and $\Delta\varphi(\mathbf{R}_{\rm last}\mathbf{A}_{\rm out}, \mathbf{R}_{\rm out})$ are the twist angles of the entry and exit DNA segments with respect to their relaxed orientations.

Thus, for the earlier discussed case of a protein with $K = 3$ binding site size, which is attached to DNA segments with indexes $j$, $j+1$ and $j+2$, we have: $\mathbf{R}_{\rm in} = \mathbf{R}_{j-1}$, $\mathbf{R}_{\rm first} = \mathbf{R}_j$, $\mathbf{R}_{\rm last} = \mathbf{R}_{j+2}$ and $\mathbf{R}_{\rm out} = \mathbf{R}_{j+3}$. Furthermore, as can be easily seen from Eq. (B4), the relaxed conformation of such a nucleoprotein complex is described by the following set of equations:

$$\begin{aligned} &E_{\rm in} = E_{\rm out} = 0, \quad \mathbf{R}^{\rm (rel)}_{j-1}\mathbf{A}_{\rm in}\mathbf{z}_0 = \mathbf{R}_j\mathbf{z}_0, \quad \mathbf{R}_{j+2}\mathbf{A}_{\rm out}\mathbf{z}_0 = \mathbf{R}^{\rm (rel)}_{j+3}\mathbf{z}_0, \\ &\Delta\varphi\Big(\mathbf{R}^{\rm (rel)}_{j-1}\mathbf{A}_{\rm in}, \mathbf{R}_j\Big) = 0 \quad \text{and} \quad \Delta\varphi\Big(\mathbf{R}_{j+2}\mathbf{A}_{\rm out}, \mathbf{R}^{\rm (rel)}_{j+3}\Big) = 0, \end{aligned} \tag{B5}$$

from which it can be immediately found that: $\mathbf{R}^{\rm (rel)}_{j-1}\mathbf{A}_{\rm in} = \mathbf{R}_j$ and $\mathbf{R}_{j+2}\mathbf{A}_{\rm out} = \mathbf{R}^{\rm (rel)}_{j+3}$ – i.e., the relaxed orientations of the entry and exit DNA segments with respect to the nucleoprotein complex are indeed described by rotation matrices $\mathbf{A}_{\rm in}$ and $\mathbf{A}_{\rm out}$.

Next, we would like to note that for the sake of formulas simplicity in this study we considered only the case when proteins form complete nucleoprotein complexes upon binding to DNA, never assembling into partially unfolded structures. I.e., all the binding sites on the surface of a protein are assumed to fully engaged with DNA during formation of a nucleoprotein complex. To this aim, we set the energy of all of the DNA-protein conformations that contain one or more partially unfolded nucleoprotein complexes equal to $+\infty$, preventing any contribution of improperly formed nucleoprotein complexes to the DNA partition function.

The only question then is how to distinguish properly formed nucleoprotein complexes from partially unfolded in model calculations? To this aim, we use the following approach. First, it is clear that in the case of properly formed nucleoprotein complexes each pair of indexes, $(k_j, k_{j+1})$, of neighbouring DNA segments can have only one of the following values: $(0,0)$, $(0,1)$, $(1,2)$, $(2,3)$, $(3,0)$, where we still consider the case of proteins, which occupy $K = 3$ DNA segments upon binding to DNA. All other pairs of $(k_j, k_{j+1})$, such as $(1,1)$, $(2,1)$, $(0,2)$, etc., correspond either to a situation when there is one or more partially unfolded nucleoprotein complexes formed on DNA, or there are two complexes bound to DNA next to each other, i.e., $(k_j, k_{j+1}) = (3,1)$. In this study, we exclude the latter pair of indexes from the set of properly formed complexes as experimental data show that nucleoprotein complexes, such as nucleosomes, are usually separated by bare DNA linkers of $\sim 10-60$ bp length [22, 30, 31].

Anyway, it is clear that in order to set the energies of improper DNA-protein conformations to infinity, all we need to do is to add the following sum $\sum_{j=1}^{N-1}\sum_{(n,m)\notin H}\delta_{k_j n}\delta_{k_{j+1} m}\times\infty$ to $E_{\rm protein}$ energy term, where $H = \{(0,0),(0,1),(1,2),(2,3),(3,0)\}$ is the set of correct combinations of the neighbouring DNA segments' states corresponding to properly formed nucleoprotein complexes. Here we use a typical mathematical convention that $0\times\infty = 0$.

By collecting all of the above energy contributions and adding to them the binding free energy of proteins to DNA, $\mu_{\rm pr}$, we finally obtain the following formula for $E_{\rm protein}$:

$$\begin{aligned} E_{\rm protein} = &-(\mu_{\rm pr} + \mu_{\rm off})\sum_{j=1}^{N-1}\delta_{k_j 0}\delta_{k_{j+1} 1} + \sum_{j=1}^{N-1}\delta_{k_j 0}\delta_{k_{j+1} 1}E_{\rm in}(\mathbf{R}_j, \mathbf{R}_{j+1}) + \sum_{j=1}^{N-1}\delta_{k_j K}\delta_{k_{j+1} 0}E_{\rm out}(\mathbf{R}_j, \mathbf{R}_{j+1}) \\ &-\frac{1}{\beta}\sum_{j=1}^{N-1}\sum_{n=1}^{K-1}\delta_{k_j n}\ln\delta(\mathbf{R}_j - \mathbf{R}_{j+1}) + \sum_{j=1}^{N-1}\sum_{(n,m)\notin H}\delta_{k_j n}\delta_{k_{j+1} m}\times\infty + (1-\delta_{k_1 0})\times\infty + (1-\delta_{k_N 0})\times\infty \end{aligned} \tag{B6}$$

Here we also included the last to terms, $(1-\delta_{k_1 0})\times\infty$ and $(1-\delta_{k_N 0})\times\infty$, to prohibit formation of partially unfolded nucleoprotein complexes on the end segments of DNA.

It should be noted that the free energy associated with proteins binding to DNA, $\mu_{\rm pr}$, not only comprises the binding energy, $\mu$, of a protein to DNA, but also change in the free energies of other molecules, which switch their state upon formation of a nucleoprotein complex. For example, in the case of nucleosomes, which form from histone dimers delivered by histone-binding chaperones, we must add to $\mu$ logarithms of all of the prefactors from Eq. (A19) under the sum sign related to histone-binding chaperones:

$$\mu_{\rm pr} = \mu - 2k_{\rm B}T\ln\left(\frac{Z^{\psi}_{{\rm c}_1{\rm b}}}{n^0_{{\rm c}_1{\rm b}}}\right) - 2k_{\rm B}T\ln\left(\frac{Z^{\psi}_{{\rm c}_2{\rm b}}}{n^0_{{\rm c}_2{\rm b}}}\right) + 2k_{\rm B}T\ln\left(\frac{Z^{\psi}_{{\rm c}_1{\rm u}}}{n^0_{{\rm c}_1{\rm u}}}\right) + 2k_{\rm B}T\ln\left(\frac{Z^{\psi}_{{\rm c}_2{\rm u}}}{n^0_{{\rm c}_2{\rm u}}}\right) \tag{B7}$$

In the absence of the electrostatic field ($\psi = 0$), the above formula turns into a simple difference between the chemical potentials of the initial reactants and final products: $\mu_{\rm pr} = \mu^0_{\rm pr} + 2k_{\rm B}T\ln c^0_{{\rm c}_1{\rm b}} + 2k_{\rm B}T\ln c^0_{{\rm c}_2{\rm b}} - 2k_{\rm B}T\ln c^0_{{\rm c}_1{\rm u}} - 2k_{\rm B}T\ln c^0_{{\rm c}_2{\rm u}}$, where $c^0_{{\rm c}_1{\rm u}}$, $c^0_{{\rm c}_2{\rm u}}$, $c^0_{{\rm c}_1{\rm b}}$ and $c^0_{{\rm c}_2{\rm b}}$ are the average concentrations of unloaded, H2A·H2B-bound and H3·H4-bound chaperones $c_1$ and $c_2$, respectively. Here $\mu^0_{\rm pr} = \mu - 2k_{\rm B}T\ln(Z^{\rm in}_{{\rm c}_1{\rm b}}/\Lambda^3_{{\rm c}_1{\rm b}}) - 2k_{\rm B}T\ln(Z^{\rm in}_{{\rm c}_2{\rm b}}/\Lambda^3_{{\rm c}_2{\rm b}}) + 2k_{\rm B}T\ln(Z^{\rm in}_{{\rm c}_1{\rm u}}/\Lambda^3_{{\rm c}_1{\rm u}}) + 2k_{\rm B}T\ln(Z^{\rm in}_{{\rm c}_2{\rm u}}/\Lambda^3_{{\rm c}_2{\rm u}})$ is the standard Gibbs free energy of the chemical reaction describing nucleosome assembly on DNA, which can be derived from the definition of $Z^{\psi}_{{\rm c}_1{\rm b}}$, $Z^{\psi}_{{\rm c}_2{\rm b}}$, $Z^{\psi}_{{\rm c}_1{\rm u}}$ and $Z^{\psi}_{{\rm c}_2{\rm u}}$ partition functions [Eq. (A15)]. Experimental measurements show that in the case of Nap1, which is one of the major histone-binding chaperones in living cells [17–19], the average value of $\mu^0_{\rm pr}$ is of the order of $\sim 7\ k_{\rm B}T$ [32, 33].

By using the above definition of the standard Gibbs free energy, $\mu^0_{\rm pr}$, and Eq. (A15), it is convenient to rewrite Eq. (B7) in a slightly different form:

$$\mu_{\rm pr} = \mu^0_{\rm pr} + 2k_{\rm B}T\ln\theta_{{\rm c}_1} + 2k_{\rm B}T\ln\theta_{{\rm c}_2} + 2k_{\rm B}T\ln\left(\frac{\int_{V^{\rm osm}_{\rm cell}} e^{-\beta q_{{\rm c}_1{\rm u}}\psi(\mathbf{r})}\,d\mathbf{r}\cdot\int_{V^{\rm osm}_{\rm cell}} e^{-\beta q_{{\rm c}_2{\rm u}}\psi(\mathbf{r})}\,d\mathbf{r}}{\int_{V^{\rm osm}_{\rm cell}} e^{-\beta q_{{\rm c}_1{\rm b}}\psi(\mathbf{r})}\,d\mathbf{r}\cdot\int_{V^{\rm osm}_{\rm cell}} e^{-\beta q_{{\rm c}_2{\rm b}}\psi(\mathbf{r})}\,d\mathbf{r}}\right) \tag{B8}$$

Where $\theta_{{\rm c}_1} = n^0_{{\rm c}_1{\rm b}}/n^0_{{\rm c}_1{\rm u}} = c^0_{{\rm c}_1{\rm b}}/c^0_{{\rm c}_1{\rm u}}$ and $\theta_{{\rm c}_2} = n^0_{{\rm c}_2{\rm b}}/n^0_{{\rm c}_2{\rm u}} = c^0_{{\rm c}_2{\rm b}}/c^0_{{\rm c}_2{\rm u}}$ are occupancy ratios of histone-binding chaperones, $c_1$ and $c_2$. $q_{{\rm c}_1{\rm u}}$, $q_{{\rm c}_2{\rm u}}$, $q_{{\rm c}_1{\rm b}}$ and $q_{{\rm c}_2{\rm b}}$ are the total electric charges of unloaded as well as histone-bound chaperones $c_1$ and $c_2$, respectively.

It should be noted that the total binding free energy of histone octamers to DNA, $\mu^{\rm tot}_{\rm pr}$, which is defined by Eq. (9) in the main text, differ from $\mu_{\rm pr}$ only by the value of the average potential energy of a histone octamer in the electrostatic field of the cell nucleus: $\mu^{\rm tot}_{\rm pr} \approx \mu_{\rm pr} - q_{\rm oct}\langle\psi\rangle$. Here $q_{\rm oct}$ is the electrical charge of a histone octamer and $\langle\psi\rangle$ is the average value of the nuclear electrostatic potential. The main reason why this potential energy is absent in $\mu_{\rm pr}$ is that in the model calculations it is taken into account in a separate term, $E_\psi$, see Eq. (B15) below.

Anyway, since Eq. (B4), Eq. (B6) and Eq. (B8) completely define $E_{\rm protein}$ energy from Eq. (B1), we then can move to description of the next term, $E_{\rm conf}$, corresponding to the confinement energy of chromosomal DNA in the cell nucleus.

In the case of the polymer chain model of DNA, this term can be represented by a disretized version of the third integral in Eq. (A1):

$$E_{\rm conf} = \frac{1}{L}\sum_{j=1}^{N} b_{k_j} U_{R_{\rm nucl}}(\mathbf{r}_j) \tag{B9}$$

Where $L$ is the contour length of DNA and $b_{k_j}$ is the length of the $j^{\rm th}$ DNA segment in state $k_j$: $b_{k_j} = b_0 = b$ if $k_j = 0$ (i.e., the $j^{\rm th}$ DNA segment is not bound to a protein) and $b_{k_j} = r_{\rm pr}/K$, otherwise. In fact, the exact values of $b_{k_j}$ parameters do not make any difference in Eq. (B9) as soon as they are positive ($b_{k_j} > 0$) because the confinement potential, $U_{R_{\rm nucl}}$, takes only two values: 0 or $+\infty$. On the other hand, $b_{k_j}$ lengths have a strong effect on the remaining two energy terms in Eq. (B1), $E_\psi$ and $E_f$, as they define the position vectors of the DNA segments, $(\mathbf{r}_1, ..., \mathbf{r}_N)$, with respect to the global coordinate frame:

$$\begin{cases} \mathbf{r}_1 &= \mathbf{r}_0 + b_{k_1}\mathbf{R}_1\mathbf{z}_0 \\ \mathbf{r}_2 &= \mathbf{r}_0 + \sum_{j=1}^{j=2} b_{k_j}\mathbf{R}_j\mathbf{z}_0 \\ \vdots & \\ \mathbf{r}_N &= \mathbf{r}_0 + \sum_{j=1}^{j=N} b_{k_j}\mathbf{R}_j\mathbf{z}_0 \end{cases} \tag{B10}$$

Here $\mathbf{r}_0$ is the position vector of the starting point of the DNA.

While Eq. (B9) describes the confinement energy of DNA in the general case, we are going to use a slightly simpler version of it to make easier later derivation of a formula for the DNA partition function. Specifically, we will assume in our model that only those joints of the polygonal chain representing the DNA interact with the NE which are followed by a bare DNA segment. In other words, protein-bound DNA segments feel the physical presence of the NE only through flanking bare DNA segments, see schematic Figure S9. This approximation introduces a negligible error into the partition function calculations as the size of the cell nucleus is by several orders of magnitude larger than the size of individual nucleoprotein complexes. On the other hand, such simplification leads to a much faster computational algorithm for the DNA partition function, which will be constructed in Appendices C-I, justifying such an approximation.

Thus, instead of Eq. (B9), we are going to use the following formula for $E_{\mathrm{conf}}$ energy:

$$E_{\mathrm{conf}} = \frac{1}{L}\sum_{j=1}^{N-1} b_{k_j} U_{R_{\mathrm{nucl}}}(\mathbf{r}_j)\, \delta_{k_{j+1}0} + \frac{1}{L} b_{k_N} U_{R_{\mathrm{nucl}}}(\mathbf{r}_N) \tag{B11}$$

For the sake of simplicity, it is convenient to rewrite Eq. (B11) as:

$$E_{\mathrm{conf}} = \sum_{j=1}^{N} \sigma_{k_j k_{j+1}} U_{R_{\mathrm{nucl}}}(\mathbf{r}_j) \tag{B12}$$

Where $\sigma_{k_j k_{j+1}} = b_{k_j}/L$ in the case when the $j^{\mathrm{th}}$ DNA segment is followed by one being in the bare DNA state (i.e., $k_{j+1} = 0$), or when $j = N$, in all other cases $\sigma_{k_j k_{j+1}} = 0$.

As for the last two terms from Eq. (B1), $E_\psi$ and $E_f$, it is straightforward to obtain a mathematical expression for them based on the continuous polymer model of DNA, from which we have:

$$E_\psi + E_f = \int_0^L \rho_{\mathrm{d}}(s)\psi(\mathbf{r}_{\mathrm{d}}(s))\mathrm{d}s - \int_0^L \big[\mathbf{f}\cdot\mathbf{t}(s)\big]\mathrm{d}s \tag{B13}$$

Here $\mathbf{r}_{\mathrm{d}}(s)$ is the position vector of a point on the DNA contour, which corresponds to the arc length $s$. $\psi(\mathbf{r}_{\mathrm{d}}(s))$ is the value of the electrostatic potential, $\psi$, at the point described by the position vector $\mathbf{r}_{\mathrm{d}}(s)$. $\rho_{\mathrm{d}}(s)$ is the net linear charge density of protein-covered DNA that determines the strength of DNA interaction with the potential $\psi$. $\mathbf{t}(s)$ is the unit tangent vector to the DNA contour at the point corresponding to the arc length $s$. $\mathbf{f} = f\mathbf{z}_0$ is the stretching force applied to the DNA along $\mathbf{z}_0$-axis of the global coordinate system, where $f = \|\mathbf{f}\|$ is the force magnitude.

In the case of the discretized polymer chain model of DNA, the above expression for the potential energies, $E_\psi$ and $E_f$, turns into:

$$E_\psi + E_f = \sum_{j=1}^{N} b_{k_j}\rho_{k_j}\psi(\mathbf{r}_j) - f\sum_{j=1}^{N} b_{k_j}\big[\mathbf{z}_0\cdot\mathbf{R}_j\mathbf{z}_0\big] = \sum_{j=1}^{N} q_{k_j}\psi(\mathbf{r}_j) - f\sum_{j=1}^{N} b_{k_j}\big[\mathbf{z}_0\cdot\mathbf{R}_j\mathbf{z}_0\big] \tag{B14}$$

Where $\rho_{k_j}$ is the linear charge density of the $j^{\mathrm{th}}$ DNA segment in the corresponding state, $k_j$. For bare DNA segments ($k_j = 0$): $\rho_0 = \rho_{\mathrm{d}} = -2q_{\mathrm{e}}/\,0.33$ nm – i.e., $2e^-$ per single DNA base-pair; whereas, in the case of protein-bound DNA segments ($k_j > 0$) this value will be generally closer to zero due to the electrostatic screening effect created by positively charged proteins attached to the DNA. $q_{k_j} = b_{k_j}\rho_{k_j}$ is the electrical charge of the $j^{\mathrm{th}}$ DNA segment in state $k_j$.

Previous theoretical studies as well as our own calculations show that the mean electrostatic potential inside viral particles created by packed DNA and ions diffusing in solution is usually described by a smooth function, see ref. [34, 35] and Figure 2(d). Similarly, it is natural to expected that the nuclear electrostatic potential in living cells, $\psi$, also has a smooth shape, which is indeed the case as can be seen from Figure 4(a) and Figure S2. As the characteristic length of $\psi$ field variations is much larger than the size of individual nucleoprotein complexes, it can be concluded that the strength of interaction between the field and nucleoprotein complexes formed on DNA is practically insensitive to the exact distribution of electrical charges within the complexes. Thus, for the sake of faster model calculations, it can be assumed that the electrical charge of each nucleoprotein complex is concentrated at a single point on the polymer chain representing the DNA, as schematically shown in Figure S9.

In such a case, the electrical charge of each DNA segment will be a function of its own physical state as well as the state of the downstream DNA segment. I.e., electrical charge of the $j^{\text{th}}$ DNA segment, $q_{k_j k_{j+1}}$, will generally depend on physical states of both $j^{\text{th}}$ and $(j+1)^{\text{th}}$ DNA segments, $k_j$ and $k_{j+1}$. For example, in the case of nucleosomes: $q_{00} = b\rho_{\text{d}} = -2bq_{\text{e}}/\,0.34$ nm; $q_{01} = q_{12} = ... = q_{K-1,K} = 0$, and $q_{K0} = b\rho_{\text{d}} + q_{\text{nucl}}$, see Figure S9. Here $q_{\text{nucl}} = q_{\text{oct}} - 2\cdot 147\, q_{\text{e}} = -145.2\, q_{\text{e}}$ is the total electrical charge of a nucleosome (i.e., the net electrical charge of a histone octamer, $q_{\text{oct}}$, and DNA wrapped around it). As a result, we obtain the following formula for $E_\psi$ and $E_f$ energies:

$$E_\psi + E_f = \sum_{j=1}^{N} q_{k_j k_{j+1}} \psi(\mathbf{r}_j) - f \sum_{j=1}^{N} b_{k_j} \left[\mathbf{z}_0 \cdot \mathbf{R}_j \mathbf{z}_0\right] \tag{B15}$$

Substituting Eq. (B2), (B6), (B12) and (B15) into Eq. (B1), it is not hard to see that the total energy of DNA can be represented as a sum of local energy contributions, $E_{k_j k_{j+1}}$, made by neighbouring DNA segments whose states are defined by indexes $k_j$ and $k_{j+1}$, respectively:

$$\begin{aligned} E_{\text{tot}}(k_1...k_N, \mathbf{R}_1...\mathbf{R}_N, \mathbf{r}_0) = \sum_{j=1}^{N-1} E_{k_j k_{j+1}}(\mathbf{R}_j, \mathbf{R}_{j+1}) + \sum_{j=1}^{N} \hat{\psi}_{k_j k_{j+1}}(\mathbf{r}_j) - \\ - bf\left[\mathbf{z}_0 \cdot \mathbf{R}_N \mathbf{z}_0\right] + (1-\delta_{k_1 0}) \times \infty + (1-\delta_{k_N 0}) \times \infty \end{aligned} \tag{B16}$$

Where $\hat{\psi}_{k_j k_{j+1}}(\mathbf{r}_j) = \sigma_{k_j k_{j+1}} U_{R_{\text{nucl}}}(\mathbf{r}_j) + q_{k_j k_{j+1}} \psi(\mathbf{r}_j)$ is the net action of $U_{R_{\text{nucl}}}$ and $\psi$ fields on the $j^{\text{th}}$ DNA segment. For $j = N$ case, we put $\hat{\psi}_{k_j k_{j+1}} = \hat{\psi}_{00}$.

From Eq. (B1)-Eq. (B16), it is not hard to find a mathematical expression for $E_{k_j k_{j+1}}(\mathbf{R}_j, \mathbf{R}_{j+1})$ energy terms, which in the case of nucleoprotein complexes occupying three DNA segments ($K = 3$ scenario) takes the following form:

$$\begin{cases} E_{00}(\mathbf{R}_j, \mathbf{R}_{j+1}) = \frac{a}{2\beta}\left(\mathbf{R}_j \mathbf{z}_0 - \mathbf{R}_{j+1}\mathbf{z}_0\right)^2 + \frac{c}{2\beta}\left[\Delta\varphi_j(\mathbf{R}_j, \mathbf{R}_{j+1})\right]^2 - bf\left[\mathbf{z}_0 \cdot \mathbf{R}_j \mathbf{z}_0\right] \\ E_{01}(\mathbf{R}_j, \mathbf{R}_{j+1}) = -\mu_{\text{pr}} - \mu_{\text{off}} + \frac{a_{\text{pr}}}{2\beta}\left(\mathbf{R}_j \mathbf{A}_{\text{in}} \mathbf{z}_0 - \mathbf{R}_{j+1}\mathbf{z}_0\right)^2 + \frac{c_{\text{pr}}}{2\beta}\left[\Delta\varphi_j\left(\mathbf{R}_j \mathbf{A}_{\text{in}}, \mathbf{R}_{j+1}\right)\right]^2 - bf\left[\mathbf{z}_0 \cdot \mathbf{R}_j \mathbf{z}_0\right] \\ E_{12}(\mathbf{R}_j, \mathbf{R}_{j+1}) = -\frac{1}{\beta}\ln\delta(\mathbf{R}_j - \mathbf{R}_{j+1}) - \frac{r_{\text{pr}}}{K} f\left[\mathbf{z}_0 \cdot \mathbf{R}_j \mathbf{z}_0\right] \\ E_{23}(\mathbf{R}_j, \mathbf{R}_{j+1}) = -\frac{1}{\beta}\ln\delta(\mathbf{R}_j - \mathbf{R}_{j+1}) - \frac{r_{\text{pr}}}{K} f\left[\mathbf{z}_0 \cdot \mathbf{R}_j \mathbf{z}_0\right] \\ E_{30}(\mathbf{R}_j, \mathbf{R}_{j+1}) = \frac{a_{\text{pr}}}{2\beta}\left(\mathbf{R}_j \mathbf{A}_{\text{out}} \mathbf{z}_0 - \mathbf{R}_{j+1}\mathbf{z}_0\right)^2 + \frac{c_{\text{pr}}}{2\beta}\left[\Delta\varphi_j\left(\mathbf{R}_j \mathbf{A}_{\text{out}}, \mathbf{R}_{j+1}\right)\right]^2 - \frac{r_{\text{pr}}}{K} f\left[\mathbf{z}_0 \cdot \mathbf{R}_j \mathbf{z}_0\right] \\ E_{02} = E_{03} = E_{10} = E_{11} = E_{13} = E_{20} = E_{21} = E_{22} = E_{31} = E_{32} = E_{33} = +\infty \end{cases} \tag{B17}$$

Despite the daunting look of the above formula, it can be seen that most of $E_{k_j k_{j+1}}$ energy terms are very similar to each other, only slightly varying from one line of the equation to another. This makes it easy to obtain formulas for all of the elements of the DNA transfer-matrix, $\mathbf{L}$, described in the next Appendix section, as soon as we know a mathematical expression only for one of them.

However, before proceeding to description of the transfer-matrix formalism, we need to make a couple of important notes.

First, as was mentioned in ref. [3], all nucleoprotein complexes formed on DNA have a certain orientational freedom – upon binding to DNA, proteins may form nucleoprotein complexes on either side of the DNA duplex due to its double-stranded helical structure. This introduces a new degree of freedom into the model calculations, which we have not considered so far.

From a physical point of view, such positional freedom means that in Eq. (B17) we need to replace $\mathbf{R}_j$ rotation matrix in the formula for $E_{01}$ term with $\mathbf{R}_j\mathbf{R}_{\eta_{\rm in}}$ matrix product, where $\mathbf{R}_{\eta_{\rm in}} = \mathbf{R}_{\eta_{\rm in}}(\eta_{\rm in}, 0, 0)$ is a Euler matrix corresponding to the revolution about **z**-axis by an angle $\eta_{\rm in} \in [0, 2\pi]$. The latter describes the relative orientation of the nucleoprotein complex with respect to the axis of the DNA segment entering it. I.e., matrix $\mathbf{R}_{\eta_{\rm in}}$ and angle $\eta_{\rm in}$ basically tell on which side of the DNA the nucleoprotein complex is formed. Indeed, since $\mathbf{R}_j\mathbf{R}_{\eta_{\rm in}} = \mathbf{R}_{\alpha_j}\mathbf{R}_{\beta_j}\mathbf{R}_{\gamma_j}\mathbf{R}_{\eta_{\rm in}} = \mathbf{R}_{\alpha_j}\mathbf{R}_{\beta_j}\mathbf{R}_{\gamma_j+\eta_{\rm in}}$, it is clear that angle $\eta_{\rm in}$ simply introduces rotation of the nucleoprotein complex with respect to the axis of the DNA segment entering it. Here $(\alpha_j, \beta_j, \gamma_j)$ are Euler angles corresponding to the rotation matrix $\mathbf{R}_j = \mathbf{R}_j(\alpha_j, \beta_j, \gamma_j)$; and $\mathbf{R}_{\alpha_j}$, $\mathbf{R}_{\beta_j}$ and $\mathbf{R}_{\gamma_j}$ are rotation matrices describing the respective coordinate system revolutions through angles $\alpha_j$, $\beta_j$ and $\gamma_j$.

It should be noted that the above matrix product, $\mathbf{R}_j\mathbf{R}_{\eta_{\rm in}}$, has to be used only in $E_{01}$ energy term, without making similar changes in other energy terms corresponding to the downstream DNA segments in a nucleoprotein complex since orientations of the latter will be completely defined by the orientation of the first DNA segment bound to the protein.

Another note, which we wanted to make, concerns viral particle calculations shown in Figure 2. So far, in this Appendix section, we have considered only the case of chromosomal DNA packed in the cell nucleus. However, it is not very hard to obtain very similar formulas for the total energy of DNA confined inside a viral capsid, which can be done simply by removing $E_{\rm protein}$ and $E_f$ terms from Eq. (B1).

Anyway, since we have now all of the formulas necessary for introduction of the transfer-matrix formalism, let's proceed to its description, applying it to obtain a formula for computation of the partition function of DNA confined in the cell nucleus.

#### Appendix C: DNA partition function

In this section, we will further consider the special case of DNA interacting with proteins that occupy 3 DNA segments upon binding to DNA (i.e., $K = 3$), which has been introduced in Appendix B. By utilizing the transfer-matrix approach and gradually increasing its complexity, we are aiming at the end of this section to obtain a mathematical expression for the partition function of a mechanically stretched DNA in the presence of the potential field $\psi$ in a form of a transfer-matrix product.

To this aim, we start first with a general definition of the partition function of DNA, $Z_{\psi,f}$, which is stretched by mechanical force, $f$, in the electrostatic potential field, $\psi$:

$$Z_{\psi,f} = \sum_{k_1...k_N=0}^{K} \int_{\mathbb{R}^3} \mathrm{d}\mathbf{r}_0 \int \mathrm{d}\mathbf{R}_1...\mathrm{d}\mathbf{R}_N\, \mathrm{d}[\eta_{\rm in}]\, e^{-\beta E_{\rm tot}(k_1...k_N,\mathbf{R}_1...\mathbf{R}_N,\mathbf{r}_0)} \xi(\mathbf{R}_N, \mathbf{R}_1) \tag{C1}$$

Here $E_{\rm tot}$ is the total energy of DNA defined by Eq. (B16)–Eq. (B17). $\xi(\mathbf{R}_N, \mathbf{R}_1)$ is a function that imposes specific boundary conditions on orientations of the DNA end segments. Integration in the above formula is carried out over all possible orientations of DNA segments (i.e., $\int \mathrm{d}\mathbf{R}_1...\mathrm{d}\mathbf{R}_N = \int_0^{2\pi}\mathrm{d}\alpha_1...\mathrm{d}\alpha_N \int_0^{2\pi}\mathrm{d}\gamma_1...\mathrm{d}\gamma_N \int_0^{\pi}\sin\beta_1\,\mathrm{d}\beta_1...\sin\beta_N\,\mathrm{d}\beta_N$) as well as over the set $[\eta_{\rm in}]$ of angles $\eta_{{\rm in},j}$ describing the relative orientations of nucleoprotein complexes with respect to the DNA segments entering them, see the end of Appendix B for details. Here subscript $j$ is used to enumerate $\eta_{\rm in}$ angles according to positions of the corresponding nucleoprotein complexes on the DNA. I.e., angle $\eta_{{\rm in},j}$ refers to a nucleoprotein complex occupying DNA segments with indexes $j+1$, $j+2$ and $j+3$.

Applying the vector-valued integration technique described in ref. [1], it is not hard to show that the DNA partition function defined by Eq. (C1) obeys a number of recurrence relations, which are very similar to those derived in ref. [1, 3]. Using these relations, it is then possible to greatly simplify the expression for the DNA partition function by utilizing the transfer-matrix formalism.

To demonstrate this, let's first rewrite Eq. (C1) in a slightly different form by noting that integrals $\int \mathrm{d}[\eta_{\rm in}]$ are relevant only for DNA segments being in state $k_j = 0$, which are followed by a DNA segment in $k_{j+1} = 1$ state.

Indeed, from the comments after Eq. (B17) it can be seen that angle $\eta_{\rm in}$ appears only in $E_{01}$ energy term in the form of the rotation matrix $\mathbf{R}_{\eta_{\rm in}}$. This makes it possible to rewrite Eq. (C1) as:

$$Z_{\psi,f} = \sum_{k_1...k_N=0}^{K} \int_{\mathrm{IR}^3} \mathrm{d}\mathbf{r}_0 \int \mathrm{d}\mathbf{R}_1...\mathrm{d}\mathbf{R}_N \, \mathrm{d}\theta_1...\mathrm{d}\theta_N \, e^{-\beta E_{\rm tot}(k_1...k_N,\mathbf{R}_1...\mathbf{R}_N,\mathbf{r}_0)} \xi(\mathbf{R}_N, \mathbf{R}_1) \tag{C2}$$

Here $\int \mathrm{d}\theta_1...\mathrm{d}\theta_N$ is a shorthand notation for $\int \mathrm{d}[\eta_{\rm in}]$ integrals, which are calculated only for DNA segments being in the respective states:

$$\mathrm{d}\theta_j = \begin{cases} \mathrm{d}\eta_{\mathrm{in},j}, & \text{if } k_j = 0 \text{ and } k_{j+1} = 1 \\ 1 & , \text{ otherwise} \end{cases} \tag{C3}$$

To further simplify Eq. (C2), it is convenient to introduce level-one transfer-functions $T^{(1)}_{k_j k_{j+1}}(\mathbf{R}_j, \mathbf{R}_{j+1})$ defined as:

$$T^{(1)}_{k_j k_{j+1}}(\mathbf{R}_j, \mathbf{R}_{j+1}) = \begin{cases} \displaystyle\int_0^{2\pi} \mathrm{d}\eta_{\mathrm{in},j} \, e^{-\beta E_{k_j k_{j+1}}(\mathbf{R}_j,\mathbf{R}_{j+1})}, & \text{if } (k_j, k_{j+1}) = (0,1) \\ e^{-\beta E_{k_j k_{j+1}}(\mathbf{R}_j,\mathbf{R}_{j+1})} & , \text{ otherwise} \end{cases} \tag{C4}$$

Where $E_{k_j k_{j+1}}(\mathbf{R}_j, \mathbf{R}_{j+1})$ are local energy contributions made by DNA segments into the total energy of DNA, see Eq. (B16) and Eq. (B17).

Substituting Eq. (B16) into Eq. (C2) and using Eq. (C3)-Eq. (C4), we get:

$$Z_{\psi,f} = \sum_{k_1...k_N=0}^{K} \delta_{k_1 0} \times \int_{\mathrm{IR}^3} \mathrm{d}\mathbf{r}_0 \int \mathrm{d}\mathbf{R}_1...\mathrm{d}\mathbf{R}_N \prod_{j=1}^{N-1} \left[ T^{(1)}_{k_j k_{j+1}}(\mathbf{R}_j, \mathbf{R}_{j+1}) \, e^{-\beta \hat{\psi}_{k_j k_{j+1}}(\mathbf{r}_j)} \right] \times \xi^{(1)}_{k_N}(\mathbf{R}_N, \mathbf{R}_1) \, e^{-\beta \hat{\psi}_{00}(\mathbf{r}_N)} \tag{C5}$$

Where in the case of chromosomal DNA confined inside the cell nucleus $\hat{\psi}_{k_j k_{j+1}}$ describes the net action of $U_{R_{\rm nucl}}$ and $\psi$ fields on the $j^{\rm th}$ DNA segment: $\hat{\psi}_{k_j k_{j+1}}(\mathbf{r}_j) = \sigma_{k_j k_{j+1}} U_{R_{\rm nucl}}(\mathbf{r}_j) + q_{k_j k_{j+1}} \psi(\mathbf{r}_j)$. In the above formula, $\xi^{(1)}_{k_N}(\mathbf{R}_N, \mathbf{R}_1)$ is the level-one boundary condition function defined as:

$$\xi^{(1)}_{k_N}(\mathbf{R}_N, \mathbf{R}_1) = \delta_{k_N 0} \times \xi(\mathbf{R}_N, \mathbf{R}_1) \, e^{\beta b f [\mathbf{z}_0 \cdot \mathbf{R}_N \mathbf{z}_0]} \tag{C6}$$

Let's consider in more detail $e^{-\beta \hat{\psi}_{k_j k_{j+1}}(\mathbf{r}_j)}$ exponential function. Since $\mathbf{r}_j = \mathbf{r}_0 + \sum_{n=1}^{j} b_{k_n} \mathbf{R}_n \mathbf{z}_0$ [Eq. (B10)], it is not hard to see that it can be expressed in the following form with the help of the integral representation of the Dirac $\delta$-function:

$$e^{-\beta \hat{\psi}_{k_j k_{j+1}}(\mathbf{r}_j)} = \int_{\mathrm{IR}^3} \mathrm{d}\mathbf{r}'_j \, e^{-\beta \hat{\psi}_{k_j k_{j+1}}(\mathbf{r}'_j)} \, \delta\Big(\mathbf{r}_0 + \sum_{n=1}^{j} b_{k_n} \mathbf{R}_n \mathbf{z}_0 - \mathbf{r}'_j\Big) =$$
$$= \frac{1}{(2\pi)^3} \int_{\mathrm{IR}^3} \mathrm{d}\mathbf{r}'_j \int_{\mathrm{IR}^3} \mathrm{d}\boldsymbol{\omega}_j \, e^{-\beta \hat{\psi}_{k_j k_{j+1}}(\mathbf{r}'_j)} \, e^{-i[\boldsymbol{\omega}_j \cdot \mathbf{r}'_j] + i[\boldsymbol{\omega}_j \cdot \sum_{n=1}^{j} b_{k_n} \mathbf{R}_n \mathbf{z}_0] + i[\boldsymbol{\omega}_j \cdot \mathbf{r}_0]} \tag{C7}$$

Here $\int_{\mathrm{IR}^3} \mathrm{d}\mathbf{r}'_j = \int_{\mathrm{IR}^3} \mathrm{d}x'_j \mathrm{d}y'_j \mathrm{d}z'_j$, where $(x'_j, y'_j, z'_j)$ are coordinates of the position vector $\mathbf{r}'_j$. Similarly, for the wave-vector $\boldsymbol{\omega}_j = (\omega^x_j, \omega^y_j, \omega^z_j)$, we have: $\int_{\mathrm{IR}^3} \mathrm{d}\boldsymbol{\omega}_j = \int_{\mathrm{IR}^3} \mathrm{d}\omega^x_j \mathrm{d}\omega^y_j \mathrm{d}\omega^z_j$.

Substituting Eq. (C7) into Eq. (C5), it can be shown that the formula for the DNA partition function can be rewritten as:

$$Z_{\psi,f} = \sum_{k_1...k_N=0}^{K} \delta_{k_1 0} \times \int_{\rm I\!R^3} d\mathbf{r}_0 \int d\mathbf{R}_1...d\mathbf{R}_N \int_{\rm I\!R^{3N}} d\boldsymbol{\omega}_1...d\boldsymbol{\omega}_N \prod_{j=1}^{N-1} \Big[ T^{(1)}_{k_j k_{j+1}}(\mathbf{R}_j, \mathbf{R}_{j+1})\, \Phi_{k_j k_{j+1}}(\boldsymbol{\omega}_j) \times$$
$$\times\, e^{i[\boldsymbol{\omega}_j \cdot \sum_{n=1}^{j} b_{k_n} \mathbf{R}_n \mathbf{z}_0] + i[\boldsymbol{\omega}_j \cdot \mathbf{r}_0]} \Big] \times \xi^{(1)}_{k_N}(\mathbf{R}_N, \mathbf{R}_1)\, \Phi_{00}(\boldsymbol{\omega}_N)\, e^{i[\boldsymbol{\omega}_N \cdot \sum_{n=1}^{N} b_{k_n} \mathbf{R}_n \mathbf{z}_0] + i[\boldsymbol{\omega}_N \cdot \mathbf{r}_0]} \qquad \text{(C8)}$$

Where $\Phi_{k_j k_{j+1}}(\boldsymbol{\omega}_j)$ is the Fourier-transform of the field exponential function:

$$\Phi_{k_j k_{j+1}}(\boldsymbol{\omega}_j) = \frac{1}{(2\pi)^3} \int_{\rm I\!R^3} d\mathbf{r}'_j\, e^{-\beta \hat{\psi}_{k_j k_{j+1}}(\mathbf{r}'_j)}\, e^{-i[\boldsymbol{\omega}_j \cdot \mathbf{r}'_j]} \qquad \text{(C9)}$$

For further derivations, it is convenient to make a change in the order of summation of the exponential factors in Eq. (C8): $\sum_{j=1}^{N} \left[\boldsymbol{\omega}_j \cdot \sum_{n=1}^{j} b_{k_n} \mathbf{R}_n \mathbf{z}_0\right] = \sum_{n=1}^{N} \left[b_{k_n} \mathbf{R}_n \mathbf{z}_0 \cdot \sum_{j=n}^{N} \boldsymbol{\omega}_j\right] = [j \leftrightarrow n] = \sum_{j=1}^{N} \left[b_{k_j} \mathbf{R}_j \mathbf{z}_0 \cdot \sum_{n=j}^{N} \boldsymbol{\omega}_n\right]$, yielding:

$$Z_{\psi,f} = \sum_{k_1...k_N=0}^{K} \delta_{k_1 0} \times \int_{\rm I\!R^3} d\mathbf{r}_0 \int d\mathbf{R}_1...d\mathbf{R}_N \int_{\rm I\!R^{3N}} d\boldsymbol{\omega}_1...d\boldsymbol{\omega}_N \prod_{j=1}^{N-1} \Big[ T^{(1)}_{k_j k_{j+1}}(\mathbf{R}_j, \mathbf{R}_{j+1})\, \Phi_{k_j k_{j+1}}(\boldsymbol{\omega}_j) \times$$
$$\times\, e^{i b_{k_j} [\mathbf{R}_j \mathbf{z}_0 \cdot \sum_{n=j}^{N} \boldsymbol{\omega}_n]} \Big] \times e^{i[\mathbf{r}_0 \cdot \sum_{n=1}^{N} \boldsymbol{\omega}_n]} \times \xi^{(1)}_{k_N}(\mathbf{R}_N, \mathbf{R}_1)\, \Phi_{00}(\boldsymbol{\omega}_N)\, e^{i b [\mathbf{R}_N \mathbf{z}_0 \cdot \boldsymbol{\omega}_N]} \qquad \text{(C10)}$$

To simplify Eq. (C10), let's introduce new wave-vectors, $(\boldsymbol{\nu}_1, ..., \boldsymbol{\nu}_N)$, instead of old ones, $(\boldsymbol{\omega}_1, ..., \boldsymbol{\omega}_N)$:

$$\boldsymbol{\nu}_j = \sum_{n=j}^{N} \boldsymbol{\omega}_n \qquad \text{(C11)}$$

It is easy to see that the determinant of $\frac{\partial(\boldsymbol{\omega}_1,...,\boldsymbol{\omega}_N)}{\partial(\boldsymbol{\nu}_1,...,\boldsymbol{\nu}_N)}$ Jacobian matrix equals to unity: $\left|\frac{\partial(\boldsymbol{\omega}_1,...,\boldsymbol{\omega}_N)}{\partial(\boldsymbol{\nu}_1,...,\boldsymbol{\nu}_N)}\right| = 1$. Thus, after substituting Eq. (C11) into Eq. (C10) and integrating over $\mathbf{r}_0$, we get:

$$Z_{\psi,f} = (2\pi)^3 \sum_{k_1...k_N=0}^{K} \delta_{k_1 0} \times \int d\mathbf{R}_1...d\mathbf{R}_N \int_{\rm I\!R^{3N}} d\boldsymbol{\nu}_1...d\boldsymbol{\nu}_N \prod_{j=1}^{N-1} \Big[ T^{(1)}_{k_j k_{j+1}}(\mathbf{R}_j, \mathbf{R}_{j+1})\, \Phi_{k_j k_{j+1}}(\boldsymbol{\nu}_j - \boldsymbol{\nu}_{j+1})\, e^{i b_{k_j} [\mathbf{R}_j \mathbf{z}_0 \cdot \boldsymbol{\nu}_j]} \Big] \times$$
$$\times\, \xi^{(1)}_{k_N}(\mathbf{R}_N, \mathbf{R}_1)\, \delta(\boldsymbol{\nu}_1)\, \Phi_{00}(\boldsymbol{\nu}_N)\, e^{i b [\mathbf{R}_N \mathbf{z}_0 \cdot \boldsymbol{\nu}_N]} \qquad \text{(C12)}$$

Finally, by combining $\Phi_{k_j k_{j+1}}(\boldsymbol{\nu}_j - \boldsymbol{\nu}_{j+1})$ and $e^{i b_{k_j} [\mathbf{R}_j \mathbf{z}_0 \cdot \boldsymbol{\nu}_j]}$ terms with the transfer-functions, $T^{(1)}_{k_j k_{j+1}}(\mathbf{R}_j, \mathbf{R}_{j+1})$, and the boundary condition function, $\xi^{(1)}_{k_N}(\mathbf{R}_N, \mathbf{R}_1)$, one can further streamline the above expression for the DNA partition function:

$$Z_{\psi,f} = (2\pi)^3 \sum_{k_1...k_N=0}^{K} \delta_{k_1 0} \times \int d\mathbf{R}_1...d\mathbf{R}_N \int_{\rm I\!R^{3N}} d\boldsymbol{\nu}_1...d\boldsymbol{\nu}_N \prod_{j=1}^{N-1} T^{(3)}_{k_j k_{j+1}}(\mathbf{R}_j, \mathbf{R}_{j+1}, \boldsymbol{\nu}_j, \boldsymbol{\nu}_{j+1}) \times \xi^{(3)}_{k_N}(\mathbf{R}_N, \mathbf{R}_1, \boldsymbol{\nu}_N, \boldsymbol{\nu}_1) \qquad \text{(C13)}$$

Here $T^{(3)}_{k_j k_{j+1}}(\mathbf{R}_j, \mathbf{R}_{j+1}, \boldsymbol{\nu}_j, \boldsymbol{\nu}_{j+1})$ and $\xi^{(3)}_{k_N}(\mathbf{R}_N, \mathbf{R}_1, \boldsymbol{\nu}_N, \boldsymbol{\nu}_1)$ are level-three transfer-functions and the boundary condition function, which are defined as:

$$T^{(3)}_{k_j k_{j+1}}(\mathbf{R}_j, \mathbf{R}_{j+1}, \boldsymbol{\nu}_j, \boldsymbol{\nu}_{j+1}) = T^{(2)}_{k_j k_{j+1}}(\mathbf{R}_j, \mathbf{R}_{j+1}, \boldsymbol{\nu}_j)\ \Phi_{k_j k_{j+1}}(\boldsymbol{\nu}_j - \boldsymbol{\nu}_{j+1})$$
$$\xi^{(3)}_{k_N}(\mathbf{R}_N, \mathbf{R}_1, \boldsymbol{\nu}_N, \boldsymbol{\nu}_1) = \xi^{(2)}_{k_N}(\mathbf{R}_N, \mathbf{R}_1, \boldsymbol{\nu}_N)\ \delta(\boldsymbol{\nu}_1)\ \Phi_{00}(\boldsymbol{\nu}_N) \qquad \text{(C14)}$$

Where:

$$T^{(2)}_{k_j k_{j+1}}(\mathbf{R}_j, \mathbf{R}_{j+1}, \boldsymbol{\nu}_j) = T^{(1)}_{k_j k_{j+1}}(\mathbf{R}_j, \mathbf{R}_{j+1})\, e^{ib_{k_j}[\mathbf{R}_j \mathbf{z}_0 \cdot \boldsymbol{\nu}_j]}$$
$$\xi^{(2)}_{k_N}(\mathbf{R}_N, \mathbf{R}_1, \boldsymbol{\nu}_N) = \xi^{(1)}_{k_N}(\mathbf{R}_N, \mathbf{R}_1)\, e^{ib[\mathbf{R}_N \mathbf{z}_0 \cdot \boldsymbol{\nu}_N]} \tag{C15}$$

Later, in Appendix G, we are going to use $T^{(2)}_{k_j k_{j+1}}(\mathbf{R}_j, \mathbf{R}_{j+1}, \boldsymbol{\nu}_j, \boldsymbol{\nu}_{j+1})$ and $\xi^{(2)}_{k_N}(\mathbf{R}_N, \mathbf{R}_1, \boldsymbol{\nu}_N, \boldsymbol{\nu}_1)$ functions to derive formulas for the elements of the DNA transfer-matrix, $\mathbf{L}$, and the boundary condition matrix, $\mathbf{Y}$, which play the central role in the DNA partition function calculations. However, before proceeding to the construction of the DNA transfer-matrix, we want make several comments regarding $T^{(3)}_{01}$, $T^{(3)}_{12}$, $T^{(3)}_{23}$ and $T^{(3)}_{30}$ functions in Eq. (C13)-Eq. (C14), which describe the contribution of nucleoprotein complexes into the DNA partition function. As mentioned in Appendix B, $U_{R_{\text{nucl}}}$ and $\psi$ potential fields are introduced into the model in such a way that they act only on those joints of the polygonal chain representing DNA which are located outside of nucleoprotein complexes, see Figure S9. This leads to an extremely simple formula for $\Phi_{k_j k_{j+1}}(\boldsymbol{\nu}_j - \boldsymbol{\nu}_{j+1})$ function in the case of protein-bound DNA segments. Specifically, if the $j^{\text{th}}$ DNA segment is located near, or inside a nucleoprotein complex, such that it is followed by a DNA segment bound to a protein (i.e., $k_{j+1} \neq 0$), then from Eq. (C9) and definition of $\hat{\psi}_{k_j k_{j+1}}$ field action (Appendix B) it follows that:

$$k_{j+1} \neq 0 \rightarrow \hat{\psi}_{k_j k_{j+1}} = 0, \text{ and thus: } \Phi_{k_j k_{j+1}}(\boldsymbol{\nu}_j - \boldsymbol{\nu}_{j+1}) = \frac{1}{(2\pi)^3} \int_{\mathbb{R}^3} d\mathbf{r}'_j\, e^{-i[(\boldsymbol{\nu}_j - \boldsymbol{\nu}_{j+1}) \cdot \mathbf{r}'_j]} = \delta(\boldsymbol{\nu}_j - \boldsymbol{\nu}_{j+1}) \tag{C16}$$

As a result, all of the transfer-functions associated with DNA segments, which are bound to the same nucleoprotein complex, can be conveniently combined into a single term. Namely, if a nucleoprotein complex occupies DNA segments with indexes $j$, $j+1$ and $j+2$, then it is clear that the following three products of transfer-functions under the integral sign in Eq. (C13) lead to the same result in terms of the DNA partition function value:

$$\begin{aligned}
&... \times T^{(3)}_{00}(\boldsymbol{\nu}_{j-2}, \boldsymbol{\nu}_{j-1}) \times T^{(3)}_{01}(\boldsymbol{\nu}_{j-1}, \boldsymbol{\nu}_j) \times T^{(3)}_{12}(\boldsymbol{\nu}_j, \boldsymbol{\nu}_{j+1}) \times T^{(3)}_{23}(\boldsymbol{\nu}_{j+1}, \boldsymbol{\nu}_{j+2}) \times T^{(3)}_{30}(\boldsymbol{\nu}_{j+2}, \boldsymbol{\nu}_{j+3}) \times \\
&\times T^{(3)}_{00}(\boldsymbol{\nu}_{j+3}, \boldsymbol{\nu}_{j+4}) \times ... \equiv \\
\equiv\ &... \times T^{(3)}_{00}(\boldsymbol{\nu}_{j-2}, \boldsymbol{\nu}_{j-1}) \times \left[ T^{(2)}_{01}(\boldsymbol{\nu}_{j-1})\, T^{(2)}_{12}(\boldsymbol{\nu}_{j-1})\, T^{(2)}_{23}(\boldsymbol{\nu}_{j-1})\, T^{(3)}_{30}(\boldsymbol{\nu}_{j-1}, \boldsymbol{\nu}_{j+3}) \right] \times \\
&\times \delta(\boldsymbol{\nu}_{j-1} - \boldsymbol{\nu}_j) \times \delta(\boldsymbol{\nu}_j - \boldsymbol{\nu}_{j+1}) \times \delta(\boldsymbol{\nu}_{j+1} - \boldsymbol{\nu}_{j+2}) \times T^{(3)}_{00}(\boldsymbol{\nu}_{j+3}, \boldsymbol{\nu}_{j+4}) \times ... \\
\equiv\ &... \times T^{(3)}_{00}(\boldsymbol{\nu}_{j-2}, \boldsymbol{\nu}_{j-1}) \times \left[ T^{(2)}_{01}(\boldsymbol{\nu}_{j-1})\, T^{(2)}_{12}(\boldsymbol{\nu}_{j-1})\, T^{(2)}_{23}(\boldsymbol{\nu}_{j-1})\, T^{(3)}_{30}(\boldsymbol{\nu}_{j-1}, \boldsymbol{\nu}_j) \right] \times \\
&\times \delta(\boldsymbol{\nu}_j - \boldsymbol{\nu}_{j+1}) \times \delta(\boldsymbol{\nu}_{j+1} - \boldsymbol{\nu}_{j+2}) \times \delta(\boldsymbol{\nu}_{j+2} - \boldsymbol{\nu}_{j+3}) \times T^{(3)}_{00}(\boldsymbol{\nu}_{j+3}, \boldsymbol{\nu}_{j+4}) \times ...
\end{aligned} \tag{C17}$$

Here we applied Eq. (C14) and Eq. (C16) to $T^{(3)}_{01}$, $T^{(3)}_{12}$ and $T^{(3)}_{23}$ terms. In addition, for the sake of formula simplicity, we omitted $\mathbf{R}_{j-2}$, $\mathbf{R}_{j-1}$, ..., $\mathbf{R}_{j+4}$ arguments of the transfer-functions.

Based on this observation, it follows that one can simplify most of the terms in Eq. (C13) related to nucleoprotein complexes by introducing level-four transfer-functions defined as:

$$\begin{cases} T^{(4)}_{00}(\mathbf{R}_j,\mathbf{R}_{j+1},\boldsymbol{\nu}_j,\boldsymbol{\nu}_{j+1}) = T^{(3)}_{00}(\mathbf{R}_j,\mathbf{R}_{j+1},\boldsymbol{\nu}_j,\boldsymbol{\nu}_{j+1}) \\ T^{(4)}_{01}(\mathbf{R}_j,\mathbf{R}_{j+1},\boldsymbol{\nu}_j,\boldsymbol{\nu}_{j+1}) = \int \mathrm{d}\mathbf{R}'\mathrm{d}\mathbf{R}''\mathrm{d}\mathbf{R}'''\, T^{(2)}_{01}(\mathbf{R}_j,\mathbf{R}',\boldsymbol{\nu}_j)\, T^{(2)}_{12}(\mathbf{R}',\mathbf{R}'',\boldsymbol{\nu}_j)\, T^{(2)}_{23}(\mathbf{R}'',\mathbf{R}''',\boldsymbol{\nu}_j) \times \\ \qquad \times\, T^{(3)}_{30}(\mathbf{R}''',\mathbf{R}_{j+1},\boldsymbol{\nu}_j,\boldsymbol{\nu}_{j+1}) \\ T^{(4)}_{12}(\mathbf{R}_j,\mathbf{R}_{j+1},\boldsymbol{\nu}_j,\boldsymbol{\nu}_{j+1}) = T^{(4)}_{23}(\mathbf{R}_j,\mathbf{R}_{j+1},\boldsymbol{\nu}_j,\boldsymbol{\nu}_{j+1}) = T^{(4)}_{30}(\mathbf{R}_j,\mathbf{R}_{j+1},\boldsymbol{\nu}_j,\boldsymbol{\nu}_{j+1}) = \delta(\mathbf{R}_j-\mathbf{R}_{j+1})\,\delta(\boldsymbol{\nu}_j-\boldsymbol{\nu}_{j+1}) \\ T^{(4)}_{02} = T^{(4)}_{03} = T^{(4)}_{10} = T^{(4)}_{11} = T^{(4)}_{13} = T^{(4)}_{20} = T^{(4)}_{21} = T^{(4)}_{22} = T^{(4)}_{31} = T^{(4)}_{32} = T^{(4)}_{33} = 0 \end{cases} \tag{C18}$$

Substituting Eq. (C17) and Eq. (C18) into Eq. (C13), it is not hard to see that:

$$Z_{\psi,f} = (2\pi)^3 \sum_{k_1\ldots k_N=0}^{K} \delta_{k_1 0} \times \int \mathrm{d}\mathbf{R}_1\ldots\mathrm{d}\mathbf{R}_N \int_{\mathrm{I\!R}^{3N}} \mathrm{d}\boldsymbol{\nu}_1\ldots\mathrm{d}\boldsymbol{\nu}_N \prod_{j=1}^{N-1} T^{(4)}_{k_j k_{j+1}}(\mathbf{R}_j,\mathbf{R}_{j+1},\boldsymbol{\nu}_j,\boldsymbol{\nu}_{j+1}) \times \xi^{(3)}_{k_N}(\mathbf{R}_N,\mathbf{R}_1,\boldsymbol{\nu}_N,\boldsymbol{\nu}_1) \tag{C19}$$

It should be noted that despite a very similar appearance of Eq. (C13) and Eq. (C19), introduction of the level-four transfer-functions has multiple benefits, leading to construction of the conveyor belt algorithm for calculation of the DNA partition function (see Appendix I), which considerably reduces the amount of computational resources needed for evaluation of the DNA partition function.

Anyway, to derive a formula for the DNA partition function in terms of a transfer-matrix product, we will follow the same steps as in the case of a semiflexible polymer chain model of DNA, which has been described in our previous studies [1, 3]. Indeed, starting from Eq. (C19), it is not very hard to show that the DNA partition function obeys a number of recurrence relations, which can be obtained by using intermediary partition functions defined as:

$$Z_s(k_s,\mathbf{R}_s,\mathbf{R}_1,\boldsymbol{\nu}_s,\boldsymbol{\nu}_1) = \sum_{k_{s+1}\ldots k_N=0}^{K} \int \mathrm{d}\mathbf{R}_{s+1}\ldots\mathrm{d}\mathbf{R}_N \int_{\mathrm{I\!R}^{3(N-s)}} \mathrm{d}\boldsymbol{\nu}_{s+1}\ldots\mathrm{d}\boldsymbol{\nu}_N \prod_{j=s}^{N-1} T^{(4)}_{k_j k_{j+1}}(\mathbf{R}_j,\mathbf{R}_{j+1},\boldsymbol{\nu}_j,\boldsymbol{\nu}_{j+1}) \times \\ \times\, \xi^{(3)}_{k_N}(\mathbf{R}_N,\mathbf{R}_1,\boldsymbol{\nu}_N,\boldsymbol{\nu}_1) \tag{C20}$$

Where $1 \le s \le N-1$.

From Eq. (C19) and (C20), it can be seen that the DNA partition function, $Z_{\psi,f}$, equals to:

$$Z_{\psi,f} = (2\pi)^3 \sum_{k_1=0}^{K} \delta_{k_1 0} \times \int \mathrm{d}\mathbf{R}_1 \int_{\mathrm{I\!R}^3} \mathrm{d}\boldsymbol{\nu}_1\, Z_1(k_1,\mathbf{R}_1,\mathbf{R}_1,\boldsymbol{\nu}_1,\boldsymbol{\nu}_1) = \\ = (2\pi)^3 \int \mathrm{d}\mathbf{R}_1 \int_{\mathrm{I\!R}^3} \mathrm{d}\boldsymbol{\nu}_1 \begin{pmatrix} 1 & 0 & 0 & 0 \end{pmatrix} \times \begin{pmatrix} Z_1(0,\mathbf{R}_1,\mathbf{R}_1,\boldsymbol{\nu}_1,\boldsymbol{\nu}_1) \\ Z_1(1,\mathbf{R}_1,\mathbf{R}_1,\boldsymbol{\nu}_1,\boldsymbol{\nu}_1) \\ Z_1(2,\mathbf{R}_1,\mathbf{R}_1,\boldsymbol{\nu}_1,\boldsymbol{\nu}_1) \\ Z_1(3,\mathbf{R}_1,\mathbf{R}_1,\boldsymbol{\nu}_1,\boldsymbol{\nu}_1) \end{pmatrix} \tag{C21}$$

What is even more important, from the definition of the intermediary partition functions it follows that they obey the following recurrence relation:

$$Z_{s-1}(k_{s-1},\mathbf{R}_{s-1},\mathbf{R}_1,\boldsymbol{\nu}_{s-1},\boldsymbol{\nu}_1) = \int \mathrm{d}\mathbf{R}_s \int_{\mathrm{I\!R}^3} \mathrm{d}\boldsymbol{\nu}_s \begin{pmatrix} T^{(4)}_{k_{s-1}0} & T^{(4)}_{k_{s-1}1} & T^{(4)}_{k_{s-1}2} & T^{(4)}_{k_{s-1}3} \end{pmatrix} \times \begin{pmatrix} Z_s(0,\mathbf{R}_s,\mathbf{R}_1,\boldsymbol{\nu}_s,\boldsymbol{\nu}_1) \\ Z_s(1,\mathbf{R}_s,\mathbf{R}_1,\boldsymbol{\nu}_s,\boldsymbol{\nu}_1) \\ Z_s(2,\mathbf{R}_s,\mathbf{R}_1,\boldsymbol{\nu}_s,\boldsymbol{\nu}_1) \\ Z_s(3,\mathbf{R}_s,\mathbf{R}_1,\boldsymbol{\nu}_s,\boldsymbol{\nu}_1) \end{pmatrix}, \tag{C22}$$

which can be conveniently rewritten in a more compact form by using the vector-valued integration technique (see Appendix F in ref. [1]):

$$\begin{pmatrix} Z_{s-1}(0,\mathbf{R}_{s-1},\mathbf{R}_1,\boldsymbol{\nu}_{s-1},\boldsymbol{\nu}_1) \\ Z_{s-1}(1,\mathbf{R}_{s-1},\mathbf{R}_1,\boldsymbol{\nu}_{s-1},\boldsymbol{\nu}_1) \\ Z_{s-1}(2,\mathbf{R}_{s-1},\mathbf{R}_1,\boldsymbol{\nu}_{s-1},\boldsymbol{\nu}_1) \\ Z_{s-1}(3,\mathbf{R}_{s-1},\mathbf{R}_1,\boldsymbol{\nu}_{s-1},\boldsymbol{\nu}_1) \end{pmatrix} = \int d\mathbf{R}_s \int_{\mathrm{I\!R}^3} d\boldsymbol{\nu}_s\, \mathbf{T}^{(4)}(\mathbf{R}_{s-1},\mathbf{R}_s,\boldsymbol{\nu}_{s-1},\boldsymbol{\nu}_s) \times \begin{pmatrix} Z_{s}(0,\mathbf{R}_{s},\mathbf{R}_1,\boldsymbol{\nu}_{s},\boldsymbol{\nu}_1) \\ Z_{s}(1,\mathbf{R}_{s},\mathbf{R}_1,\boldsymbol{\nu}_{s},\boldsymbol{\nu}_1) \\ Z_{s}(2,\mathbf{R}_{s},\mathbf{R}_1,\boldsymbol{\nu}_{s},\boldsymbol{\nu}_1) \\ Z_{s}(3,\mathbf{R}_{s},\mathbf{R}_1,\boldsymbol{\nu}_{s},\boldsymbol{\nu}_1) \end{pmatrix} \tag{C23}$$

Here $\mathbf{T}^{(4)}(\mathbf{R}_{s-1},\mathbf{R}_s,\boldsymbol{\nu}_{s-1},\boldsymbol{\nu}_s)$ is the DNA transfer-matrix, which is defined as:

$$\mathbf{T}^{(4)}(\mathbf{R}_{s-1},\mathbf{R}_s,\boldsymbol{\nu}_{s-1},\boldsymbol{\nu}_s) = \begin{pmatrix} T^{(4)}_{00} & T^{(4)}_{01} & T^{(4)}_{02} & T^{(4)}_{03} \\ T^{(4)}_{10} & T^{(4)}_{11} & T^{(4)}_{12} & T^{(4)}_{13} \\ T^{(4)}_{20} & T^{(4)}_{21} & T^{(4)}_{22} & T^{(4)}_{23} \\ T^{(4)}_{30} & T^{(4)}_{31} & T^{(4)}_{32} & T^{(4)}_{33} \end{pmatrix} = \begin{pmatrix} T^{(4)}_{00} & T^{(4)}_{01} & 0 & 0 \\ 0 & 0 & T^{(4)}_{12} & 0 \\ 0 & 0 & 0 & T^{(4)}_{23} \\ T^{(4)}_{30} & 0 & 0 & 0 \end{pmatrix} \tag{C24}$$

In the right part of the above formula, we took into account the last line of Eq. (C18), which basically tells that all of the matrix entries corresponding to partially unfolded nucleoprotein complexes are equal to zero due to the infinitely high energy of such DNA-protein conformations, see Eq. (B6) and Eq. (B17). For the sake of formulas simplicity, the arguments of $T^{(4)}_{nm}(\mathbf{R}_{s-1},\mathbf{R}_s,\boldsymbol{\nu}_{s-1},\boldsymbol{\nu}_s)$ transfer-functions are omitted in Eq. (C22) and (C24).

Combining together Eq. (C21) and (C23), we finally obtain a mathematical expression for the DNA partition function in terms of the DNA transfer-matrix product:

$$Z_{\psi,f} = (2\pi)^3 \begin{pmatrix} 1 & 0 & 0 & 0 \end{pmatrix} \times \int d\mathbf{R}_1 ... d\mathbf{R}_N \int_{\mathrm{I\!R}^{3N}} d\boldsymbol{\nu}_1 ... d\boldsymbol{\nu}_N \prod_{j=1}^{N-1} \mathbf{T}^{(4)}(\mathbf{R}_j,\mathbf{R}_{j+1},\boldsymbol{\nu}_j,\boldsymbol{\nu}_{j+1}) \times \boldsymbol{\xi}^{(3)}(\mathbf{R}_N,\mathbf{R}_1,\boldsymbol{\nu}_N,\boldsymbol{\nu}_1) \tag{C25}$$

Where the boundary condition vector, $\boldsymbol{\xi}^{(3)}(\mathbf{R}_N,\mathbf{R}_1,\boldsymbol{\nu}_N,\boldsymbol{\nu}_1)$, is:

$$\boldsymbol{\xi}^{(3)}(\mathbf{R}_N,\mathbf{R}_1,\boldsymbol{\nu}_N,\boldsymbol{\nu}_1) = \begin{pmatrix} \xi^{(3)}_0(\mathbf{R}_N,\mathbf{R}_1,\boldsymbol{\nu}_N,\boldsymbol{\nu}_1) \\ 0 \\ 0 \\ 0 \end{pmatrix} \tag{C26}$$

To further streamline Eq. (C25), one may recall that any square-integrable function defined on SO(3) group of 3D rotation matrices parametrized by Euler angles $(\alpha,\beta,\gamma)$ can be expanded into a series of the Wigner D-functions, $D^s_{p,q}(\alpha,\beta,\gamma)$ [36]. Performing such an expansion with respect to $\mathbf{R}_j$ and $\mathbf{R}_{j+1}$ arguments of $T^{(4)}_{nm}(\mathbf{R}_j,\mathbf{R}_{j+1},\boldsymbol{\nu}_j,\boldsymbol{\nu}_{j+1})$ elements of the DNA transfer-matrix, $\mathbf{T}^{(4)}(\mathbf{R}_j,\mathbf{R}_{j+1},\boldsymbol{\nu}_j,\boldsymbol{\nu}_{j+1})$, we obtain the following series of $D^s_{p,q}$ functions [see Appendices F-G for more details]:

$$T^{(4)}_{nm}(\mathbf{R}_j,\mathbf{R}_{j+1},\boldsymbol{\nu}_j,\boldsymbol{\nu}_{j+1}) = \frac{1}{8\pi^2} \sum_{\substack{p_j,p_{j+1}\\ q_j,q_{j+1}\\ s_j,s_{j+1}}} \sqrt{(2s_j+1)(2s_{j+1}+1)}\, [T^{(4)}_{nm}(\boldsymbol{\nu}_j,\boldsymbol{\nu}_{j+1})]^{p_{j+1},\,q_{j+1},\,s_{j+1}}_{p_j,\,q_j,\,s_j} D^{s_j}_{p_j,\,q_j}(\mathbf{R}_j)\, \overline{D}^{s_{j+1}}_{p_{j+1},\,q_{j+1}}(\mathbf{R}_{j+1}) \tag{C27}$$

Where $[T^{(4)}_{nm}(\boldsymbol{\nu}_j,\boldsymbol{\nu}_{j+1})]^{p_{j+1},\,q_{j+1},\,s_{j+1}}_{p_j,\,q_j,\,s_j}$ are expansion coefficients, whose values in the general case depend on $\boldsymbol{\nu}_j$ and $\boldsymbol{\nu}_{j+1}$ wave-vectors.

Then, by taking into account the linear property of matrices, it is not hard to see that $\mathbf{T}^{(4)}(\mathbf{R}_j,\mathbf{R}_{j+1},\boldsymbol{\nu}_j,\boldsymbol{\nu}_{j+1})$

transfer-matrix itself can be presented as:

$$\mathbf{T}^{(4)}(\mathbf{R}_j,\mathbf{R}_{j+1},\boldsymbol{\nu}_j,\boldsymbol{\nu}_{j+1}) = \frac{1}{8\pi^2}\sum_{\substack{p_j,p_{j+1}\\ q_j,q_{j+1}\\ s_j,s_{j+1}}}\sqrt{(2s_j+1)(2s_{j+1}+1)}\;\big[\mathbf{T}^{(4)}(\boldsymbol{\nu}_j,\boldsymbol{\nu}_{j+1})\big]^{p_{j+1},\,q_{j+1},\,s_{j+1}}_{p_j,\,q_j,\,s_j} D^{s_j}_{p_j,\,q_j}(\mathbf{R}_j)\,\overline{D}^{s_{j+1}}_{p_{j+1},\,q_{j+1}}(\mathbf{R}_{j+1}) \tag{C28}$$

Where each $[\mathbf{T}^{(4)}(\boldsymbol{\nu}_j,\boldsymbol{\nu}_{j+1})]^{p_{j+1},\,q_{j+1},\,s_{j+1}}_{p_j,\,q_j,\,s_j}$ coefficient denotes a matrix built of the expansion coefficients of the level-four transfer-functions, $T^{(4)}_{nm}(\mathbf{R}_j,\mathbf{R}_{j+1},\boldsymbol{\nu}_j,\boldsymbol{\nu}_{j+1})$:

$$\big[\mathbf{T}^{(4)}(\boldsymbol{\nu}_j,\boldsymbol{\nu}_{j+1})\big]^{p_{j+1},\,q_{j+1},\,s_{j+1}}_{p_j,\,q_j,\,s_j} = \begin{pmatrix} \big[T^{(4)}_{00}\big]^{p_{j+1},\,q_{j+1},\,s_{j+1}}_{p_j,\,q_j,\,s_j} & \big[T^{(4)}_{01}\big]^{p_{j+1},\,q_{j+1},\,s_{j+1}}_{p_j,\,q_j,\,s_j} & 0 & 0\\ 0 & 0 & \big[T^{(4)}_{12}\big]^{p_{j+1},\,q_{j+1},\,s_{j+1}}_{p_j,\,q_j,\,s_j} & 0\\ 0 & 0 & 0 & \big[T^{(4)}_{23}\big]^{p_{j+1},\,q_{j+1},\,s_{j+1}}_{p_j,\,q_j,\,s_j}\\ \big[T^{(4)}_{30}\big]^{p_{j+1},\,q_{j+1},\,s_{j+1}}_{p_j,\,q_j,\,s_j} & 0 & 0 & 0 \end{pmatrix} \tag{C29}$$

Here, as before, we omitted $\boldsymbol{\nu}_j$ and $\boldsymbol{\nu}_{j+1}$ arguments of the expansion coefficients of $T^{(4)}_{nm}(\mathbf{R}_j,\mathbf{R}_{j+1},\boldsymbol{\nu}_j,\boldsymbol{\nu}_{j+1})$ transfer-functions for the sake of formula simplicity.

Analogously, for the boundary condition vector, $\boldsymbol{\xi}^{(3)}(\mathbf{R}_N,\mathbf{R}_1,\boldsymbol{\nu}_N,\boldsymbol{\nu}_1)$, we have:

$$\boldsymbol{\xi}^{(3)}(\mathbf{R}_N,\mathbf{R}_1,\boldsymbol{\nu}_N,\boldsymbol{\nu}_1) = \frac{1}{8\pi^2}\sum_{\substack{p_1,\,q_1,\,s_1\\ p_N,\,q_N,\,s_N}}\sqrt{(2s_1+1)(2s_N+1)}\;\big[\boldsymbol{\xi}^{(3)}(\boldsymbol{\nu}_N,\boldsymbol{\nu}_1)\big]^{p_1,\,q_1,\,s_1}_{p_N,\,q_N,\,s_N} D^{s_N}_{p_N,q_N}(\mathbf{R}_N)\,\overline{D}^{s_1}_{p_1,q_1}(\mathbf{R}_1) \tag{C30}$$

Where $[\boldsymbol{\xi}^{(3)}(\boldsymbol{\nu}_N,\boldsymbol{\nu}_1)]^{p_1,\,q_1,\,s_1}_{p_N,\,q_N,\,s_N}$ is the following vector of expansion coefficients:

$$\big[\boldsymbol{\xi}^{(3)}(\boldsymbol{\nu}_N,\boldsymbol{\nu}_1)\big]^{p_1,\,q_1,\,s_1}_{p_N,\,q_N,\,s_N} = \begin{pmatrix} \big[\xi^{(3)}_0(\boldsymbol{\nu}_N,\boldsymbol{\nu}_1)\big]^{p_1,\,q_1,\,s_1}_{p_N,\,q_N,\,s_N}\\ 0\\ 0\\ 0 \end{pmatrix} \tag{C31}$$

After substituting Eq. (C28) and (C30) into Eq. (C25), and using orthogonality of $D^s_{p,q}$ functions [Eq. (F9)], it can be shown that all $\int \mathrm{d}\mathbf{R}_j$ integrals in Eq. (C25) reduce to a mere summation over the indexes of the expansion coefficients:

$$Z_{\psi,f} = (2\pi)^3\begin{pmatrix}1 & 0 & 0 & 0\end{pmatrix}\times\sum_{\substack{p_1\dots p_N\\ q_1\dots q_N\\ s_1\dots s_N}}\int_{\mathrm{I\!R}^{3N}}\mathrm{d}\boldsymbol{\nu}_1\dots\mathrm{d}\boldsymbol{\nu}_N\prod_{j=1}^{N-1}\big[\mathbf{T}^{(4)}(\boldsymbol{\nu}_j,\boldsymbol{\nu}_{j+1})\big]^{p_{j+1},\,q_{j+1},\,s_{j+1}}_{p_j,\,q_j,\,s_j}\times\big[\boldsymbol{\xi}^{(3)}(\boldsymbol{\nu}_N,\boldsymbol{\nu}_1)\big]^{p_1,\,q_1,\,s_1}_{p_N,\,q_N,\,s_N} \tag{C32}$$

The above expression can be further simplified, by noting that all of the expansion coefficients of $T^{(4)}_{00}$, $T^{(4)}_{12}$, $T^{(4)}_{23}$ and $T^{(4)}_{30}$ transfer-functions contain Kronecker $\delta_{q_jq_{j+1}}$ prefactor; whereas, expansion coefficients of $T^{(4)}_{01}$ transfer-function and $\xi^{(3)}_0$ boundary condition function have a $\delta_{q_j0}$ term, see Eq. (G40), Eq. (G44), Eq. (G49) and Eq. (G55). This is where the work done on reorganization of transfer-functions in Eq. (C17) starts to pay-off as all these Kronecker deltas combined together lead to nullification of $q_j$ indexes in Eq. (C32) via a domino-like effect, which is similar to the one discussed in Appendix C of ref. [1, 3].

By taking into account the above notes and putting $q_j = 0$ for all $j = 1,\dots,N$, we obtain the following expression for the DNA partition function:

$$Z_{\psi,f} = (2\pi)^3\begin{pmatrix}1 & 0 & 0 & 0\end{pmatrix}\times\sum_{\substack{p_1\dots p_N\\ s_1\dots s_N}}\int_{\mathrm{I\!R}^{3N}}\mathrm{d}\boldsymbol{\nu}_1\dots\mathrm{d}\boldsymbol{\nu}_N\prod_{j=1}^{N-1}\big[\mathbf{T}^{(4)}(\boldsymbol{\nu}_j,\boldsymbol{\nu}_{j+1})\big]^{p_{j+1},\,0,\,s_{j+1}}_{p_j,\,0,\,s_j}\times\big[\boldsymbol{\xi}^{(3)}(\boldsymbol{\nu}_N,\boldsymbol{\nu}_1)\big]^{p_1,\,0,\,s_1}_{p_N,\,0,\,s_N} \tag{C33}$$

It is convenient to slightly rearrange multidimensional arrays $[\mathbf{T}^{(4)}(\boldsymbol{\nu}_j,\boldsymbol{\nu}_{j+1})]^{p_{j+1},\,0,\,s_{j+1}}_{p_j,\,0,\,s_j}$ and $[\boldsymbol{\xi}^{(3)}(\boldsymbol{\nu}_N,\boldsymbol{\nu}_1)]^{p_1,\,0,\,s_1}_{p_N,0,\,s_N}$ in

Eq. (C33) by recalling from the definition of $D^{s_j}_{p_j,q_j}$ functions that index $p_j$ varies in the range of $-s_j \le p_j \le s_j$ for any given value of index $s_j$ (see Appendix F for more details). Similarly, for $p_{j+1}$ we have: $-s_{j+1} \le p_{j+1} \le s_{j+1}$. As a result, it can be shown that all $[T^{(4)}_{nm}(\boldsymbol{\nu}_j, \boldsymbol{\nu}_{j+1})]^{p_{j+1},\,0,\,s_{j+1}}_{p_j,\,0,\,s_j}$ arrays of the expansion coefficients can be organized in a form of $\mathbf{T}^{(5)}_{nm}(\boldsymbol{\nu}_j, \boldsymbol{\nu}_{j+1})$ matrices, whose elements are enumerated by a pair of indexes, $v_j = p_j + s_j(s_j+1)$ and $v_{j+1} = p_{j+1} + s_{j+1}(s_{j+1}+1)$:

$$\big[\mathbf{T}^{(5)}_{nm}(\boldsymbol{\nu}_j, \boldsymbol{\nu}_{j+1})\big]_{v_j v_{j+1}} = \big[T^{(4)}_{nm}(\boldsymbol{\nu}_j, \boldsymbol{\nu}_{j+1})\big]^{p_{j+1},\,0,\,s_{j+1}}_{p_j,\,0,\,s_j} \tag{C34}$$

Similarly, it is possible to arrange $[\xi^{(3)}_0(\boldsymbol{\nu}_N, \boldsymbol{\nu}_1)]^{p_1,\,0,\,s_1}_{p_N,\,0,\,s_N}$ expansion coefficients in a form of $\boldsymbol{\xi}^{(4)}_0(\boldsymbol{\nu}_N, \boldsymbol{\nu}_1)$ boundary condition matrix defined as:

$$\big[\boldsymbol{\xi}^{(4)}_0(\boldsymbol{\nu}_N, \boldsymbol{\nu}_1)\big]_{v_N v_1} = \big[\xi^{(3)}_0(\boldsymbol{\nu}_N, \boldsymbol{\nu}_1)\big]^{p_1,\,0,\,s_1}_{p_N,\,0,\,s_N} \tag{C35}$$

Where indexes $v_1$ and $v_N$ are equal to: $v_1 = p_1 + s_1(s_1+1)$ and $v_N = p_N + s_N(s_N+1)$.

Substituting Eq. (C34) and Eq. (C35) into Eq. (C29) and (C31), it is not hard to see that:

$$\big[\mathbf{T}^{(4)}(\boldsymbol{\nu}_j, \boldsymbol{\nu}_{j+1})\big]^{p_{j+1},\,0,\,s_{j+1}}_{p_j,\,0,\,s_j} = \begin{pmatrix} [\mathbf{T}^{(5)}_{00}]_{v_j v_{j+1}} & [\mathbf{T}^{(5)}_{01}]_{v_j v_{j+1}} & 0 & 0 \\ 0 & 0 & [\mathbf{T}^{(5)}_{12}]_{v_j v_{j+1}} & 0 \\ 0 & 0 & 0 & [\mathbf{T}^{(5)}_{23}]_{v_j v_{j+1}} \\ [\mathbf{T}^{(5)}_{30}]_{v_j v_{j+1}} & 0 & 0 & 0 \end{pmatrix} = \mathbf{T}^{(5)}(\boldsymbol{\nu}_j, \boldsymbol{\nu}_{j+1})\Big|_{v_j v_{j+1}}$$

$$\big[\boldsymbol{\xi}^{(3)}(\boldsymbol{\nu}_N, \boldsymbol{\nu}_1)\big]^{p_1,\,0,\,s_1}_{p_N,\,0,\,s_N} = \begin{pmatrix} [\boldsymbol{\xi}^{(4)}_0(\boldsymbol{\nu}_N, \boldsymbol{\nu}_1)]_{v_N v_1} \\ 0 \\ 0 \\ 0 \end{pmatrix} = \boldsymbol{\xi}^{(4)}(\boldsymbol{\nu}_N, \boldsymbol{\nu}_1)\Big|_{v_N v_1} \tag{C36}$$

Where $\mathbf{T}^{(5)}(\boldsymbol{\nu}_j, \boldsymbol{\nu}_{j+1})$ and $\boldsymbol{\xi}^{(4)}(\boldsymbol{\nu}_N, \boldsymbol{\nu}_1)$ are matrices composed of $\mathbf{T}^{(5)}_{nm}(\boldsymbol{\nu}_j, \boldsymbol{\nu}_{j+1})$ and $\boldsymbol{\xi}^{(4)}_0(\boldsymbol{\nu}_N, \boldsymbol{\nu}_1)$ blocks:

$$\mathbf{T}^{(5)}(\boldsymbol{\nu}_j, \boldsymbol{\nu}_{j+1}) = \begin{pmatrix} \mathbf{T}^{(5)}_{00}(\boldsymbol{\nu}_j, \boldsymbol{\nu}_{j+1}) & \mathbf{T}^{(5)}_{01}(\boldsymbol{\nu}_j, \boldsymbol{\nu}_{j+1}) & 0 & 0 \\ 0 & 0 & \mathbf{T}^{(5)}_{12}(\boldsymbol{\nu}_j, \boldsymbol{\nu}_{j+1}) & 0 \\ 0 & 0 & 0 & \mathbf{T}^{(5)}_{23}(\boldsymbol{\nu}_j, \boldsymbol{\nu}_{j+1}) \\ \mathbf{T}^{(5)}_{30}(\boldsymbol{\nu}_j, \boldsymbol{\nu}_{j+1}) & 0 & 0 & 0 \end{pmatrix} \text{ and } \boldsymbol{\xi}^{(4)}(\boldsymbol{\nu}_N, \boldsymbol{\nu}_1) = \begin{pmatrix} \boldsymbol{\xi}^{(4)}_0(\boldsymbol{\nu}_N, \boldsymbol{\nu}_1) \\ 0 \\ 0 \\ 0 \end{pmatrix}, \tag{C37}$$

and where we used the following shorthand notation:

$$\begin{pmatrix} \mathbf{A} & \cdots & \mathbf{B} \\ \vdots & \ddots & \vdots \\ \mathbf{C} & \cdots & \mathbf{D} \end{pmatrix}\Bigg|_{vv'} = \begin{pmatrix} [\mathbf{A}]_{vv'} & \cdots & [\mathbf{B}]_{vv'} \\ \vdots & \ddots & \vdots \\ [\mathbf{C}]_{vv'} & \cdots & [\mathbf{D}]_{vv'} \end{pmatrix} \tag{C38}$$

Here $\mathbf{A}$, $\mathbf{B}$, $\mathbf{C}$ and $\mathbf{D}$ are arbitrary matrices of the same size. $[\mathbf{A}]_{vv'}$, $[\mathbf{B}]_{vv'}$, $[\mathbf{C}]_{vv'}$ and $[\mathbf{D}]_{vv'}$ are their elements corresponding to a specific pair of indexes, $v$ and $v'$.

It should be noted that, in the general case, $\mathbf{T}^{(5)}_{nm}(\boldsymbol{\nu}_j, \boldsymbol{\nu}_{j+1})$ and $\boldsymbol{\xi}^{(4)}_0(\boldsymbol{\nu}_N, \boldsymbol{\nu}_1)$ blocks from Eq. (C37) are of infinite size. However, calculations show that the value of the DNA partition function is typically determined by several first harmonics corresponding to indexes $-s_j \le p_j \le s_j$, $-s_{j+1} \le p_{j+1} \le s_{j+1}$ and $0 \le s_j, s_{j+1} \le s_{\max}$, where $s_{\max} \sim 10$, see ref. [1–3]. Thus, it makes sense to use in computations finite $(s_{\max}{+}1)^2 \times (s_{\max}{+}1)^2$ square matrices, $\mathbf{T}^{(5)}_{nm}(\boldsymbol{\nu}_j, \boldsymbol{\nu}_{j+1})$ and $\boldsymbol{\xi}^{(4)}_0(\boldsymbol{\nu}_N, \boldsymbol{\nu}_1)$, which include only the first $(s_{\max}{+}1)^2$ rows and columns describing the above harmonics.

By utilizing $\mathbf{T}^{(5)}(\boldsymbol{\nu}_j, \boldsymbol{\nu}_{j+1})$ and $\boldsymbol{\xi}^{(4)}(\boldsymbol{\nu}_N, \boldsymbol{\nu}_1)$ block-matrices, it is then straightforward to apply the mathematical technique based on the generalized matrix multiplication formula described in Appendix E of ref. [1] in order to simplify Eq. (C33). Indeed, by substituting Eq. (C36) into Eq. (C33), one can obtain the following expression for the DNA partition function:

$$Z_{\psi,f} = (2\pi)^3 \begin{pmatrix} 1 & 0 & 0 & 0 \end{pmatrix} \times \sum_{v_1 \ldots v_N} \int_{\mathbb{R}^{3N}} \mathrm{d}\boldsymbol{\nu}_1 \ldots \mathrm{d}\boldsymbol{\nu}_N \prod_{j=1}^{N-1} \mathbf{T}^{(5)}(\boldsymbol{\nu}_j, \boldsymbol{\nu}_{j+1}) \Big|_{v_j v_{j+1}} \times \boldsymbol{\xi}^{(4)}(\boldsymbol{\nu}_N, \boldsymbol{\nu}_1) \Big|_{v_N v_1} \tag{C39}$$

Now, by using the usual matrix product definition, it is not hard to check that each of the sums over indexes $v_2, .., v_{N-1}$ in Eq. (C39) reduces to a mere multiplication of $\mathbf{T}^{(5)}(\boldsymbol{\nu}_j, \boldsymbol{\nu}_{j+1})$ matrices:

$$\sum_{v_j} \begin{pmatrix} \mathbf{T}_{00}^{(5)} & \mathbf{T}_{01}^{(5)} & 0 & 0 \\ 0 & 0 & \mathbf{T}_{12}^{(5)} & 0 \\ 0 & 0 & 0 & \mathbf{T}_{23}^{(5)} \\ \mathbf{T}_{30}^{(5)} & 0 & 0 & 0 \end{pmatrix} \Bigg|_{v_{j-1} v_j} \times \begin{pmatrix} \mathbf{T}_{00}^{(5)} & \mathbf{T}_{01}^{(5)} & 0 & 0 \\ 0 & 0 & \mathbf{T}_{12}^{(5)} & 0 \\ 0 & 0 & 0 & \mathbf{T}_{23}^{(5)} \\ \mathbf{T}_{30}^{(5)} & 0 & 0 & 0 \end{pmatrix} \Bigg|_{v_j v_{j+1}} = \begin{pmatrix} \mathbf{T}_{00}^{(5)} & \mathbf{T}_{01}^{(5)} & 0 & 0 \\ 0 & 0 & \mathbf{T}_{12}^{(5)} & 0 \\ 0 & 0 & 0 & \mathbf{T}_{23}^{(5)} \\ \mathbf{T}_{30}^{(5)} & 0 & 0 & 0 \end{pmatrix}^2 \Bigg|_{v_{j-1} v_{j+1}} \tag{C40}$$

Where we omitted $\boldsymbol{\nu}_{j-1}$, $\boldsymbol{\nu}_j$ and $\boldsymbol{\nu}_{j+1}$ arguments of matrices $\mathbf{T}_{nm}^{(5)}$ for the sake of formula simplicity.

In other words, Eq. (C40) says that matrix multiplication and $\big|_{v_j v_{j+1}}$ matrix operator commute with each other.

Applying Eq. (C40) $N-2$ times to Eq. (C39), it is not hard to see that the mathematical expression for the DNA partition function takes the following form:

$$Z_{\psi,f} = (2\pi)^3 \begin{pmatrix} 1 & 0 & 0 & 0 \end{pmatrix} \times \sum_{v_1,\, v_N} \int_{\mathbb{R}^{3N}} \mathrm{d}\boldsymbol{\nu}_1 \ldots \mathrm{d}\boldsymbol{\nu}_N \left[ \prod_{j=1}^{N-1} \mathbf{T}^{(5)}(\boldsymbol{\nu}_j, \boldsymbol{\nu}_{j+1}) \right] \Bigg|_{v_1 v_N} \times \boldsymbol{\xi}^{(4)}(\boldsymbol{\nu}_N, \boldsymbol{\nu}_1) \Big|_{v_N v_1} \tag{C41}$$

Finally, after a few simple algebraic rearrangements, Eq. (C41) can be reduced to:

$$Z_{\psi,f} = (2\pi)^3 \int_{\mathbb{R}^{3N}} \mathrm{d}\boldsymbol{\nu}_1 \ldots \mathrm{d}\boldsymbol{\nu}_N \, \mathrm{Tr} \left\{ \mathbf{W} \times \prod_{j=1}^{N-1} \mathbf{T}^{(5)}(\boldsymbol{\nu}_j, \boldsymbol{\nu}_{j+1}) \times \boldsymbol{\xi}^{(4)}(\boldsymbol{\nu}_N, \boldsymbol{\nu}_1) \right\} \tag{C42}$$

Here Tr{...} is the trace of a matrix. $\mathbf{W}$ is a block-matrix, which equals to: $\mathbf{W} = \begin{pmatrix} \mathbf{I} & 0 & 0 & 0 \end{pmatrix}$, where $\mathbf{I}$ is the identity matrix of $(s_{\max}+1)^2 \times (s_{\max}+1)^2$ size (i.e., $[\,\mathbf{I}\,]_{nm} = \delta_{nm}$).

So far, we have considered only the special case of $K = 3$, when upon binding to DNA proteins occupy three DNA segments. However, in the general case, $K$ may have an arbitrary value. For example, histone octamers bind on average to $\sim 147$ bp of DNA during nucleosome formation [37, 38]. This corresponds to $K \approx 14$ DNA segments of $b = 3.4$ nm length used in our study, see Table I.

It is not very hard to generalize the above formulas for any value of $K$ by noting that Eq. (C42), which defines the partition function of DNA, as well as Eq. (C4), Eq. (C6), Eq. (C14), Eq. (C15), Eq. (C34) and Eq. (C35) are independent of $K$, and thus remain valid in the general case. On the other hand, Eq. (C18), Eq. (C36) and Eq. (C37) need to be slightly adjusted by adding a number of new functions / elements / blocks. Specifically, Eq. (C18) changes to:

$$\begin{cases} T_{00}^{(4)}(\mathbf{R}_j, \mathbf{R}_{j+1}, \boldsymbol{\nu}_j, \boldsymbol{\nu}_{j+1}) = T_{00}^{(3)}(\mathbf{R}_j, \mathbf{R}_{j+1}, \boldsymbol{\nu}_j, \boldsymbol{\nu}_{j+1}) \\ T_{01}^{(4)}(\mathbf{R}_j, \mathbf{R}_{j+1}, \boldsymbol{\nu}_j, \boldsymbol{\nu}_{j+1}) = \displaystyle\int \mathrm{d}\mathbf{R}'_1 \ldots \mathbf{R}'_K \, T_{01}^{(2)}(\mathbf{R}_j, \mathbf{R}'_1, \boldsymbol{\nu}_j) \, T_{12}^{(2)}(\mathbf{R}'_1, \mathbf{R}'_2, \boldsymbol{\nu}_j) \times \cdots \times \\ \qquad\qquad \times T_{K-1\,K}^{(2)}(\mathbf{R}'_{K-1}, \mathbf{R}'_K, \boldsymbol{\nu}_j) \, T_{K0}^{(3)}(\mathbf{R}'_K, \mathbf{R}_{j+1}, \boldsymbol{\nu}_j, \boldsymbol{\nu}_{j+1}) \\ T_{12}^{(4)}(\mathbf{R}_j, \mathbf{R}_{j+1}, \boldsymbol{\nu}_j, \boldsymbol{\nu}_{j+1}) = \cdots = T_{K0}^{(4)}(\mathbf{R}_j, \mathbf{R}_{j+1}, \boldsymbol{\nu}_j, \boldsymbol{\nu}_{j+1}) = \delta(\mathbf{R}_j - \mathbf{R}_{j+1}) \, \delta(\boldsymbol{\nu}_j - \boldsymbol{\nu}_{j+1}) \\ \text{All other } T_{nm}^{(4)}(\mathbf{R}_j, \mathbf{R}_{j+1}, \boldsymbol{\nu}_j, \boldsymbol{\nu}_{j+1}) = 0 \end{cases} \tag{C43}$$

As for Eq. (C36), it becomes:

$$\left[\mathbf{T}^{(4)}(\boldsymbol{\nu}_j,\boldsymbol{\nu}_{j+1})\right]^{p_{j+1},\,0,\,s_{j+1}}_{p_j,\,0,\,s_j} = \begin{pmatrix} \left[\mathbf{T}^{(5)}_{00}\right]_{v_j v_{j+1}} & \left[\mathbf{T}^{(5)}_{01}\right]_{v_j v_{j+1}} & 0 & \cdots & 0 \\ 0 & 0 & \left[\mathbf{T}^{(5)}_{12}\right]_{v_j v_{j+1}} & \cdots & 0 \\ \vdots & \vdots & \vdots & \ddots & \vdots \\ 0 & 0 & 0 & \cdots & \left[\mathbf{T}^{(5)}_{K-1\,K}\right]_{v_j v_{j+1}} \\ \left[\mathbf{T}^{(5)}_{K0}\right]_{v_j v_{j+1}} & 0 & 0 & \cdots & 0 \end{pmatrix} = \mathbf{T}^{(5)}(\boldsymbol{\nu}_j,\boldsymbol{\nu}_{j+1})\Big|_{v_j v_{j+1}}$$

$$\left[\boldsymbol{\xi}^{(3)}(\boldsymbol{\nu}_N,\boldsymbol{\nu}_1)\right]^{p_1,\,0,\,s_1}_{p_N,\,0,\,s_N} = \begin{pmatrix} \left[\boldsymbol{\xi}^{(4)}_0(\boldsymbol{\nu}_N,\boldsymbol{\nu}_1)\right]_{v_N v_1} \\ 0 \\ \vdots \\ 0 \end{pmatrix} = \boldsymbol{\xi}^{(4)}(\boldsymbol{\nu}_N,\boldsymbol{\nu}_1)\Big|_{v_N v_1} \tag{C44}$$

Where the above matrix $\mathbf{T}^{(5)}(\boldsymbol{\nu}_j,\boldsymbol{\nu}_{j+1})\big|_{v_j v_{j+1}}$ is composed of $(K+1)\times(K+1)$ elements, and vector $\boldsymbol{\xi}^{(4)}(\boldsymbol{\nu}_N,\boldsymbol{\nu}_1)\big|_{v_N v_1}$ – of $K+1$ elements arranged in a column.

Consequently, it is not hard to see that Eq. (C37) transforms into:

$$\mathbf{T}^{(5)}(\boldsymbol{\nu}_j,\boldsymbol{\nu}_{j+1}) = \begin{pmatrix} \mathbf{T}^{(5)}_{00} & \mathbf{T}^{(5)}_{01} & 0 & \cdots & 0 \\ 0 & 0 & \mathbf{T}^{(5)}_{12} & \cdots & 0 \\ \vdots & \vdots & \vdots & \ddots & \vdots \\ 0 & 0 & 0 & \cdots & \mathbf{T}^{(5)}_{K-1\,K} \\ \mathbf{T}^{(5)}_{K0} & 0 & 0 & \cdots & 0 \end{pmatrix} \text{ and } \boldsymbol{\xi}^{(4)}(\boldsymbol{\nu}_N,\boldsymbol{\nu}_1) = \begin{pmatrix} \boldsymbol{\xi}^{(4)}_0(\boldsymbol{\nu}_N,\boldsymbol{\nu}_1) \\ 0 \\ \vdots \\ 0 \end{pmatrix}, \tag{C45}$$

Where we omitted $(\boldsymbol{\nu}_j,\boldsymbol{\nu}_{j+1})$ arguments of $\mathbf{T}^{(5)}_{nm}(\boldsymbol{\nu}_j,\boldsymbol{\nu}_{j+1})$ matrix-blocks for the sake of formula simplicity.

Finally, matrix $\mathbf{W}$ turns into: $\mathbf{W} = \begin{pmatrix} \mathbf{I} & 0 & 0 & \cdots & 0 \end{pmatrix}$, where the total number of 0 blocks equals to $K$.

As Eq. (C42) holds in the general case, all that we need to do to calculate the DNA partition function is to derive mathematical expressions for the elements of the above matrix-blocks, $\mathbf{T}^{(5)}_{nm}(\boldsymbol{\nu}_j,\boldsymbol{\nu}_{j+1})$ and $\boldsymbol{\xi}^{(4)}_0(\boldsymbol{\nu}_N,\boldsymbol{\nu}_1)$, see Appendices F-G. In addition, we also need to find a way to evaluate $\int_{\mathrm{I\!R}^{3N}} \mathrm{d}\boldsymbol{\nu}_1 ... \mathrm{d}\boldsymbol{\nu}_N$ multiple integral in Eq. (C42), and this is exactly what we are going to discuss in the next section, Appendix D.

#### Appendix D: Dirac comb function

By having in hands Eq. (C42), calculation of the DNA partition function seems to be a rather straightforward procedure if not for the fact that we need to compute $\int_{\mathrm{I\!R}^{3N}} \mathrm{d}\boldsymbol{\nu}_1 ... \mathrm{d}\boldsymbol{\nu}_N$ integral. While direct evaluation of this integral is an impossible task for modern computational systems, we still can obtain an accurate estimation of the DNA partition function by replacing $\int_{\mathrm{I\!R}^{3N}} \mathrm{d}\boldsymbol{\nu}_1 ... \mathrm{d}\boldsymbol{\nu}_N$ integral in Eq. (C42) with a sum. Indeed, $\int \mathrm{d}\boldsymbol{\nu}_j$ integrals were introduced into the partition function by using the following well-known representation of the Dirac $\delta$-function [see Eq. (C7)-Eq. (C13)]:

$$\delta(\mathbf{r}) = \frac{1}{(2\pi)^3} \int_{\mathrm{I\!R}^3} e^{i[\boldsymbol{\omega}\cdot\mathbf{r}]} \mathrm{d}\boldsymbol{\omega} \tag{D1}$$

Where $\mathbf{r}$ is a 3D position vector.

However, from the practical point of view, it is more convenient to work instead with the Dirac comb function,

III($\mathbf{r}$), which is defined as an infinite sum of Dirac $\delta$-functions located at the nodes of a Bravais lattice:

$$\mathrm{III}(\mathbf{r}) = \sum_{\mathbf{n}} \delta(\mathbf{r} + \mathbf{r}_{\mathbf{n}}) \tag{D2}$$

Here $\mathbf{r}_{\mathbf{n}} = n_1\mathbf{a}_1 + n_2\mathbf{a}_2 + n_2\mathbf{a}_2$, $\mathbf{n} = (n_1, n_2, n_3) \in \mathbb{Z}^3$, are the position vectors of the nodes of the Bravais lattice spanned by primitive vectors $\mathbf{a}_1$, $\mathbf{a}_2$ and $\mathbf{a}_3$. $\sum_{\mathbf{n}}$ summation in the above formula is carried out over all possible values of the vector $\mathbf{n}$.

From the first look at Eq. (D2) it seems that by using the Dirac comb functions, we make things much more complex than in the original case as we have to consider now an infinite number of Dirac $\delta$-functions instead of a single one. However, this is rather deceptive impression as it can be shown that by properly selecting the length of the primitive vectors, $(\mathbf{a}_1, \mathbf{a}_2, \mathbf{a}_3)$, we can achieve a situation where for each position vector, $\mathbf{r}$, inside the cell nucleus (i.e., $\mathbf{r} \in B_{R_{\mathrm{nucl}}}$) there is exactly one Dirac $\delta$-function, which is located at the point corresponding to this vector, with the rest of $\delta$-functions from Eq. (D2) sitting outside of the cell nucleus. For example, in this study we utilize a hexagonal Bravais lattice with the lengths of the primitive vectors being slightly larger than two radii of the cell nucleus ($R_{\mathrm{nucl}}$): $\|\mathbf{a}_1\| = \|\mathbf{a}_2\| = \|\mathbf{a}_3\| = 2.4\,R_{\mathrm{nucl}}$, such that the cell nucleus can be completely covered by a single primitive unit cell of the Bravais lattice as schematically shown in Figure S10(a). It can be easily checked that in this case for each position vector, $\mathbf{r}$, in the cell nucleus there is a unique Dirac $\delta$-function located at the point corresponding to it; whereas, all other $\delta$-functions from Eq. (D2) lie outside of the cell nucleus. Thus, from a physical point of view, there is no difference whether to use a single Dirac $\delta$-function in Eq. (C7) or the Dirac comb function composed of an infinite number of Dirac $\delta$-functions.

What is more important, it can be shown that by using the Dirac comb function, we do not need to deal anymore with the multiple integral from Eq. (C42) as it becomes replaced by a sum. Indeed, from the definition of III($\mathbf{r}$) it follows that this is a periodic function: $\mathrm{III}(\mathbf{r} + \mathbf{r}_{\mathbf{n}'}) = \mathrm{III}(\mathbf{r})$, $\forall \mathbf{n}' \in \mathbb{Z}^3$ (i.e., the Dirac comb function is periodic on the Bravais lattice). Thus, by expanding it into Fourier series, we get:

$$\mathrm{III}(\mathbf{r}) = \sum_{\mathbf{n}} \delta(\mathbf{r} + \mathbf{r}_{\mathbf{n}}) = \frac{1}{V_{\mathrm{B}}} \sum_{\mathbf{m}} e^{i[\mathbf{G}_{\mathbf{m}} \cdot \mathbf{r}]} \tag{D3}$$

Where $V_{\mathrm{B}} = \mathbf{a}_1 \cdot [\mathbf{a}_2 \times \mathbf{a}_3]$ is the volume of a primitive unit cell of the Bravais lattice. $\mathbf{G}_{\mathbf{m}} = m_1\mathbf{b}_1 + m_2\mathbf{b}_2 + m_3\mathbf{b}_3$, $\mathbf{m} = (m_1, m_2, m_3) \in \mathbb{Z}^3$, is an array of points in 3D space describing the nodes of so-called reciprocal lattice, which is spanned by the following primitive vectors:

$$\mathbf{b}_1 = \frac{2\pi}{V_{\mathrm{B}}}[\mathbf{a}_2 \times \mathbf{a}_3] \quad \text{and} \quad \mathbf{b}_2 = \frac{2\pi}{V_{\mathrm{B}}}[\mathbf{a}_3 \times \mathbf{a}_1] \quad \text{and} \quad \mathbf{b}_3 = \frac{2\pi}{V_{\mathrm{B}}}[\mathbf{a}_1 \times \mathbf{a}_2] \tag{D4}$$

By comparing Eq. (D1) to Eq. (D3), it can be seen that Eq. (D3) is just a discrete analogue of the integral representation of the Dirac $\delta$-function given by Eq. (D1), which, however, has an advantage over the latter as it contains a sum instead of the integral. By substituting Eq. (D3) into Eq. (C7) instead of the integral representation of the Dirac $\delta$-function [Eq. (D1)], it is straightforward to see that:

$$\begin{aligned} e^{-\beta\hat{\psi}_{k_j k_{j+1}}(\mathbf{r}_j)} &= \int_{\mathrm{P.C.}} \mathrm{d}\mathbf{r}'_j \, e^{-\beta\hat{\psi}_{k_j k_{j+1}}(\mathbf{r}'_j)} \, \mathrm{III}\Big(\mathbf{r}_0 + \sum_{n=1}^{j} b_{k_n} \mathbf{R}_n \mathbf{z}_0 - \mathbf{r}'_j\Big) = \\ &= \frac{1}{V_{\mathrm{B}}} \sum_{\mathbf{m}_j} \int_{\mathrm{P.C.}} \mathrm{d}\mathbf{r}'_j \, e^{-\beta\hat{\psi}_{k_j k_{j+1}}(\mathbf{r}'_j)} \, e^{-i[\mathbf{G}_{\mathbf{m}_j} \cdot \mathbf{r}'_j] + i[\mathbf{G}_{\mathbf{m}_j} \cdot \sum_{n=1}^{j} b_{k_n} \mathbf{R}_n \mathbf{z}_0] + i[\mathbf{G}_{\mathbf{m}_j} \cdot \mathbf{r}_0]} \end{aligned} \tag{D5}$$

Here $\int_{\mathrm{P.C.}} \mathrm{d}\mathbf{r}'_j$ integration is carried out over a primitive unit cell (P.C.) of the Bravais lattice. In the case of an earlier discussed hexagonal lattice, one can simply use the primitive unit cell schematically shown in Figure S10(a), which covers the entire cell nucleus.

In addition to Eq. (D5), it is not hard to obtain the following useful expression for a discrete version of the Dirac

$\delta$-function:

$$\delta^{3D}_{\mathbf{m}0} = \frac{1}{V_B} \int_{P.C.} d\mathbf{r}\, e^{-i[\mathbf{G_m} \cdot \mathbf{r}]} \tag{D6}$$

Where $\delta^{3D}_{\mathbf{mm}'}$ is a 3D Kronecker delta: $\delta^{3D}_{\mathbf{mm}'} = \delta_{m_1 m'_1}\delta_{m_2 m'_2}\delta_{m_3 m'_3} = 1$ if $\mathbf{m} = (m_1, m_2, m_3) = (m'_1, m'_2, m'_3) = \mathbf{m}'$ and $\delta^{3D}_{\mathbf{mm}'} = 0$, otherwise.

As a result, after substituting Eq. (D5) and Eq. (D6) into Eq. (C5), one can get the following formula for the DNA partition function, which is analogous to Eq. (C12):

$$Z_{\psi,f} = V_B \sum_{k_1...k_N=0}^{K} \delta_{k_1 0} \times \sum_{\mathbf{m}_1...\mathbf{m}_N} \int d\mathbf{R}_1...d\mathbf{R}_N \prod_{j=1}^{N-1} \left[ T^{(1)}_{k_j k_{j+1}}(\mathbf{R}_j, \mathbf{R}_{j+1})\, \Phi_{k_j k_{j+1}}(\mathbf{G}_{\mathbf{m}_j} - \mathbf{G}_{\mathbf{m}_{j+1}})\, e^{ib_{k_j}[\mathbf{R}_j \mathbf{z}_0 \cdot \mathbf{G}_{\mathbf{m}_j}]} \right] \times$$
$$\times\, \xi^{(1)}_{k_N}(\mathbf{R}_N, \mathbf{R}_1)\, \delta^{3D}_{\mathbf{m}_1 0}\, \Phi_{00}(\mathbf{G}_{\mathbf{m}_N})\, e^{ib[\mathbf{R}_N \mathbf{z}_0 \cdot \mathbf{G}_{\mathbf{m}_N}]}, \tag{D7}$$

Where this time $\Phi_{k_j k_{j+1}}(\mathbf{G}_{\mathbf{m}_j})$ function is defined as:

$$\Phi_{k_j k_{j+1}}(\mathbf{G}_{\mathbf{m}_j}) = \frac{1}{V_B} \int_{P.C.} d\mathbf{r}'_j\, e^{-\beta \hat{\psi}_{k_j k_{j+1}}(\mathbf{r}'_j)}\, e^{-i[\mathbf{G}_{\mathbf{m}_j} \cdot \mathbf{r}'_j]} \tag{D8}$$

Then, by following all of the derivations described in Appendix C, it can be shown that the final Eq. (C42) for the DNA partition function changes to:

$$Z_{\psi,f} = V_B \sum_{\mathbf{m}_1...\mathbf{m}_N} \mathrm{Tr}\left\{ \mathbf{W} \times \prod_{j=1}^{N-1} \mathbf{T}^{(5)}(\mathbf{G}_{\mathbf{m}_j}, \mathbf{G}_{\mathbf{m}_{j+1}}) \times \boldsymbol{\xi}^{(4)}(\mathbf{G}_{\mathbf{m}_N}, \mathbf{G}_{\mathbf{m}_1}) \right\} \tag{D9}$$

Here all matrices retain their original definition with the only difference being replacement of $\boldsymbol{\nu}_j$ vectors by $\mathbf{G}_{\mathbf{m}_j}$ vectors and $\delta(\boldsymbol{\nu}_j - \boldsymbol{\nu}_{j+1})$ functions by $\delta^{3D}_{\mathbf{m}_j \mathbf{m}_{j+1}}$ Kronecker symbols in the corresponding formulas, such as Eq. (C43).

From Eq. (D9) it can be seen that in order to calculate the DNA partition function all we need to do is to find $\sum_{\mathbf{m}_1...\mathbf{m}_N}$ sum, which is a much more simpler task then computation of $\int_{\mathbb{R}^{3N}} d\boldsymbol{\nu}_1...d\boldsymbol{\nu}_N$ multiple integral in Eq. (C42). The main reason why did not use the Dirac comb function in our derivations from the very beginning is that later we will need both Eq. (C42) and Eq. (D9) in order to construct an efficient algorithm for calculation of the DNA partition function, see Appendix H.

In any case, replacement of the multiple integral in Eq. (C42) by a sum makes it possible to further simplify Eq. (D9) by applying the same technique as the one described in Appendix C, which is based on the construction of higher-level matrices.

To this aim, the first thing we need to do is to enumerate $\mathbf{m}_j$ vectors by using a new index, $l_j = 1, 2, ..., l_{\max}$: $\mathbf{m}_j = \mathbf{m}_j(l_j)$, where $l_{\max}$ is the total number of the reciprocal lattice nodes used in evaluation of the DNA partition function. One of the possible examples of how this can be done is shown schematically in Figure S10(b). Numeric calculations indicate that $l_{\max}$ does not have a strong effect on the DNA partition function value as soon as $l_{\max} \gtrsim 2500$, see Figure S8(b). As for the rest of physical quantities investigated in our study, some of them may have a slightly higher requirement to the total number of the reciprocal lattice nodes, which, however, can be easily fulfilled by modern computational systems without much of a problem in most of the cases, see, for example, Appendix J.

After enumerating $\mathbf{m}_j$ vectors, it is clear that Eq. (D9) can be represented in the following form:

$$Z_{\psi,f} = V_B \sum_{l_1...l_N} \mathrm{Tr}\left\{ \mathbf{W} \times \prod_{j=1}^{N-1} \mathbf{T}^{(5)}(l_j, l_{j+1}) \times \boldsymbol{\xi}^{(4)}(l_N, l_1) \right\} \tag{D10}$$

Where $\mathbf{T}^{(5)}(l_j, l_{j+1}) = \mathbf{T}^{(5)}(\mathbf{G}_{\mathbf{m}_j(l_j)}, \mathbf{G}_{\mathbf{m}_{j+1}(l_{j+1})})$ and $\boldsymbol{\xi}^{(4)}(l_N, l_1) = \boldsymbol{\xi}^{(4)}(\mathbf{G}_{\mathbf{m}_N(l_N)}, \mathbf{G}_{\mathbf{m}_1(l_1)})$.

At this point construction of the higher-level transfer-matrix, $\mathbf{L}$, becomes rather trivial – by using $\mathbf{T}^{(5)}_{nm}(l_j, l_{j+1})$

matrix-blocks that comprise $\mathbf{T}^{(5)}(l_j, l_{j+1})$ transfer-matrix, we can define the next level blocks, $\mathbf{T}^{(6)}_{nm}$, as:

$$\mathbf{T}^{(6)}_{nm} = \begin{pmatrix} \mathbf{T}^{(5)}_{nm}(1,1) & \cdots\ \cdots & \mathbf{T}^{(5)}_{nm}(1,l_{\max}) \\ \vdots & \ddots & \vdots \\ \vdots & \ddots & \vdots \\ \mathbf{T}^{(5)}_{nm}(l_{\max},1) & \cdots\ \cdots & \mathbf{T}^{(5)}_{nm}(l_{\max},l_{\max}) \end{pmatrix} \tag{D11}$$

Here $\mathbf{T}^{(5)}_{nm}(l_j, l_{j+1}) = \mathbf{T}^{(5)}_{nm}(\mathbf{G}_{\mathbf{m}_j(l_j)}, \mathbf{G}_{\mathbf{m}_{j+1}(l_{j+1})})$.

Then we can assemble the final transfer-matrix, $\mathbf{L}$, by following the same pattern as in Eq. (C45):

$$\mathbf{L} = \begin{pmatrix} \mathbf{T}^{(6)}_{00} & \mathbf{T}^{(6)}_{01} & 0 & \cdots & 0 \\ 0 & 0 & \mathbf{T}^{(6)}_{12} & \cdots & 0 \\ \vdots & \vdots & \vdots & \ddots & \vdots \\ 0 & 0 & 0 & \cdots & \mathbf{T}^{(6)}_{K-1\,K} \\ \mathbf{T}^{(6)}_{K0} & 0 & 0 & \cdots & 0 \end{pmatrix} = \begin{pmatrix} \mathbf{T}^{(6)}_{00} & \mathbf{T}^{(6)}_{01} & 0 & \cdots & 0 \\ 0 & 0 & \mathbf{I} & \cdots & 0 \\ \vdots & \vdots & \vdots & \ddots & \vdots \\ 0 & 0 & 0 & \cdots & \mathbf{I} \\ \mathbf{I} & 0 & 0 & \cdots & 0 \end{pmatrix} \tag{D12}$$

Here we have taken into account that from Eq. (D11) and Eq. (G45) it follows that $\mathbf{T}^{(6)}_{12} = \mathbf{T}^{(6)}_{23} = ... = \mathbf{T}^{(6)}_{K0} = \mathbf{I}$, where the identity matrix, $\mathbf{I}$, has the same size as $\mathbf{T}^{(6)}_{00}$ and $\mathbf{T}^{(6)}_{01}$ matrix-blocks: $l_{\max}(s_{\max}+1)^2 \times l_{\max}(s_{\max}+1)^2$.

As for the boundary condition matrix, $\mathbf{P}$, it is defined in a very similar way to the transfer-matrix, $\mathbf{L}$:

$$\mathbf{P} = \begin{pmatrix} \boldsymbol{\xi}^{(5)}_0 \\ 0 \\ 0 \\ \vdots \\ 0 \end{pmatrix}, \quad \text{where} \quad \boldsymbol{\xi}^{(5)}_0 = \begin{pmatrix} \boldsymbol{\xi}^{(4)}_0(1,1) & \cdots\ \cdots & \boldsymbol{\xi}^{(4)}_0(1,l_{\max}) \\ \vdots & \ddots & \vdots \\ \vdots & \ddots & \vdots \\ \boldsymbol{\xi}^{(4)}_0(l_{\max},1) & \cdots\ \cdots & \boldsymbol{\xi}^{(4)}_0(l_{\max},l_{\max}) \end{pmatrix} \tag{D13}$$

Where, as before, $\boldsymbol{\xi}^{(4)}_0(l_N, l_1) = \boldsymbol{\xi}^{(4)}_0(\mathbf{G}_{\mathbf{m}_N(l_N)}, \mathbf{G}_{\mathbf{m}_1(l_1)})$.

Combining together Eq. (C45), Eq. (D11) and Eq. (D12), it is not hard to see that:

$$\mathbf{L}\Big|_{l_j l_{j+1}} = \left.\begin{pmatrix} \mathbf{T}^{(6)}_{00} & \mathbf{T}^{(6)}_{01} & 0 & \cdots & 0 \\ 0 & 0 & \mathbf{T}^{(6)}_{12} & \cdots & 0 \\ \vdots & \vdots & \vdots & \ddots & \vdots \\ 0 & 0 & 0 & \cdots & \mathbf{T}^{(6)}_{K-1\,K} \\ \mathbf{T}^{(6)}_{K0} & 0 & 0 & \cdots & 0 \end{pmatrix}\right|_{l_j l_{j+1}} =$$

$$= \begin{pmatrix} \mathbf{T}^{(5)}_{00}(l_j,l_{j+1}) & \mathbf{T}^{(5)}_{01}(l_j,l_{j+1}) & 0 & \cdots & 0 \\ 0 & 0 & \mathbf{T}^{(5)}_{12}(l_j,l_{j+1}) & \cdots & 0 \\ \vdots & \vdots & \vdots & \ddots & \vdots \\ 0 & 0 & 0 & \cdots & \mathbf{T}^{(5)}_{K-1\,K}(l_j,l_{j+1}) \\ \mathbf{T}^{(5)}_{K0}(l_j,l_{j+1}) & 0 & 0 & \cdots & 0 \end{pmatrix} = \mathbf{T}^{(5)}(l_j, l_{j+1}) \tag{D14}$$

Analogously, from Eq. (C45) and Eq. (D13) we have: $\mathbf{P}\Big|_{l_N l_1} = \boldsymbol{\xi}^{(4)}(l_N, l_1)$. By substituting these two formulas into Eq. (D10) and by using the same sequence of the DNA partition function transformations as the one described at the end of Appendix C [see Eq. (C39)-Eq. (C42)], it is straightforward to obtain the following expression for the DNA partition function:

$$Z_{\psi,f} = V_{\rm B} \sum_{l_1 \dots l_N} \mathrm{Tr}\left\{ \mathbf{W} \times \prod_{j=1}^{N-1} \mathbf{L}\big|_{l_j l_{j+1}} \times \mathbf{P}\big|_{l_N l_1} \right\} = V_{\rm B} \sum_{l_1, l_N} \mathrm{Tr}\left\{ \mathbf{W} \times \mathbf{L}^{N-1}\big|_{l_1 l_N} \times \mathbf{P}\big|_{l_N l_1} \right\} = V_{\rm B}\, \mathrm{Tr}\left\{ \mathbf{S}\mathbf{L}^{N-1}\mathbf{P} \right\} \quad \text{(D15)}$$

Here $\mathbf{S} = \begin{pmatrix} \mathbf{I} & 0 & 0 & \cdots & 0 \end{pmatrix}$ is a matrix composed of a single identity matrix, $\mathbf{I}$, and $K$ zero blocks, which all have the same size as $\mathbf{T}^{(6)}_{nm}$ matrices.

Eq. (D15) can be further simplified by noting that due to the presence of $\delta_{s'0}$, $\delta_{p'0}$ and $\delta(\boldsymbol{\nu}_1)$ terms in Eq. (G56), only a single column of matrix $\mathbf{P}$ will contain non-zero elements – the one comprised of the first columns of $\mathbf{G}_{\mathbf{m}_1} = 0$ matrix blocks; whereas, all other columns will be composed of zero entries. Thus, it makes sense to use in calculations only this single column instead of the whole boundary matrix, $\mathbf{P}$, which allows one to considerably improve the rate of the DNA partition function computation.

Let $n_0$ be the index of the column containing non-zero elements in the boundary condition matrix, $\mathbf{P}$. Then it is not hard to see that Eq. (D15) is equivalent to the following one:

$$Z_{\psi,f} = V_{\rm B}\, \mathbf{U}\mathbf{L}^{N-1}\mathbf{Y}, \quad \text{(D16)}$$

Where $\mathbf{U}$ and $\mathbf{Y}$ vectors are:

$$\mathbf{U} = \mathbf{S}\big|^{n_0}_{\rm row} = \begin{pmatrix} 0 & \cdots & 0 & 1 & 0 & \cdots & 0 \end{pmatrix} \quad \text{and} \quad \mathbf{Y} = \begin{pmatrix} \boldsymbol{\xi}^{(5)}_0\big|^{n_0}_{\rm column} \\ 0 \\ 0 \\ \vdots \\ 0 \end{pmatrix} \quad \text{(D17)}$$

Here $\mathbf{S}\big|^{n_0}_{\rm row}$ and $\boldsymbol{\xi}^{(5)}_0\big|^{n_0}_{\rm column}$ are the $n_0^{\rm th}$ row and column of $\mathbf{S}$ and $\boldsymbol{\xi}^{(5)}_0$ matrices, respectively.

Comparing Eq. (D16) to Eq. (C42), it is easy to see that by using the Dirac comb function we have reduced the DNA partition function calculation to a pure matrix multiplication procedure, which is at the core of all modern GPU computational systems, making our theory well-adapted to them. Furthermore, Eq. (D17) allows one to easily find the functional derivative, $\frac{\delta \ln Z_\psi}{\delta \psi(r)}$, which is needed to solve Eq. (A30). This is the goal of the next section, Appendix E.

#### Appendix E: $\frac{\delta \ln Z_\psi}{\delta \psi(r)}$ functional derivative

Having at hand Eq. (D16) for the partition function of DNA, we can now derive a mathematical expression for $\frac{\delta \ln Z_\psi}{\delta \psi(r)}$ functional derivative.

To this aim, it should be noted that in the case of $Q$ chromosomes constrained inside the cell nucleus, Eq. (A19) and Eq. (D16) transform into:

$$Z_\psi = \prod_{u=1}^{Q} \left[ V_{\rm B} \mathbf{U}\mathbf{L}^{N_u - 1}\mathbf{Y} \right] \quad \text{(E1)}$$

Here $N_u = \lfloor L_u / b \rfloor$ is the total number of DNA segments in the $u^{\rm th}$ chromosome. $\mathbf{L}$, $\mathbf{U}$ and $\mathbf{Y}$ are the transfer-matrix and the boundary condition vectors, which are defined in Appendix D.

By applying logarithm to the both sides of Eq. (E1) and taking $\frac{\delta}{\delta \psi(r)}$ functional derivative, it is not hard to see that:

$$\frac{\delta \ln Z_\psi}{\delta \psi(r)} = \sum_{u=1}^{Q} \frac{\frac{\delta}{\delta \psi(r)} \left[ \mathbf{U}\mathbf{L}^{N_u-1}\mathbf{Y} \right]}{\mathbf{U}\mathbf{L}^{N_u-1}\mathbf{Y}} = \sum_{u=1}^{Q} \frac{\sum_{n=0}^{N_u-2} \left[ \mathbf{U}\mathbf{L}^{n} \times \frac{\delta}{\delta \psi(r)} \mathbf{L} \times \mathbf{L}^{N_u-n-2}\mathbf{Y} \right] + \mathbf{U}\mathbf{L}^{N_u-1} \times \frac{\delta}{\delta \psi(r)} \mathbf{Y}}{\mathbf{U}\mathbf{L}^{N_u-1}\mathbf{Y}} \quad \text{(E2)}$$

Where $\frac{\delta}{\delta\psi(r)}\mathbf{L}$ and $\frac{\delta}{\delta\psi(r)}\mathbf{Y}$ designate matrices whose entries are functional derivatives of the respective elements of the matrices $\mathbf{L}$ and $\mathbf{Y}$.

By using the Jordan decomposition of the transfer-matrix, $\mathbf{L}$, and following the same formula derivation as in the proof of the power iteration method (also known as the power method) [39], it is possible to greatly simplify Eq. (E2). Specifically, one can show that the partition function of chromosomal DNA with a good accuracy equals to $Z_\psi \approx C_0 \lambda_{\max}^{N_{\rm tot}}$. Here $N_{\rm tot} = \sum_{u=1}^{Q} N_u$ is the total number of DNA segments in chromosomes; $C_0$ is a positive constant, which is independent from $N_{\rm tot}$ as soon as $N_u \gg 1$, $\forall u = 1, ..., Q$; and $\lambda_{\max}$ is the dominant eigenvalue of the transfer-matrix, $\mathbf{L}$. Since the value of the partition function is a positive real number, it follows from this equation that $\lambda_{\max}$ is also a positive real number. Based on this result, it then can be demonstrated that:

$$\lim_{n\to\infty} \frac{\mathbf{L}^n\mathbf{Y}}{\|\mathbf{L}^n\mathbf{Y}\|} = \frac{\mathrm{Pr}^{\rm R}_{\lambda_{\max}}\mathbf{Y}}{\|\mathrm{Pr}^{\rm R}_{\lambda_{\max}}\mathbf{Y}\|} \quad \text{and} \quad \lim_{n\to\infty} \frac{\mathbf{U}\mathbf{L}^n}{\|\mathbf{U}\mathbf{L}^n\|} = \frac{\mathrm{Pr}^{\rm L}_{\lambda_{\max}}\mathbf{U}}{\|\mathrm{Pr}^{\rm L}_{\lambda_{\max}}\mathbf{U}\|} \tag{E3}$$

Here $\mathrm{Pr}^{\rm R}_{\lambda_{\max}}\mathbf{Y}$ and $\mathrm{Pr}^{\rm L}_{\lambda_{\max}}\mathbf{U}$ are the projections of $\mathbf{U}$ and $\mathbf{Y}$ vectors onto the right and the left eigenspaces of the transfer-matrix, $\mathbf{L}$, respectively, which correspond to the dominant eigenvalue, $\lambda_{\max}$.

From Eq. (E3) it is then easy to see that in the case of a sufficiently large $n \in \mathbb{N}$:

$$\mathbf{L}\left(\mathbf{L}^n\mathbf{Y}\right) = \mathbf{L}^{n+1}\mathbf{Y} \approx \lambda_{\max}\mathbf{L}^n\mathbf{Y} \quad \text{and} \quad \left(\mathbf{U}\mathbf{L}^n\right)\mathbf{L} = \mathbf{U}\mathbf{L}^{n+1} \approx \lambda_{\max}\mathbf{U}\mathbf{L}^n \tag{E4}$$

Indeed, numeric calculations indicate that in the case of the bare DNA segments' length $b = 3.4$ nm, the deviation of $\frac{1}{\lambda_{\max}}\frac{\|\mathbf{L}^{n+1}\mathbf{Y}\|}{\|\mathbf{L}^n\mathbf{Y}\|}$ and $\frac{1}{\lambda_{\max}}\frac{\|\mathbf{U}\mathbf{L}^{n+1}\|}{\|\mathbf{U}\mathbf{L}^n\|}$ from the unit is $< 10^{-5}$ after $n = 200 - 8000$ matrix multiplications. Thus, by taking into account Eq. (E4) and neglecting the end effects, which can be safely done for long DNA molecules, we can rewrite Eq. (E2) in the following simpler form:

$$\frac{\delta \ln Z_\psi}{\delta\psi(r)} \approx \frac{N_{\rm tot}}{\lambda_{\max}} \frac{\mathrm{Pr}^{\rm L}_{\lambda_{\max}}\mathbf{U} \times \frac{\delta}{\delta\psi(r)}\mathbf{L} \times \mathrm{Pr}^{\rm R}_{\lambda_{\max}}\mathbf{Y}}{\mathrm{Pr}^{\rm L}_{\lambda_{\max}}\mathbf{U} \times \mathrm{Pr}^{\rm R}_{\lambda_{\max}}\mathbf{Y}} \tag{E5}$$

The value of $\lambda_{\max}$ as well as the projections of $\mathbf{U}$ and $\mathbf{Y}$ vectors on the corresponding eigenspaces can be easily found by using the aforementioned power iteration method, whose central idea is expressed by Eq. (E3) and Eq. (E4).

To complete Eq. (E5), all that remains is to find a formula for $\frac{\delta\mathbf{L}}{\delta\psi(r)}$ functional derivative of the DNA transfer-matrix, $\mathbf{L}$. This can be easily done by noting that the only way the elements of matrix $\mathbf{L}$ may depend on the potential field $\psi$ is either through the protein binding free energy to DNA, $\mu_{\rm pr}$ [see Eq. (B8)], or via $\Phi_{00}$ and $\Phi_{K0}$ functions defined by Eq. (D8), where $\hat{\psi}_{k_j k_{j+1}}(\mathbf{r}_j) = \sigma_{k_j k_{j+1}} U_{R_{\rm nucl}}(\mathbf{r}_j) + q_{k_j k_{j+1}}\psi(\mathbf{r}_j)$ is the net action of $U_{R_{\rm nucl}}$ and $\psi$ potential fields on the $j^{\rm th}$ DNA segment. Namely, from Eq. (C14), Eq. (C43), Eq. (C44), Eq. (D11) and Eq. (D12) it can be seen that each entry of the matrix $\mathbf{L}$ has at most one $\Phi_{k_j k_{j+1}}$ function as a multiplier factor. Furthermore, Eq. (C16) and notes after Eq. (D9) imply that only $\Phi_{00}$ and $\Phi_{K0}$ functions will have non-zero functional derivatives with respect to $\psi(r)$ field. Indeed, in the case of $k_{j+1} \neq 0$, from Eq. (D6) and Eq. (D8) we have:

$$\Phi_{k_j k_{j+1}}(\mathbf{G}_{\mathbf{m}_j}) = \frac{1}{V_{\rm B}} \int_{\rm P.C.} \mathrm{d}\mathbf{r}'_j\, e^{-\beta\hat{\psi}_{k_j k_{j+1}}(\mathbf{r}'_j)}\, e^{-i[\mathbf{G}_{\mathbf{m}_j}\cdot\mathbf{r}'_j]} = \left[k_{j+1} \neq 0 \to \hat{\psi}_{k_j k_{j+1}} = 0\right] = \frac{1}{V_{\rm B}} \int_{\rm P.C.} \mathrm{d}\mathbf{r}'_j\, e^{-i[\mathbf{G}_{\mathbf{m}_j}\cdot\mathbf{r}'_j]} = \delta^{\rm 3D}_{\mathbf{m}_j 0}, \tag{E6}$$

and it immediately follows that $\frac{\delta\Phi_{k_j k_{j+1}}}{\delta\psi(r)} = 0$.

In addition, by generalizing Eq. (B17) to the case of an arbitrary value of $K$ and by using Eq. (C4) and Eq. (C15), it becomes clear that $\mu_{\rm pr}$ enters Eq. (E1) only in the form of $e^{\beta\mu_{\rm pr}}$ prefactor in $T^{(2)}_{01}$ transfer-function. Thus, to construct $\frac{\delta\mathbf{L}}{\delta\psi(r)}$ matrix, all we need to do is to repeat the same assembly steps as in the case of the transfer-matrix, $\mathbf{L}$, – starting from Eq. (C4) to Eq. (D12), only this time $\Phi_{00}$ and $\Phi_{K0}$ functions must be replaced in Eq. (C14) with $\frac{\delta\Phi_{00}}{\delta\psi(r)}$ and $\frac{\delta\Phi_{K0}}{\delta\psi(r)} + \beta\Phi_{K0}\frac{\delta\mu_{\rm pr}}{\delta\psi(r)}$ functional derivatives, respectively. As for the rest of $\Phi_{k_j k_{j+1}}$ functions, they must be simply put equal to zero.

Altogether, the above notes lead us to the following formula for the functional derivative of the transfer-matrix, $\mathbf{L}$:

$$\frac{\delta}{\delta\psi(r)}\mathbf{L} = \begin{pmatrix} \frac{\delta}{\delta\psi(r)}\mathbf{T}^{(6)}_{00} & \frac{\delta}{\delta\psi(r)}\mathbf{T}^{(6)}_{01} & 0 & \cdots & 0 \\ 0 & 0 & 0 & \cdots & 0 \\ \vdots & \vdots & \vdots & \ddots & \vdots \\ 0 & 0 & 0 & \cdots & 0 \\ 0 & 0 & 0 & \cdots & 0 \end{pmatrix} \tag{E7}$$

Where matrices $\frac{\delta}{\delta\psi(r)}\mathbf{T}^{(6)}_{00}$ and $\frac{\delta}{\delta\psi(r)}\mathbf{T}^{(6)}_{01}$ are constructed in the same way as matrices $\mathbf{T}^{(6)}_{00}$ and $\mathbf{T}^{(6)}_{01}$ with the only difference being replacement of $\Phi_{00}$ and $\Phi_{K0}$ functions by $\frac{\delta\Phi_{00}}{\delta\psi(r)}$ and $\frac{\delta\Phi_{K0}}{\delta\psi(r)} + \beta\Phi_{K0}\frac{\delta\mu_{\mathrm{pr}}}{\delta\psi(r)}$ functional derivatives.

To find these derivatives, the first thing we need to do is to rewrite Eq. (D8) for $\Phi_{00}$ and $\Phi_{K0}$ functions in terms of a spherically symmetric potential, $\psi(r)$:

$$\begin{aligned}\Phi_{k_j0}\big(\mathbf{G}_{\mathbf{m}_j}\big) &= \frac{1}{V_{\mathrm{B}}}\int_{\mathrm{P.C.}} \mathrm{d}\mathbf{r}'_j\, e^{-\beta\hat{\psi}_{k_j0}(\mathbf{r}'_j)}\, e^{-i[\mathbf{G}_{\mathbf{m}_j}\cdot\mathbf{r}'_j]} = \frac{1}{V_{\mathrm{B}}}\int_{V_{\mathrm{nucl}}} \mathrm{d}\mathbf{r}'_j\, e^{-\beta q_{k_j0}\psi(\mathbf{r}'_j)}\, e^{-i[\mathbf{G}_{\mathbf{m}_j}\cdot\mathbf{r}'_j]} = \\ &= \frac{1}{V_{\mathrm{B}}}\int_0^{2\pi}\mathrm{d}\varphi'_j\int_0^{\pi}\sin\theta'_j\,\mathrm{d}\theta'_j\int_0^{R_{\mathrm{nucl}}}\big(r'_j\big)^2\, e^{-\beta q_{k_j0}\psi(r'_j)}\, e^{-iG_{\mathbf{m}_j}r'_j\cos\theta'_j}\,\mathrm{d}r'_j = \Big[u=\cos\theta'_j\Big] = \\ &= \frac{2\pi}{V_{\mathrm{B}}}\int_{-1}^{1}\mathrm{d}u\int_0^{R_{\mathrm{nucl}}}\big(r'_j\big)^2\, e^{-\beta q_{k_j0}\psi(r'_j)}\, e^{-iG_{\mathbf{m}_j}r'_j u}\,\mathrm{d}r'_j = \frac{4\pi}{V_{\mathrm{B}}G_{\mathbf{m}_j}}\int_0^{R_{\mathrm{nucl}}} r'_j\, e^{-\beta q_{k_j0}\psi(r'_j)}\times \\ &\times \sin\big(G_{\mathbf{m}_j}r'_j\big)\,\mathrm{d}r'_j = \frac{4\pi}{V_{\mathrm{B}}}\int_0^{R_{\mathrm{nucl}}}\big(r'_j\big)^2\, e^{-\beta q_{k_j0}\psi(r'_j)}\, j_0\big(G_{\mathbf{m}_j}r'_j\big)\,\mathrm{d}r'_j\end{aligned} \tag{E8}$$

Here $k_j = 0$ or $k_j = K$. $G_{\mathbf{m}_j} = \|\mathbf{G}_{\mathbf{m}_j}\|$ is the length of the vector $\mathbf{G}_{\mathbf{m}_j}$. $r'_j$, $\theta'_j$ and $\varphi'_j$ are the spherical coordinates of a coordinate system, which is selected in such a way that its $\mathbf{z}$-axis is collinear to $\mathbf{G}_{\mathbf{m}_j}$ wave-vector. $j_0(x) = \frac{\sin x}{x}$ is a spherical Bessel function of the first kind.

By applying $\frac{\delta}{\delta\psi(r)}$ functional derivative to Eq. (E6) and Eq. (E8), it is not hard to see that in the general case we have:

$$\frac{\delta\Phi_{k_jk_{j+1}}}{\delta\psi(r)} = \begin{cases} -\dfrac{4\pi}{V_{\mathrm{B}}}\beta r^2 q_{k_jk_{j+1}}e^{-\beta q_{k_jk_{j+1}}\psi(r)}\, j_0\big(G_{\mathbf{m}_j}r\big)\,, & \text{if } (k_j,k_{j+1}) = (0,0) \text{ or } (K,0), \text{ and } r < R_{\mathrm{nucl}} \\ 0 & , \text{ otherwise}\end{cases} \tag{E9}$$

As for the functional derivative of the protein binding free energy to DNA, $\mu_{\mathrm{pr}}$, from Eq. (B7) it follows that:

$$\frac{\delta\mu_{\mathrm{pr}}}{\delta\psi(r)} = 2k_{\mathrm{B}}T\frac{\delta}{\delta\psi(r)}\Big[\ln Z^{\psi}_{\mathrm{c_1u}} + \ln Z^{\psi}_{\mathrm{c_2u}} - \ln Z^{\psi}_{\mathrm{c_1b}} - \ln Z^{\psi}_{\mathrm{c_2b}}\Big] \tag{E10}$$

Substituting Eq. (A29) into the above expression and taking into account that outside of the cell nucleus the electrostatic potential $\psi$ drops to zero (since the value of the cytoplasmic electrostatic potential is used as a reference point), it is straightforward to obtain the following formula for $\frac{\delta\mu_{\mathrm{pr}}}{\delta\psi(r)}$ derivative:

$$\frac{\delta\mu_{\mathrm{pr}}}{\delta\psi(r)} = \begin{cases} -8\pi r^2\left[\frac{q_{\mathrm{c_1u}}}{I^{\psi}_{\mathrm{c_1u}}}e^{-\beta q_{\mathrm{c_1u}}\psi(r)} + \frac{q_{\mathrm{c_2u}}}{I^{\psi}_{\mathrm{c_2u}}}e^{-\beta q_{\mathrm{c_2u}}\psi(r)} - \frac{q_{\mathrm{c_1b}}}{I^{\psi}_{\mathrm{c_1b}}}e^{-\beta q_{\mathrm{c_1b}}\psi(r)} - \frac{q_{\mathrm{c_2b}}}{I^{\psi}_{\mathrm{c_2b}}}e^{-\beta q_{\mathrm{c_2b}}\psi(r)}\right], & \text{if } r < R_{\mathrm{nucl}} \\ 0 & , \text{ otherwise}\end{cases} \tag{E11}$$

Here we neglected a narrow boundary layer of a few Debye lengths in width (about several nanometers) right next to the NE, where the potential $\psi$ drops exponentially to the cytoplasmic level. This, however, does not influence any of the results presented in the main text as the boundary layer is much smaller than the size of the cell nucleus or the cell cytoplasm.

In summary, we now have practically all of the necessary information to calculate the functional derivative of the

DNA partition function, $\frac{\delta \ln Z_\psi}{\delta \psi(r)}$, which is defined by Eq. (E5). The only missing piece are the expansion coefficients of $T^{(4)}_{k_j k_{j+1}}$ transfer-functions required for the construction of $\mathbf{L}$ and $\frac{\delta \mathbf{L}}{\delta \psi(r)}$ matrices. In the next two sections, Appendix F and Appendix G, we will show how to derive formulas for their computation.

#### Appendix F: Orthogonal D-functions

To find out the elements of the transfer-matrix, $\mathbf{L}$, and the boundary condition vector, $\mathbf{Y}$, we will use several famous results from the group theory, which have been described in our previous work [1] and which we are going to repeat in this Appendix section as they will be exploited extensively in our derivations.

First of all, we would like to recall that from the group theory it is known that any square-integrable function defined on SO(3) group of rotation matrices can be expanded into a series of orthogonal functions, $D^s_{p,q}$, which have the following canonical form, see p. 101 in [36]:

$$D^s_{p,q}(\alpha,\beta,\gamma) = e^{-ip\alpha} P^s_{p,q}(\cos\beta)\, e^{-iq\gamma} \tag{F1}$$

Here $(\alpha,\beta,\gamma)$ are Euler rotation angles, which usually used to parametrize SO(3) group ($\alpha,\gamma \in [0,2\pi]$ and $\beta \in [0,\pi]$); $s$, $p$, $q$ are integers such that $s \geq 0$ and $-s \leq p,q \leq s$; finally, $P^s_{p,q}$ are polynomials, which relate to the elements of so-called small Wigner d-matrix, $d^s_{p,q}$, as: $P^s_{p,q}(\cos\beta) = i^{p-q} d^s_{p,q}(\beta)$.

Functions $D^s_{p,q}$ and polynomials $P^s_{p,q}$ possess a number of important properties, which will come in handy later.

First, by substituting $(\alpha,\beta,\gamma) = (0,0,0)$ into Eq. (F1) and taking into account that $d^s_{p,q}(0) = \delta_{pq}$, we get:

$$D^s_{p,q}(0,0,0) = P^s_{p,q}(1) = i^{p-q} d^s_{p,q}(0) = \delta_{pq} \tag{F2}$$

Here, as before, $\delta_{pq}$ is the Kronecker delta ($\delta_{pq} = 1$ if $p = q$ and $\delta_{pq} = 0$, otherwise).

Furthermore, since for any indexes $s \geq 0$ and $-s \leq p,q \leq s$: $d^s_{p,q}(\beta)$ are real functions obeying the following symmetric relations $d^s_{p,q}(\beta) = (-1)^{q-p} d^s_{q,p}(\beta) = d^s_{-q,-p}(\beta)$, it is not very hard to see that:

$$(-1)^{q-p}\, \overline{P}^s_{p,q}(x) = P^s_{p,q}(x) = P^s_{q,p}(x) = P^s_{-p,-q}(x) \tag{F3}$$

Where the bar denotes complex conjugation.

Combining together Eq. (F1) and Eq. (F3), we obtain:

$$\overline{D}^s_{p,q}(\mathbf{R}) = (-1)^{p-q}\, D^s_{-p,-q}(\mathbf{R}) \tag{F4}$$

Here and below for the sake of formulas simplicity we use $D^s_{p,q}(\mathbf{R})$ notation to address $D^s_{p,q}(\alpha,\beta,\gamma)$ functions, where $\mathbf{R}$ is the Euler rotation matrix corresponding to $(\alpha,\beta,\gamma)$ angles.

Using Eq. (F1) and Eq. (F3), it is also straightforward to show that:

$$D^s_{p,q}\left(\mathbf{R}^{-1}\right) = \overline{D}^s_{q,p}(\mathbf{R}) \tag{F5}$$

Here, as before, $s$, $p$ and $q$ are integer indexes. $\mathbf{R}^{-1}$ is the inverse of matrix $\mathbf{R}$ (i.e., $\mathbf{R}^{-1}\mathbf{R} = \mathbf{R}\mathbf{R}^{-1} = \mathbf{I}$, where $\mathbf{I}$ is the identity matrix), which corresponds to Euler angles $(\pi-\gamma,\beta,\pi-\alpha)$: $\mathbf{R}^{-1} = \mathbf{R}^{-1}(\pi-\gamma,\beta,\pi-\alpha)$.

Next, functions $D^s_{p,q}$ obey the following two multiplication rules [36]:

$$D^s_{p,q}(\mathbf{R}_1\mathbf{R}_2) = \sum_{t=-s}^{s} D^s_{p,t}(\mathbf{R}_1)\, D^s_{t,q}(\mathbf{R}_2) \tag{F6}$$

and

$$D^{s_1}_{p_1,q_1}(\mathbf{R})\, D^{s_2}_{p_2,q_2}(\mathbf{R}) = \sum_s \langle s_1 s_2 p_1 p_2 | s(p_1{+}p_2)\rangle \, \langle s_1 s_2 q_1 q_2 | s(q_1{+}q_2)\rangle \, D^{s}_{p_1+p_2,q_1+q_2}(\mathbf{R}) \tag{F7}$$

Where $\langle s_1 s_2 p_1 p_2 | s_3 p_3\rangle$ are Clebsch-Gordan coefficients. For the sake of formula simplicity, we will use below Wigner 3-j symbols instead of Clebsch-Gordan coefficients, which relate to each other as:

$$\begin{pmatrix} s_1 & s_2 & s_3 \\ p_1 & p_2 & p_3 \end{pmatrix} = \frac{(-1)^{s_1-s_2-p_3}}{\sqrt{2s_3{+}1}} \, \langle s_1 s_2 p_1 p_2 | s_3(-p_3)\rangle \tag{F8}$$

The final important property of $D^s_{p,q}$ functions required for our derivations is their orthogonality, which was mentioned in the beginning of this Appendix section. Specifically, it can be shown that [36]:

$$\int \mathrm{d}\mathbf{R}\, \overline{D}^{s_1}_{p_1,q_1}(\mathbf{R})\, D^{s_2}_{p_2,q_2}(\mathbf{R}) = \frac{8\pi^2}{2s_1{+}1}\, \delta_{s_1 s_2}\delta_{p_1 p_2}\delta_{q_1 q_2} \tag{F9}$$

Where the integration in the above formula is carried out over all possible values of the Euler angles, $(\alpha, \beta, \gamma)$:

$$\int \mathrm{d}\mathbf{R} = \int_0^{2\pi} \mathrm{d}\alpha \int_0^{2\pi} \mathrm{d}\gamma \int_0^{\pi} \sin\beta \, \mathrm{d}\beta \tag{F10}$$

Orthogonality and completeness of $D^s_{p,q}$ functions make it possible to use them as a Hilbert basis in the space of square-integrable functions, $F(\alpha,\beta,\gamma) = F(\mathbf{R})$, defined on SO(3) group [36]. Therefore, any such function can be expanded into the following series:

$$F(\mathbf{R}) = \sum_{s=0}^{\infty} \sum_{p,q=-s}^{s} F_{p,q,s} D^s_{p,q}(\mathbf{R}) \tag{F11}$$

Where $F_{p,q,s}$ expansion coefficients are:

$$F_{p,q,s} = \frac{2s{+}1}{8\pi^2} \int \mathrm{d}\mathbf{R}\, \overline{D}^s_{p,q}(\mathbf{R})\, F(\mathbf{R}) \tag{F12}$$

Analogously, for any square-integrable function $F(\mathbf{R},\mathbf{R}')$, where $\mathbf{R}$ and $\mathbf{R}'$ are two rotation matrices, we have:

$$F\big(\mathbf{R},\mathbf{R}'\big) = \sum_{s,s'=0}^{\infty} \sum_{p,q=-s}^{s} \sum_{p',q'=-s'}^{s'} F^{p',\,q',\,s'}_{p,\,q,\,s}\, D^s_{p,q}(\mathbf{R})\, \overline{D}^{s'}_{p',q'}\big(\mathbf{R}'\big) \tag{F13}$$

Where $F^{p',\,q',\,s'}_{p,\,q,\,s}$ expansion coefficients are:

$$F^{p',\,q',\,s'}_{p,\,q,\,s} = \frac{(2s{+}1)(2s'{+}1)}{\big(8\pi^2\big)^2} \int \mathrm{d}\mathbf{R}\mathrm{d}\mathbf{R}'\, \overline{D}^s_{p,q}(\mathbf{R})\, F\big(\mathbf{R},\mathbf{R}'\big)\, D^{s'}_{p',q'}\big(\mathbf{R}'\big) \tag{F14}$$

In fact, in order to simplify construction of the DNA transfer-matrix, $\mathbf{L}$, it is more convenient to use in Eq. (F13) and Eq. (F14) normalized $D^s_{p,q}$ functions instead of the original ones. Indeed, from Eq. (F9) it can be seen that the $L_2$-norm of $D^s_{p,q}$ functions, $\|D^s_{p,q}\|$, equals to:

$$\|D^s_{p,q}\|^2 = \int \mathrm{d}\mathbf{R}\, \overline{D}^s_{p,q}(\mathbf{R})\, D^s_{p,q}(\mathbf{R}) = \frac{8\pi^2}{2s{+}1} \tag{F15}$$

Hence, while being orthogonal, the basis formed by $D^s_{p,q}$ functions is not orthonormal. Using Eq. (F15), we can easily normalize it by switching from $D^s_{p,q}$ to $\sqrt{\frac{2s+1}{8\pi^2}}D^s_{p,q}$ functions. By doing so, Eq. (F13) turns into:

$$F\big(\mathbf{R},\mathbf{R}'\big) = \frac{1}{8\pi^2}\sum_{s,s'=0}^{\infty}\sum_{p,q=-s}^{s}\sum_{p',q'=-s'}^{s'}\sqrt{(2s+1)(2s'+1)}\,F^{p',q',s'}_{p,\,q,\,s}\,D^s_{p,q}(\mathbf{R})\,\overline{D}^{s'}_{p',q'}\big(\mathbf{R}'\big) \tag{F16}$$

Where $F^{p',q',s'}_{p,\,q,\,s}$ expansion coefficients are:

$$F^{p',q',s'}_{p,\,q,\,s} = \frac{\sqrt{(2s+1)(2s'+1)}}{8\pi^2}\int \mathrm{d}\mathbf{R}\mathrm{d}\mathbf{R}'\,\overline{D}^s_{p,q}(\mathbf{R})\,F\big(\mathbf{R},\mathbf{R}'\big)\,D^{s'}_{p',q'}\big(\mathbf{R}'\big) \tag{F17}$$

With all of the above formulas at hand, we can finally begin to deduce mathematical expressions for the elements of the DNA transfer-matrix, $\mathbf{L}$, and the boundary condition vector, $\mathbf{Y}$, discussed in Appendices C-D.

#### Appendix G: Derivation of the transfer-matrix and the boundary condition vector elements

##### 1. General strategy

To find the elements of the transfer-matrix, $\mathbf{L}$, and the boundary condition vector, $\mathbf{Y}$, we are going to adapt a multi-step approach by obtaining first expansion coefficients of low-level functions, $T^{(2)}_{k_jk_{j+1}}$ and $\xi^{(2)}_0$, and then using them to derive mathematical expressions for the expansion coefficients of higher-level functions, $T^{(4)}_{k_jk_{j+1}}$ and $\xi^{(3)}_0$. Eventually, this will lead us to formulas for the entries of $\mathbf{T}^{(5)}_{k_jk_{j+1}}$ and $\boldsymbol{\xi}^{(4)}_0$ matrix-blocks, which play the central role in the construction of the transfer-matrix, $\mathbf{L}$, and the boundary condition vector, $\mathbf{Y}$, see Eq. (D11)-Eq. (D13).

To acquire expansion coefficients of $T^{(2)}_{k_jk_{j+1}}$ and $\xi^{(2)}_0$, it is convenient first to write down explicit formulas of these two functions, which can be done by substituting Eq. (C4) and Eq. (C6) into Eq. (C15):

$$T^{(2)}_{k_jk_{j+1}}(\mathbf{R}_j,\mathbf{R}_{j+1},\boldsymbol{\nu}_j) = e^{ib_{k_j}[\mathbf{R}_j\mathbf{z}_0\cdot\boldsymbol{\nu}_j]}\times\begin{cases}\displaystyle\int_0^{2\pi}\mathrm{d}\eta_{\mathrm{in},j}\,e^{-\beta E_{k_jk_{j+1}}(\mathbf{R}_j,\mathbf{R}_{j+1})}, & \text{if } (k_j,k_{j+1})=(0,1)\\ e^{-\beta E_{k_jk_{j+1}}(\mathbf{R}_j,\mathbf{R}_{j+1})}, & \text{otherwise}\end{cases}$$

$$\xi^{(2)}_{k_N}(\mathbf{R}_N,\mathbf{R}_1,\boldsymbol{\nu}_N) = \delta_{k_N0}\times\xi(\mathbf{R}_N,\mathbf{R}_1)\,e^{\beta bf[\mathbf{z}_0\cdot\mathbf{R}_N\mathbf{z}_0]}e^{ib[\mathbf{R}_N\mathbf{z}_0\cdot\boldsymbol{\nu}_N]} \tag{G1}$$

It should be noted that all $E_{k_jk_{j+1}}(\mathbf{R}_j,\mathbf{R}_{j+1})$ energy terms, which describe local contributions of neighbouring DNA segments into the total energy of DNA, in the general case can be divided into two groups: 1) $E_{00}$, $E_{01}$ and $E_{K0}$ terms that look like a function $\frac{a}{2\beta}(\mathbf{R}_j\mathbf{A}\mathbf{z}_0-\mathbf{R}_{j+1}\mathbf{z}_0)^2+\frac{c}{2\beta}[\Delta\varphi_j(\mathbf{R}_j\mathbf{A},\mathbf{R}_{j+1})]^2-b_{k_j}f\big[\mathbf{z}_0\cdot\mathbf{R}_j\mathbf{z}_0\big]+\mathrm{const}$, see, for example, $E_{00}$, $E_{01}$ and $E_{30}$ in Eq. (B17); and 2) $E_{12}$, $E_{23}$, ..., $E_{K-1\,K}$ terms, which have the following mathematical form: $-\frac{1}{\beta}\ln\delta(\mathbf{R}_j-\mathbf{R}_{j+1})-b_{k_j}f\big[\mathbf{z}_0\cdot\mathbf{R}_j\mathbf{z}_0\big]+\mathrm{const}$, see, for example, $E_{12}$ and $E_{23}$ in Eq. (B17). As a result, it can be seen from Eq. (G1) that expansion coefficients of $T^{(2)}_{k_jk_{j+1}}$ transfer-functions can be easily obtained from the expansion series of the following two functions:

$$F_1\big(\mathbf{R},\mathbf{R}',\boldsymbol{\nu}\big) = e^{-\frac{a}{2}(\mathbf{R}\mathbf{A}\mathbf{z}_0-\mathbf{R}'\mathbf{z}_0)^2-\frac{c}{2}[\Delta\varphi(\mathbf{R}\mathbf{A},\mathbf{R}')]^2+\beta bf[\mathbf{z}_0\cdot\mathbf{R}\mathbf{z}_0]+ib[\boldsymbol{\nu}\cdot\mathbf{R}\mathbf{z}_0]}$$

$$F_2\big(\mathbf{R},\mathbf{R}',\boldsymbol{\nu}\big) = \delta\big(\mathbf{R}-\mathbf{R}'\big)\;e^{\beta bf[\mathbf{z}_0\cdot\mathbf{R}\mathbf{z}_0]+ib[\boldsymbol{\nu}\cdot\mathbf{R}\mathbf{z}_0]} \tag{G2}$$

Here $\mathbf{R}$ and $\mathbf{R}'$ are Euler rotation matrices, which describe orientations of neighbouring DNA segments in the polygonal chain representing the DNA molecule with respect to the global coordinate system; whereas, matrix $\mathbf{A}$ denotes an equilibrium orientation of the DNA segments with respect to each other in the absence of $\psi$ and $U_{R_{\mathrm{nucl}}}$ potentials, and zero stretching force ($f=0$).

In the process of deriving expansion series of $F_1$ and $F_2$ functions, we will obtain several useful formulas, which will also allow us to get a mathematical expression for the expansion coefficients of $\xi^{(2)}_0$ function, thus providing a full set of information needed for the second step – finding expansion coefficients of $T^{(4)}_{k_jk_{j+1}}$ and $\xi^{(3)}_0$ functions based on Eq. (C14) and Eq. (C43). The third and the final step – acquiring formulas for the entries of $\mathbf{T}^{(5)}_{k_j k_{j+1}}$ and $\boldsymbol{\xi}^{(4)}_0$ matrix-blocks, will be done by using Eq. (C34) and Eq. (C35), concluding this Appendix section.

##### 2. Expansion series of $F_1$ function

Let's begin with $F_1$ function. To obtain its expansion series, we will follow the same procedure as the one described in our previous theoretical study, see Appendix E of ref. [3]. Namely, we will first derive separate expansion series of the exponential functions comprising the global and local energy terms, $e^{\beta bf[\mathbf{z}_0\cdot\mathbf{R}\mathbf{z}_0]+ib[\boldsymbol{\nu}\cdot\mathbf{R}\mathbf{z}_0]}$ and $e^{-\frac{a}{2}(\mathbf{R}\mathbf{A}\mathbf{z}_0-\mathbf{R}'\mathbf{z}_0)^2-\frac{c}{2}[\Delta\varphi(\mathbf{R}\mathbf{A},\mathbf{R}')]^2}$, respectively, and then combine them together by using the second multiplication rule for the Wigner D-functions [Eq. (F7)] to get the final result.

To acquire expansion series of $e^{\beta bf[\mathbf{z}_0\cdot\mathbf{R}\mathbf{z}_0]+ib[\boldsymbol{\nu}\cdot\mathbf{R}\mathbf{z}_0]}$ function, we will start with a somewhat simpler case of $f=0$, considering only $e^{ib[\boldsymbol{\nu}\cdot\mathbf{R}\mathbf{z}_0]}$ term at the moment. Let $\mathbf{A}_\nu$ be a Euler rotation matrix such that $\boldsymbol{\nu}=\nu\times\mathbf{A}_\nu\mathbf{z}_0$, where $\nu=\|\boldsymbol{\nu}\|$. I.e., matrix $\mathbf{A}_\nu$ rotates the unit basis vector $\mathbf{z}_0$ of the global coordinate system into a unit vector collinear to $\boldsymbol{\nu}$. Then, by utilizing Eq. (F12), it is straightforward to obtain the following formula for the expansion coefficients of $e^{ib[\boldsymbol{\nu}\cdot\mathbf{R}\mathbf{z}_0]}$:

$$\begin{aligned}F_{p,\,q,\,s} &= \frac{2s{+}1}{8\pi^2}\int \mathrm{d}\mathbf{R}\,\overline{D}^s_{p,q}(\mathbf{R})\,e^{ib[\boldsymbol{\nu}\cdot\mathbf{R}\mathbf{z}_0]} = \frac{2s{+}1}{8\pi^2}\int \mathrm{d}\mathbf{R}\,\overline{D}^s_{p,q}(\mathbf{R})\,e^{ib\nu[\mathbf{A}_\nu\mathbf{z}_0\cdot\mathbf{R}\mathbf{z}_0]} = \Big[\mathbf{R}\to\mathbf{A}_\nu\mathbf{R}\Big] = \\ &= \frac{2s{+}1}{8\pi^2}\int \mathrm{d}\mathbf{R}\,\overline{D}^s_{p,q}(\mathbf{A}_\nu\mathbf{R})\,e^{ib\nu[\mathbf{z}_0\cdot\mathbf{R}\mathbf{z}_0]} = \frac{2s{+}1}{8\pi^2}\sum_m \overline{D}^s_{p,m}(\mathbf{A}_\nu)\int \mathrm{d}\mathbf{R}\,\overline{D}^s_{m,q}(\mathbf{R})\,e^{ib\nu[\mathbf{z}_0\cdot\mathbf{R}\mathbf{z}_0]} = \\ &= \frac{2s{+}1}{8\pi^2}\sum_m \overline{D}^s_{p,m}(\mathbf{A}_\nu)\int_0^{2\pi}\mathrm{d}\alpha\int_0^{\pi}\sin\beta\,\mathrm{d}\beta\int_0^{2\pi}\mathrm{d}\gamma\,e^{im\alpha+iq\gamma}\overline{P}^s_{m,q}(\cos\beta)\,e^{ib\nu\cos\beta} = \\ &= \frac{2s{+}1}{8\pi^2}\sum_m \overline{D}^s_{p,m}(\mathbf{A}_\nu)\times 4\pi^2\,\delta_{m0}\,\delta_{q0}\int_{-1}^{1}P_s(u)\,e^{ib\nu u}\mathrm{d}u = \delta_{q0}\times(2s{+}1)\,i^s j_s(b\nu)\,\overline{D}^s_{p,0}(\mathbf{A}_\nu)\end{aligned} \qquad \text{(G3)}$$

Here we used left-invariance of the Haar measure of SO(3) group (i.e., that $\int \mathrm{d}(\mathbf{A}_\nu\mathbf{R})=\int\mathrm{d}\mathbf{R}$, see ref. [36]), and the fact that $[\mathbf{A}_\nu\mathbf{z}_0\cdot\mathbf{A}_\nu\mathbf{R}\mathbf{z}_0]=[\mathbf{z}_0\cdot\mathbf{R}\mathbf{z}_0]=\cos\beta$, where $(\alpha,\beta,\gamma)$ are Euler angles of the rotation matrix $\mathbf{R}$. Furthermore, in the above equation, we applied the first multiplication rule for the Wigner D-functions [Eq. (F6)] and have taken into account that $\overline{P}^s_{0,0}(\cos\beta)=P^s_{0,0}(\cos\beta)=P_s(\cos\beta)$, where $P_s(x)$ are Legendre polynomials whose inverse Fourier-transform is:

$$\int_{-1}^{1}P_s(x)\,e^{i\nu x}\mathrm{d}x = 2\,i^s j_s(\nu) \qquad \text{(G4)}$$

Where $j_s(\nu)$ is the spherical Bessel function of the first kind.

Thus, from Eq. (F11) and Eq. (G3) it follows that:

$$e^{ib[\boldsymbol{\nu}\cdot\mathbf{R}\mathbf{z}_0]} = \sum_{s,p}(2s{+}1)\,i^s j_s(b\nu)\,\overline{D}^s_{p,0}(\mathbf{A}_\nu)\,D^s_{p,0}(\mathbf{R}) \qquad \text{(G5)}$$

Substituting $-i\beta\mathbf{f}=-i\beta f\mathbf{z}_0$ for $\boldsymbol{\nu}$ in Eq. (G5), it is not hard to find an expansion formula for the second part, $e^{bf[\mathbf{z}_0\cdot\mathbf{R}\mathbf{z}_0]}$, of the exponential function containing the global energy terms:

$$e^{\beta bf[\mathbf{z}_0\cdot\mathbf{R}\mathbf{z}_0]} = \sum_{s',p'}(2s'{+}1)\,i_{s'}(\beta bf)\,\overline{D}^{s'}_{p',0}(\mathbf{A}_f)\,D^{s'}_{p',0}(\mathbf{R}) \qquad \text{(G6)}$$

Where $i_s(x)$ is the modified spherical Bessel function of the first kind, which relates to $j_s(x)$ function as: $i_s(x)=i^s j_s(-ix)$. In the above formula, matrix $\mathbf{A}_f$ is defined in a very similar way as matrix $\mathbf{A}_\nu$: $\mathbf{f}=f\times\mathbf{A}_f\mathbf{z}_0$. Since in all our calculations the stretching force, $\mathbf{f}$, applied to DNA is collinear to $\mathbf{z}_0$-axis of the global coordinate system, we can simply put $\mathbf{A}_f = \mathbf{I}$, where $\mathbf{I}$ is the identity $3\times 3$ matrix. As a result, with the help of Eq. (F2), Eq. (G6) simplifies to:

$$e^{\beta b f[\mathbf{z}_0\cdot\mathbf{R}\mathbf{z}_0]} = \sum_{s'} (2s'{+}1)\, i_{s'}(\beta b f)\, D^{s'}_{0,0}(\mathbf{R}) \tag{G7}$$

In order to obtain an expansion formula for the combined $e^{\beta b f[\mathbf{z}_0\cdot\mathbf{R}\mathbf{z}_0]+ib[\boldsymbol{\nu}\cdot\mathbf{R}\mathbf{z}_0]}$ function, all we need to do is to multiply Eq. (G5) by Eq. (G7) and then use the second multiplication rule for the Wigner D-functions [Eq. (F7)]:

$$\begin{aligned} e^{\beta b f[\mathbf{z}_0\cdot\mathbf{R}\mathbf{z}_0]+ib[\boldsymbol{\nu}\cdot\mathbf{R}\mathbf{z}_0]} &= \sum_{s,s',p} (2s{+}1)\,(2s'{+}1)\, i^s j_s(b\nu)\, i_{s'}(\beta b f)\, \overline{D}^s_{p,0}(\mathbf{A}_\nu)\, D^s_{p,0}(\mathbf{R})\, D^{s'}_{0,0}(\mathbf{R}) = \\ &= \sum_{s,s',u,p} (2s{+}1)\,(2s'{+}1)\,(2u{+}1)\,(-1)^p\, i^s j_s(b\nu)\, i_{s'}(\beta b f)\, \overline{D}^s_{p,0}(\mathbf{A}_\nu)\times \\ &\quad\times \begin{pmatrix} s & s' & u \\ p & 0 & -p \end{pmatrix} \begin{pmatrix} s & s' & u \\ 0 & 0 & 0 \end{pmatrix} \times D^u_{p,0}(\mathbf{R}) \end{aligned} \tag{G8}$$

Next, to derive expansion series of $e^{-\frac{a}{2}(\mathbf{R}\mathbf{A}\mathbf{z}_0-\mathbf{R}'\mathbf{z}_0)^2-\frac{c}{2}[\Delta\varphi(\mathbf{R}\mathbf{A},\mathbf{R}')]^2}$ function, we will again follow the logic from ref. [1]. Specifically, by taking a look at this function, it is clear that it actually depends only on $(\mathbf{R}\mathbf{A})^{-1}\mathbf{R}'$ product of Euler rotation matrices. Indeed, it is not hard to see that $(\mathbf{R}\mathbf{A}\mathbf{z}_0-\mathbf{R}'\mathbf{z}_0)^2 = 2-2\,[\mathbf{R}\mathbf{A}\mathbf{z}_0\cdot\mathbf{R}'\mathbf{z}_0] = 2-2\,[\mathbf{z}_0\cdot(\mathbf{R}\mathbf{A})^{-1}\mathbf{R}'\mathbf{z}_0]$ and $\Delta\varphi(\mathbf{R}\mathbf{A},\mathbf{R}') = \Delta\varphi(\mathbf{I},(\mathbf{R}\mathbf{A})^{-1}\mathbf{R}')$. In other words, the twist angle between the coordinate systems corresponding to $\mathbf{R}\mathbf{A}$ and $\mathbf{R}'$ Euler rotation matrices as well as the bending angle between their $\mathbf{z}$-axes depend only on the relative orientation of the two coordinate systems with respect to each other and is independent from their exact alignments with respect to the global coordinate frame, $(\mathbf{x}_0,\mathbf{y}_0,\mathbf{z}_0)$.

As a result, the expansion series of $e^{-\frac{a}{2}(\mathbf{R}\mathbf{A}\mathbf{z}_0-\mathbf{R}'\mathbf{z}_0)^2-\frac{c}{2}[\Delta\varphi(\mathbf{R}\mathbf{A},\mathbf{R}')]^2}$ function can be found in two steps. First, we will consider a special case in which the coordinate system corresponding to $\mathbf{R}\mathbf{A}$ matrix product is identical to the global coordinate frame (i.e., $\mathbf{R}\mathbf{A}=\mathbf{I}$), and the coordinate system corresponding to the matrix $\mathbf{R}'$ is only slightly rotated relative to it (due to $a \gg 1$ and $c \gg 1$, see Table I). Second, by substituting $\mathbf{R}' \to (\mathbf{R}\mathbf{A})^{-1}\mathbf{R}'$ into the formula obtained in the special case and by using Eq. (F6), we will get the desired expansion series for the above exponential function in the general case.

Let $(\alpha',\beta',\gamma')$ be the Euler angles corresponding to matrix $\mathbf{R}'$. Then, by taking into account the above notes, for the special case of $\mathbf{R}\mathbf{A}=\mathbf{I}$ we have: $(\mathbf{R}\mathbf{A}\mathbf{z}_0-\mathbf{R}'\mathbf{z}_0)^2 = 2-2\,[\mathbf{z}_0\cdot\mathbf{R}'\mathbf{z}_0] = 2-2\cos\beta'$ and $\Delta\varphi(\mathbf{R}\mathbf{A},\mathbf{R}') = \Delta\varphi(\mathbf{I},\mathbf{R}')$. Furthermore, since the coordinate frame corresponding to $\mathbf{R}'$ matrix is only slightly rotated relative to the global coordinate system, it is clear that:

$$\Delta\varphi(\mathbf{I},\mathbf{R}') \approx \alpha'+\gamma' \approx \sin(\alpha'+\gamma') \quad \text{and} \quad \big[\Delta\varphi(\mathbf{I},\mathbf{R}')\big]^2 \approx 2-2\cos(\alpha'+\gamma') \tag{G9}$$

Thus, in the special case of $\mathbf{R}\mathbf{A}=\mathbf{I}$:

$$e^{-\frac{a}{2}\left(\mathbf{R}\mathbf{A}\mathbf{z}_0-\mathbf{R}'\mathbf{z}_0\right)^2-\frac{c}{2}\left[\Delta\varphi\left(\mathbf{R}\mathbf{A},\mathbf{R}'\right)\right]^2} = e^{-\frac{a}{2}\left(\mathbf{z}_0-\mathbf{R}'\mathbf{z}_0\right)^2-\frac{c}{2}\left[\Delta\varphi\left(\mathbf{I},\mathbf{R}'\right)\right]^2} = e^{-a-c}\, e^{a\cos\beta'+c\cos\left(\alpha'+\gamma'\right)} \tag{G10}$$

To proceed further, we will need the following form of the Jacobi-Anger expansion formula (see Eq. (E7) in ref. [3]):

$$e^{\rho\cos\varphi} = \sum_{m=-\infty}^{+\infty} I_m(\rho)\, e^{im\varphi} \tag{G11}$$

Where $i$ is imaginary unit and $\rho$ is an arbitrary constant; $I_m(x)$ is the modified Bessel function of the first kind, which have the following property: $I_{-m}(x) = I_m(x)$, see p. 714 in ref. [40].

By applying Eq. (F12) and Eq. (G11) to Eq. (G10), it is not hard to obtain the following expansion coefficients for

$e^{-\frac{a}{2}(\mathbf{z}_0-\mathbf{R}'\mathbf{z}_0)^2-\frac{c}{2}[\Delta\varphi(\mathbf{I},\mathbf{R}')]^2}$ function:

$$\begin{aligned} F_{p'',q'',s''} &= \frac{2s''+1}{8\pi^2}\int \mathrm{d}\mathbf{R}'\,\overline{D}^{s''}_{p'',q''}\big(\mathbf{R}'\big)\,e^{-\frac{a}{2}\left(\mathbf{z}_0-\mathbf{R}'\mathbf{z}_0\right)^2-\frac{c}{2}\left[\Delta\varphi\left(\mathbf{I},\mathbf{R}'\right)\right]^2} = \\ &= \frac{2s''+1}{8\pi^2}\,e^{-a-c}\int \mathrm{d}\mathbf{R}'\,\overline{P}^{s''}_{p'',q''}(\cos\beta')\,e^{a\cos\beta'}\times\sum_{m=-\infty}^{+\infty} I_m(c)\,e^{i\left(p''+m\right)\alpha'+i\left(q''+m\right)\gamma'} = \\ &= \delta_{p''q''}\times\frac{1}{2}\,(2s''+1)\,e^{-a-c}\,I_{p''}(c)\,\mathscr{L}^{s''}_{p''}(-a) \end{aligned} \tag{G12}$$

Where $\mathscr{L}^{s}_{p}(x)$ designates bilateral Laplace transform of $P^{s}_{p,p}$ polynomial (or, which is the same thing, a diagonal element, $d^{s}_{p,p}$, of the Wigner small d-matrix):

$$\mathscr{L}^{s}_{p}(x) = \int_{-1}^{1} P^{s}_{p,p}(y)\,e^{-xy}\mathrm{d}y = \int_{-1}^{1} d^{s}_{p,p}\big(\cos^{-1}y\big)\,e^{-xy}\mathrm{d}y \tag{G13}$$

By substituting Eq. (G12) into Eq. (F11), we finally get the desired expansion series for the special case of $\mathbf{RA}=\mathbf{I}$:

$$e^{-\frac{a}{2}\left(\mathbf{z}_0-\mathbf{R}'\mathbf{z}_0\right)^2-\frac{c}{2}\left[\Delta\varphi\left(\mathbf{I},\mathbf{R}'\right)\right]^2} = \frac{1}{2}\,e^{-a-c}\sum_{s'',p''}(2s''+1)\,I_{p''}(c)\,\mathscr{L}^{s''}_{p''}(-a)\,D^{s''}_{p'',p''}\big(\mathbf{R}'\big) \tag{G14}$$

To extend the above formula to the general case, we simply need to put $\mathbf{R}'\to(\mathbf{RA})^{-1}\mathbf{R}'$ and use the first multiplication rule for $D^{s}_{p,q}$ functions [Eq. (F6)]. As a result, we have:

$$e^{-\frac{a}{2}\left(\mathbf{RAz}_0-\mathbf{R}'\mathbf{z}_0\right)^2-\frac{c}{2}\left[\Delta\varphi\left(\mathbf{RA},\mathbf{R}'\right)\right]^2} = \frac{1}{2}\,e^{-a-c}\sum_{s'',p'',q,v}(2s''+1)\,I_{p''}(c)\,\mathscr{L}^{s''}_{p''}(-a)\,D^{s''}_{v,p''}(\mathbf{A})\,D^{s''}_{q,v}(\mathbf{R})\,\overline{D}^{s''}_{q,p''}\big(\mathbf{R}'\big) \tag{G15}$$

Here, in addition to the index change $p''\to-p''$, we also used Eq. (F4)-Eq. (F5) and Eq. (G13) in order to write down the expansion series in the form of Eq. (F13).

Finally, by multiplying Eq. (G8) and Eq. (G15), and applying the second multiplication rule for the Wigner D-functions [Eq. (F7)], it can be shown that:

$$\begin{aligned} F_1\big(\mathbf{R},\mathbf{R}',\boldsymbol{\nu}\big) &= \frac{1}{2}\,e^{-a-c}\sum_{\substack{s,s',s'',q\\ p,p'',u,v}}(2s+1)\,(2s'+1)\,(2s''+1)\,(2u+1)\,(-1)^p\,i^s j_s(b\nu)\,i_{s'}(\beta bf)\,I_{p''}(c)\,\mathscr{L}^{s''}_{p''}(-a)\times \\ &\quad\times\overline{D}^{s}_{p,0}(\mathbf{A}_\nu)\,D^{s''}_{v,p''}(\mathbf{A})\times\begin{pmatrix} s & s' & u\\ p & 0 & -p\end{pmatrix}\begin{pmatrix} s & s' & u\\ 0 & 0 & 0\end{pmatrix}\times D^{u}_{p,0}(\mathbf{R})\,D^{s''}_{q,v}(\mathbf{R})\,\overline{D}^{s''}_{q,p''}\big(\mathbf{R}'\big) = \\ &= \frac{1}{2}\,e^{-a-c}\sum_{\substack{s,s',s'',q,t\\ p,p'',u,v}}(2s+1)\,(2s'+1)\,(2s''+1)\,(2u+1)\,(2t+1)(-1)^{q+v}\,i^s j_s(b\nu)\,i_{s'}(\beta bf)\,I_{p''}(c)\,\mathscr{L}^{s''}_{p''}(-a)\times \\ &\quad\times\overline{D}^{s}_{p,0}(\mathbf{A}_\nu)\,D^{s''}_{v,p''}(\mathbf{A})\times\begin{pmatrix} s & s' & u\\ p & 0 & -p\end{pmatrix}\begin{pmatrix} s & s' & u\\ 0 & 0 & 0\end{pmatrix}\begin{pmatrix} u & s'' & t\\ p & q & -p-q\end{pmatrix}\begin{pmatrix} u & s'' & t\\ 0 & v & -v\end{pmatrix}\times \\ &\quad\times D^{t}_{p+q,v}(\mathbf{R})\,\overline{D}^{s''}_{q,p''}\big(\mathbf{R}'\big) \end{aligned} \tag{G16}$$

By making the following change of indexes: $s''\to s'$, $s'\to t$, $t\to s$, $s\to r$, $v\to q$, $q\to p'$, $p\to p-p'$ and $p''\to q'$, it is straightforward to rewrite the above expansion formula for $F_1$ function in the form of Eq. (F16):

$$F_1\big(\mathbf{R},\mathbf{R}',\boldsymbol{\nu}\big) = \frac{1}{8\pi^2}\sum_{p,p',q,q',s,s'}\sqrt{(2s+1)(2s'+1)}\;\big[F_1(\boldsymbol{\nu})\big]^{p',q',s'}_{p,\,q,\,s}\,D^{s}_{p,q}(\mathbf{R})\,\overline{D}^{s'}_{p',q'}\big(\mathbf{R}'\big) \tag{G17}$$

Where the expansion coefficients $[F_1(\boldsymbol{\nu})]_{p,\,q,\,s}^{p',\,q',\,s'}$ are:

$$[F_1(\boldsymbol{\nu})]_{p,\,q,\,s}^{p',\,q',\,s'} = 4\pi^2\sqrt{(2s{+}1)(2s'{+}1)} \times (-1)^{p'+q}\, e^{-a-c} I_{q'}(c)\, \mathscr{L}_{q'}^{s'}(-a)\, D_{q,q'}^{s'}(\mathbf{A}) \times \sum_{r,t,u} (2r{+}1)\,(2t{+}1)\,(2u{+}1)\, i^r \times$$
$$\times\, j_r(b\nu)\, i_t(\beta b f)\, \overline{D}_{p-p',0}^{r}(\mathbf{A}_\nu) \times \begin{pmatrix} r & t & u \\ p-p' & 0 & p'-p \end{pmatrix} \begin{pmatrix} r & t & u \\ 0 & 0 & 0 \end{pmatrix} \begin{pmatrix} u & s' & s \\ p-p' & p' & -p \end{pmatrix} \begin{pmatrix} u & s' & s \\ 0 & q & -q \end{pmatrix} \tag{G18}$$

##### 3. Expansion series of $F_2$ function

In order to find expansion series of $F_2$ function defined by Eq. (G2), we will use the same strategy as above, deriving first separate expansion formulas for the two parts of $F_2$ function, $e^{\beta b f[\mathbf{z}_0\cdot\mathbf{R}\mathbf{z}_0]+ib[\boldsymbol{\nu}\cdot\mathbf{R}\mathbf{z}_0]}$ and $\delta(\mathbf{R}-\mathbf{R}')$, and then combining them together by using the second multiplication rule for the Wigner D-functions [Eq. (F7)] to obtain the final result.

The best part of such approach is that we already have the expansion series of $e^{\beta b f[\mathbf{z}_0\cdot\mathbf{R}\mathbf{z}_0]+ib[\boldsymbol{\nu}\cdot\mathbf{R}\mathbf{z}_0]}$ exponent, see Eq. (G8). Thus, all that remains to do is to find an expansion formula for the remaining $\delta(\mathbf{R}-\mathbf{R}')$ function. This can be easily done by applying Eq. (F13) and Eq. (F14):

$$F_{p,\,q,\,s}^{p',\,q',\,s'} = \frac{(2s{+}1)(2s'{+}1)}{(8\pi^2)^2} \int \mathrm{d}\mathbf{R}\mathrm{d}\mathbf{R}'\, \overline{D}_{p,q}^{s}(\mathbf{R})\, \delta\big(\mathbf{R}-\mathbf{R}'\big)\, D_{p',q'}^{s'}\big(\mathbf{R}'\big) =$$
$$= \frac{(2s{+}1)(2s'{+}1)}{(8\pi^2)^2} \int \mathrm{d}\mathbf{R}\, \overline{D}_{p,q}^{s}(\mathbf{R})\, D_{p',q'}^{s'}(\mathbf{R}) = \delta_{ss'}\delta_{pp'}\delta_{qq'} \times \frac{2s{+}1}{8\pi^2} \tag{G19}$$

Substituting the above expansion coefficients into Eq. (F13), it immediately follows that:

$$\delta\big(\mathbf{R}-\mathbf{R}'\big) = \frac{1}{8\pi^2}\sum_{s,p,q}(2s{+}1)\, D_{p,q}^{s}(\mathbf{R})\, \overline{D}_{p,q}^{s}\big(\mathbf{R}'\big) = \Big[s\to s''; p\to p'\Big] = \frac{1}{8\pi^2}\sum_{s'',p',q}(2s''{+}1)\, D_{p',q}^{s''}(\mathbf{R})\, \overline{D}_{p',q}^{s''}\big(\mathbf{R}'\big) \tag{G20}$$

Finally, by multiplying Eq. (G8) and Eq. (G20), and using Eq. (F7), we have:

$$F_2\big(\mathbf{R},\mathbf{R}',\boldsymbol{\nu}\big) = \frac{1}{8\pi^2}\sum_{\substack{s,s',s'',u\\ p,p',q}} (2s{+}1)\,(2s'{+}1)\,(2s''{+}1)\,(2u{+}1)\,(-1)^p\, i^s j_s(b\nu)\, i_{s'}(\beta b f)\, \overline{D}_{p,0}^{s}(\mathbf{A}_\nu) \times$$
$$\times \begin{pmatrix} s & s' & u \\ p & 0 & -p \end{pmatrix} \begin{pmatrix} s & s' & u \\ 0 & 0 & 0 \end{pmatrix} \times D_{p,0}^{u}(\mathbf{R})\, D_{p',q}^{s''}(\mathbf{R})\, \overline{D}_{p',q}^{s''}\big(\mathbf{R}'\big) =$$

$$= \frac{1}{8\pi^2}\sum_{\substack{s,s',s'',u\\ v,p,p',q}} (2s{+}1)\,(2s'{+}1)\,(2s''{+}1)\,(2u{+}1)\,(2v{+}1)\,(-1)^{p'+q}\, i^s j_s(b\nu)\, i_{s'}(\beta b f)\, \overline{D}_{p,0}^{s}(\mathbf{A}_\nu) \times$$
$$\times \begin{pmatrix} s & s' & u \\ p & 0 & -p \end{pmatrix} \begin{pmatrix} s & s' & u \\ 0 & 0 & 0 \end{pmatrix} \begin{pmatrix} u & s'' & v \\ p & p' & -p-p' \end{pmatrix} \begin{pmatrix} u & s'' & v \\ 0 & q & -q \end{pmatrix} \times D_{p+p',q}^{v}(\mathbf{R})\, \overline{D}_{p',q}^{s''}\big(\mathbf{R}'\big) \tag{G21}$$

In order to rewrite Eq. (G21) in the form of Eq. (F16), we just need to make a simple change of indexes: $v\to s$, $s\to r$, $s''\to s'$, $s'\to t$, and $p\to p-p'$. As a result, one can obtain the following expansion series for $F_2$ function:

$$F_2\big(\mathbf{R},\mathbf{R}',\boldsymbol{\nu}\big) = \frac{1}{8\pi^2}\sum_{p,p',\,q,q',\,s,s'} \sqrt{(2s{+}1)(2s'{+}1)}\; [F_2(\boldsymbol{\nu})]_{p,\,q,\,s}^{p',\,q',\,s'}\, D_{p,q}^{s}(\mathbf{R})\, \overline{D}_{p',q'}^{s'}\big(\mathbf{R}'\big) \tag{G22}$$

Where the expansion coefficients $[F_2(\boldsymbol{\nu})]^{p',q',s'}_{p,q,s}$ are:

$$[F_2(\boldsymbol{\nu})]^{p',q',s'}_{p,q,s} = \delta_{qq'} \times \sqrt{(2s+1)(2s'+1)} \times (-1)^{p'+q} \sum_{r,t,u} (2r+1)\,(2t+1)\,(2u+1)\, i^r j_r(b\nu)\, i_t(\beta b f) \times \\ \times \overline{D}^r_{p-p',0}(\mathbf{A}_\nu) \times \begin{pmatrix} r & t & u \\ p-p' & 0 & p'-p \end{pmatrix} \begin{pmatrix} r & t & u \\ 0 & 0 & 0 \end{pmatrix} \begin{pmatrix} u & s' & s \\ p-p' & p' & -p \end{pmatrix} \begin{pmatrix} u & s' & s \\ 0 & q & -q \end{pmatrix} \quad \text{(G23)}$$

##### 4. Expansion coefficients of $T^{(2)}_{00}$ function

By using the above expansion series of $F_1$ and $F_2$ functions, it is straightforward to derive formulas for the expansion coefficients of $T^{(2)}_{k_j k_{j+1}}$ transfer-functions. Let's start with $T^{(2)}_{00}$ function first.

Substituting $E_{00}$ energy term from Eq. (B17) into Eq. (G1) and comparing the obtained result to Eq. (G2), it becomes clear that the expansion coefficients of $T^{(2)}_{00}(\mathbf{R}_j, \mathbf{R}_{j+1}, \boldsymbol{\nu}_j)$ transfer-function can be obtained from Eq. (G18) by putting $\mathbf{A} = \mathbf{I}$ and $\boldsymbol{\nu} = \boldsymbol{\nu}_j$. Taking into account Eq. (F2) saying that $D^s_{p,q}(\mathbf{I}) = \delta_{pq}$, it is then easy to obtain the following formula for the expansion coefficients of $T^{(2)}_{00}(\mathbf{R}_j, \mathbf{R}_{j+1}, \boldsymbol{\nu}_j)$ function:

$$[T^{(2)}_{00}(\boldsymbol{\nu}_j)]^{p',q',s'}_{p,q,s} = \delta_{qq'} \times 4\pi^2 \sqrt{(2s+1)(2s'+1)}\;(-1)^{p'+q}\, e^{-a-c} I_{q'}(c)\, \mathscr{L}^{s'}_{q'}(-a) \times \\ \times \sum_{r,t,u} (2r+1)\,(2t+1)\,(2u+1)\, i^r j_r(b\nu_j)\, i_t(\beta b f)\, \overline{D}^r_{p-p',0}(\mathbf{A}_{\nu_j}) \times \\ \times \begin{pmatrix} r & t & u \\ p-p' & 0 & p'-p \end{pmatrix} \begin{pmatrix} r & t & u \\ 0 & 0 & 0 \end{pmatrix} \begin{pmatrix} u & s' & s \\ p-p' & p' & -p \end{pmatrix} \begin{pmatrix} u & s' & s \\ 0 & q & -q \end{pmatrix} \quad \text{(G24)}$$

Where for the sake of simplicity we used the following notations: $s = s_j$, $p = p_j$, $q = q_j$, $s' = s_{j+1}$, $p' = p_{j+1}$ and $q' = q_{j+1}$.

Later, in Appendix H, it will be shown that calculations of the DNA partition function can be considerably simplified by introduction of modified expansion coefficients defined by Eq. (H2). By combining Eq. (H2) with Eq. (H1) and Eq. (G24), it is then rather straightforward to get the following formula for the modified expansion coefficients:

$$[\widetilde{T}^{(2)}_{00}(\boldsymbol{\nu}_j)]^{p',q',s'}_{p,q,s} = \delta_{qq'} \times 4\pi^2 \sqrt{(2s+1)(2s'+1)}\;(-1)^{p'+q}\; i^{p-p'} e^{-a-c} I_{q'}(c)\, \mathscr{L}^{s'}_{q'}(-a) \times \\ \times \sum_{r,t,u} (2r+1)\,(2t+1)\,(2u+1)\, i^r j_r(b\nu_j)\, i_t(\beta b f)\, \overline{P}^r_{p-p',0}(\cos\beta_{\nu_j}) \times \\ \times \begin{pmatrix} r & t & u \\ p-p' & 0 & p'-p \end{pmatrix} \begin{pmatrix} r & t & u \\ 0 & 0 & 0 \end{pmatrix} \begin{pmatrix} u & s' & s \\ p-p' & p' & -p \end{pmatrix} \begin{pmatrix} u & s' & s \\ 0 & q & -q \end{pmatrix} \quad \text{(G25)}$$

Where $\beta_{\nu_j}$ is the polar Euler angle of the wave-vector $\boldsymbol{\nu}_j$.

In the case of zero stretching force ($f = 0$), Eq. (G25) turns into:

$$[\widetilde{T}^{(2)}_{00}(\boldsymbol{\nu}_j)]^{p',q',s'}_{p,q,s} = \delta_{qq'} \times 4\pi^2 \sqrt{(2s+1)(2s'+1)}\;(-1)^{p+q}\; i^{p-p'} e^{-a-c} I_{q'}(c)\, \mathscr{L}^{s'}_{q'}(-a) \times \\ \times \sum_{u} (2u+1)\, i^u j_u(b\nu_j)\, \overline{P}^u_{p-p',0}(\cos\beta_{\nu_j}) \begin{pmatrix} u & s' & s \\ p-p' & p' & -p \end{pmatrix} \begin{pmatrix} u & s' & s \\ 0 & q & -q \end{pmatrix} \quad \text{(G26)}$$

Where we have used the following property of the Wigner 3-j symbols:

$$\begin{pmatrix} r & 0 & u \\ p & 0 & -p \end{pmatrix} = \delta_{ru} \times \frac{(-1)^{u-p}}{\sqrt{2u+1}} \quad \text{(G27)}$$

##### 5. Expansion coefficients of $T_{01}^{(2)}$ function

In order to derive a mathematical expression for the expansion coefficients of $T_{01}^{(2)}(\mathbf{R}_j, \mathbf{R}_{j+1}, \boldsymbol{\nu}_j)$ transfer-function, we need to do pretty much similar manipulations with Eq. (G18) as in the above section. Indeed, from Eq. (B17), Eq. (G1), Eq. (G2) and comments at the end of Appendix B it can be seen that the expansion series of $T_{01}^{(2)}(\mathbf{R}_j, \mathbf{R}_{j+1}, \boldsymbol{\nu}_j)$ function can be acquired from Eq. (G17)-Eq. (G18) via the following two steps: 1) by putting $a = a_{\rm pr}$, $c = c_{\rm pr}$, $\boldsymbol{\nu} = \boldsymbol{\nu}_j$, $\mathbf{A} = \mathbf{A}_{\rm in}$, and substituting $\mathbf{R} \to \mathbf{R}_j \mathbf{R}_{\eta_{{\rm in},j}}$ and $\mathbf{R}' \to \mathbf{R}_{j+1}$, where $\mathbf{R}_{\eta_{{\rm in},j}} = \mathbf{R}_{\eta_{{\rm in},j}}(\eta_{{\rm in},j}, 0, 0)$, and then 2) by multiplying the whole expression by $e^{\beta(\mu_{\rm pr}+\mu_{\rm off})}$ and performing integration over the angle $\eta_{{\rm in},j}$. As can be seen from Eq. (F1) and Eq. (G17), the last integration step results in nullification of the index $q_j$ and multiplication of the whole expression by $2\pi$ prefactor, leading us to the following formula for the expansion coefficients of $T_{01}^{(2)}(\mathbf{R}_j, \mathbf{R}_{j+1}, \boldsymbol{\nu}_j)$ function:

$$\big[T_{01}^{(2)}(\boldsymbol{\nu}_j)\big]_{p,\,q,\,s}^{p',\,q',\,s'} = \delta_{q0} \times 8\pi^3 \sqrt{(2s{+}1)(2s'{+}1)}\; (-1)^{p'}\, e^{\beta(\mu_{\rm pr}+\mu_{\rm off})-a_{\rm pr}-c_{\rm pr}} I_{q'}(c_{\rm pr})\, \mathscr{L}_{q'}^{s'}(-a_{\rm pr})\, D_{0,q'}^{s'}(\mathbf{A}_{\rm in}) \times$$
$$\times \sum_{r,t,u} (2r{+}1)\,(2t{+}1)\,(2u{+}1)\, i^r j_r(b\nu_j)\, i_t(\beta b f)\, \overline{D}_{p-p',0}^{\,r}(\mathbf{A}_{\nu_j}) \times \begin{pmatrix} r & t & u \\ p-p' & 0 & p'-p \end{pmatrix} \times$$
$$\times \begin{pmatrix} r & t & u \\ 0 & 0 & 0 \end{pmatrix} \begin{pmatrix} u & s' & s \\ p-p' & p' & -p \end{pmatrix} \begin{pmatrix} u & s' & s \\ 0 & 0 & 0 \end{pmatrix} \tag{G28}$$

Where, as before, $s = s_j$, $p = p_j$, $q = q_j$, $s' = s_{j+1}$, $p' = p_{j+1}$ and $q' = q_{j+1}$.

As for modified expansion coefficients discussed in Appendix H, by applying Eq. (H1) and Eq. (H2) to the above expression, it can be shown that they are equal to:

$$\big[\widetilde{T}_{01}^{(2)}(\boldsymbol{\nu}_j)\big]_{p,\,q,\,s}^{p',\,q',\,s'} = \delta_{q0} \times 8\pi^3 \sqrt{(2s{+}1)(2s'{+}1)}\; (-1)^{p'}\, i^{p-p'} e^{\beta(\mu_{\rm pr}+\mu_{\rm off})-a_{\rm pr}-c_{\rm pr}} I_{q'}(c_{\rm pr})\, \mathscr{L}_{q'}^{s'}(-a_{\rm pr})\, D_{0,q'}^{s'}(\mathbf{A}_{\rm in}) \times$$
$$\times \sum_{r,t,u} (2r{+}1)\,(2t{+}1)\,(2u{+}1)\, i^r j_r(b\nu_j)\, i_t(\beta b f)\, \overline{P}_{p-p',0}^{\,r}(\cos\beta_{\nu_j}) \times \begin{pmatrix} r & t & u \\ p-p' & 0 & p'-p \end{pmatrix} \times$$
$$\times \begin{pmatrix} r & t & u \\ 0 & 0 & 0 \end{pmatrix} \begin{pmatrix} u & s' & s \\ p-p' & p' & -p \end{pmatrix} \begin{pmatrix} u & s' & s \\ 0 & 0 & 0 \end{pmatrix} \tag{G29}$$

Where $\beta_{\nu_j}$ is the polar Euler angle of the wave-vector $\boldsymbol{\nu}_j$.

By taking into account Eq. (G27), it is also not very hard to find that in the case of zero stretching force ($f = 0$) Eq. (G29) turns into:

$$\big[\widetilde{T}_{01}^{(2)}(\boldsymbol{\nu}_j)\big]_{p,\,q,\,s}^{p',\,q',\,s'} = \delta_{q0} \times 8\pi^3 \sqrt{(2s{+}1)(2s'{+}1)}\; (-1)^{p}\, i^{p-p'} e^{\beta(\mu_{\rm pr}+\mu_{\rm off})-a_{\rm pr}-c_{\rm pr}} I_{q'}(c_{\rm pr})\, \mathscr{L}_{q'}^{s'}(-a_{\rm pr})\, D_{0,q'}^{s'}(\mathbf{A}_{\rm in}) \times$$
$$\times \sum_{u} (2u{+}1)\, i^u j_u(b\nu_j)\, \overline{P}_{p-p',0}^{\,u}(\cos\beta_{\nu_j}) \begin{pmatrix} u & s' & s \\ p-p' & p' & -p \end{pmatrix} \begin{pmatrix} u & s' & s \\ 0 & 0 & 0 \end{pmatrix} \tag{G30}$$

##### 6. Expansion coefficients of $T_{K0}^{(2)}$ function

Formula for the expansion coefficients of $T_{K0}^{(2)}(\mathbf{R}_j, \mathbf{R}_{j+1}, \boldsymbol{\nu}_j)$ function is derived in exactly the same way as in the case of $T_{01}^{(2)}(\mathbf{R}_j, \mathbf{R}_{j+1}, \boldsymbol{\nu}_j)$ function, which has been discussed in the previous section. The only major difference is that we do not need to multiply $F_1$ function by $e^{\beta(\mu_{\rm pr}+\mu_{\rm off})}$ prefactor and perform the integration step. Indeed, since in the general case $E_{K0}$ energy term is identical to $E_{30}$ term from Eq. (B17), it can be seen from Eq. (G1) and Eq. (G2) that the expansion coefficients of $T_{K0}^{(2)}(\mathbf{R}_j, \mathbf{R}_{j+1}, \boldsymbol{\nu}_j)$ function can be found from the expansion series of $F_1$ function by making the following parameter substitutions: $a = a_{\rm pr}$, $c = c_{\rm pr}$, $\boldsymbol{\nu} = \boldsymbol{\nu}_j$, $\mathbf{A} = \mathbf{A}_{\rm out}$ and $b = \frac{r_{\rm pr}}{K}$. Thus, from Eq. (G18) we have:

$$\big[T^{(2)}_{K0}(\boldsymbol{\nu}_j)\big]^{p',q',s'}_{p,\,q,\,s} = 4\pi^2\sqrt{(2s{+}1)(2s'{+}1)}\times(-1)^{p'+q}\,e^{-a_{\rm pr}-c_{\rm pr}}I_{q'}(c_{\rm pr})\,\mathscr{L}^{s'}_{q'}(-a_{\rm pr})\,D^{s'}_{q,q'}(\mathbf{A}_{\rm out})\times$$

$$\times\sum_{r,t,u}(2r{+}1)\,(2t{+}1)\,(2u{+}1)\,i^r j_r\big(\nu_j\tfrac{r_{\rm pr}}{K}\big)\,i_t\big(\beta f\tfrac{r_{\rm pr}}{K}\big)\,\overline{D}^r_{p-p',0}\big(\mathbf{A}_{\nu_j}\big)\times$$

$$\times\begin{pmatrix} r & t & u\\ p-p' & 0 & p'-p\end{pmatrix}\begin{pmatrix} r & t & u\\ 0 & 0 & 0\end{pmatrix}\begin{pmatrix} u & s' & s\\ p-p' & p' & -p\end{pmatrix}\begin{pmatrix} u & s' & s\\ 0 & q & -q\end{pmatrix} \tag{G31}$$

Where $s=s_j$, $p=p_j$, $q=q_j$, $s'=s_{j+1}$, $p'=p_{j+1}$ and $q'=q_{j+1}$.

Based on Eq. (G31), it is easy to obtain the following mathematical expression for the modified expansion coefficients defined by Eq. (H2):

$$\big[\widetilde{T}^{(2)}_{K0}(\boldsymbol{\nu}_j)\big]^{p',q',s'}_{p,\,q,\,s} = 4\pi^2\sqrt{(2s{+}1)(2s'{+}1)}\times(-1)^{p'+q}\;i^{p-p'}e^{-a_{\rm pr}-c_{\rm pr}}I_{q'}(c_{\rm pr})\,\mathscr{L}^{s'}_{q'}(-a_{\rm pr})\,D^{s'}_{q,q'}(\mathbf{A}_{\rm out})\times$$

$$\times\sum_{r,t,u}(2r{+}1)\,(2t{+}1)\,(2u{+}1)\,i^r j_r\big(\nu_j\tfrac{r_{\rm pr}}{K}\big)\,i_t\big(\beta f\tfrac{r_{\rm pr}}{K}\big)\,\overline{P}^r_{p-p',0}\big(\cos\beta_{\nu_j}\big)\times$$

$$\times\begin{pmatrix} r & t & u\\ p-p' & 0 & p'-p\end{pmatrix}\begin{pmatrix} r & t & u\\ 0 & 0 & 0\end{pmatrix}\begin{pmatrix} u & s' & s\\ p-p' & p' & -p\end{pmatrix}\begin{pmatrix} u & s' & s\\ 0 & q & -q\end{pmatrix} \tag{G32}$$

Where $\beta_{\nu_j}$ is the polar Euler angle of the wave-vector $\boldsymbol{\nu}_j$.

Furthermore, with the help of Eq. (G27), it can be shown that in the case of zero stretching force ($f=0$) the above formula reduces to:

$$\big[\widetilde{T}^{(2)}_{K0}(\boldsymbol{\nu}_j)\big]^{p',q',s'}_{p,\,q,\,s} = 4\pi^2\sqrt{(2s{+}1)(2s'{+}1)}\times(-1)^{p+q}\;i^{p-p'}e^{-a_{\rm pr}-c_{\rm pr}}I_{q'}(c_{\rm pr})\,\mathscr{L}^{s'}_{q'}(-a_{\rm pr})\,D^{s'}_{q,q'}(\mathbf{A}_{\rm out})\times$$

$$\times\sum_{u}(2u{+}1)\,i^u j_u\big(\nu_j\tfrac{r_{\rm pr}}{K}\big)\,\overline{P}^u_{p-p',0}\big(\cos\beta_{\nu_j}\big)\begin{pmatrix} u & s' & s\\ p-p' & p' & -p\end{pmatrix}\begin{pmatrix} u & s' & s\\ 0 & q & -q\end{pmatrix} \tag{G33}$$

##### 7. Expansion coefficients of $T^{(2)}_{12}$, $T^{(2)}_{23}$, ..., $T^{(2)}_{K-1\,K}$ functions

As for expansion coefficients of $T^{(2)}_{12}(\mathbf{R}_j,\mathbf{R}_{j+1},\boldsymbol{\nu}_j) = T^{(2)}_{23}(\mathbf{R}_j,\mathbf{R}_{j+1},\boldsymbol{\nu}_j) = \ldots = T^{(2)}_{K-1\,K}(\mathbf{R}_j,\mathbf{R}_{j+1},\boldsymbol{\nu}_j)$ transfer-functions, they can be acquired from Eq. (G23) by noting that all these functions are identical to $F_2$ from Eq. (G2) up to parameter values. Indeed, in the general case, $E_{12}=E_{23}=\ldots=E_{K-1\,K}=-\frac{1}{\beta}\ln\delta(\mathbf{R}_j-\mathbf{R}_{j+1})-\frac{r_{\rm pr}}{K}f[\mathbf{z}_0\cdot\mathbf{R}_j\mathbf{z}_0]$, see, for example, Eq. (B17). After substituting this formula into Eq. (G1), it is not hard to check that $T^{(2)}_{12}$, $T^{(2)}_{23}$, ..., $T^{(2)}_{K\,K-1}$ transfer-functions can be obtained from $F_2$ function defined by Eq. (G2) by making the following change of the model parameters: $b\to\frac{r_{\rm pr}}{K}$, $\mathbf{R}\to\mathbf{R}_j$, $\mathbf{R}'\to\mathbf{R}_{j+1}$ and $\boldsymbol{\nu}\to\boldsymbol{\nu}_j$. By doing the same parameter change in Eq. (G23), we get the next result for the expansion coefficients of $T^{(2)}_{12}$, $T^{(2)}_{23}$, ..., $T^{(2)}_{K\,K-1}$ transfer-functions:

$$\big[T^{(2)}_{12}(\boldsymbol{\nu}_j)\big]^{p',q',s'}_{p,\,q,\,s} = \big[T^{(2)}_{23}(\boldsymbol{\nu}_j)\big]^{p',q',s'}_{p,\,q,\,s} = \ldots = \big[T^{(2)}_{K-1\,K}(\boldsymbol{\nu}_j)\big]^{p',q',s'}_{p,\,q,\,s} =$$

$$\delta_{qq'}\times\sqrt{(2s{+}1)(2s'{+}1)}\times(-1)^{p'+q}\sum_{r,t,u}(2r{+}1)\,(2t{+}1)\,(2u{+}1)\,i^r j_r\big(\nu_j\tfrac{r_{\rm pr}}{K}\big)\,i_t\big(\beta f\tfrac{r_{\rm pr}}{K}\big)\times$$

$$\times\overline{D}^r_{p-p',0}\big(\mathbf{A}_{\nu_j}\big)\times\begin{pmatrix} r & t & u\\ p-p' & 0 & p'-p\end{pmatrix}\begin{pmatrix} r & t & u\\ 0 & 0 & 0\end{pmatrix}\begin{pmatrix} u & s' & s\\ p-p' & p' & -p\end{pmatrix}\begin{pmatrix} u & s' & s\\ 0 & q & -q\end{pmatrix} \tag{G34}$$

Where, as before, $s=s_j$, $p=p_j$, $q=q_j$, $s'=s_{j+1}$, $p'=p_{j+1}$ and $q'=q_{j+1}$.

Correspondingly, for the modified expansion coefficients discussed in Appendix H we have:

$$\big[\widetilde{T}^{(2)}_{12}(\boldsymbol{\nu}_j)\big]^{p',\,q',\,s'}_{p,\,q,\,s} = \big[\widetilde{T}^{(2)}_{23}(\boldsymbol{\nu}_j)\big]^{p',\,q',\,s'}_{p,\,q,\,s} = \ldots = \big[\widetilde{T}^{(2)}_{K-1\,K}(\boldsymbol{\nu}_j)\big]^{p',\,q',\,s'}_{p,\,q,\,s} =$$

$$\delta_{qq'} \times \sqrt{(2s{+}1)(2s'{+}1)} \times (-1)^{p'+q}\; i^{p-p'} \sum_{r,t,u} (2r{+}1)\,(2t{+}1)\,(2u{+}1)\, i^r j_r\big(\nu_j \tfrac{r_{\rm pr}}{K}\big)\, i_t\big(\beta f \tfrac{r_{\rm pr}}{K}\big) \times$$

$$\times\, \overline{P}^{r}_{p-p',0}(\cos\beta_{\nu_j}) \times \begin{pmatrix} r & t & u \\ p-p' & 0 & p'-p \end{pmatrix} \begin{pmatrix} r & t & u \\ 0 & 0 & 0 \end{pmatrix} \begin{pmatrix} u & s' & s \\ p-p' & p' & -p \end{pmatrix} \begin{pmatrix} u & s' & s \\ 0 & q & -q \end{pmatrix} \quad \text{(G35)}$$

Where $\beta_{\nu_j}$ is the polar Euler angle of the wave-vector $\boldsymbol{\nu}_j$.

In the case of zero force ($f = 0$), it can be shown with the help of Eq. (G27) that the above formula turns into:

$$\big[\widetilde{T}^{(2)}_{12}(\boldsymbol{\nu}_j)\big]^{p',\,q',\,s'}_{p,\,q,\,s} = \big[\widetilde{T}^{(2)}_{23}(\boldsymbol{\nu}_j)\big]^{p',\,q',\,s'}_{p,\,q,\,s} = \ldots = \big[\widetilde{T}^{(2)}_{K-1\,K}(\boldsymbol{\nu}_j)\big]^{p',\,q',\,s'}_{p,\,q,\,s} = \delta_{qq'} \times \sqrt{(2s{+}1)(2s'{+}1)} \times (-1)^{p+q}\; i^{p-p'} \times$$

$$\times \sum_{u} (2u{+}1)\, i^u j_u\big(\nu_j \tfrac{r_{\rm pr}}{K}\big)\, \overline{P}^{u}_{p-p',0}(\cos\beta_{\nu_j}) \begin{pmatrix} u & s' & s \\ p-p' & p' & -p \end{pmatrix} \begin{pmatrix} u & s' & s \\ 0 & q & -q \end{pmatrix} \quad \text{(G36)}$$

##### 8. Expansion coefficients of $\xi_0^{(2)}$ function

Besides $T^{(2)}_{k_j k_{j+1}}(\mathbf{R}_j, \mathbf{R}_{j+1}, \boldsymbol{\nu}_j)$ transfer-functions, we also need to know expansion coefficients of $\xi_0^{(2)}(\mathbf{R}_N, \mathbf{R}_1, \boldsymbol{\nu}_N)$ boundary condition function defined by Eq. (G1) in order to able to calculate the DNA partition function. Since in this study we do not put any additional constraints on orientations of the DNA end segments apart from the confinement potential, $U_{R_{\rm nucl}}$, for $\xi(\mathbf{R}_N, \mathbf{R}_1)$ function from Eq. (G1) we have: $\xi(\mathbf{R}_N, \mathbf{R}_1) = 1$.

It is clear then that the expansion coefficients of $\xi_0^{(2)}(\mathbf{R}_N, \mathbf{R}_1, \boldsymbol{\nu}_N)$ function can be acquired from Eq. (G8) by putting $\boldsymbol{\nu} = \boldsymbol{\nu}_N$, $\mathbf{R} = \mathbf{R}_N$ and expressing the obtained result in a form similar to Eq. (C30). By doing so, we get:

$$\big[\xi_0^{(2)}(\boldsymbol{\nu}_N)\big]^{p',\,q',\,s'}_{p,\,q,\,s} = \delta_{s'0}\,\delta_{p'0}\,\delta_{q'0}\,\delta_{q0} \times 8\pi^2 \sqrt{2s{+}1}\;(-1)^p \times \sum_{r,t} (2r{+}1)\,(2t{+}1)\, i^r j_r(b\nu_N)\, i_t(\beta b f) \times$$

$$\times\, \overline{D}^{r}_{p,0}(\mathbf{A}_{\nu_N}) \begin{pmatrix} r & t & s \\ p & 0 & -p \end{pmatrix} \begin{pmatrix} r & t & s \\ 0 & 0 & 0 \end{pmatrix} \quad \text{(G37)}$$

Where $s = s_N$, $p = p_N$, $q = q_N$, $s' = s_1$, $p' = p_1$ and $q' = q_1$.

From Eq. (G37), it is also not very hard to find the following formula for the modified expansion coefficients defined by Eq. (H2):

$$\big[\widetilde{\xi}_0^{(2)}(\boldsymbol{\nu}_N)\big]^{p',\,q',\,s'}_{p,\,q,\,s} = \delta_{s'0}\,\delta_{p'0}\,\delta_{q'0}\,\delta_{q0} \times 8\pi^2 \sqrt{2s{+}1}\;(-1)^p\; i^p \times \sum_{r,t} (2r{+}1)\,(2t{+}1)\, i^r j_r(b\nu_N)\, i_t(\beta b f) \times$$

$$\times\, \overline{P}^{r}_{p,0}(\cos\beta_{\nu_N}) \begin{pmatrix} r & t & s \\ p & 0 & -p \end{pmatrix} \begin{pmatrix} r & t & s \\ 0 & 0 & 0 \end{pmatrix} \quad \text{(G38)}$$

Where $\beta_{\nu_N}$ is the polar Euler angle of the wave-vector $\boldsymbol{\nu}_N$.

In the case of zero stretching force ($f = 0$), the above expression reduces to:

$$\big[\widetilde{\xi}_0^{(2)}(\boldsymbol{\nu}_N)\big]^{p',\,q',\,s'}_{p,\,q,\,s} = \delta_{s'0}\,\delta_{p'0}\,\delta_{q'0}\,\delta_{q0} \times 8\pi^2 \sqrt{2s{+}1}\; i^{s+p} j_s(b\nu_N)\, \overline{P}^{s}_{p,0}(\cos\beta_{\nu_N}) \quad \text{(G39)}$$

Having at hand expansion coefficients of $T^{(2)}_{k_j k_{j+1}}$ and $\xi_0^{(2)}$ functions, we can move to the next step – derivation of mathematical formulas for the elements of the higher-level matrix-blocks, $\mathbf{T}^{(5)}_{k_j k_{j+1}}$ and $\boldsymbol{\xi}_0^{(4)}$, which constitute the DNA transfer-matrix, $\mathbf{L}$, and the boundary condition vector, $\mathbf{Y}$.

##### 9. Elements of $\mathbf{T}_{00}^{(5)}$ matrix

As before, let's start with $\mathbf{T}_{00}^{(5)}$ matrix first. From Eq. (C34) it can be seen that in order to find elements of this matrix, we need to know the expansion coefficients of $T_{00}^{(4)}(\mathbf{R}_j, \mathbf{R}_{j+1}, \boldsymbol{\nu}_j, \boldsymbol{\nu}_{j+1})$ transfer-function, which can be found by combining together Eq. (C14), Eq. (C43) and Eq. (G24):

$$\begin{aligned}\left[T_{00}^{(4)}(\boldsymbol{\nu}_j, \boldsymbol{\nu}_{j+1})\right]_{p,\,q,\,s}^{p',\,q',\,s'} = \delta_{qq'} \times \Phi_{00}(\boldsymbol{\nu}_j - \boldsymbol{\nu}_{j+1}) \times 4\pi^2 \sqrt{(2s{+}1)(2s'{+}1)}\; (-1)^{p'+q}\, e^{-a-c} I_{q'}(c)\, \mathscr{L}_{q'}^{s'}(-a) \times \\ \times \sum_{r,t,u} (2r{+}1)\,(2t{+}1)\,(2u{+}1)\, i^r j_r(b\nu_j)\, i_t(\beta b f)\, \overline{D}_{p-p',0}^{r}(\mathbf{A}_{\nu_j}) \times \\ \times \begin{pmatrix} r & t & u \\ p-p' & 0 & p'-p \end{pmatrix} \begin{pmatrix} r & t & u \\ 0 & 0 & 0 \end{pmatrix} \begin{pmatrix} u & s' & s \\ p-p' & p' & -p \end{pmatrix} \begin{pmatrix} u & s' & s \\ 0 & q & -q \end{pmatrix}\end{aligned} \tag{G40}$$

Where for the sake of simplicity we used the following notations: $s = s_j$, $p = p_j$, $q = q_j$, $s' = s_{j+1}$, $p' = p_{j+1}$ and $q' = q_{j+1}$.

From the above formula and Eq. (C34), it then easy to see that the elements of $\mathbf{T}_{00}^{(5)}(\boldsymbol{\nu}_j, \boldsymbol{\nu}_{j+1})$ matrix-block are equal to:

$$\begin{aligned}\left[\mathbf{T}_{00}^{(5)}(\boldsymbol{\nu}_j, \boldsymbol{\nu}_{j+1})\right]_{vv'} = \Phi_{00}(\boldsymbol{\nu}_j - \boldsymbol{\nu}_{j+1}) \times 4\pi^2 \sqrt{(2s{+}1)(2s'{+}1)}\; (-1)^{p'}\, e^{-a-c} I_0(c)\, \mathscr{L}_0^{s'}(-a) \times \\ \times \sum_{r,t,u} (2r{+}1)\,(2t{+}1)\,(2u{+}1)\, i^r j_r(b\nu_j)\, i_t(\beta b f)\, \overline{D}_{p-p',0}^{r}(\mathbf{A}_{\nu_j}) \times \\ \times \begin{pmatrix} r & t & u \\ p-p' & 0 & p'-p \end{pmatrix} \begin{pmatrix} r & t & u \\ 0 & 0 & 0 \end{pmatrix} \begin{pmatrix} u & s' & s \\ p-p' & p' & -p \end{pmatrix} \begin{pmatrix} u & s' & s \\ 0 & 0 & 0 \end{pmatrix}\end{aligned} \tag{G41}$$

Where $v = p + s(s+1)$ and $v' = p' + s'(s'+1)$.

By using the axial symmetry of the system and by utilizing the Dirac comb function in derivation of the DNA transfer-matrix, it can be shown that Eq. (G41) leads to the following expression for the elements of the reduced matrix-block $\widetilde{\mathbf{T}}_{00}^{(5)}(\mathbf{G}_{\mathbf{m}_j}, \mathbf{G}_{\mathbf{m}_{j+1}})$ discussed at the end of Appendix H [see Appendix D and Appendix H for more details]:

$$\begin{aligned}\left[\widetilde{\mathbf{T}}_{00}^{(5)}(\mathbf{G}_{\mathbf{m}_j}, \mathbf{G}_{\mathbf{m}_{j+1}})\right]_{vv'} = \widetilde{\Phi}_{00}(p', \mathbf{G}_{\mathbf{m}_j}, \mathbf{G}_{\mathbf{m}_{j+1}}) \times 4\pi^2 \sqrt{(2s{+}1)(2s'{+}1)}\; (-1)^{p'}\, i^{p-p'} e^{-a-c} I_0(c)\, \mathscr{L}_0^{s'}(-a) \times \\ \times \sum_{r,t,u} (2r{+}1)\,(2t{+}1)\,(2u{+}1)\, i^r j_r(b\, G_{\mathbf{m}_j})\, i_t(\beta b f)\, \overline{P}_{p-p',0}^{r}(\cos\beta_{\mathbf{G}_{\mathbf{m}_j}}) \times \\ \times \begin{pmatrix} r & t & u \\ p-p' & 0 & p'-p \end{pmatrix} \begin{pmatrix} r & t & u \\ 0 & 0 & 0 \end{pmatrix} \begin{pmatrix} u & s' & s \\ p-p' & p' & -p \end{pmatrix} \begin{pmatrix} u & s' & s \\ 0 & 0 & 0 \end{pmatrix}\end{aligned} \tag{G42}$$

Where $\widetilde{\Phi}_{00}(p_{j+1}, \mathbf{G}_{\mathbf{m}_j}, \mathbf{G}_{\mathbf{m}_{j+1}})$ is a function defined by Eq. (H15). $G_{\mathbf{m}_j} = \|\mathbf{G}_{\mathbf{m}_j}\|$ and $\beta_{\mathbf{G}_{\mathbf{m}_j}}$ are the length and the polar Euler angle of the wave-vector $\mathbf{G}_{\mathbf{m}_j}$, respectively.

In the special case of zero stretching force ($f = 0$), Eq. (G42) with the help of Eq. (G27) transforms into:

$$\begin{aligned}\left[\widetilde{\mathbf{T}}_{00}^{(5)}(\mathbf{G}_{\mathbf{m}_j}, \mathbf{G}_{\mathbf{m}_{j+1}})\right]_{vv'} = \widetilde{\Phi}_{00}(p', \mathbf{G}_{\mathbf{m}_j}, \mathbf{G}_{\mathbf{m}_{j+1}}) \times 4\pi^2 \sqrt{(2s{+}1)(2s'{+}1)}\; (-1)^{p}\, i^{p-p'} e^{-a-c} I_0(c)\, \mathscr{L}_0^{s'}(-a) \times \\ \times \sum_{u} (2u{+}1)\, i^u j_u(b\, G_{\mathbf{m}_j})\, \overline{P}_{p-p',0}^{u}(\cos\beta_{\mathbf{G}_{\mathbf{m}_j}}) \begin{pmatrix} u & s' & s \\ p-p' & p' & -p \end{pmatrix} \begin{pmatrix} u & s' & s \\ 0 & 0 & 0 \end{pmatrix}\end{aligned} \tag{G43}$$

##### 10. Elements of $\mathbf{T}^{(5)}_{12}$, $\mathbf{T}^{(5)}_{23}$, ..., $\mathbf{T}^{(5)}_{K0}$ matrices

Similarly to the previous section, to find the elements of $\mathbf{T}^{(5)}_{12}(\boldsymbol{\nu}_j,\boldsymbol{\nu}_{j+1})$, $\mathbf{T}^{(5)}_{23}(\boldsymbol{\nu}_j,\boldsymbol{\nu}_{j+1})$, ..., $\mathbf{T}^{(5)}_{K0}(\boldsymbol{\nu}_j,\boldsymbol{\nu}_{j+1})$ matrices, we need to know in the first place the expansion coefficients of the corresponding level-four transfer-functions, $T^{(4)}_{12}(\mathbf{R}_j,\mathbf{R}_{j+1},\boldsymbol{\nu}_j,\boldsymbol{\nu}_{j+1})$, $T^{(4)}_{23}(\mathbf{R}_j,\mathbf{R}_{j+1},\boldsymbol{\nu}_j,\boldsymbol{\nu}_{j+1})$, ..., $T^{(4)}_{K0}(\mathbf{R}_j,\mathbf{R}_{j+1},\boldsymbol{\nu}_j,\boldsymbol{\nu}_{j+1})$, which can be easily obtained from Eq. (C43) and Eq. (G20) by casting the latter in the form of Eq. (F16). As a result, we get:

$$\big[T^{(4)}_{12}(\boldsymbol{\nu}_j,\boldsymbol{\nu}_{j+1})\big]^{p',q',s'}_{p,q,s} = \big[T^{(4)}_{23}(\boldsymbol{\nu}_j,\boldsymbol{\nu}_{j+1})\big]^{p',q',s'}_{p,q,s} = ... = \big[T^{(4)}_{K0}(\boldsymbol{\nu}_j,\boldsymbol{\nu}_{j+1})\big]^{p',q',s'}_{p,q,s} = \delta_{pp'}\delta_{qq'}\delta_{ss'}\,\delta(\boldsymbol{\nu}_j-\boldsymbol{\nu}_{j+1}) \tag{G44}$$

Where we used the following notations: $s=s_j$, $p=p_j$, $q=q_j$, $s'=s_{j+1}$, $p'=p_{j+1}$ and $q'=q_{j+1}$.

Combining the above formula with Eq. (C34), it immediately follows that:

$$\big[\mathbf{T}^{(5)}_{12}(\boldsymbol{\nu}_j,\boldsymbol{\nu}_{j+1})\big]_{vv'} = \big[\mathbf{T}^{(5)}_{23}(\boldsymbol{\nu}_j,\boldsymbol{\nu}_{j+1})\big]_{vv'} = ... = \big[\mathbf{T}^{(5)}_{K0}(\boldsymbol{\nu}_j,\boldsymbol{\nu}_{j+1})\big]_{vv'} = \delta_{pp'}\delta_{ss'}\,\delta(\boldsymbol{\nu}_j-\boldsymbol{\nu}_{j+1}) \tag{G45}$$

Here $v=p+s(s+1)$ and $v'=p'+s'(s'+1)$.

Furthermore, by making $\boldsymbol{\nu}_j\to\mathbf{G}_{\mathbf{m}_j}$, $\boldsymbol{\nu}_{j+1}\to\mathbf{G}_{\mathbf{m}_{j+1}}$ and $\delta(\boldsymbol{\nu}_j-\boldsymbol{\nu}_{j+1})\to\delta^{3\text{D}}_{\mathbf{m}_j\mathbf{m}_{j+1}}$ substitutions in the above equation, it is not very hard to show that the elements of the corresponding reduced matrix-blocks, which are discussed in Appendix H, are equal to [see Appendix D in Appendix H for more details]:

$$\big[\widetilde{\mathbf{T}}^{(5)}_{12}(\mathbf{G}_{\mathbf{m}_j},\mathbf{G}_{\mathbf{m}_{j+1}})\big]_{vv'} = \big[\widetilde{\mathbf{T}}^{(5)}_{23}(\mathbf{G}_{\mathbf{m}_j},\mathbf{G}_{\mathbf{m}_{j+1}})\big]_{vv'} = ... = \big[\widetilde{\mathbf{T}}^{(5)}_{K0}(\mathbf{G}_{\mathbf{m}_j},\mathbf{G}_{\mathbf{m}_{j+1}})\big]_{vv'} = \delta_{pp'}\delta_{ss'}\,\delta^{3\text{D}}_{\mathbf{m}_j\mathbf{m}_{j+1}} \tag{G46}$$

In other words, these matrices are nothing else but the identity matrix, $\mathbf{I}$, multiplied by $\delta^{3\text{D}}_{\mathbf{m}_j\mathbf{m}_{j+1}}$ Kronecker delta:

$$\widetilde{\mathbf{T}}^{(5)}_{12}(\mathbf{G}_{\mathbf{m}_j},\mathbf{G}_{\mathbf{m}_{j+1}}) = \widetilde{\mathbf{T}}^{(5)}_{23}(\mathbf{G}_{\mathbf{m}_j},\mathbf{G}_{\mathbf{m}_{j+1}}) = ... = \widetilde{\mathbf{T}}^{(5)}_{K0}(\mathbf{G}_{\mathbf{m}_j},\mathbf{G}_{\mathbf{m}_{j+1}}) = \delta^{3\text{D}}_{\mathbf{m}_j\mathbf{m}_{j+1}}\,\mathbf{I} \tag{G47}$$

##### 11. Elements of $\mathbf{T}^{(5)}_{01}$ matrix

To obtain a mathematical expression for the elements of $\mathbf{T}^{(5)}_{01}(\boldsymbol{\nu}_j,\boldsymbol{\nu}_{j+1})$ matrix, we will use the same approach as in Appendix C in our derivation of the DNA partition function. Namely, from Eq. (C43) it can be seen that $T^{(4)}_{01}(\boldsymbol{\nu}_j,\boldsymbol{\nu}_{j+1})$ transfer-function is defined as a multiple integral over the product of the level-two transfer-functions, $T^{(2)}_{nm}(\mathbf{R},\mathbf{R}',\boldsymbol{\nu}_j)$. By expanding all these $T^{(2)}_{nm}(\mathbf{R},\mathbf{R}',\boldsymbol{\nu}_j)$ functions in the form of Eq. (C27) and by using orthogonality of the Wigner D-functions [Eq. (F9)], it can be shown that the multiple integral in definition of $T^{(4)}_{01}(\boldsymbol{\nu}_j,\boldsymbol{\nu}_{j+1})$ reduces to a mere summation. As a result, we get the following formula for the expansion coefficients of $T^{(4)}_{01}(\boldsymbol{\nu}_j,\boldsymbol{\nu}_{j+1})$ transfer-function:

$$\big[T^{(4)}_{01}(\boldsymbol{\nu}_j,\boldsymbol{\nu}_{j+1})\big]^{p',q',s'}_{p,q,s} = \Phi_{K0}(\boldsymbol{\nu}_j-\boldsymbol{\nu}_{j+1})\times\sum_{\substack{r_1,t_1\\u_1}}\sum_{\substack{r_2,t_2\\u_2}}...\sum_{\substack{r_K,t_K\\u_K}}\big[T^{(2)}_{01}(\boldsymbol{\nu}_j)\big]^{r_1,t_1,u_1}_{p,q,s}\big[T^{(2)}_{12}(\boldsymbol{\nu}_j)\big]^{r_2,t_2,u_2}_{r_1,t_1,u_1}...\big[T^{(2)}_{K0}(\boldsymbol{\nu}_j)\big]^{p',q',s'}_{r_K,t_K,u_K} \tag{G48}$$

Where $s=s_j$, $p=p_j$, $q=q_j$, $s'=s_{j+1}$, $p'=p_{j+1}$ and $q'=q_{j+1}$.

Eq. (G48) can be further simplified by noting that all of the expansion coefficients of $T^{(2)}_{12}(\mathbf{R},\mathbf{R}',\boldsymbol{\nu}_j)$, $T^{(2)}_{23}(\mathbf{R},\mathbf{R}',\boldsymbol{\nu}_j)$, ..., $T^{(2)}_{K-1\,K}(\mathbf{R},\mathbf{R}',\boldsymbol{\nu}_j)$ transfer-functions contain $\delta_{qq'}$ prefactor, see Eq. (G34). Similarly, expansion coefficients of $T^{(2)}_{01}(\mathbf{R},\mathbf{R}',\boldsymbol{\nu}_j)$ function include $\delta_{q0}$ prefactor, see Eq. (G28). Combining these two results together, it is not hard to check that:

$$\big[T^{(4)}_{01}(\boldsymbol{\nu}_j,\boldsymbol{\nu}_{j+1})\big]^{p',q',s'}_{p,q,s} = \delta_{q0}\times\Phi_{K0}(\boldsymbol{\nu}_j-\boldsymbol{\nu}_{j+1})\times\sum_{t,r_1,u_1}\sum_{r_2,u_2}...\sum_{r_K,u_K}\big[T^{(2)}_{01}(\boldsymbol{\nu}_j)\big]^{r_1,t,u_1}_{p,0,s}\big[T^{(2)}_{12}(\boldsymbol{\nu}_j)\big]^{r_2,t,u_2}_{r_1,t,u_1}...\big[T^{(2)}_{K0}(\boldsymbol{\nu}_j)\big]^{p',q',s'}_{r_K,t,u_K} \tag{G49}$$

From the above formula and Eq. (C34), it is then straightforward to find the elements of $\mathbf{T}_{01}^{(5)}(\boldsymbol{\nu}_j, \boldsymbol{\nu}_{j+1})$ matrix:

$$\left[\mathbf{T}_{01}^{(5)}(\boldsymbol{\nu}_j, \boldsymbol{\nu}_{j+1})\right]_{vv'} = \Phi_{K0}(\boldsymbol{\nu}_j - \boldsymbol{\nu}_{j+1}) \times \sum_{t, r_1, u_1} \sum_{r_2, u_2} \dots \sum_{r_K, u_K} \left[T_{01}^{(2)}(\boldsymbol{\nu}_j)\right]_{p,\,0,\,s}^{r_1, t, u_1} \left[T_{12}^{(2)}(\boldsymbol{\nu}_j)\right]_{r_1, t, u_1}^{r_2, t, u_2} \dots \left[T_{K0}^{(2)}(\boldsymbol{\nu}_j)\right]_{r_K, t, u_K}^{p',\,0,\,s'} \tag{G50}$$

Here $v = p + s(s+1)$ and $v' = p' + s'(s'+1)$.

For the purpose of calculations, it is convenient to recast Eq. (G50) in a more compact matrix form:

$$\mathbf{T}_{01}^{(5)}(\boldsymbol{\nu}_j, \boldsymbol{\nu}_{j+1}) = \Phi_{K0}(\boldsymbol{\nu}_j - \boldsymbol{\nu}_{j+1}) \times \sum_t \mathbf{T}_{01}^{(2)}(\boldsymbol{\nu}_j, t)\, \mathbf{T}_{12}^{(2)}(\boldsymbol{\nu}_j, t)\, \mathbf{T}_{23}^{(2)}(\boldsymbol{\nu}_j, t) \dots \mathbf{T}_{K0}^{(2)}(\boldsymbol{\nu}_j, t)\,, \tag{G51}$$

Where matrices $\mathbf{T}_{01}^{(2)}(\boldsymbol{\nu}_j, t)$, $\mathbf{T}_{12}^{(2)}(\boldsymbol{\nu}_j, t)$, $\mathbf{T}_{23}^{(2)}(\boldsymbol{\nu}_j, t)$, ..., $\mathbf{T}_{K0}^{(2)}(\boldsymbol{\nu}_j, t)$ are defined by the following equations:

$$\begin{aligned}
\left[\mathbf{T}_{01}^{(2)}(\boldsymbol{\nu}_j, t)\right]_{vv'} &= \left[T_{01}^{(2)}(\boldsymbol{\nu}_j)\right]_{p,\,0,\,s}^{p',\,t,\,s'} \\
\left[\mathbf{T}_{12}^{(2)}(\boldsymbol{\nu}_j, t)\right]_{vv'} &= \left[T_{12}^{(2)}(\boldsymbol{\nu}_j)\right]_{p,\,t,\,s}^{p',\,t,\,s'} \\
\left[\mathbf{T}_{23}^{(2)}(\boldsymbol{\nu}_j, t)\right]_{vv'} &= \left[T_{23}^{(2)}(\boldsymbol{\nu}_j)\right]_{p,\,t,\,s}^{p',\,t,\,s'} \\
&\vdots \\
\left[\mathbf{T}_{K0}^{(2)}(\boldsymbol{\nu}_j, t)\right]_{vv'} &= \left[T_{K0}^{(2)}(\boldsymbol{\nu}_j)\right]_{p,\,t,\,s}^{p',\,0,\,s'}
\end{aligned} \tag{G52}$$

Here, as before, $v = p + s(s+1)$ and $v' = p' + s'(s'+1)$.

Furthermore, by repeating the above derivation in the case of the modified expansion coefficients defined by Eq. (G29), Eq. (G32) and Eq. (G35), one can arrive to the next formula for the elements of $\widetilde{\mathbf{T}}_{01}^{(5)}(\mathbf{G}_{\mathbf{m}_j}, \mathbf{G}_{\mathbf{m}_{j+1}})$ matrix discussed in Appendix H:

$$\left[\widetilde{\mathbf{T}}_{01}^{(5)}(\mathbf{G}_{\mathbf{m}_j}, \mathbf{G}_{\mathbf{m}_{j+1}})\right]_{vv'} = \widetilde{\Phi}_{K0}(p', \mathbf{G}_{\mathbf{m}_j}, \mathbf{G}_{\mathbf{m}_{j+1}}) \times \sum_t \left[\widetilde{\mathbf{T}}_{01}^{(2)}(\mathbf{G}_{\mathbf{m}_j}, t)\, \widetilde{\mathbf{T}}_{12}^{(2)}(\mathbf{G}_{\mathbf{m}_j}, t)\, \widetilde{\mathbf{T}}_{23}^{(2)}(\mathbf{G}_{\mathbf{m}_j}, t) \dots \widetilde{\mathbf{T}}_{K0}^{(2)}(\mathbf{G}_{\mathbf{m}_j}, t)\right]_{vv'} \tag{G53}$$

Where function $\widetilde{\Phi}_{K0}(p', \mathbf{G}_{\mathbf{m}_j}, \mathbf{G}_{\mathbf{m}_{j+1}})$ is defined by Eq. (H15), and matrices $\widetilde{\mathbf{T}}_{01}^{(2)}(\mathbf{G}_{\mathbf{m}_j}, t)$, $\widetilde{\mathbf{T}}_{12}^{(2)}(\mathbf{G}_{\mathbf{m}_j}, t)$, $\widetilde{\mathbf{T}}_{23}^{(2)}(\mathbf{G}_{\mathbf{m}_j}, t)$, ... , $\widetilde{\mathbf{T}}_{K0}^{(2)}(\mathbf{G}_{\mathbf{m}_j}, t)$ are given by the following equations:

$$\begin{aligned}
\left[\widetilde{\mathbf{T}}_{01}^{(2)}(\mathbf{G}_{\mathbf{m}_j}, t)\right]_{vv'} &= \left[\widetilde{T}_{01}^{(2)}(\mathbf{G}_{\mathbf{m}_j})\right]_{p,\,0,\,s}^{p',\,t,\,s'} \\
\left[\widetilde{\mathbf{T}}_{12}^{(2)}(\mathbf{G}_{\mathbf{m}_j}, t)\right]_{vv'} &= \left[\widetilde{T}_{12}^{(2)}(\mathbf{G}_{\mathbf{m}_j})\right]_{p,\,t,\,s}^{p',\,t,\,s'} \\
\left[\widetilde{\mathbf{T}}_{23}^{(2)}(\mathbf{G}_{\mathbf{m}_j}, t)\right]_{vv'} &= \left[\widetilde{T}_{23}^{(2)}(\mathbf{G}_{\mathbf{m}_j})\right]_{p,\,t,\,s}^{p',\,t,\,s'} \\
&\vdots \\
\left[\widetilde{\mathbf{T}}_{K0}^{(2)}(\mathbf{G}_{\mathbf{m}_j}, t)\right]_{vv'} &= \left[\widetilde{T}_{K0}^{(2)}(\mathbf{G}_{\mathbf{m}_j})\right]_{p,\,t,\,s}^{p',\,0,\,s'}
\end{aligned} \tag{G54}$$

##### 12. Elements of $\boldsymbol{\xi}_0^{(4)}$ matrix

As for the elements of $\boldsymbol{\xi}_0^{(4)}(\boldsymbol{\nu}_N, \boldsymbol{\nu}_1)$ matrix, from Eq. (C35) it can be seen that to find them, we need to know first the expansion coefficients of $\xi_0^{(3)}(\mathbf{R}_N, \mathbf{R}_1, \boldsymbol{\nu}_N, \boldsymbol{\nu}_1)$ boundary condition function, which can be easily obtained by combining together Eq. (C14) and Eq. (G37):

$$\begin{aligned}
\left[\xi_0^{(3)}(\boldsymbol{\nu}_N, \boldsymbol{\nu}_1)\right]_{p,\,q,\,s}^{p',\,q',\,s'} &= \delta_{s'0}\, \delta_{p'0}\, \delta_{q'0}\, \delta_{q0} \times \delta(\boldsymbol{\nu}_1)\; \Phi_{00}(\boldsymbol{\nu}_N) \times 8\pi^2 \sqrt{2s+1}\; (-1)^p \times \\
&\times \sum_{r,t} (2r+1)\,(2t+1)\, i^r j_r(b\nu_N)\, i_t(\beta b f)\, \overline{D}_{p,0}^{r}(\mathbf{A}_{\nu_N}) \begin{pmatrix} r & t & s \\ p & 0 & -p \end{pmatrix} \begin{pmatrix} r & t & s \\ 0 & 0 & 0 \end{pmatrix}
\end{aligned} \tag{G55}$$

Where $s = s_N$, $p = p_N$, $q = q_N$, $s' = s_1$, $p' = p_1$ and $q' = q_1$.

Thus, for the elements of $\boldsymbol{\xi}_0^{(4)}(\boldsymbol{\nu}_N, \boldsymbol{\nu}_1)$ matrix we have:

$$\left[\boldsymbol{\xi}_0^{(4)}(\boldsymbol{\nu}_N, \boldsymbol{\nu}_1)\right]_{v_N v_1} = \delta_{s'0}\,\delta_{p'0} \times \delta(\boldsymbol{\nu}_1)\;\Phi_{00}(\boldsymbol{\nu}_N) \times 8\pi^2\sqrt{2s{+}1}\;(-1)^p \times \\ \times \sum_{r,t}(2r{+}1)\,(2t{+}1)\,i^r j_r(b\nu_N)\,i_t(\beta b f)\,\overline{D}^{\,r}_{p,0}(\mathbf{A}_{\nu_N}) \begin{pmatrix} r & t & s \\ p & 0 & -p \end{pmatrix} \begin{pmatrix} r & t & s \\ 0 & 0 & 0 \end{pmatrix} \tag{G56}$$

Where $v_1 = p_1 + s_1(s_1+1)$ and $v_N = p_N + s_N(s_N+1)$.

Similarly, the elements of the reduced boundary matrix, $\widetilde{\boldsymbol{\xi}}_0^{(4)}(\mathbf{G}_{\mathbf{m}_N}, \mathbf{G}_{\mathbf{m}_1})$, which is mentioned in Appendix H, are equal to:

$$\left[\widetilde{\boldsymbol{\xi}}_0^{(4)}(\mathbf{G}_{\mathbf{m}_N}, \mathbf{G}_{\mathbf{m}_1})\right]_{v_N v_1} = \delta_{s'0}\,\delta_{p'0} \times \delta^{3\mathrm{D}}_{\mathbf{m}_1 0}\;\widetilde{\Phi}_{00}(0, \mathbf{G}_{\mathbf{m}_N}, 0) \times 8\pi^2\sqrt{2s{+}1}\;(-1)^p\; i^p \times \\ \times \sum_{r,t}(2r{+}1)\,(2t{+}1)\,i^r j_r(b\,G_{\mathbf{m}_N})\,i_t(\beta b f)\,\overline{P}^{\,r}_{p,0}(\cos\beta_{\mathbf{G}_{\mathbf{m}_N}}) \begin{pmatrix} r & t & s \\ p & 0 & -p \end{pmatrix} \begin{pmatrix} r & t & s \\ 0 & 0 & 0 \end{pmatrix} \tag{G57}$$

Where $\widetilde{\Phi}_{00}(0, \mathbf{G}_{\mathbf{m}_N}, 0)$ is a function defined by Eq. (H15). $G_{\mathbf{m}_N} = \|\mathbf{G}_{\mathbf{m}_N}\|$ and $\beta_{\mathbf{G}_{\mathbf{m}_N}}$ are the length and the polar Euler angle of the wave-vector $\mathbf{G}_{\mathbf{m}_N}$, respectively.

In the special case of zero stretching force ($f = 0$), Eq. (G57) transforms into:

$$\left[\widetilde{\boldsymbol{\xi}}_0^{(4)}(\mathbf{G}_{\mathbf{m}_N}, \mathbf{G}_{\mathbf{m}_1})\right]_{v_N v_1} = \delta_{s'0}\,\delta_{p'0} \times \delta^{3\mathrm{D}}_{\mathbf{m}_1 0}\;\widetilde{\Phi}_{00}(0, \mathbf{G}_{\mathbf{m}_N}, 0) \times 8\pi^2\sqrt{2s{+}1}\; i^{s+p} j_s(b\,G_{\mathbf{m}_N})\,\overline{P}^{\,s}_{p,0}(\cos\beta_{\mathbf{G}_{\mathbf{m}_N}}) \tag{G58}$$

The above formula concludes the derivation of the elements of the DNA transfer-matrix, $\mathbf{L}$, and the boundary condition vector, $\mathbf{Y}$, which can now be constructed by using Eq. (D11)-Eq. (D13) and Eq. (D17) together with Eq. (G41), Eq. (G51)-Eq. (G52) and Eq. (G56), where wave-vectors $\boldsymbol{\nu}_j$ ($j = 1, ..., N$) should be replaced by the corresponding $\mathbf{G}_{\mathbf{m}_j}$ vectors. Yet, as will be shown in the next Appendix section, it is possible to further simplify the DNA transfer-matrix and the boundary condition vector by taking into account the axial symmetry of the system, greatly reducing the amount of computational resources required for calculation of the DNA partition function.

#### Appendix H: Reduction of the transfer-matrix size based on the axial symmetry of the system

While the above formulas for the blocks of the DNA transfer-matrix and the boundary condition vector provide a complete theoretical basis necessary for study of molecular processes responsible for the nucleus size regulation and chromatin organization, it should be noted that from a practical point of view, these formulas need to be further streamlined to make it possible to use them in real computations. Indeed, it is not hard to check that a typical transfer-matrix constructed by using Eq. (D12) will be usually so large that no computer will be able to handle the DNA partition function calculations. For example, simple estimations show that in the case of $s_{\max} = 7$ and $l_{\max} \approx 2500$, each of the transfer-matrix blocks in Eq. (D12) contains $\sim$ 26 billion complex elements, which is equivalent to staggering $\sim$ 420 Gb of computer memory needed to store information about a single matrix-block. It is clear that in such a situation it would be very hard to calculate the DNA partition function in a short time. However, by using the axial symmetry of the system, it is possible to solve this problem by greatly reducing the size of the DNA transfer-matrix and the boundary condition vector.

To this aim, the first thing we need to do is to note that from Eq. (G24), Eq. (G28), Eq. (G31), Eq. (G34) and Eq. (G37) it follows that the azimuthal angle, $\alpha_{\nu_j}$, of the wave-vector $\boldsymbol{\nu}_j$ enters the DNA transfer-matrix, $\mathbf{L}$, and the boundary condition vector, $\mathbf{Y}$, via a single term: $\overline{D}^{\,u}_{p-p',0}(\mathbf{A}_{\nu_j})$. Indeed, by taking into account that $\mathbf{A}_{\nu_j} = \mathbf{A}_{\nu_j}(\frac{\pi}{2} + \alpha_{\nu_j}, \beta_{\nu_j}, 0)$, from Eq. (F1) we have:

$$\overline{D}^{\,u}_{p-p',0}\big(\mathbf{A}_{\nu_j}\big) = i^{p-p'} e^{i(p-p')\alpha_{\nu_j}}\,\overline{P}^{\,u}_{p-p',0}\big(\cos\beta_{\nu_j}\big) \tag{H1}$$

Where $p = p_j$ and $p' = p_{j+1}$.

To proceed further, let's introduce modified expansion coefficients for the DNA transfer-functions and the boundary condition function, $\big[\widetilde{T}^{(2)}_{k_j k_{j+1}}(\boldsymbol{\nu}_j)\big]^{p_{j+1},\,q_{j+1},\,s_{j+1}}_{p_j,\,q_j,\,s_j}$ and $\big[\widetilde{\xi}^{(2)}_0(\boldsymbol{\nu}_N)\big]^{p_1,\,q_1,\,s_1}_{p_N,\,q_N,\,s_N}$, defined as:

$$\begin{aligned} \big[T^{(2)}_{k_j k_{j+1}}(\boldsymbol{\nu}_j)\big]^{p_{j+1},\,q_{j+1},\,s_{j+1}}_{p_j,\,q_j,\,s_j} &= e^{i(p_j - p_{j+1})\alpha_{\nu_j}}\,\big[\widetilde{T}^{(2)}_{k_j k_{j+1}}(\boldsymbol{\nu}_j)\big]^{p_{j+1},\,q_{j+1},\,s_{j+1}}_{p_j,\,q_j,\,s_j} \\ \big[\xi^{(2)}_0(\boldsymbol{\nu}_N)\big]^{p_1,\,q_1,\,s_1}_{p_N,\,q_N,\,s_N} &= e^{ip_N\alpha_{\nu_N}}\,\big[\widetilde{\xi}^{(2)}_0(\boldsymbol{\nu}_N)\big]^{p_1,\,q_1,\,s_1}_{p_N,\,q_N,\,s_N} \end{aligned} \tag{H2}$$

Then it is easy to see from Eq. (G25), Eq. (G29), Eq. (G32), Eq. (G35) and Eq. (G38) that these modified expansion coefficients are independent from the azimuthal angle, $\alpha_{\nu_j}$.

Substituting Eq. (H2) together with Eq. (C14) and Eq. (C43) into the formula for the DNA partition function [i.e., into Eq. (C32) generalized to an arbitrary value of $K$], it can be shown that the latter turns into:

$$\begin{aligned} Z_{\psi,f} = (2\pi)^3 \int_{\mathrm{IR}^{3N}} \delta(\boldsymbol{\nu}_1)\,\mathrm{d}\boldsymbol{\nu}_1 \ldots \mathrm{d}\boldsymbol{\nu}_N \sum_{\{k_j,p_j,q_j,s_j\}_{j=1}^N} \Big\{ \delta_{k_1 0}\delta_{k_N 0} \times e^{i(p_1-p_2)\alpha_{\nu_1}}\,\big[\widetilde{T}^{(2)}_{k_1 k_2}(\boldsymbol{\nu}_1)\big]^{p_2,q_2,s_2}_{p_1,q_1,s_1}\,\Phi_{k_1 k_2}(\boldsymbol{\nu}_1 - \boldsymbol{\nu}_2)\,\times \\ \times\, e^{i(p_2-p_3)\alpha_{\nu_2}}\,\big[\widetilde{T}^{(2)}_{k_2 k_3}(\boldsymbol{\nu}_2)\big]^{p_3,q_3,s_3}_{p_2,q_2,s_2}\,\Phi_{k_2 k_3}(\boldsymbol{\nu}_2 - \boldsymbol{\nu}_3) \times \ldots \times e^{ip_N\alpha_{\nu_N}}\,\big[\widetilde{\xi}^{(2)}_0(\boldsymbol{\nu}_N)\big]^{p_1,\,q_1,\,s_1}_{p_N,\,q_N,\,s_N}\,\Phi_{00}(\boldsymbol{\nu}_N)\Big\} \end{aligned} \tag{H3}$$

By slightly rearranging the exponential terms in Eq. (H3), it is possible to rewrite it in the following form:

$$\begin{aligned} Z_{\psi,f} = (2\pi)^3 \int_{\mathrm{IR}^{3N}} \delta(\boldsymbol{\nu}_1)\,\mathrm{d}\boldsymbol{\nu}_1 \ldots \mathrm{d}\boldsymbol{\nu}_N \sum_{\{k_j,p_j,q_j,s_j\}_{j=1}^N} \Big\{ \delta_{k_1 0}\delta_{k_N 0} \times e^{ip_1\alpha_{\nu_1}}\,\big[\widetilde{T}^{(2)}_{k_1 k_2}(\boldsymbol{\nu}_1)\big]^{p_2,q_2,s_2}_{p_1,q_1,s_1}\,\Phi_{k_1 k_2}(\boldsymbol{\nu}_1 - \boldsymbol{\nu}_2)\,e^{-ip_2(\alpha_{\nu_1}-\alpha_{\nu_2})} \times \\ \times\, \big[\widetilde{T}^{(2)}_{k_2 k_3}(\boldsymbol{\nu}_2)\big]^{p_3,q_3,s_3}_{p_2,q_2,s_2}\,\Phi_{k_2 k_3}(\boldsymbol{\nu}_2 - \boldsymbol{\nu}_3)\,e^{-ip_3(\alpha_{\nu_2}-\alpha_{\nu_3})} \times \ldots \times \big[\widetilde{\xi}^{(2)}_0(\boldsymbol{\nu}_N)\big]^{p_1,\,q_1,\,s_1}_{p_N,\,q_N,\,s_N}\,\Phi_{00}(\boldsymbol{\nu}_N)\Big\} \end{aligned} \tag{H4}$$

In addition, due to the presence of $\delta_{p'0}$ factor in the expression for the elements of the boundary condition matrix [Eq. (G38)], it follows that the index $p_1$ will be nullified in Eq. (H4). As a result, it is safe to remove $e^{ip_1\alpha_{\nu_1}}$ multiplier from Eq. (H4) as it does not have any effect on the value of the DNA partition function:

$$\begin{aligned} Z_{\psi,f} = (2\pi)^3 \int_{\mathrm{IR}^{3N}} \delta(\boldsymbol{\nu}_1)\,\mathrm{d}\boldsymbol{\nu}_1 \ldots \mathrm{d}\boldsymbol{\nu}_N \sum_{\{k_j,p_j,q_j,s_j\}_{j=1}^N} \Big\{ \delta_{k_1 0}\delta_{k_N 0} \times \big[\widetilde{T}^{(2)}_{k_1 k_2}(\boldsymbol{\nu}_1)\big]^{p_2,q_2,s_2}_{p_1,q_1,s_1}\,\Phi_{k_1 k_2}(\boldsymbol{\nu}_1 - \boldsymbol{\nu}_2)\,e^{-ip_2(\alpha_{\nu_1}-\alpha_{\nu_2})} \times \\ \times\, \big[\widetilde{T}^{(2)}_{k_2 k_3}(\boldsymbol{\nu}_2)\big]^{p_3,q_3,s_3}_{p_2,q_2,s_2}\,\Phi_{k_2 k_3}(\boldsymbol{\nu}_2 - \boldsymbol{\nu}_3)\,e^{-ip_3(\alpha_{\nu_2}-\alpha_{\nu_3})} \times \ldots \times \big[\widetilde{\xi}^{(2)}_0(\boldsymbol{\nu}_N)\big]^{p_1,\,q_1,\,s_1}_{p_N,\,q_N,\,s_N}\,\Phi_{00}(\boldsymbol{\nu}_N)\Big\} \end{aligned} \tag{H5}$$

One can further simplify Eq. (H5) by noting that the imaginary parts of $e^{-ip_{j+1}(\alpha_{\nu_j}-\alpha_{\nu_{j+1}})}$ exponents ($j = 1, \ldots, N$) cancel out during the integration over the corresponding wave-vectors, leaving only the real parts, $\cos(p_{j+1}[\alpha_{\nu_j} - \alpha_{\nu_{j+1}}])$. To see this, let's denote the integrand sum in Eq. (H5) as $\sum(\boldsymbol{\nu}_1, \ldots, \boldsymbol{\nu}_N)$. With this notation, Eq. (H5) transforms into:

$$Z_{\psi,f} = (2\pi)^3 \int_{\mathrm{IR}^{3N}} \delta(\boldsymbol{\nu}_1) \sum(\boldsymbol{\nu}_1, \ldots, \boldsymbol{\nu}_N)\,\mathrm{d}\boldsymbol{\nu}_1 \ldots \mathrm{d}\boldsymbol{\nu}_N \tag{H6}$$

Then, we can split all of the wave-vectors, $(\boldsymbol{\nu}_1, \ldots, \boldsymbol{\nu}_N)$, into two parts: $(\boldsymbol{\nu}_1, \ldots, \boldsymbol{\nu}_j)$ and $(\boldsymbol{\nu}_{j+1}, \ldots, \boldsymbol{\nu}_N)$. While keeping the first part intact, we can rotate wave-vectors from the second part over the same angle, $\Delta\alpha$, in counter-clockwise direction about $\mathbf{z}_0$-axis of the global coordinate system: $(\boldsymbol{\nu}_{j+1}, \ldots, \boldsymbol{\nu}_N) \to (\mathbf{R}_{\Delta\alpha}\boldsymbol{\nu}_{j+1}, \ldots, \mathbf{R}_{\Delta\alpha}\boldsymbol{\nu}_N)$, where $\mathbf{R}_{\Delta\alpha} = \mathbf{R}_{\Delta\alpha}(\Delta\alpha, 0, 0)$ is the rotation matrix corresponding to the angle $\Delta\alpha$. It is easy to check that the determinant of $\frac{\partial(\boldsymbol{\nu}_1,\ldots,\boldsymbol{\nu}_N)}{\partial(\boldsymbol{\nu}_1,\ldots,\boldsymbol{\nu}_j,\mathbf{R}_{\Delta\alpha}\boldsymbol{\nu}_{j+1},\ldots,\mathbf{R}_{\Delta\alpha}\boldsymbol{\nu}_N)}$ Jacobian matrix equals to unity: $\left|\frac{\partial(\boldsymbol{\nu}_1,\ldots,\boldsymbol{\nu}_N)}{\partial(\boldsymbol{\nu}_1,\ldots,\boldsymbol{\nu}_j,\mathbf{R}_{\Delta\alpha}\boldsymbol{\nu}_{j+1},\ldots,\mathbf{R}_{\Delta\alpha}\boldsymbol{\nu}_N)}\right| = 1$. Thus, it can be concluded that Eq. (H6) is equivalent to the following one:

$$Z_{\psi,f} = (2\pi)^3 \int_{\mathrm{IR}^{3N}} \delta(\boldsymbol{\nu}_1) \sum(\boldsymbol{\nu}_1, \ldots, \boldsymbol{\nu}_j, \mathbf{R}_{\Delta\alpha}\boldsymbol{\nu}_{j+1}, \ldots, \mathbf{R}_{\Delta\alpha}\boldsymbol{\nu}_N)\,\mathrm{d}\boldsymbol{\nu}_1 \ldots \mathrm{d}\boldsymbol{\nu}_N \tag{H7}$$

Where $\Delta\alpha \in [0, 2\pi]$ is an arbitrary angle.

Since Eq. (H7) holds for any $\Delta\alpha$, it is clear that we can perform $\frac{1}{2\pi}\int_0^{2\pi} \mathrm{d}\Delta\alpha$ integration of the right side of the above expression over all possible values of $\Delta\alpha$, without affecting its left side:

$$Z_{\psi,f} = \frac{1}{2\pi}\,(2\pi)^3 \int_0^{2\pi} \mathrm{d}\Delta\alpha \int_{\mathrm{IR}^{3N}} \delta(\boldsymbol{\nu}_1) \sum (\boldsymbol{\nu}_1, ..., \boldsymbol{\nu}_j, \mathbf{R}_{\Delta\alpha}\boldsymbol{\nu}_{j+1}, ..., \mathbf{R}_{\Delta\alpha}\boldsymbol{\nu}_N)\; \mathrm{d}\boldsymbol{\nu}_1...\mathrm{d}\boldsymbol{\nu}_N \tag{H8}$$

From the first look at Eq. (H6) and Eq. (H8), it seems that we have arrived to the formula for the DNA partition function, which is more complex than the original one. However, as we will see in a moment, this is a deceptive feeling. Specifically, let's consider any set of wave-vectors, $(\boldsymbol{\nu}_1, ..., \boldsymbol{\nu}_N)$, under the integral sign in Eq. (H8) and put $\Delta\alpha = 2(\alpha_{\nu_j} - \alpha_{\nu_{j+1}})$. Obviously, rotation by the angle $\Delta\alpha$ does not have any effect on $\big[\widetilde{T}^{(2)}_{k_n k_{n+1}}(\boldsymbol{\nu}_n)\big]^{p_{n+1},\, q_{n+1},\, s_{n+1}}_{p_n,\, q_n,\, s_n}$ and $\big[\widetilde{\xi}^{(2)}_0(\boldsymbol{\nu}_N)\big]^{p_1,\, q_1,\, s_1}_{p_N,\, q_N,\, s_N}$ expansion coefficients $(n = 1, ..., N-1)$, which are independent from the azimuthal angles of the wave-vectors. $\Phi_{k_n k_{n+1}}(\boldsymbol{\nu}_n - \boldsymbol{\nu}_{n+1})$ functions, where $n = 1, ..., j-1, j+1, ..., N-1$, also do not change their values as $(\boldsymbol{\nu}_n, \boldsymbol{\nu}_{n+1})$ pairs of wave-vectors either stay intact or rotate together by the same angle, $\Delta\alpha$. As for $\Phi_{k_j k_{j+1}}(\boldsymbol{\nu}_j - \boldsymbol{\nu}_{j+1})$ function, it keeps its value due to a different reason – based on Eq. (C9), it can be shown that the value of this function stay unchanged upon rotation by $\Delta\alpha = 2(\alpha_{\nu_j} - \alpha_{\nu_{j+1}})$ angle due to the axial symmetry of the system. Similarly, all of the exponents, $e^{-ip_{n+1}(\alpha_{\nu_n} - \alpha_{\nu_{n+1}})}$, stay unchanged except for the one: $e^{-ip_{j+1}(\alpha_{\nu_j} - \alpha_{\nu_{j+1}})}$. As for the latter, upon rotation, its imaginary part simply reverses the sign, while the real part remains untouched. Thus, for any set of the wave-vectors under the integral sign of Eq. (H8) there always exists a dual one, which can be obtain by rotation about $\mathbf{z}_0$-axis of the global coordinate system, that reverses the sign of the imaginary part of $e^{-ip_{j+1}(\alpha_{\nu_j} - \alpha_{\nu_{j+1}})}$ exponent without affecting other terms.

It is then clear that the imaginary part of $e^{-ip_{j+1}(\alpha_{\nu_j} - \alpha_{\mathbf{R}_{\Delta\alpha}\nu_{j+1}})}$ exponent in Eq. (H8) will be cancelled out due to $\frac{1}{2\pi}\int_0^{2\pi} \mathrm{d}\Delta\alpha$ integration, without making any contribution to the DNA partition function. In other words, we need to keep track only of the real part of $e^{-ip_{j+1}(\alpha_{\nu_j} - \alpha_{\mathbf{R}_{\Delta\alpha}\nu_{j+1}})}$ term in Eq. (H8), while its imaginary part can be safely omitted:

$$Z_{\psi,f} = \frac{1}{2\pi}\,(2\pi)^3 \int_0^{2\pi} \mathrm{d}\Delta\alpha \int_{\mathrm{IR}^{3N}} \delta(\boldsymbol{\nu}_1)\; \mathrm{Re}_j\Big[\sum (\boldsymbol{\nu}_1, ..., \boldsymbol{\nu}_j, \mathbf{R}_{\Delta\alpha}\boldsymbol{\nu}_{j+1}, ..., \mathbf{R}_{\Delta\alpha}\boldsymbol{\nu}_N)\Big]\; \mathrm{d}\boldsymbol{\nu}_1...\mathrm{d}\boldsymbol{\nu}_N \tag{H9}$$

Where $\mathrm{Re}_j\big[\sum(\boldsymbol{\nu}_1, ..., \boldsymbol{\nu}_j, \mathbf{R}_{\Delta\alpha}\boldsymbol{\nu}_{j+1}, ..., \mathbf{R}_{\Delta\alpha}\boldsymbol{\nu}_N)\big]$ simply means the sum $\sum(\boldsymbol{\nu}_1, ..., \boldsymbol{\nu}_j, \mathbf{R}_{\Delta\alpha}\boldsymbol{\nu}_{j+1}, ..., \mathbf{R}_{\Delta\alpha}\boldsymbol{\nu}_N)$ in which $e^{-ip_{j+1}(\alpha_{\nu_j} - \alpha_{\mathbf{R}_{\Delta\alpha}\nu_{j+1}})}$ function is replaced by its real part, $\cos(p_{j+1}[\alpha_{\nu_j} - \alpha_{\mathbf{R}_{\Delta\alpha}\nu_{j+1}}])$.

Similarly to Eq. (H6) and Eq. (H7), it is not hard to show that the value of the multiple integral, $\int_{\mathrm{IR}^{3N}} \mathrm{d}\boldsymbol{\nu}_1...\mathrm{d}\boldsymbol{\nu}_N$, in the above formula is independent from the value of the rotation angle, $\Delta\alpha$. Thus, after performing $\frac{1}{2\pi}\int_0^{2\pi} \mathrm{d}\Delta\alpha$ integration, we arrive to the following formula:

$$Z_{\psi,f} = (2\pi)^3 \int_{\mathrm{IR}^{3N}} \delta(\boldsymbol{\nu}_1)\; \mathrm{Re}_j\Big[\sum (\boldsymbol{\nu}_1, ..., \boldsymbol{\nu}_N)\Big]\; \mathrm{d}\boldsymbol{\nu}_1...\mathrm{d}\boldsymbol{\nu}_N, \tag{H10}$$

in which only the real part of $e^{-ip_{j+1}(\alpha_{\nu_j} - \alpha_{\mathbf{R}_{\Delta\alpha}\nu_{j+1}})}$ exponent is remained.

Since the above argument can be applied to any $j$ from 1 to $N$, it then easy to see that Eq. (H5) can be reduced to:

$$\begin{aligned} Z_{\psi,f} = (2\pi)^3 \int_{\mathrm{IR}^{3N}} \delta(\boldsymbol{\nu}_1)\, \mathrm{d}\boldsymbol{\nu}_1...\mathrm{d}\boldsymbol{\nu}_N \sum_{\{k_j, p_j, q_j, s_j\}_{j=1}^N} \Big\{ & \delta_{k_1 0}\delta_{k_N 0} \times \big[\widetilde{T}^{(2)}_{k_1 k_2}(\boldsymbol{\nu}_1)\big]^{p_2, q_2, s_2}_{p_1, q_1, s_1} \Phi_{k_1 k_2}(\boldsymbol{\nu}_1 - \boldsymbol{\nu}_2) \cos(p_2[\alpha_{\nu_1} - \alpha_{\nu_2}]) \times \\ & \times \big[\widetilde{T}^{(2)}_{k_2 k_3}(\boldsymbol{\nu}_2)\big]^{p_3, q_3, s_3}_{p_2, q_2, s_2} \Phi_{k_2 k_3}(\boldsymbol{\nu}_2 - \boldsymbol{\nu}_3) \cos(p_3[\alpha_{\nu_2} - \alpha_{\nu_3}]) \times \,...\, \times \big[\widetilde{\xi}^{(2)}_0(\boldsymbol{\nu}_N)\big]^{p_1,\, q_1,\, s_1}_{p_N,\, q_N,\, s_N} \Phi_{00}(\boldsymbol{\nu}_N) \Big\} \end{aligned} \tag{H11}$$

As Eq. (D9) has a very similar look to Eq. (C42), which is derived from Eq. (C32), it is natural to expect that the above approach can be also used to simplify Eq. (D9). Indeed, by substituting Eq. (C14), Eq. (C34), Eq. (C37), Eq. (C43) and Eq. (H2) into Eq. (D9), where all $\boldsymbol{\nu}_j$ wave-vectors are replaced by their discrete counterparts, $\mathbf{G}_{\mathbf{m}_j}$, one can obtain the next formula for the DNA partition function:

$$Z_{\psi,f} = V_{\rm B} \times \sum_{\{\mathbf{m}_j,k_j,p_j,q_j,s_j\}_{j=1}^N} \Big\{ \delta^{\rm 3D}_{\mathbf{m}_1 0}\delta_{k_1 0}\delta_{k_N 0} \times \big[\widetilde{T}^{(2)}_{k_1 k_2}(\mathbf{G}_{\mathbf{m}_1})\big]^{p_2,q_2,s_2}_{p_1,q_1,s_1} \Phi_{k_1 k_2}(\mathbf{G}_{\mathbf{m}_1} - \mathbf{G}_{\mathbf{m}_2})\, e^{-ip_2(\alpha_{\mathbf{G}_{\mathbf{m}_1}} - \alpha_{\mathbf{G}_{\mathbf{m}_2}})} \times$$
$$\times \big[\widetilde{T}^{(2)}_{k_2 k_3}(\mathbf{G}_{\mathbf{m}_2})\big]^{p_3,q_3,s_3}_{p_2,q_2,s_2} \Phi_{k_2 k_3}(\mathbf{G}_{\mathbf{m}_2} - \mathbf{G}_{\mathbf{m}_3})\, e^{-ip_3(\alpha_{\mathbf{G}_{\mathbf{m}_2}} - \alpha_{\mathbf{G}_{\mathbf{m}_3}})} \times \ldots \times \big[\widetilde{\xi}^{(2)}_0(\mathbf{G}_{\mathbf{m}_N})\big]^{p_1,\,q_1,\,s_1}_{p_N,\,q_N,\,s_N} \Phi_{00}(\mathbf{G}_{\mathbf{m}_N}) \Big\}, \quad \text{(H12)}$$

which closely resembles Eq. (H5).

Based on the analogy with Eq. (H5), it is then clear that all of the imaginary parts of the exponents present in Eq. (H12) have to cancel out somehow, resulting in the following expression:

$$Z_{\psi,f} = V_{\rm B} \times \sum_{\{\mathbf{m}_j,k_j,p_j,q_j,s_j\}_{j=1}^N} \Big\{ \delta^{\rm 3D}_{\mathbf{m}_1 0}\delta_{k_1 0}\delta_{k_N 0} \times \big[\widetilde{T}^{(2)}_{k_1 k_2}(\mathbf{G}_{\mathbf{m}_1})\big]^{p_2,q_2,s_2}_{p_1,q_1,s_1} \Phi_{k_1 k_2}(\mathbf{G}_{\mathbf{m}_1} - \mathbf{G}_{\mathbf{m}_2}) \cos(p_2[\alpha_{\mathbf{G}_{\mathbf{m}_1}} - \alpha_{\mathbf{G}_{\mathbf{m}_2}}]) \times$$
$$\times \big[\widetilde{T}^{(2)}_{k_2 k_3}(\mathbf{G}_{\mathbf{m}_2})\big]^{p_3,q_3,s_3}_{p_2,q_2,s_2} \Phi_{k_2 k_3}(\mathbf{G}_{\mathbf{m}_2} - \mathbf{G}_{\mathbf{m}_3}) \cos(p_3[\alpha_{\mathbf{G}_{\mathbf{m}_2}} - \alpha_{\mathbf{G}_{\mathbf{m}_3}}]) \times \ldots \times \big[\widetilde{\xi}^{(2)}_0(\mathbf{G}_{\mathbf{m}_N})\big]^{p_1,\,q_1,\,s_1}_{p_N,\,q_N,\,s_N} \Phi_{00}(\mathbf{G}_{\mathbf{m}_N}) \Big\} \quad \text{(H13)}$$

Yet, it is not hard to see that the aforementioned proof based on the rotation of wave-vectors cannot be directly applied to the Eq. (H12) in its original form due to the absence of the corresponding nodes in the reciprocal lattice, see, for example, Figure S11(a). This is exactly why we had to go through all of the trouble to derive Eq. (H5) and Eq. (H11) in the first place since by using them one can prove equivalence of Eq. (H12) and Eq. (H13).

Indeed, from the formulas derived in this section as well as in Appendices C-D it follows that Eq. (H5), Eq. (H11) and Eq. (H12) provide the same value of the DNA partition function. In turn, by utilizing the Dirac comb function, it can be shown that Eq. (H11) is equivalent to its discrete version, Eq. (H13). Thus, even though the wave-vector rotation argument cannot be directly applied to Eq. (H12), it is possible to prove correctness of Eq. (H13) by using a detour through Eq. (H5) and Eq. (H11). Numeric calculations confirm this result by showing that Eq. (H13) indeed outputs the same value of the DNA partition function as Eq. (H12).

The main reason why we are interested in Eq. (H13) is because from this equation it follows that in the case of an axially-symmetric nucleus, one does not have to use all of the wave-vectors, $\mathbf{G}_{\mathbf{m}_j}$, in the construction of the transfer-matrix, $\mathbf{L}$, and the boundary condition vector, $\mathbf{Y}$. Instead only a small subset of wave-vectors corresponding to the reciprocal lattice nodes highlighted in red color in Figure S11(b), which we will refer to as the unique-value nodes, suffice for this purpose.

To see this, we just need to split $\sum_{\{\mathbf{m}_j\}_{j=1}^N}$ sum in Eq. (H13) into the following parts: $\sum_{\{\mathbf{m}_j\}_{j=1}^N} = \sum_{j=1}^N \sum_{\mathbf{m}_j} = \sum_{j=1}^N \sum'_{\mathbf{m}_j} \sum_{{\rm SR}(\mathbf{m}_j)}$, where $\sum'_{\mathbf{m}_j}$ sum is calculated over the unique-value nodes depicted in Figure S11(b), and $\sum_{{\rm SR}(\mathbf{m}_j)}$ sum is performed over all possible rotations of a given vector, $\mathbf{m}_j$, about $\mathbf{z}_0$-axis of the global coordinate system as schematically shown in Figure S11(c, d).

Then, since $\big[\widetilde{T}^{(2)}_{k_j k_{j+1}}(\boldsymbol{\nu}_j)\big]^{p_{j+1},\,q_{j+1},\,s_{j+1}}_{p_j,\,q_j,\,s_j}$ and $\big[\widetilde{\xi}^{(2)}_0(\boldsymbol{\nu}_N)\big]^{p_1,\,q_1,\,s_1}_{p_N,\,q_N,\,s_N}$ expansion coefficients are independent from the azimuthal angles of $\mathbf{G}_{\mathbf{m}_j}$ wave-vectors, it is clear that after performing the above splitting, these coefficients can be moved outside of the right-most sum, $\sum_{{\rm SR}(\mathbf{m}_j)}$. By doing so, Eq. (H13) turns into:

$$Z_{\psi,f} = V_{\rm B} \times \sum'_{\{\mathbf{m}_j,k_j,p_j,q_j,s_j\}_{j=1}^N} \Big\{ \delta^{\rm 3D}_{\mathbf{m}_1 0}\delta_{k_1 0}\delta_{k_N 0} \times \big[\widetilde{T}^{(2)}_{k_1 k_2}(\mathbf{G}_{\mathbf{m}_1})\big]^{p_2,q_2,s_2}_{p_1,q_1,s_1} \widetilde{\Phi}_{k_1 k_2}(p_2, \mathbf{G}_{\mathbf{m}_1}, \mathbf{G}_{\mathbf{m}_2}) \times$$
$$\times \big[\widetilde{T}^{(2)}_{k_2 k_3}(\mathbf{G}_{\mathbf{m}_2})\big]^{p_3,q_3,s_3}_{p_2,q_2,s_2} \widetilde{\Phi}_{k_2 k_3}(p_3, \mathbf{G}_{\mathbf{m}_2}, \mathbf{G}_{\mathbf{m}_3}) \times \ldots \times \big[\widetilde{\xi}^{(2)}_0(\mathbf{G}_{\mathbf{m}_N})\big]^{p_1,\,q_1,\,s_1}_{p_N,\,q_N,\,s_N} \widetilde{\Phi}_{00}(0, \mathbf{G}_{\mathbf{m}_N}, 0) \Big\} \quad \text{(H14)}$$

Where the prime indicates that $\sum'_{\{\mathbf{m}_j\}_{j=1}^N}$ summation is performed only over the unique-value nodes. As for the modified $\widetilde{\Phi}_{k_j k_{j+1}}(p_{j+1}, \mathbf{G}_{\mathbf{m}_j}, \mathbf{G}_{\mathbf{m}_{j+1}})$ functions in Eq. (H14), they are defined as:

$$\widetilde{\Phi}_{k_j k_{j+1}}(p_{j+1}, \mathbf{G}_{\mathbf{m}_j}, \mathbf{G}_{\mathbf{m}_{j+1}}) = \sum_{{\rm SR}(\mathbf{m}_{j+1})} \Phi_{k_j k_{j+1}}(\mathbf{G}_{\mathbf{m}_j} - \mathbf{G}_{\mathbf{m}_{j+1}}) \cos(p_{j+1}[\alpha_{\mathbf{G}_{\mathbf{m}_j}} - \alpha_{\mathbf{G}_{\mathbf{m}_{j+1}}}]) \quad \text{(H15)}$$

Here, as before, summation $\sum_{{\rm SR}(\mathbf{m}_{j+1})}$ is carried out over all possible rotations of the vector $\mathbf{m}_{j+1}$ about $\mathbf{z}_0$-axis of the global coordinate system as schematically shown in Figure S11(c, d).

By following the same derivation as in Appendices C-D, it is not hard to see that Eq. (H14) leads to the following formula for the DNA partition function, which is similar to Eq. (D16):

$$Z_{\psi,f} = V_{\mathrm{B}}\, \widetilde{\mathbf{U}}\widetilde{\mathbf{L}}^{N-1}\widetilde{\mathbf{Y}} \tag{H16}$$

Where this time matrices $\widetilde{\mathbf{U}}$, $\widetilde{\mathbf{L}}$ and $\widetilde{\mathbf{Y}}$ include only the blocks corresponding to the unique-value nodes of the reciprocal lattice. Specifically, for these matrices we have:

$$\widetilde{\mathbf{U}} = (0\ \cdots\ 0\ \ 1\ \ 0\ \cdots\ 0) \quad \text{and} \quad \widetilde{\mathbf{L}} = \begin{pmatrix} \widetilde{\mathbf{T}}^{(6)}_{00} & \widetilde{\mathbf{T}}^{(6)}_{01} & 0 & \cdots & 0 \\ 0 & 0 & \mathbf{I} & \cdots & 0 \\ \vdots & \vdots & \vdots & \ddots & \vdots \\ 0 & 0 & 0 & \cdots & \mathbf{I} \\ \mathbf{I} & 0 & 0 & \cdots & 0 \end{pmatrix} \quad \text{and} \quad \widetilde{\mathbf{Y}} = \begin{pmatrix} \widetilde{\boldsymbol{\xi}}^{(5)}_{0}\big|^{\widetilde{n}_0}_{\text{column}} \\ 0 \\ 0 \\ \vdots \\ 0 \end{pmatrix} \tag{H17}$$

Where $\widetilde{\mathbf{T}}^{(6)}_{nm}$ and $\widetilde{\boldsymbol{\xi}}^{(5)}_{0}$ matrix-blocks are constructed in the same way as the original blocks, $\mathbf{T}^{(6)}_{nm}$ and $\boldsymbol{\xi}^{(5)}_{0}$, except that the former have a much smaller size of $\widetilde{l}_{\max} \times \widetilde{l}_{\max}$ in comparison to the latter, whose size is $l_{\max} \times l_{\max}$ [see Eq. (D11) and Eq. (D13)]:

$$\widetilde{\mathbf{T}}^{(6)}_{nm} = \begin{pmatrix} \widetilde{\mathbf{T}}^{(5)}_{nm}(1,1) & \cdots & \cdots & \widetilde{\mathbf{T}}^{(5)}_{nm}(1,\widetilde{l}_{\max}) \\ \vdots & \ddots & & \vdots \\ \vdots & & \ddots & \vdots \\ \widetilde{\mathbf{T}}^{(5)}_{nm}(\widetilde{l}_{\max},1) & \cdots & \cdots & \widetilde{\mathbf{T}}^{(5)}_{nm}(\widetilde{l}_{\max},\widetilde{l}_{\max}) \end{pmatrix} \quad \text{and} \quad \widetilde{\boldsymbol{\xi}}^{(5)}_{0} = \begin{pmatrix} \widetilde{\boldsymbol{\xi}}^{(4)}_{0}(1,1) & \cdots & \cdots & \widetilde{\boldsymbol{\xi}}^{(4)}_{0}(1,\widetilde{l}_{\max}) \\ \vdots & \ddots & & \vdots \\ \vdots & & \ddots & \vdots \\ \widetilde{\boldsymbol{\xi}}^{(4)}_{0}(\widetilde{l}_{\max},1) & \cdots & \cdots & \widetilde{\boldsymbol{\xi}}^{(4)}_{0}(\widetilde{l}_{\max},\widetilde{l}_{\max}) \end{pmatrix} \tag{H18}$$

Here $\widetilde{l}_{\max}$ is the total number of the unique-value nodes of the reciprocal lattice used in the DNA partition function calculations.

Indeed, simple estimations indicate that in contrast to the original matrix-blocks, $\mathbf{T}^{(6)}_{00}$ and $\mathbf{T}^{(6)}_{01}$, each of which occupies $\sim$ 420 Gb of the computer memory in the case of $s_{\max} = 7$ and $l_{\max} \approx 2500$, their size-reduced counterparts defined by Eq. (H18) take up $\sim$ 6 Gb of the computer memory each because of much smaller value of $\widetilde{l}_{\max}$ ($\widetilde{l}_{\max} \approx 300$ when $l_{\max} \approx 2500$). In fact, it can be shown that the amount of the computer memory needed to calculate the DNA partition function can be further decreased by $\sim$ 2 times by using $\big[\widetilde{T}^{(2)}_{k_j k_{j+1}}(\nu_x,\nu_y,\nu_z)\big]^{p',\,q',\,s'}_{p,\,q,\,s} = (-1)^{p-p'}\big[\widetilde{T}^{(2)}_{k_j k_{j+1}}(\nu_x,\nu_y,-\nu_z)\big]^{p',\,q',\,s'}_{p,\,q,\,s}$ symmetry of the modified expansion coefficients, which can be derived from Eq. (G25), Eq. (G29), Eq. (G32), Eq. (G35) and Eq. (F3), with a very similar formula holding for $\big[\widetilde{\xi}^{(2)}_{0}(\boldsymbol{\nu})\big]^{p',\,q',\,s'}_{p,\,q,\,s}$ coefficients as well. Hence, the axial symmetry of the cell nucleus assumed in this study as well as the symmetry of $P^{s}_{p,q}$ polynomials make it possible to save a lot of computational resources, rendering calculations of the partition function of DNA to be manageable by modern computer systems.

To complete Eq. (H16)-Eq. (H18), it should be noted that $\widetilde{\mathbf{T}}^{(5)}_{nm}$ and $\widetilde{\boldsymbol{\xi}}^{(4)}_{0}$ matrix-blocks in Eq. (H18) are defined in the same way as their original counterparts [see comments after Eq. (D10)]: $\widetilde{\mathbf{T}}^{(5)}_{nm}(l_j,l_{j+1}) = \widetilde{\mathbf{T}}^{(5)}_{nm}(\mathbf{G}_{\mathbf{m}_j(l_j)},\mathbf{G}_{\mathbf{m}_{j+1}(l_{j+1})})$ and $\widetilde{\boldsymbol{\xi}}^{(4)}_{0}(l_N,l_1) = \widetilde{\boldsymbol{\xi}}^{(4)}_{0}(\mathbf{G}_{\mathbf{m}_N(l_N)},\mathbf{G}_{\mathbf{m}_1(l_1)})$, where the elements of the matrices are given by Eq. (G42), Eq. (G47), Eq. (G53) and Eq. (G57), accordingly. As for $\widetilde{\boldsymbol{\xi}}^{(5)}_{0}\big|^{\widetilde{n}_0}_{\text{column}}$ in Eq. (H17), this is the $\widetilde{n}_0^{\text{th}}$ column of the matrix $\widetilde{\boldsymbol{\xi}}^{(5)}_{0}$, which is the only column of the matrix that contains non-zero elements, see Appendix D for more details. Similarly, the only non-zero element, 1, of the vector $\widetilde{\mathbf{U}}$ in Eq. (H16) is also located in the $\widetilde{n}_0^{\text{th}}$ column.

Having at hand the above formulas, it is then rather straightforward to find a corresponding mathematical expression for the functional derivative of the DNA partition function, $\frac{\delta \ln Z_\psi}{\delta\psi(r)}$, by repeating the derivation from Appendix E. As a result, it can be shown that:

$$\frac{\delta \ln Z_\psi}{\delta\psi(r)} \approx \frac{N_{\text{tot}}}{\lambda_{\max}}\,\frac{\mathrm{Pr}^{\mathrm{L}}_{\lambda_{\max}}\widetilde{\mathbf{U}} \times \frac{\delta}{\delta\psi(r)}\widetilde{\mathbf{L}} \times \mathrm{Pr}^{\mathrm{R}}_{\lambda_{\max}}\widetilde{\mathbf{Y}}}{\mathrm{Pr}^{\mathrm{L}}_{\lambda_{\max}}\widetilde{\mathbf{U}} \times \mathrm{Pr}^{\mathrm{R}}_{\lambda_{\max}}\widetilde{\mathbf{Y}}} \tag{H19}$$

Where $\frac{\delta}{\delta\psi(r)}\widetilde{\mathbf{L}}$ functional derivative of the transfer-matrix $\widetilde{\mathbf{L}}$ equals to:

$$\frac{\delta}{\delta\psi(r)}\widetilde{\mathbf{L}} = \begin{pmatrix} \frac{\delta}{\delta\psi(r)}\widetilde{\mathbf{T}}^{(6)}_{00} & \frac{\delta}{\delta\psi(r)}\widetilde{\mathbf{T}}^{(6)}_{01} & 0 & \cdots & 0 \\ 0 & 0 & 0 & \cdots & 0 \\ \vdots & \vdots & \vdots & \ddots & \vdots \\ 0 & 0 & 0 & \cdots & 0 \\ 0 & 0 & 0 & \cdots & 0 \end{pmatrix} \tag{H20}$$

Here the matrix-blocks $\frac{\delta}{\delta\psi(r)}\widetilde{\mathbf{T}}^{(6)}_{00}$ and $\frac{\delta}{\delta\psi(r)}\widetilde{\mathbf{T}}^{(6)}_{01}$ are constructed in the same way as matrices $\widetilde{\mathbf{T}}^{(6)}_{00}$ and $\widetilde{\mathbf{T}}^{(6)}_{01}$ with the only difference being that the modified functions, $\widetilde{\Phi}_{00}$ and $\widetilde{\Phi}_{K0}$, in Eq. (G42) and Eq. (G53) must be replaced with the respective functional derivatives: $\frac{\delta\widetilde{\Phi}_{00}}{\delta\psi(r)}$ and $\frac{\delta\widetilde{\Phi}_{K0}}{\delta\psi(r)} + \beta\widetilde{\Phi}_{K0}\frac{\delta\mu_{\rm pr}}{\delta\psi(r)}$, where $\frac{\delta\mu_{\rm pr}}{\delta\psi(r)}$ is defined by Eq. (E11); whereas, for $\frac{\delta\widetilde{\Phi}_{00}}{\delta\psi(r)}$ and $\frac{\delta\widetilde{\Phi}_{K0}}{\delta\psi(r)}$ functional derivatives from Eq. (E9) and Eq. (H15) we have:

$$\begin{aligned}
\frac{\delta\widetilde{\Phi}_{00}}{\delta\psi(r)} &= \begin{cases} -\dfrac{4\pi}{V_{\rm B}}\beta r^2 q_{00} e^{-\beta q_{00}\psi(r)} \displaystyle\sum_{\mathrm{SR}(\mathbf{m}_{j+1})} j_0\big(r\,\|\mathbf{G}_{\mathbf{m}_j}-\mathbf{G}_{\mathbf{m}_{j+1}}\|\big)\cos(p_{j+1}[\alpha_{\mathbf{G}_{\mathbf{m}_j}}-\alpha_{\mathbf{G}_{\mathbf{m}_{j+1}}}]), & \text{if } r<R_{\rm nucl} \\ 0 & \text{, otherwise} \end{cases} \\
\frac{\delta\widetilde{\Phi}_{K0}}{\delta\psi(r)} &= \begin{cases} -\dfrac{4\pi}{V_{\rm B}}\beta r^2 q_{K0} e^{-\beta q_{K0}\psi(r)} \displaystyle\sum_{\mathrm{SR}(\mathbf{m}_{j+1})} j_0\big(r\,\|\mathbf{G}_{\mathbf{m}_j}-\mathbf{G}_{\mathbf{m}_{j+1}}\|\big)\cos(p_{j+1}[\alpha_{\mathbf{G}_{\mathbf{m}_j}}-\alpha_{\mathbf{G}_{\mathbf{m}_{j+1}}}]), & \text{if } r<R_{\rm nucl} \\ 0 & \text{, otherwise} \end{cases}
\end{aligned} \tag{H21}$$

Since now we have all the necessary formulas for calculation of the DNA partition function, we can proceed to the description of the computational algorithm in the next Appendix section.

#### Appendix I: Calculation algorithm for the DNA partition function

By using formulas derived in Appendix G and Appendix H, one can find the partition function of DNA via the following three steps:

1) *Calculation of the offset free energy of nucleosomes,* $\mu_{\rm off}$.

   To obtain accurate estimations of the DNA partition function, it is necessary in the first place to know the exact value of the offset free energy of nucleosomes, $\mu_{\rm off}$ (see Appendix B), which, in contrast to other model parameters that can be measured in experiments, has to be determined numerically.

   Specifically, the value of $\mu_{\rm off}$ can be acquired by performing several iterative transfer-matrix calculations, in which the parameter $\mu_{\rm off}$ is varied until the average occupancy fraction of DNA by nucleosomes, $O_{\rm nucl}(\psi, f)$, starts to obey the classical exponential relation, $O_{\rm nucl}(\psi, f) \approx e^{\beta\mu_{\rm pr}}$, when the nucleus radius is set to infinity ($R_{\rm nucl} \to +\infty$) and the stretching force, $f$, and the electrostatic potential, $\psi$, are put equal to zero ($f = 0$ and $\psi = 0$) to imitate typical *in vitro* bulk experimental conditions.

   It should be noted that such iterative calculations must be done at sufficiently large negative values of the protein binding free energy to DNA (i.e., for $\mu_{\rm pr} < -5\ k_{\rm B}T$) to keep the amount of protein-bound DNA segments at a low level, since otherwise behaviour of the average DNA occupancy fraction will strongly deviate from the exponential law.

   To calculate $O_{\rm nucl}(\psi, f)$, one can use the following formula:

$$O_{\rm nucl}(\psi, f) = \frac{K}{\beta N}\frac{\partial \ln Z_{\psi,f}}{\partial \mu_{\rm pr}}, \tag{I1}$$

   which can be easily derived from Eq. (B1), Eq. (B6) and Eq. (C1). Here $K$ is the number of DNA segments constrained inside a single nucleosome, $\mu_{\rm pr}$ is the binding free energy of a histone octamer to DNA and $N$ is the total number of DNA segments in a simulated DNA molecule, see Appendix B for more details.

In order to find $\frac{\partial \ln Z_{\psi,f}}{\partial \mu_{\rm pr}}$ derivative from Eq. (I1), it is possible to apply the standard finite difference method. To this aim, the DNA partition function can be obtained via Eq. (H16), with the easiest way of calculating the matrix product being the one by using the power iteration method described by Eq. (E3)-Eq. (E4). Namely, it can be shown that in the general case:

$$Z_{\psi,f} = V_{\rm B}\,\widetilde{\mathbf{U}}\widetilde{\mathbf{L}}^{N-1}\widetilde{\mathbf{Y}} = C_0\,\lambda_{\rm max}^{N}\,[\mathrm{Pr}^{\rm L}_{\lambda_{\rm max}}\widetilde{\mathbf{U}} \times \mathrm{Pr}^{\rm R}_{\lambda_{\rm max}}\widetilde{\mathbf{Y}}] \tag{I2}$$

Where the prefactor $C_0$, the dominant eigenvalue of the transfer-matrix, $\lambda_{\rm max}$, and the corresponding projections of the vectors $\widetilde{\mathbf{U}}$ and $\widetilde{\mathbf{Y}}$: $\mathrm{Pr}^{\rm L}_{\lambda_{\rm max}}\widetilde{\mathbf{U}}$ and $\mathrm{Pr}^{\rm R}_{\lambda_{\rm max}}\widetilde{\mathbf{Y}}$, – all can be obtained with the help of the power iteration algorithm.

In the above formula, the DNA transfer-matrix, $\widetilde{\mathbf{L}}$, and the boundary condition vectors, $\widetilde{\mathbf{U}}$ and $\widetilde{\mathbf{Y}}$, are constructed with the help of Eq. (H17) and Eq. (H18) by putting $f = 0$, $\psi = 0$ and $R_{\rm nucl} \to +\infty$ in the corresponding formulas for the elements of $\widetilde{\mathbf{T}}^{(5)}_{nm}$ and $\boldsymbol{\xi}^{(4)}_0$ matrix-blocks. From these equations, it is clear that the transfer-matrix $\widetilde{\mathbf{L}}$ is composed of $(K+1)\times(K+1)$ matrix-blocks, $\widetilde{\mathbf{T}}^{(6)}_{nm}$, with each of the blocks containing $\widetilde{l}_{\rm max}(s_{\rm max}+1)^2 \times \widetilde{l}_{\rm max}(s_{\rm max}+1)^2$ elements, where $\widetilde{l}_{\rm max}$ is the total number of the unique-value nodes of the reciprocal lattice used in the DNA partition function calculations [see Figure S11(b)]; and $s_{\rm max}$ is the upper bound on the principal index of the Wigner D-functions used in the expansion series of the DNA transfer-functions [see comments after Eq. (C38) and Table I]. Thus, the total size of the transfer-matrix $\widetilde{\mathbf{L}}$ is $(K+1)\,\widetilde{l}_{\rm max}(s_{\rm max}+1)^2 \times (K+1)\,\widetilde{l}_{\rm max}(s_{\rm max}+1)^2$. Correspondingly, the size of the boundary condition vector $\widetilde{\mathbf{Y}}$ is $(K+1)\,\widetilde{l}_{\rm max}(s_{\rm max}+1)^2 \times 1$ and the size of the vector $\widetilde{\mathbf{U}}$ is $1 \times (K+1)\,\widetilde{l}_{\rm max}(s_{\rm max}+1)^2$.

It should be noted that in order to optimize the use of computational resources, it is possible to work only with $\widetilde{\mathbf{T}}^{(6)}_{00}$ and $\widetilde{\mathbf{T}}^{(6)}_{01}$ matrix-blocks instead of the whole matrix $\widetilde{\mathbf{L}}$ to calculate the DNA partition function.

To see this, let's say that we have a vector, $\mathbf{V}$, composed of $(K+1)\,\widetilde{l}_{\rm max}(s_{\rm max}+1)^2$ elements, and we want to find the result of this vector multiplication by the matrix $\widetilde{\mathbf{L}}$: $\mathbf{V}' = \widetilde{\mathbf{L}}\mathbf{V}$. Since the transfer-matrix $\widetilde{\mathbf{L}}$ comprises $(K+1)\times(K+1)$ matrix blocks, it makes sense to split the both vectors, $\mathbf{V}$ and $\mathbf{V}'$, into the same number of blocks, $\mathbf{V}_n$ and $\mathbf{V}'_n$ $(n = 1, ..., K+1)$:

$$\mathbf{V} = \begin{pmatrix} \mathbf{V}_1 \\ \vdots \\ \mathbf{V}_{K+1} \end{pmatrix} \quad \text{and} \quad \mathbf{V}' = \begin{pmatrix} \mathbf{V}'_1 \\ \vdots \\ \mathbf{V}'_{K+1} \end{pmatrix} \tag{I3}$$

Then from Eq. (H17) it is clear that:

$$\mathbf{V}' = \begin{pmatrix} \mathbf{V}'_1 \\ \mathbf{V}'_2 \\ \mathbf{V}'_3 \\ \vdots \\ \mathbf{V}'_{K-1} \\ \mathbf{V}'_K \\ \mathbf{V}'_{K+1} \end{pmatrix} = \widetilde{\mathbf{L}}\mathbf{V} = \begin{pmatrix} \widetilde{\mathbf{T}}^{(6)}_{00}\mathbf{V}_1 + \widetilde{\mathbf{T}}^{(6)}_{01}\mathbf{V}_2 \\ \mathbf{V}_3 \\ \mathbf{V}_4 \\ \vdots \\ \mathbf{V}_K \\ \mathbf{V}_{K+1} \\ \mathbf{V}_1 \end{pmatrix} \tag{I4}$$

Hence, the transfer-matrix $\widetilde{\mathbf{L}}$ works like a conveyor belt by shifting the vector blocks in a cyclic manner by one position up and processing the two top-most blocks by multiplying them by $\widetilde{\mathbf{T}}^{(6)}_{00}$ and $\widetilde{\mathbf{T}}^{(6)}_{01}$ matrices, respectively.

Combining this technique with the power iteration method, it is not hard then to get the value of the DNA partition function, which is defined by Eq. (I2).

By using this approach, it has been found in our study that in the case of $b = 3.4$ nm DNA segment size and $K = 14$ DNA segments constrained inside a single nucleosome (i.e., $\sim 145 - 147$ bp, see Table I), the offset free energy of nucleosomes equals to: $\mu_{\rm off} = -25.2\ k_{\rm B}T$.

2) *Finding the stationary phase electrostatic potential,* $\psi_{\rm sp}$.

The next step of the algorithm is to obtain a solution to Eq. (A37) describing the electrostatic potential, $\psi_{\rm sp}$, which corresponds to the stationary phase of the integrand in Eq. (A20).

As has been mentioned in Appendix A, this can be done by numerically minimizing the value of the integral shown in Eq. (A39) with respect to the parameters $(r, R_0, \omega, \psi_0, ..., \psi_{n_{\rm max}})$ characterizing the shape of $\psi$ electrostatic potential. The only small drawback of this method is that it requires knowledge of $\frac{\delta \ln Z_\psi}{\delta \psi(r)}$ functional derivative, which makes a contribution to the integrand, as can be seen from Eq. (A40).

One of the simplest ways to calculate $\frac{\delta \ln Z_\psi}{\delta \psi(r)}$ derivative is by using Eq. (H19). Indeed, with the help of Eq. (H17)-Eq. (H18) and Eq. (H20), it is not hard to construct $\widetilde{\mathbf{L}}$ and $\frac{\delta}{\delta\psi(r)}\widetilde{\mathbf{L}}$ matrices and the boundary condition vectors, $\widetilde{\mathbf{U}}$ and $\widetilde{\mathbf{Y}}$. Then by utilizing the power iteration method, one can easily find the dominant eigenvalue of the transfer-matrix, $\lambda_{\rm max}$, as well as ${\rm Pr}^{\rm L}_{\lambda_{\rm max}}\widetilde{\mathbf{U}}$ and ${\rm Pr}^{\rm R}_{\lambda_{\rm max}}\widetilde{\mathbf{Y}}$ projections of the vectors $\widetilde{\mathbf{U}}$ and $\widetilde{\mathbf{Y}}$. Finally, by substituting the acquired vector projections, the matrix $\frac{\delta}{\delta\psi(r)}\widetilde{\mathbf{L}}$ and the dominant eigenvalue, $\lambda_{\rm max}$, into Eq. (H19), it is straightforward to obtain the desired functional derivative, $\frac{\delta \ln Z_\psi}{\delta \psi(r)}$.

The only thing that should be noted here is that during the construction of the matrices $\widetilde{\mathbf{L}}$ and $\frac{\delta}{\delta\psi(r)}\widetilde{\mathbf{L}}$ as well as the vector $\widetilde{\mathbf{Y}}$, one has to set the force $f$ to be equal to zero ($f = 0$) in the respective formulas for the elements of $\widetilde{\mathbf{T}}^{(5)}_{nm}$ and $\widetilde{\boldsymbol{\xi}}^{(4)}_0$ matrix-blocks. As for the electrostatic potential $\psi$ present in Eq. (E8) and Eq. (H21), it should be calculated according to Eq. (A38).

Finally, it should be mentioned that similarly to the case of the DNA partition function, one does not actually need to work with the whole matrices $\widetilde{\mathbf{L}}$ and $\frac{\delta}{\delta\psi(r)}\widetilde{\mathbf{L}}$ to run the above computations. Instead, it is possible to utilize the same conveyor belt method described by Eq. (I3)-Eq. (I4), which, due to Eq. (H20), takes the following form:

$$\mathbf{V}' = \begin{pmatrix} \mathbf{V}'_1 \\ \mathbf{V}'_2 \\ \mathbf{V}'_3 \\ \vdots \\ \mathbf{V}'_{K+1} \end{pmatrix} = \frac{\delta}{\delta\psi(r)}\widetilde{\mathbf{L}} \times \mathbf{V} = \begin{pmatrix} \frac{\delta}{\delta\psi(r)}\widetilde{\mathbf{T}}^{(6)}_{00} \times \mathbf{V}_1 + \frac{\delta}{\delta\psi(r)}\widetilde{\mathbf{T}}^{(6)}_{01} \times \mathbf{V}_2 \\ 0 \\ 0 \\ \vdots \\ 0 \end{pmatrix} \tag{I5}$$

3) *Calculation of the partition function of DNA,* $Z_{\psi_{\rm sp}}$, *in the stationary phase electrostatic potential,* $\psi_{\rm sp}$.

As soon as the stationary phase electrostatic potential, $\psi_{\rm sp}$, is known, it is straightforward to find the partition function of DNA, $Z_{\psi_{\rm sp}}$, by using a combination of Eq. (E1) and Eq. (H16).

Alternatively, in the case when the chromosomal DNA is sufficiently long and there are no specific interactions between different chromosomes, one can ignore the boundary effects introduced by $\widetilde{\mathbf{U}}$ and $\widetilde{\mathbf{Y}}$ vectors and calculate the DNA partition function solely with the help of Eq. (H16) by putting $N = N_{\rm tot}$, where $N_{\rm tot}$ is the total number of DNA segments in all the chromosomes, see Appendix E.

In both cases, the DNA transfer-matrix, $\widetilde{\mathbf{L}}$, and the boundary condition vectors, $\widetilde{\mathbf{U}}$ and $\widetilde{\mathbf{Y}}$, are constructed by utilizing the same Eq. (H17)-Eq. (H18), with the stationary phase electrostatic potential, $\psi_{\rm sp}$, being put in the place of the field $\psi$. By applying the power iteration method, it is then straightforward to obtain ${\rm Pr}^{\rm L}_{\lambda_{\rm max}}\widetilde{\mathbf{U}}$ and ${\rm Pr}^{\rm R}_{\lambda_{\rm max}}\widetilde{\mathbf{Y}}$ projections of the vectors $\widetilde{\mathbf{U}}$ and $\widetilde{\mathbf{Y}}$, and, as a result, to evaluate the DNA partition function, $Z_{\psi_{\rm sp}}$.

In *Results* section of the main text, we also consider a scenario, in which a small part in the middle of chromosomal DNA is stretched by a mechanical force, $F$. It is clear that in order to find the DNA partition function in this case, it is necessary to construct two transfer-matrices, $\widetilde{\mathbf{L}}$ and $\widetilde{\mathbf{L}}_F$, for mechanically relaxed and stretched parts of the DNA, respectively. To this aim, it is still possible to use Eq. (H17) and Eq. (H18), where for the relaxed and stretched parts of DNA one needs to put $f = 0$ and $f = F$, accordingly, in the respective formulas for $\widetilde{\mathbf{T}}^{(5)}_{nm}$ transfer-matrix blocks. Furthermore, since stretching of a small part of DNA is unlikely to cause large changes in the distribution of the rest of the chromosomal DNA, it is clear that to construct $\widetilde{\mathbf{L}}$ and $\widetilde{\mathbf{L}}_F$ matrices we still can exploit the same stationary phase electrostatic potential, $\psi_{\text{sp}}$, as before. As for the vectors $\widetilde{\mathbf{U}}$ and $\widetilde{\mathbf{Y}}$, they can be found via Eq. (H17) and Eq. (H18).

By utilizing the power iteration algorithm in combination with the conveyor belt technique, it is not hard then to obtain the value of the DNA partition function. However, it should be noted that this time the formula for the DNA partition function takes a slightly different form from Eq. (I2):

$$Z_{\psi_{\text{sp}}} = V_{\text{B}}\, \widetilde{\mathbf{U}}\, \widetilde{\mathbf{L}}^{N'}\, \widetilde{\mathbf{L}}_F^{N_F}\, \widetilde{\mathbf{L}}^{N''} \widetilde{\mathbf{Y}} = C_0\, \lambda_{\max}^{N_{\text{tot}}-N_F}\, [\text{Pr}^{\text{L}}_{\lambda_{\max}} \widetilde{\mathbf{U}} \times \widetilde{\mathbf{L}}_F^{N_F} \times \text{Pr}^{\text{R}}_{\lambda_{\max}} \widetilde{\mathbf{Y}}] \tag{I6}$$

Where $N_F$ is the total number of mechanically stretched DNA segments; $N'$ and $N''$ are the numbers of relaxed DNA segments upstream and downstream of the stretched segments, such that $N' + N_F + N'' = N_{\text{tot}} - 1$. Prefactor $C_0$, the dominant eigenvalue, $\lambda_{\max}$, of the transfer-matrix $\widetilde{\mathbf{L}}$, and $\text{Pr}^{\text{L}}_{\lambda_{\max}} \widetilde{\mathbf{U}}$ and $\text{Pr}^{\text{R}}_{\lambda_{\max}} \widetilde{\mathbf{Y}}$ projections of the vectors $\widetilde{\mathbf{U}}$ and $\widetilde{\mathbf{Y}}$ in the above formula – are all calculated with the help of the power iteration algorithm. As for $\text{Pr}^{\text{L}}_{\lambda_{\max}} \widetilde{\mathbf{U}} \times (\widetilde{\mathbf{L}}_F)^{N_F} \times \text{Pr}^{\text{R}}_{\lambda_{\max}} \widetilde{\mathbf{Y}}$ product, it can be either directly computed if $N_F$ number is not very large, or it can be found by using the power iteration method, otherwise.

Having in hand the above algorithm for the DNA partition function, it is then straightforward to get the total free energy of the system, $G$, via Eq. (A44). By differentiating it with respect to the model parameters, one can obtain estimations of various observables characterizing the physical state of the system. For example, the average pressures of the chromosomal DNA, monovalent ions and cytosolic macromolecules on the NE, $p_{\text{DNA}}$, $p_{\text{ions}}$ and $p_{\text{macro}}$, can be found via the following formulas [see more details in Appendix K]:

$$p_{\text{DNA}} = -\frac{\partial G_{\text{DNA}}}{\partial V_{\text{nucl}}}\bigg|_{\substack{T=\text{const}\\ \forall j:\, n_j=\text{const}}} \quad \text{and} \quad p_{\text{ions}} = -\frac{\partial G_{\text{ions}}}{\partial V_{\text{nucl}}}\bigg|_{\substack{T=\text{const}\\ \forall j:\, n_j=\text{const}}} \quad \text{and} \quad p_{\text{macro}} = -\frac{\partial G_{\text{macro}}}{\partial V_{\text{nucl}}}\bigg|_{\substack{T=\text{const}\\ \forall j:\, n_j=\text{const}}} \tag{I7}$$

Here $G_{\text{DNA}}$, $G_{\text{ions}}$ and $G_{\text{macro}}$ are respectively the free energies of DNA, monovalent ions and cytosolic macromolecules, which are defined by Eq. (K1).

Furthermore, similarly to Eq. (I1), for the average occupancy fraction of the chromosomal DNA by nucleosomes, $O_{\text{nucl}}$, we have:

$$O_{\text{nucl}} = -\frac{K}{N_{\text{tot}}}\frac{\partial G}{\partial \mu_{\text{pr}}} = \frac{K}{\beta N_{\text{tot}}}\frac{\partial \ln Z_{\psi_{\text{sp}}}}{\partial \mu_{\text{pr}}} \tag{I8}$$

In the case when a small part of DNA is stretched by a mechanical force, $F$, Eq. (I8) can be applied directly to the stretched part of DNA to find out how the force affects stability of nucleosomes. By doing so, it is easy to obtain the following formula:

$$O_{\text{nucl}}(F) = -\frac{K}{N_F}\frac{\partial G}{\partial \mu_{\text{pr}}}\bigg|_{\widetilde{\mathbf{L}}=\text{const}} = \frac{K}{\beta N_F}\frac{\partial \ln Z_{\psi_{\text{sp}}}}{\partial \mu_{\text{pr}}}\bigg|_{\widetilde{\mathbf{L}}=\text{const}} \tag{I9}$$

Here $N_F$ is the total number of stretched DNA segments. In the above equation, the transfer-matrix $\widetilde{\mathbf{L}}$ describing mechanically relaxed DNA segments is treated as a constant during the differentiation process, and only the matrix $\widetilde{\mathbf{L}}_F$, which corresponds to mechanically stretched DNA segments, is assumed to be dependent on the protein binding free energy, $\mu_{\text{pr}}$, see Eq. (I6) for more details regarding $\widetilde{\mathbf{L}}$ and $\widetilde{\mathbf{L}}_F$ matrices.

In addition to the average occupancy fraction of DNA, one can also find the force-extension curve of the stretched DNA part by using the next formula, which can be obtained by substituting Eq. (B17) generalized to an arbitrary value of $K$ into Eq. (C5) with subsequent differentiation of the DNA partition function with respect to the force:

$$z(F) = -\frac{\partial G}{\partial F} = k_{\mathrm{B}}T\frac{\partial \ln Z_{\psi_{\mathrm{sp}}}}{\partial F} \tag{I10}$$

Where $z(F)$ is the extension of the stretched DNA part along the force direction.

In a very similar way, it can be shown that the mean squared displacement (MSD) between any two points residing on the DNA contour equals to:

$$\mathrm{MSD} = \langle \Delta x^2 + \Delta y^2 + \Delta z^2 \rangle = -\frac{3}{\beta}\frac{\partial^2 G}{\partial F^2}\bigg|_{F=0} = \frac{3}{\beta^2}\frac{\partial^2 \ln Z_{\psi_{\mathrm{sp}}}}{\partial F^2}\bigg|_{F=0} \tag{I11}$$

Where the stretching force, $F$, is applied only to the DNA part connecting these two points. Here we have taken into account that $\langle \Delta x^2 + \Delta y^2 + \Delta z^2 \rangle = 3\langle \Delta z^2 \rangle$ in the case of a spherically symmetric cell nucleus or a viral capsid. See also Appendix J for more details regarding the MSD calculations.

From Eq. (A13), Eq. (A19), Eq. (A44) and Eq. (B7) one can also obtain the following formula for the net electrical charge density of protein-covered DNA, $\rho_{\mathrm{DNA}}(r)$, as a function of the radial distance, $r$:

$$\rho_{\mathrm{DNA}}(r) = \frac{1}{4\pi r^2}\frac{\delta G}{\delta\psi(r)}\bigg|_{\substack{\mu_{\mathrm{pr}}=\mathrm{const}\\ \psi=\psi_{\mathrm{sp}}}} = -\frac{1}{4\pi\beta r^2}\frac{\delta \ln Z_{\psi}}{\delta\psi(r)}\bigg|_{\substack{\mu_{\mathrm{pr}}=\mathrm{const}\\ \psi=\psi_{\mathrm{sp}}}} \tag{I12}$$

Where $Z_{\psi}$ is the partition function of DNA in the electrostatic potential $\psi$. In the above expression, $\frac{\delta \ln Z_{\psi}}{\delta\psi(r)}$ functional derivative is calculated at the stationary phase electrostatic potential, $\psi_{\mathrm{sp}}(r)$, by treating the protein binding free energy to DNA, $\mu_{\mathrm{pr}}$, as a fixed constant. I.e., $\frac{\delta\mu_{\mathrm{pr}}}{\delta\psi(r)}$ functional derivative must be set equal to zero in Eq. (H19) and Eq. (H20) to get a correct spatial distribution of the net electrical charge of chromatin inside the cell nucleus.

Similarly, by using Eq. (I12) and Eq. (H19)-Eq. (H21), it is not very hard to derive formulas for the spatial densities of bare DNA segments, $\rho_{\mathrm{bare}}(r)$, and nucleosomes, $\rho_{\mathrm{nucl}}(r)$, inside the cell nucleus as functions of the radial distance, $r$:

$$\rho_{\mathrm{bare}}(r) = -\frac{1}{4\pi\beta r^2 q_{00}}\frac{\delta \ln Z_{\psi}}{\delta\psi(r)}\bigg|_{\substack{\Phi_{K0}=\mathrm{const}\\ \mu_{\mathrm{pr}}=\mathrm{const}\\ \psi=\psi_{\mathrm{sp}}}} \quad \text{and} \quad \rho_{\mathrm{nucl}}(r) = -\frac{1}{4\pi\beta r^2 q_{K0}}\frac{\delta \ln Z_{\psi}}{\delta\psi(r)}\bigg|_{\substack{\Phi_{00}=\mathrm{const}\\ \mu_{\mathrm{pr}}=\mathrm{const}\\ \psi=\psi_{\mathrm{sp}}}} \tag{I13}$$

Where $\rho_{\mathrm{bare}}(r)$ equals to the average number of bare DNA segments (each of length $b$) per unit of the cell nucleus volume, and $\rho_{\mathrm{nucl}}(r)$ – to the average number of nucleosomes per unit of the cell nucleus volume. In the above equation, notations $\Phi_{00} = \mathrm{const}$ and $\Phi_{K0} = \mathrm{const}$ mean that these functions must be treated as fixed constants, i.e., their functional derivatives must be set equal to zero in the corresponding formulas for the transfer-matrix blocks: $\frac{\delta\Phi_{00}}{\delta\psi(r)} = 0$ and $\frac{\delta\Phi_{K0}}{\delta\psi(r)} = 0$, accordingly.

By following the above examples, it is possible to obtain the average values of other observable parameters as well. However, for our purposes, Eq. (I7)-Eq. (I13) will suffice as they provide all the necessary information for this study. The only thing that should be mentioned here is that some care must be taken during numeric calculations based on these formulas, especially in the case of Eq. (I11) which deals with a higher-order derivative of the DNA partition function. For more details, see the next Appendix section.

##### Appendix J: Mean squared displacement between two points on DNA

In this section, we would like to discuss some of the difficulties that arise in computation of the mean squared distance (MSD) as well as the root mean squared distance (RMSD) between two points on DNA and a possible way by which they can be resolved. To this aim, we will use the RMSD-genomic distance curves displayed in Figure 2(h) as an example since this will allow us to gain deeper insight into the problem.

Specifically, direct application of Eq. (I11) to calculation of the MSD- and RMSD-genomic distance curves shows that while this formula provides physically meaningful results for sufficiently large viral capsids ($R_{\mathrm{vir}} \gtrsim 200$ nm), in the case of smaller capsid radii it may lead to wrong values of the MSD and RMSD between two points on DNA. Indeed, for a viral capsid of several nanometers in size, it is reasonable to expect that the MSD and RMSD curves must reach a plateau at sufficiently large genomic distances as shown in Figure 2(h), and, in fact, direct numeric simulations of polymers confined in a tight space support such expectations, see ref. [41]. However, calculations based on Eq. (I11) predict a completely different picture, demonstrating the linear change of the MSD curve at $\geq 1.5$ kbp genomic distances in the case of $R_{\text{vir}} < 200$ nm, which never reaches a plateau, with equally incorrect behaviour of the RMSD curve, see, for example, Figures S12(a, b).

By varying the model parameters, it is not hard to find out the origin of such an erroneous behaviour of the MSD and RMSD curves, which seems to be high sensitivity of Eq. (I11) to the total number of reciprocal lattice nodes, $l_{\text{max}}$, used in the model computations – compare, for example, the red and the blue curves in Figures S12(a, b). From these graphs it can be seen that $l_{\text{max}}$ number has a very strong effect on the right sides of the MSD and RMSD curves corresponding to large genomic distances. At the same time, it can be noted that the left-most parts of the MSD and RMSD curves corresponding to short genomic distances do not change much with $l_{\text{max}}$. Altogether, these results indicate that while Eq. (I11) seems to accurately predict the MSD between two points residing on DNA at short genomic distances, for longer ones it provides wrong estimations, which may be caused by a limited number of reciprocal lattice nodes that can be taken into account in model calculations.

While this may look like an intrinsic problem of Eq. (I11), which does not have a simple solution, it is nevertheless possible to make necessary corrections in the model to obtain accurate predictions of the MSD and RMSD curves. To see this, the first thing we need to do is to find out more details regarding the origin of the MSD curve instability with respect to $l_{\text{max}}$ parameter, which can be done by deriving a more explicit formula for the MSD between two points on DNA.

After substituting the first lane of Eq. (B17) describing the bare DNA energy into Eq. (C5) and then applying Eq. (I11), it can be shown that the MSD formula takes the following form:

$$\text{MSD} = b^2 \sum_{n,m=0}^{N_F-1} \langle \mathbf{z}_{n_0+n} \cdot \mathbf{z}_{n_0+m} \rangle = b^2 N_F + 2b^2 \sum_{n=0}^{N_F-1} \sum_{m=n+1}^{N_F-1} \langle \mathbf{z}_{n_0+n} \cdot \mathbf{z}_{n_0+m} \rangle \tag{J1}$$

Where $b$ is the size of bare DNA segments; $N_F$ is the number of DNA segments connecting the two points of interest for which the MSD is calculated; $n_0$ is the index of the first DNA segment in the DNA part linking these points; $\mathbf{z}_{n_0+n} = \mathbf{R}_{n_0+n}\mathbf{z}_0$ and $\mathbf{z}_{n_0+m} = \mathbf{R}_{n_0+m}\mathbf{z}_0$ are unit vectors tangent to the $(n_0+n)^{\text{th}}$ and $(n_0+m)^{\text{th}}$ DNA segments, respectively; finally, $\langle \mathbf{z}_{n_0+n} \cdot \mathbf{z}_{n_0+m} \rangle$ is the ensemble average correlation between the orientations of the $(n_0+n)^{\text{th}}$ and $(n_0+m)^{\text{th}}$ DNA segments $(m > n)$, which, with the help of Eq. (I6), can be found to be:

$$\langle \mathbf{z}_{n_0+n} \cdot \mathbf{z}_{n_0+m} \rangle = \frac{3}{\beta^2 b^2} \frac{1}{\lambda_{\max}^{m-n+1}} \frac{\text{Pr}^{\text{L}}_{\lambda_{\max}} \widetilde{\mathbf{U}} \times \frac{\partial}{\partial F} \widetilde{\mathbf{L}}_F \Big|_{F=0} \times \widetilde{\mathbf{L}}^{m-n-1} \times \frac{\partial}{\partial F} \widetilde{\mathbf{L}}_F \Big|_{F=0} \times \text{Pr}^{\text{R}}_{\lambda_{\max}} \widetilde{\mathbf{Y}}}{\text{Pr}^{\text{L}}_{\lambda_{\max}} \widetilde{\mathbf{U}} \times \text{Pr}^{\text{R}}_{\lambda_{\max}} \widetilde{\mathbf{Y}}} \tag{J2}$$

From Eq. (J1) it is clear that in the case of zero tangent vector correlations ($\forall n \forall m,\ 0 \leq n, m \leq N_F - 1 : \langle \mathbf{z}_{n_0+n} \cdot \mathbf{z}_{n_0+m} \rangle = 0$) the formula for the MSD reduces to: $\text{MSD} = b^2 N_F$, which is exactly the result predicted by the freely joint chain model of polymers [42]. Thus, it can be concluded that all the problems with the MSD curve sensitivity to $l_{\text{max}}$ number stem from the right-hand sum in Eq. (J1) over the tangent vector correlations.

By taking into account the spherical symmetry of the system, it is clear that the average extension between any two points on DNA is equal to zero. As a result, it immediately follows from Eq. (I10) that:

$$\text{Pr}^{\text{L}}_{\lambda_{\max}} \widetilde{\mathbf{U}} \times \frac{\partial}{\partial F} \widetilde{\mathbf{L}}_F \Big|_{F=0} \times \text{Pr}^{\text{R}}_{\lambda_{\max}} \widetilde{\mathbf{Y}} = 0 \tag{J3}$$

Substituting Eq. (J3) into Eq. (J2) and by using the same derivation as in the proof of the power iteration method [39], it is then not hard to see that all the tangent vector correlations, $\langle \mathbf{z}_{n_0+n} \cdot \mathbf{z}_{n_0+m} \rangle$, are in fact proportional to $(\lambda'_{\max}/\lambda_{\max})^{m-n-1}$ ratio at large separation distances $(m - n \gg 1)$, where $\lambda'_{\max}$ is the second dominant eigenvalue of the transfer-matrix $\mathbf{L}$. Based on this observation, it becomes clear why the DNA partition function is robust with respect to $l_{\max}$ number, while MSD and RMSD curves show high sensitivity to it. Indeed, direct numeric calculations indicate that $\lambda_{\max}$ and $\lambda'_{\max}$ typically have very close values. As a result, while $l_{\max}$ may have a sufficiently weak effect on these two eigenvalues, their tiny changes become more pronounced in $\lambda_{\max} - \lambda'_{\max}$ difference, further amplifying via $(\lambda'_{\max}/\lambda_{\max})^{m-n-1}$ ratio at large separation distances in the formula for the tangent vector correlations [Eq. (J2)]. Eventually, by accumulating in the right-hand sum of Eq. (J1), these small errors lead to the observed deviation of the MSD and RMSD curves from their correct behaviour. In contrast, the DNA partition function value is determined mainly by the dominant eigenvalue of the transfer-matrix, $\lambda_{\max}$ [see, for example, Eq. (I2)], and is practically independent from the second dominant eigenvalue $\lambda'_{\max}$, making it much more robust quantity towards changes in $l_{\max}$ number.

To provide additional insights into the problem, we have also plotted a couple of curves for $R_{\text{vir}} = 30$ nm case demonstrating change in $\langle \mathbf{z}_{n_0+n} \cdot \mathbf{z}_{n_0+m} \rangle$ tangent vector correlation vs the genomic distance separating the $(n_0+n)^{\text{th}}$ and the $(n_0+m)^{\text{th}}$ DNA segments for two different values of $l_{\max}$, see the red and the blue curves in Figure S12(c). Surprisingly, it has been found that the tangent vector correlation function is rather robust towards the number of the unique-value nodes used in the model calculations, suggesting that the largest error accumulation takes place only during the last step – summation over all tangent vector correlations in Eq. (J1).

To see this, one just needs to rewrite Eq. (J1) in the following alternative form:

$$\text{MSD} = b^2 N_F + 2b^2 \sum_{n=1}^{N_F-1} (N_F - n) \langle \mathbf{z}_{n_0} \cdot \mathbf{z}_{n_0+n} \rangle \tag{J4}$$

Where we have taken into account that $\langle \mathbf{z}_{n_1} \cdot \mathbf{z}_{m_1} \rangle = \langle \mathbf{z}_{n_2} \cdot \mathbf{z}_{m_2} \rangle$ as soon as $m_1 - n_1 = m_2 - n_2$ and the points between which the MSD is measured are located sufficiently far away from the DNA ends (at least by a few DNA persistence lengths, $A = 50$ nm), such that the DNA boundary effects can be neglected.

Then from Eq. (J4) and the exponential decay of the tangent vector correlation function that can be seen in Figure S12(c) it follows that the asymptotic linear slope of the MSD curve at large genomic distances ($N_F \gg 1$) approximately equals to the following sum:

$$\text{MSD slope} \approx b^2 + 2b^2 \sum_{n=1}^{N_F-1} \langle \mathbf{z}_{n_0} \cdot \mathbf{z}_{n_0+n} \rangle \tag{J5}$$

After plotting the right-hand side of Eq. (J5) as a function of the genomic distance, $bN_F$, it becomes clear that the linear change of the MSD observed in Figure S12(a) results from the failure of the above sum to converge to zero at large genomic distances, see Figure S12(d).

In fact, it is not hard to obtain even more detailed insights into the mathematical origin of this problem by noting that the tangent vector correlation graphs shown in Figure S12(d) can be perfectly fitted to the following damped oscillation function [see the yellow and cyan dotted fitting curves in Figure S12(d)]:

$$g(x) = e^{-\lambda x} \, \frac{\cos(\frac{2\pi x}{T} + \phi)}{\cos\phi} \tag{J6}$$

Where $x$ is the genomic distance along the DNA contour; $\lambda$ and $T$ are the decay constant and periodicity of the curve oscillations, respectively; $\phi$ is a phase shift of the function.

It is then easy to see that the sum over the tangent vector correlations in Eq. (J5) can be approximated by the next integral:

$$I = \frac{1}{\cos\phi} \int_0^{+\infty} e^{-\lambda x} \, \cos\left(\frac{2\pi x}{T} + \phi\right) \mathrm{d}x = \frac{\lambda T^2 - 2\pi T \tan\phi}{4\pi^2 + \lambda^2 T^2}, \tag{J7}$$

from which it immediately follows that the integral can take any value in $(-\infty, +\infty)$ range due to the presence of

$\tan\phi$ term in the numerator of the above formula. Furthermore, because of a high sensitivity of $\tan\phi$ function to its argument, $\phi$, it is clear that the integral will be also sensitive to the phase shift of the damped oscillation function described by Eq. (J6). And while the sum in Eq. (J5) is not exactly equal to the integral in Eq. (J7), this simple example demonstrates that even sufficiently small horizontal displacements in the tangent vector correlation function can lead to drastic variations in the MSD curve behaviour at large genomic distances. On the other hand, it can be concluded that in order to correct non-physical linear behaviour of the MSD and RMSD curves at large genomic distances, all we need to do is to introduce a small phase shift into the tangent vector correlation function along the horizontal axis.

The only thing that we need to keep in mind while doing this is that the tangent vector correlation function has to pass through unit at zero genomic distance since the ensemble average of the squared $\mathbf{z}_{n_0}$ tangent unit vector equals to: $g(x=0) = \langle \mathbf{z}_{n_0}^2 \rangle = \langle 1 \rangle = 1$. Thus, a proper renormalization of the tangent vector correlation function needs to be done after the phase shift introduction into it. Following the above notes, it is not hard to arrive to the next formula:

$$\widetilde{g}(x) = \mathrm{Renorm}[\, g(x), x_0 \,] = \frac{g(x - x_0)}{g(-x_0)} \tag{J8}$$

Where $\widetilde{g}(x)$ is the modified tangent vector correlation function, and $x_0$ is the introduced shift of the function along the horizontal axis.

It should be noted that Eq. (J8) possesses several nice features.

First, it is known that in the case of the worm-like chain (WLC) model of semiflexible polymers, the tangent vector correlation is described by a simple exponential decay function: $\langle \mathbf{z}_n \cdot \mathbf{z}_m \rangle = e^{-b|m-n|/A} = g(x) = e^{-x/A}$, see ref. [42]. Here $b$ is the DNA segment size, $x = b\,|m-n|$ is the genomic distance along the DNA contour, and $A$ is the DNA persistence length. By applying Eq. (J8) to this function, it is easy to see that:

$$\forall x_0 \in \mathbb{R} : \mathrm{Renorm}[\, e^{-x/A}, x_0 \,] = e^{-x/A} \tag{J9}$$

I.e., the renormalization procedure does not change the exponential function describing tangent vector correlations in the WLC model of semiflexible polymers, such as DNA. It then can be concluded that for viral particles with $R_{\mathrm{vir}} \gtrsim 200$ nm radius, whose tangent vector correlation curves pass near the correlation function of the WLC model [see Figure 2(g)], the above renormalization will barely introduce any corrections. As a result, there will be practically no change in the respective MSD- and RMSD-genomic distance curves. Therefore, the most significant corrections of the MSD and RMSD curves will take place only for small viral particles ($R_{\mathrm{vir}} \lesssim 100$ nm), which is exactly where we want them to be.

Second, in the case of the damped oscillation function defined by Eq. (J6), which has been found to perfectly describe the tangent vector correlation curves of small viral particles [see, for example, Figure S12(c)], it is not hard to show that:

$$\mathrm{Renorm}\left[ e^{-\lambda x}\, \frac{\cos(\frac{2\pi x}{T} + \phi)}{\cos\phi}, x_0 \right] = e^{-\lambda x}\, \frac{\cos(\frac{2\pi x}{T} + \phi_0)}{\cos\phi_0} \tag{J10}$$

Where $\phi_0$ is a modified phase shift of the function, which equals to: $\phi_0 = \phi - \frac{2\pi x_0}{T}$.

From Eq. (J10) it can be seen that even if for some reason one gets carried away in his search for the correct value of the parameter $x_0$, missing it by an integer number of the period $T$ (i.e., $x_0' = x_0 + nT$, $n \in \mathbb{Z}$), he will still get a mathematically accurate final result due to the periodicity of the cosine function. In application to the tangent vector correlation function, this simply means that the renormalization procedure described by Eq. (J8) is rather stable towards different types of errors that may appear in pursue of the right value of the parameter $x_0$.

Anyway, by applying such a renormalization to the tangent vector correlation curve shown in Figure S12(c), it is easy to see that only a tiny horizontal shift of the curve is needed to obtain MSD and RMSD graphs demonstrating physically correct behaviour, which this time reach plateaus at large genomic distances, suggesting that the proposed renormalization procedure works rather well [see the black curves in Figures S12(a, b)]. This approach has been used in our study to plot all the RMSD-genetic distance curves for viral particles shown in Figure 2(h). The total number of reciprocal lattice nodes used in the corresponding model calculations was $l_{\max} = 14769$ (i.e., $\widetilde{l}_{\max} = 1512$).

##### Appendix K: Pressure balance on the nuclear envelope

As discussed in the main text, the volume of the cell nucleus is generally determined by mechanical equilibrium [Eq. (6)] between pressures acting on the nuclear envelope (NE), which are created by different cell components. This includes the pressure generated by chromosomal DNA confined inside the nucleus as well as an outside osmotic pressure resulting from the gradient of electrically charged ions and small cell metabolites across the NE. Furthermore, cytosolic macromolecules that cannot move through nuclear pore complexes into the nucleus create an additional osmotic pressure onto the NE [43, 44]. Finally, it has been previously proposed that a tug-of-war between the NE and ER, which share the same lipid membrane, may also contribute to regulation of the cell nucleus volume [45]. Unfortunately, due to complete absence of quantitative models able to describe relative contributions of the above factors into mechanical equilibrium of the cell nucleus, there have been a little progress in understanding of physical mechanisms controlling the cell nucleus size.

The first thing we need to do to fill the above gap is to consider free energies of the aforementioned cell elements. Let $G_{\mathrm{NE}}$, $G_{\mathrm{ER}}$, $G_{\mathrm{macro}}$, $G_{\mathrm{DNA}}$, $G_{\mathrm{ions}}$ be the free energies of the NE, ER, cytosolic macromolecules, DNA and monovalent ions, respectively. It is clear that all these energies in the general case must be functions of the nucleus volume, $V_{\mathrm{nucl}}$. Indeed, free energies of DNA, monovalent ions and cytosolic macromolecules are equal to:

$$G_{\mathrm{DNA}} = -k_{\mathrm{B}}T \ln Z_{\psi_{\mathrm{sp}}} \quad \text{and} \quad G_{\mathrm{ions}} = \sum_{\substack{\text{monovalent}\\ \text{ions}}} n_j \mu_j^{\psi_{\mathrm{sp}}} \quad \text{and} \quad G_{\mathrm{macro}} = \sum_{\substack{\text{macro-}\\ \text{molecules}}} n_j \mu_j^{\psi_{\mathrm{sp}}} \tag{K1}$$

Where $Z_{\psi_{\mathrm{sp}}}$ and $\mu_j^{\psi_{\mathrm{sp}}}$ are the partition function of DNA and the electrostatic chemical potential of the $j^{\mathrm{th}}$ type of particles in the stationary phase field, $\psi_{\mathrm{sp}}$, given by Eq. (E1) and Eq. (A42), respectively. $n_j$ is the total number of the $j^{\mathrm{th}}$ type of particles in a living cell. Summation in the formula for $G_{\mathrm{ions}}$ energy is carried out over all cytosolic monovalent ions ($K^+$, $Na^+$, $Cl^-$) and small electrically charged metabolites. Similarly, in the formula for $G_{\mathrm{macro}}$ energy, summation is performed over all types of cytosolic macromolecules, which cannot move through nuclear pore complexes.

Since $Z_{\psi_{\mathrm{sp}}}$ and $\mu_j^{\psi_{\mathrm{sp}}}$ are functions of the nuclear volume, it immediately follows that all of the above free energies, $G_{\mathrm{DNA}}$, $G_{\mathrm{ions}}$ and $G_{\mathrm{macro}}$, also depend on the volume of the cell nucleus, leading to emergence of the corresponding pressures, $p_{\mathrm{DNA}}$, $p_{\mathrm{ions}}$ and $p_{\mathrm{macro}}$, acting on the NE, which are defined by Eq. (I7).

In this study, $p_{\mathrm{DNA}}$ was calculated by interpolating the free energy of DNA, $G_{\mathrm{DNA}}$, which was computed on a fixed set of $R_{\mathrm{nucl}}$ values, with a smoothing spline, as shown in Figure S13, and then applying Eq. (I7) to the obtained fitting curve.

As for the remaining pressures, $p_{\mathrm{ions}}$ and $p_{\mathrm{macro}}$, it is not hard to obtain simple analytic formulas for them by utilizing previously derived Eq. (A42) for the electrochemical potential. Namely, from Eq. (A42), Eq. (I7) and Eq. (K1) it can be seen that:

$$p_{\mathrm{macro}} = -\left.\frac{\partial G_{\mathrm{macro}}}{\partial V_{\mathrm{nucl}}}\right|_{\substack{T=\mathrm{const}\\ \forall j:\, n_j=\mathrm{const}}} = k_{\mathrm{B}}T \sum_{\substack{\text{macro-}\\ \text{molecules}}} n_j \frac{\partial \ln I_j^{\psi_{\mathrm{sp}}}}{\partial V_{\mathrm{nucl}}} \tag{K2}$$

Here $I_j^{\psi_{\mathrm{sp}}}$ is the volume integral of the $j^{\mathrm{th}}$ type of particles, which is defined by Eq. (A15), in the stationary phase field, $\psi_{\mathrm{sp}}$.

By recalling that in this study the electrostatic potential of the cell cytoplasm is used as a reference point (and thus, $\psi_{\mathrm{cyto}} = 0$), it is not hard to see from Eq. (A15) that for cytosolic macromolecules we have: $I_j^{\psi_{\mathrm{sp}}} = V_{\mathrm{cell}}^{\mathrm{osm}} - V_{\mathrm{nucl}}$, where $V_{\mathrm{cell}}^{\mathrm{osm}}$ is the osmotically active volume of a cell. As a result, it is easy to obtain the following expression for the osmotic pressure created by cytosolic macromolecules on the NE:

$$p_{\text{macro}} = -\frac{n_{\text{macro}} k_{\text{B}} T}{V_{\text{cell}}^{\text{osm}} - V_{\text{nucl}}} \tag{K3}$$

Here $n_{\text{macro}}$ is the total number of macromolecules in the cell cytosol:

$$n_{\text{macro}} = \sum_{\substack{\text{macro-}\\\text{molecules}}} n_j \tag{K4}$$

In Eq. (K3), the minus sign in the front of the ratio means that cytosolic macromolecules create an osmotic pressure which is applied to the NE from outside, trying to reduce the volume of the cell nucleus.

In a similar way, by using Eq. (A35), Eq. (A36), Eq. (A42), Eq. (I7) and Eq. (K1), it is not hard to find the pressure created on the NE by monovalent ions and electrically charged small cell metabolites:

$$p_{\text{ions}} = -\left.\frac{\partial G_{\text{ions}}}{\partial V_{\text{nucl}}}\right|_{\substack{T=\text{const}\\ \forall j:\, n_j=\text{const}}} = k_{\text{B}}T \sum_{\substack{\text{monovalent}\\\text{ions}}} n_j \frac{\partial \ln I_j^{\psi_{\text{sp}}}}{\partial V_{\text{nucl}}} = k_{\text{B}}T \frac{\partial \ln I_+^{\psi_{\text{sp}}}}{\partial V_{\text{nucl}}} \sum_{\substack{\text{positive}\\\text{monovalent}}} n_j + k_{\text{B}}T \frac{\partial \ln I_-^{\psi_{\text{sp}}}}{\partial V_{\text{nucl}}} \sum_{\substack{\text{negative}\\\text{monovalent}}} n_j =$$
$$= c_{\text{ions}} k_{\text{B}} T \left[\frac{\partial I_+^{\psi_{\text{sp}}}}{\partial V_{\text{nucl}}} + \frac{\partial I_-^{\psi_{\text{sp}}}}{\partial V_{\text{nucl}}}\right] = c_{\text{ions}} k_{\text{B}} T \frac{\partial}{\partial V_{\text{nucl}}} \int_0^{R_{\text{nucl}}} 8\pi r^2 \left[\cosh(\beta q_{\text{e}} \psi_{\text{sp}}(r)) - 1\right] \mathrm{d}r \tag{K5}$$

Here $\cosh(x)$ is the hyperbolic cosine; $R_{\text{nucl}}$ is the nucleus radius, and $c_{\text{ions}}$ is the total cytosolic concentration of positive and negative monovalent ions as well as electrically charged small cell metabolites in a living cell, see comments before Eq. (A34).

The last partial derivative in Eq. (K5) with respect to $V_{\text{nucl}}$ was calculated in this study by interpolating the radial integral evaluated on a fixed set of nucleus radii with a smoothing spline, as shown in Figure S13, followed by application of $\frac{\partial}{\partial V_{\text{nucl}}}$ derivative to the obtained fitting curve.

It should be noted that such an approach provides an accurate estimation of $p_{\text{ions}}$ even though the total numbers of particles, $n_j$, are not fixed in these computations as required by the first equality in Eq. (K5). Specifically, as mentioned in Appendix A, $\psi_{\text{sp}}$ electrostatic potential is obtained based on the assumption that monovalent ions and electrically charged small cell metabolites follow the Boltzmann distribution described by Eq. (A34) and Eq. (A35). Thus, the corresponding particle numbers, $n_j$, in general will be dependent on the nucleus size. However, since the magnitude of the electrostatic potential considered in the model calculations is sufficiently low [from $-4$ mV to 0 mV, see Figure 4(a-b)] and since the nucleus volume is significantly smaller than the volume of the cell cytosol, it can be estimated that slight changes in the total numbers of monovalent ions and small cell metabolites at different values of $R_{\text{nucl}}$ result in an insignificant error of $< 5\% - 10\%$ in the value of $p_{\text{ions}}$.

As for viral particles, in their case the volume of the buffer solution is so much bigger than the volume of an individual viral particle that even large negative electrostatic potential of the viral capsid (up to $-100$ mV) will not results in considerable change in the total number of monovalent ions in the system due to the Boltzmann distribution described by Eq. (A34) and Eq. (A35). Thus, Eq. (K5) can be used to calculate the exact value of the osmotic pressure created by monovalent ions on the capsid wall.

So far, we have been focusing our attention on $G_{\text{DNA}}$, $G_{\text{ions}}$ and $G_{\text{macro}}$ free energies without touching upon the free energies associated with the NE and ER membrane, $G_{\text{NE}}$ and $G_{\text{ER}}$. Yet, as discussed in the main text, these energies play the central role in the nucleus size regulation due to emergence of associated pressures, $p_{\text{NE}}$ and $p_{\text{ER}}$, applied to the NE:

$$p_{\text{NE}} = -\left.\frac{\partial G_{\text{NE}}}{\partial V_{\text{nucl}}}\right|_{\substack{T=\text{const}\\ \forall j:\, n_j=\text{const}}} \quad \text{and} \quad p_{\text{ER}} = -\left.\frac{\partial G_{\text{ER}}}{\partial V_{\text{nucl}}}\right|_{\substack{T=\text{const}\\ \forall j:\, n_j=\text{const}}} \tag{K6}$$

In the general case, both $G_{\text{NE}}$ and $G_{\text{ER}}$ include several terms representing elastic bending and stretching energies of the membrane as well as its interaction with membrane-binding proteins, such as lamins or reticulons, see ref. [46–48]. As a result, formulas for $G_{\mathrm{NE}}$ and $G_{\mathrm{ER}}$ energies look rather complicated. However, it is still possible to express $p_{\mathrm{NE}}$ and $p_{\mathrm{ER}}$ pressures via the surface tensions of the NE and ER membrane, $\sigma_{\mathrm{NE}}$ and $\sigma_{\mathrm{ER}}$, which are defined as:

$$\sigma_{\mathrm{NE}} = \frac{1}{2}\frac{\partial G_{\mathrm{NE}}}{\partial S_{\mathrm{nucl}}}\bigg|_{\substack{T=\mathrm{const}\\ \forall j:\, n_j=\mathrm{const}}} \quad \text{and} \quad \sigma_{\mathrm{ER}} = \frac{\partial G_{\mathrm{ER}}}{\partial S_{\mathrm{ER}}}\bigg|_{\substack{T=\mathrm{const}\\ \forall j:\, n_j=\mathrm{const}}} \tag{K7}$$

Where $S_{\mathrm{nucl}}$ and $S_{\mathrm{ER}}$ are the surface areas of the cell nucleus and ER, respectively. Prefactor $\frac{1}{2}$ in the above formula for $\sigma_{\mathrm{NE}}$ appears due to the fact that the NE is composed of two bilayer lipid membranes as schematically shown in Figure 3(e).

The only thing that we need to keep in mind is that since the NE and ER share with each other the lipid membrane, Eq. (K7) has to be complemented with a formula describing the conservation law of the membrane surface area:

$$2S_{\mathrm{nucl}} + S_{\mathrm{ER}} = \mathrm{const} \tag{K8}$$

Where prefactor 2 again emphasizes the fact that the NE is comprised of two sheets of the bilayer lipid membrane in contrast to ER, which is enveloped only by a single lipid bilayer.

Combining Eq. (K6)-Eq. (K8), it is then straightforward to arrive to the following mathematical expressions for $p_{\mathrm{NE}}$ and $p_{\mathrm{ER}}$:

$$\begin{aligned} p_{\mathrm{NE}} &= -\frac{\partial G_{\mathrm{NE}}}{\partial V_{\mathrm{nucl}}}\bigg|_{\substack{T=\mathrm{const}\\ \forall j:\, n_j=\mathrm{const}}} = -\frac{\partial G_{\mathrm{NE}}}{\partial S_{\mathrm{nucl}}}\frac{\partial S_{\mathrm{nucl}}}{\partial R_{\mathrm{nucl}}}\left(\frac{\partial V_{\mathrm{nucl}}}{\partial R_{\mathrm{nucl}}}\right)^{-1}\bigg|_{\substack{T=\mathrm{const}\\ \forall j:\, n_j=\mathrm{const}}} = -\frac{4\sigma_{\mathrm{NE}}}{R_{\mathrm{nucl}}} \\ p_{\mathrm{ER}} &= -\frac{\partial G_{\mathrm{ER}}}{\partial V_{\mathrm{nucl}}}\bigg|_{\substack{T=\mathrm{const}\\ \forall j:\, n_j=\mathrm{const}}} = -\frac{\partial G_{\mathrm{ER}}}{\partial S_{\mathrm{ER}}}\frac{\partial S_{\mathrm{ER}}}{\partial S_{\mathrm{nucl}}}\frac{\partial S_{\mathrm{nucl}}}{\partial R_{\mathrm{nucl}}}\left(\frac{\partial V_{\mathrm{nucl}}}{\partial R_{\mathrm{nucl}}}\right)^{-1}\bigg|_{\substack{T=\mathrm{const}\\ \forall j:\, n_j=\mathrm{const}}} = \frac{4\sigma_{\mathrm{ER}}}{R_{\mathrm{nucl}}} \end{aligned} \tag{K9}$$

The physical meaning of $p_{\mathrm{NE}}$ and $p_{\mathrm{ER}}$ pressures is rather simple. In thermodynamic equilibrium, the free energy of the system reaches its minimum. As a result, in the case of positive $\sigma_{\mathrm{ER}}$ the system tries to reduce $G_{\mathrm{ER}}$ free energy by contracting the surface area of ER via expansion of the NE, – a process, which, from a physical point of view, can be described by an effective positive pressure ($p_{\mathrm{ER}} > 0$) acting on the NE. On the other hand, in the case of $\sigma_{\mathrm{ER}} < 0$, ER tries to maximize its surface area at the expense of the NE, thus effectively creating a negative pressure ($p_{\mathrm{ER}} < 0$) on the NE. Very similar arguments apply to $\sigma_{\mathrm{NE}}$ surface tension as well with the only difference being the opposite sign of the resulting effects on the NE.

It should be noted that values and signs of the both surface tensions, $\sigma_{\mathrm{NE}}$ and $\sigma_{\mathrm{ER}}$, in the general case depend on the lipid compositions of the NE and ER as well as the presence of proteins interacting with them. For example, from experiments it is known that lamins polymerize into a dense nuclear lamina network inlaying the inner side of the NE in metazoan cells [49, 50], resulting in decrease of $G_{\mathrm{NE}}$ free energy. Namely, upon binding to the NE each lamin molecule reduces $G_{\mathrm{NE}}$ free energy by $\Delta G_{\mathrm{NE}} = k_{\mathrm{B}}T\ln(c_{\mathrm{lamin}}/K_d)$, where $c_{\mathrm{lamin}}$ is the nucleoplasmic concentration of lamins and $K_d$ is the equilibrium dissociation constant of lamins from the nuclear lamina network.

From the above notes, it is clear that in the case of a sufficiently high nucleoplasmic concentration of lamins, $\sigma_{\mathrm{NE}}$ will generally have a negative value. According to Eq. (K9), this will result in appearance of positive pressure $p_{\mathrm{NE}}$ ($p_{\mathrm{NE}} > 0$) applied to the NE, indicating that lamin polymerization into the nuclear lamina network promotes expansion of the NE by performing work against other pressures, such as $p_{\mathrm{ion}}$, $p_{\mathrm{macro}}$ and $p_{\mathrm{ER}}$, contributing to the mechanical equilibrium of the cell nucleus. Indeed, it has been found that immunodepletion of lamins or downregulation of their expression in living cells lead to decrease in the nucleus size in a wide variety of eukaryotic cells, with a very similar effect taking place in the case of inhibition of the nuclear lamina assembly, suggesting that the nuclear lamina network counteracts the outside pressure created by the rest of cell components on the NE, see ref. [51–54].

Similarly, proteins binding to the ER membrane tend to decrease $\sigma_{\mathrm{ER}}$ surface tension, which at sufficiently high concentrations of the proteins may attain a negative value, resulting in a negative pressure, $p_{\mathrm{ER}}$ ($p_{\mathrm{ER}} < 0$), acting on the NE. Thus, NE- and ER-binding proteins generate pressures of opposite signs on the NE, leading to a tug-of-war between the NE and ER for the shared lipid membrane.

Anyway, by having formulas for calculation of the pressures generated by various cell components on the NE, it is then straightforward to find the resulting equilibrium size of the cell nucleus by employing Eq. (6) as discussed in the main text.

##### Appendix L: Electrostatic DNA-DNA interactions vs the volume-exclusion effect

In this Appendix section, we would like to find out the energy contributions made by electrostatic forces and the volume-exclusion effect into DNA-DNA interactions in order to understand their potential roles in DNA organization. To this aim, we are going to consider a viral particle scenario, for which analytic energy estimations can be easily obtained. As for the case of living cells, – it is not very hard to get very similar results for this scenario as well, however, due to complexity of the problem, one has to rely on numeric calculations in this situation rather than analytic formulas to calculate the energies of the electrostatic and volume-exclusion DNA-DNA interactions. For this reason, we will mainly focus here on a more informative example of a viral particle.

To begin with, let's start by recalling that in the absence of the volume-exclusion effect the partition function of DNA confined inside a viral capsid is described by the following formula, which is an analogue of Eq. (A5):

$$Z = \prod_j \left[ \frac{1}{n_j!} \prod_{k=1}^{n_j} \int_{V_{\rm tot}} \frac{Z_j^{\rm in} {\rm d}\mathbf{r}_{jk}}{\Lambda_j^3} \right] \int_{\mathbb{R}^3} {\rm d}\mathbf{r}_0 \int \mathscr{D}\mathbf{R}\, e^{-\beta E[\mathbf{R}] - \beta E_{\rm e}[\mathbf{R}, \mathbf{r}_{jk}]} \tag{L1}$$

Where $V_{\rm tot}$ is the total volume of the system. As for the rest of variables, they have absolutely the same meaning as in Eq. (A5) with the only difference being that we consider here a single viral DNA molecule instead of several DNA polymers. Finally, $E_{\rm e}[\mathbf{R}, \mathbf{r}_{jk}]$ in the above equation is the electrostatic interaction energy between the system components, including DNA and charged ions diffusing in surrounding solution:

$$E_{\rm e}[\mathbf{R}, \mathbf{r}_{jk}] = \frac{1}{2} \int_{\mathbb{R}^3} {\rm d}\mathbf{r} \int_{\mathbb{R}^3} {\rm d}\mathbf{r}'\, \rho_{\rm e}(\mathbf{r}) U_{\rm e}(\mathbf{r} - \mathbf{r}') \rho_{\rm e}(\mathbf{r}') \tag{L2}$$

Where $U_{\rm e}(\mathbf{r}) = \frac{1}{4\pi\varepsilon r}$ is the core part of the electrostatic potential and $\rho_{\rm e}(\mathbf{r})$ is the electrical charge density:

$$\rho_{\rm e}(\mathbf{r}) = \sum_j \sum_{k=1}^{n_j} q_j \delta(\mathbf{r} - \mathbf{r}_{jk}) + \rho_{\rm d} \int_0^L \delta(\mathbf{r} - \mathbf{r}_{\rm d}(s))\, {\rm d}s \tag{L3}$$

Here, as before, $\rho_{\rm d}$ and $L$ are the linear charge density and the length of the DNA molecule, respectively.

To introduce the volume-exclusion effect into the DNA partition function described by Eq. (L1), we need to define first the spatial density of DNA segments, $\rho_{\rm m}(\mathbf{r})$, which is also frequently referred to as the density of the polymer monomers:

$$\rho_{\rm m}(\mathbf{r}) = \frac{1}{b} \int_0^L \delta(\mathbf{r} - \mathbf{r}_{\rm d}(s))\, {\rm d}s \tag{L4}$$

Where $b$ is the size of DNA segments.

It then can be shown that DNA-DNA interaction energy associated with the volume-exclusion effect equals to (see [5, 34, 42, 55]):

$$E_{\rm v}[\mathbf{R}] = \frac{1}{2} \int_{\mathbb{R}^3} {\rm d}\mathbf{r} \int_{\mathbb{R}^3} {\rm d}\mathbf{r}'\, \rho_{\rm m}(\mathbf{r}) U_{\rm v}(\mathbf{r} - \mathbf{r}') \rho_{\rm m}(\mathbf{r}') = \frac{v}{2\beta} \int_{\mathbb{R}^3} \rho_{\rm m}^2(\mathbf{r})\, {\rm d}\mathbf{r} \tag{L5}$$

Where $v$ is the exclusion volume of a single DNA segment and $\beta = 1/k_{\rm B}T$ is the inverse thermodynamic temperature. As for the volume-exclusion potential, $U_{\rm v}(\mathbf{r})$, it is simply proportional to the Dirac $\delta$-function: $U_{\rm v}(\mathbf{r}) = \frac{v}{\beta}\delta(\mathbf{r})$.

By adding $E_{\rm v}[\mathbf{R}]$ energy term into Eq. (L1), one can obtain the following formula for the partition function, which accounts for both electrostatic and volume-exclusion interactions between the system components:

$$Z=\prod_j\left[\frac{1}{n_j!}\prod_{k=1}^{n_j}\int_{V_{\rm tot}}\frac{Z_j^{\rm in}\mathrm{d}\mathbf{r}_{jk}}{\Lambda_j^3}\right]\int_{\rm I\!R^3}\mathrm{d}\mathbf{r}_0\int\mathscr{D}\mathbf{R}\,e^{-\beta E[\mathbf{R}]-\beta E_{\rm e}[\mathbf{R},\mathbf{r}_{jk}]-\beta E_{\rm v}[\mathbf{R}]} \tag{L6}$$

In order to find the relative contributions of electrostatic forces and the volume-exclusion effect into DNA-DNA interactions based on Eq. (L6), we can follow the same derivation as in Appendix A by introducing $S_\psi[\mathbf{R},\mathbf{r}_{jk}]$ and $S_\Xi[\mathbf{R}]$ physical actions associated with randomly fluctuating electrostatic field, $\psi$, and volume-exclusion field, $\Xi$:

$$\begin{aligned} S_\psi[\mathbf{R},\mathbf{r}_{jk}]&=\int_{\rm I\!R^3}\mathrm{d}\mathbf{r}\,\rho_{\rm e}(\mathbf{r})\,\psi(\mathbf{r})-\frac{1}{2}\int_{\rm I\!R^3}\mathrm{d}\mathbf{r}\int_{\rm I\!R^3}\mathrm{d}\mathbf{r}'\,\psi(\mathbf{r})\,U_{\rm e}^{-1}(\mathbf{r}-\mathbf{r}')\,\psi(\mathbf{r}')\\ S_\Xi[\mathbf{R}]&=\int_{\rm I\!R^3}\mathrm{d}\mathbf{r}\,\rho_{\rm m}(\mathbf{r})\,\Xi(\mathbf{r})-\frac{1}{2}\int_{\rm I\!R^3}\mathrm{d}\mathbf{r}\int_{\rm I\!R^3}\mathrm{d}\mathbf{r}'\,\Xi(\mathbf{r})\,U_{\rm v}^{-1}(\mathbf{r}-\mathbf{r}')\,\Xi(\mathbf{r}')=\int_{\rm I\!R^3}\mathrm{d}\mathbf{r}\,\rho_{\rm m}(\mathbf{r})\,\Xi(\mathbf{r})-\frac{\beta}{2v}\int_{\rm I\!R^3}\mathrm{d}\mathbf{r}\,\Xi^2(\mathbf{r}) \end{aligned} \tag{L7}$$

Where $U_{\rm e}^{-1}(\mathbf{r})=-\varepsilon\Delta\delta(\mathbf{r})$ and $U_{\rm v}^{-1}(\mathbf{r})=\frac{\beta}{v}\delta(\mathbf{r})$ are functional inverse of $U_{\rm e}(\mathbf{r})$ and $U_{\rm v}(\mathbf{r})$ potentials, see Eq. (A7) and Eq. (A27).

Indeed, by using Eq. (L2) and Eq. (L7), it is not hard to obtain the next form of Eq. (A8):

$$E_{\rm e}[\mathbf{R},\mathbf{r}_{jk}]=\frac{1}{2}\int_{\rm I\!R^3}\mathrm{d}\mathbf{r}\int_{\rm I\!R^3}\mathrm{d}\mathbf{r}'\,\rho_{\rm e}(\mathbf{r})U_{\rm e}(\mathbf{r}-\mathbf{r}')\rho_{\rm e}(\mathbf{r}')=S_\psi[\mathbf{R},\mathbf{r}_{jk}]+\frac{1}{2}\int_{\rm I\!R^3}\mathrm{d}\mathbf{r}\int_{\rm I\!R^3}\mathrm{d}\mathbf{r}'\,\widetilde{\psi}(\mathbf{r})\,U_{\rm e}^{-1}(\mathbf{r}-\mathbf{r}')\,\widetilde{\psi}(\mathbf{r}') \tag{L8}$$

Where, as before, $\widetilde{\psi}(\mathbf{r})$ is a shifted electrostatic potential defined by Eq. (A9).

Similarly, from Eq. (L5) and Eq. (L7), we get:

$$\begin{aligned} E_{\rm v}[\mathbf{R}]&=\frac{1}{2}\int_{\rm I\!R^3}\mathrm{d}\mathbf{r}\int_{\rm I\!R^3}\mathrm{d}\mathbf{r}'\,\rho_{\rm m}(\mathbf{r})U_{\rm v}(\mathbf{r}-\mathbf{r}')\rho_{\rm m}(\mathbf{r}')=\frac{v}{2\beta}\int_{\rm I\!R^3}\mathrm{d}\mathbf{r}\,\rho_{\rm m}^2(\mathbf{r})=\\ &=S_\Xi[\mathbf{R}]+\frac{1}{2}\int_{\rm I\!R^3}\mathrm{d}\mathbf{r}\int_{\rm I\!R^3}\mathrm{d}\mathbf{r}'\,\widetilde{\Xi}(\mathbf{r})\,U_{\rm v}^{-1}(\mathbf{r}-\mathbf{r}')\,\widetilde{\Xi}(\mathbf{r}')=S_\Xi[\mathbf{R}]+\frac{\beta}{2v}\int_{\rm I\!R^3}\mathrm{d}\mathbf{r}\,\widetilde{\Xi}^2(\mathbf{r}) \end{aligned} \tag{L9}$$

Where $\widetilde{\Xi}(\mathbf{r})$ is a shifted volume-exclusion field:

$$\widetilde{\Xi}(\mathbf{r})=\Xi(\mathbf{r})-\int_{\rm I\!R^3}\mathrm{d}\mathbf{r}'\,U_{\rm v}(\mathbf{r}-\mathbf{r}')\,\rho_{\rm m}(\mathbf{r}')=\Xi(\mathbf{r})-\frac{v}{\beta}\rho_{\rm m}(\mathbf{r}) \tag{L10}$$

By multiplying Eq. (L8) by $-\beta$, exponentiating it and applying $\int\mathscr{D}\psi$ field integral to the both sides, it is then easy to arrive to the following mathematical expression:

$$\begin{aligned} \int e^{-\beta S_\psi[\mathbf{R},\mathbf{r}_{jk}]}\mathscr{D}\psi&=e^{-\frac{\beta}{2}\int_{\rm I\!R^3}\mathrm{d}\mathbf{r}\int_{\rm I\!R^3}\mathrm{d}\mathbf{r}'\,\rho_{\rm e}(\mathbf{r})U_{\rm e}(\mathbf{r}-\mathbf{r}')\rho_{\rm e}(\mathbf{r}')}\int e^{\frac{\beta}{2}\int_{\rm I\!R^3}\mathrm{d}\mathbf{r}\int_{\rm I\!R^3}\mathrm{d}\mathbf{r}'\,\widetilde{\psi}(\mathbf{r})\,U_{\rm e}^{-1}(\mathbf{r}-\mathbf{r}')\,\widetilde{\psi}(\mathbf{r}')}\mathscr{D}\widetilde{\psi}=\\ &=\mathrm{const}\times e^{-\frac{\beta}{2}\int_{\rm I\!R^3}\mathrm{d}\mathbf{r}\int_{\rm I\!R^3}\mathrm{d}\mathbf{r}'\,\rho_{\rm e}(\mathbf{r})U_{\rm e}(\mathbf{r}-\mathbf{r}')\rho_{\rm e}(\mathbf{r}')}=\mathrm{const}\times e^{-\beta E_{\rm e}[\mathbf{R},\mathbf{r}_{jk}]} \end{aligned} \tag{L11}$$

Here we have taken into account that $\int\mathscr{D}\widetilde{\psi}=\int\mathscr{D}\psi$, where $\int\mathscr{D}\psi$ and $\int\mathscr{D}\widetilde{\psi}$ are field integrals over all possible configurations of the electrostatic potentials $\psi$ and $\widetilde{\psi}$, respectively.

Similarly, from Eq. (L9) we have:

$$\int e^{-\beta S_\Xi[\mathbf{R}]}\mathscr{D}\Xi=e^{-\frac{v}{2}\int_{\rm I\!R^3}\mathrm{d}\mathbf{r}\,\rho_{\rm m}^2(\mathbf{r})}\int e^{\frac{\beta^2}{2v}\int_{\rm I\!R^3}\mathrm{d}\mathbf{r}\,\widetilde{\Xi}^2(\mathbf{r})}\mathscr{D}\widetilde{\Xi}=\mathrm{const}\times e^{-\frac{v}{2}\int_{\rm I\!R^3}\mathrm{d}\mathbf{r}\,\rho_{\rm m}^2(\mathbf{r})}=\mathrm{const}\times e^{-\beta E_{\rm v}[\mathbf{R}]} \tag{L12}$$

Where again it was taken into account that $\int\mathscr{D}\widetilde{\Xi}=\int\mathscr{D}\Xi$.

Substituting Eq. (L7), Eq. (L11) and Eq. (L12) into Eq. (L6), and ignoring the constant prefactor, it can be seen that the formula for the partition function turns into:

$$Z = \int \mathscr{D}\psi \int \mathscr{D}\Xi \prod_j \left[ \frac{1}{n_j!} \prod_{k=1}^{n_j} \int_{V_{\text{tot}}} \frac{Z_j^{\text{in}} \mathrm{d}\mathbf{r}_{jk}}{\Lambda_j^3} \right] \int_{\mathrm{I\!R}^3} \mathrm{d}\mathbf{r}_0 \int \mathscr{D}\mathbf{R}\, e^{-\beta E[\mathbf{R}] - \beta S_\psi[\mathbf{R}, \mathbf{r}_{jk}] - \beta S_\Xi[\mathbf{R}]} =$$
$$= \int \mathscr{D}\psi \int \mathscr{D}\Xi \prod_j \frac{(Z_j^\psi)^{n_j}}{n_j!} \times Z_{\psi\Xi}\, e^{\frac{\beta^2}{2v} \int_{\mathrm{I\!R}^3} \mathrm{d}\mathbf{r}\, \Xi^2(\mathbf{r}) + \frac{\beta}{2} \int_{\mathrm{I\!R}^3} \mathrm{d}\mathbf{r} \int_{\mathrm{I\!R}^3} \mathrm{d}\mathbf{r}'\, \psi(\mathbf{r})\, U_{\mathrm{e}}^{-1}(\mathbf{r} - \mathbf{r}')\, \psi(\mathbf{r}')} \quad \text{(L13)}$$

Here $Z_j^\psi$ is the partition function of an individual particle of type $j$ in the field $\psi$ described by Eq. (A15), where $V_j = V_{\text{tot}}$ for all types of ions, $j$, since based on experimental findings it is assumed in the model that ions can freely shuttle between the buffer solution and the inner space of the viral capsid, see Appendix A for details. As for $Z_{\psi\Xi}$, this is the partition function of DNA being a subject to action of the both fields, $\psi$ and $\Xi$:

$$Z_{\psi\Xi} = \int_{\mathrm{I\!R}^3} \mathrm{d}\mathbf{r}_0 \int \mathscr{D}\mathbf{R}\, e^{-\beta \left\{ E[\mathbf{R}] + \rho_{\mathrm{d}} \int_0^L \psi(\mathbf{r}_{\mathrm{d}}(s))\, \mathrm{d}s + \frac{1}{b} \int_0^L \Xi(\mathbf{r}_{\mathrm{d}}(s))\, \mathrm{d}s \right\}} \quad \text{(L14)}$$

Following Appendix A, it should be noted that the value of the double field integral in Eq. (L13) is mainly determined by the system behaviour near the stationary phase of the integrand. Thus, by applying the stationary phase approximation, for the partition function, $Z$, we have:

$$Z \propto \prod_j \left( \frac{Z_j^{\psi_{\text{sp}}}}{n_j} \right)^{n_j} \times Z_{\psi_{\text{sp}}\Xi_{\text{sp}}}\, e^{\frac{\beta^2}{2v} \int_{\mathrm{I\!R}^3} \mathrm{d}\mathbf{r}\, \Xi_{\text{sp}}^2(\mathbf{r}) + \frac{\beta}{2} \int_{\mathrm{I\!R}^3} \mathrm{d}\mathbf{r} \int_{\mathrm{I\!R}^3} \mathrm{d}\mathbf{r}'\, \psi_{\text{sp}}(\mathbf{r})\, U_{\mathrm{e}}^{-1}(\mathbf{r} - \mathbf{r}')\, \psi_{\text{sp}}(\mathbf{r}')} \quad \text{(L15)}$$

Where the partition function is defined up to a non-essential multiplicative constant, whose exact value is not important since it results only in a constant offset of the system free energy that does not affect any physical characteristic of the system. In the above formula, $\psi_{\text{sp}}$ and $\Xi_{\text{sp}}$ are potential fields that extremize the integrand function in Eq. (L13), which can be found by solving the following system of equations:

$$\begin{cases} \dfrac{\delta}{\delta\psi} \left[ \displaystyle\sum_j n_j \ln Z_j^\psi + \ln Z_{\psi\Xi} + \frac{\beta}{2} \int_{\mathrm{I\!R}^3} \mathrm{d}\mathbf{r} \int_{\mathrm{I\!R}^3} \mathrm{d}\mathbf{r}'\, \psi(\mathbf{r}) U_{\mathrm{e}}^{-1}(\mathbf{r} - \mathbf{r}') \psi(\mathbf{r}') \right] = 0 \\ \dfrac{\delta}{\delta\Xi} \left[ \ln Z_{\psi\Xi} + \dfrac{\beta^2}{2v} \displaystyle\int_{\mathrm{I\!R}^3} \mathrm{d}\mathbf{r}\, \Xi^2(\mathbf{r}) \right] = 0 \end{cases} \quad \text{(L16)}$$

Here $\sum_j$ is the sum over all types of particles, $j$, diffusing in solution, and $\frac{\delta}{\delta\psi}$ and $\frac{\delta}{\delta\Xi}$ are functional derivatives with respect to the corresponding potential fields, $\psi$ and $\Xi$. Since the system possesses a spherical symmetry, it is clear that both fields, $\psi$ and $\Xi$, in the general case will depend only on the radial distance, $r = \|\mathbf{r}\|$, measured from the center of the viral capsid. As a result, it can be shown that $\frac{\delta}{\delta\psi}$ and $\frac{\delta}{\delta\Xi}$ functional derivatives can be taken with respect to the radial distance: $\frac{\delta}{\delta\psi} = \frac{\delta}{\delta\psi(r)}$ and $\frac{\delta}{\delta\Xi} = \frac{\delta}{\delta\Xi(r)}$, with the final result being absolutely the same as in the case when the functional derivatives are taken with respect to the position vector, $\mathbf{r}$.

Let's now consider these functional derivatives one by one. As has been previously demonstrated in Appendix A, the first line of Eq. (L16) can be reduced to [see Eq. (A32)]:

$$\frac{1}{4\pi\beta r^2} \frac{\delta \ln Z_{\psi\Xi}}{\delta\psi(r)} - \sum_j c_j q_j e^{-\beta q_j \psi(r)} - \varepsilon \Delta\psi(r) = 0 \quad \text{(L17)}$$

Furthermore, from Eq. (E5), which can be shown to hold in the case of the DNA partition function defined by

Eq. (L14), we have:

$$\frac{\delta \ln Z_{\psi\Xi}}{\delta\psi(r)} \approx \frac{N_{\rm tot}}{\lambda_{\rm max}} \frac{\Pr^{\rm L}_{\lambda_{\rm max}} \mathbf{U} \times \frac{\delta}{\delta\psi(r)} \mathbf{L} \times \Pr^{\rm R}_{\lambda_{\rm max}} \mathbf{Y}}{\Pr^{\rm L}_{\lambda_{\rm max}} \mathbf{U} \times \Pr^{\rm R}_{\lambda_{\rm max}} \mathbf{Y}} = \frac{N_{\rm tot}}{\lambda_{\rm max}} \frac{\Pr^{\rm L}_{\lambda_{\rm max}} \mathbf{U} \times \frac{\delta}{\delta\psi(r)} \mathbf{T}^{(6)}_{00} \times \Pr^{\rm R}_{\lambda_{\rm max}} \mathbf{Y}}{\Pr^{\rm L}_{\lambda_{\rm max}} \mathbf{U} \times \Pr^{\rm R}_{\lambda_{\rm max}} \mathbf{Y}} \tag{L18}$$

Where the transfer-matrix $\mathbf{L}$ takes a very simple form: $\mathbf{L} = \mathbf{T}^{(6)}_{00}$, with the matrix $\mathbf{T}^{(6)}_{00}$ being defined by the same Eq. (D11) and Eq. (G41). The only major difference here is that we have to take into account the volume-exclusion field, $\Xi$, in definition of $\Phi_{00}$ function in addition to the electrostatic potential, $\psi$, and, in fact, it is not hard to see from Eq. (L14) and Eq. (E8) that:

$$\begin{aligned}
\Phi_{00}\big(\mathbf{G}_{\mathbf{m}_j}\big) &= \frac{1}{V_{\rm B}} \int_{V_{\rm vir}} {\rm d}\mathbf{r}'_j \, e^{-\beta b \rho_{\rm d} \psi(\mathbf{r}'_j) - \beta \Xi(\mathbf{r}'_j)} \, e^{-i[\mathbf{G}_{\mathbf{m}_j} \cdot \mathbf{r}'_j]} = \\
&= \frac{1}{V_{\rm B}} \int_0^{2\pi} {\rm d}\varphi'_j \int_0^{\pi} \sin\theta'_j \, {\rm d}\theta'_j \int_0^{R_{\rm vir}} \big(r'_j\big)^2 e^{-\beta b \rho_{\rm d} \psi(r'_j) - \beta \Xi(r'_j)} \, e^{-i G_{\mathbf{m}_j} r'_j \cos\theta'_j} \, {\rm d}r'_j = \Big[u = \cos\theta'_j\Big] = \\
&= \frac{2\pi}{V_{\rm B}} \int_{-1}^{1} {\rm d}u \int_0^{R_{\rm vir}} \big(r'_j\big)^2 e^{-\beta b \rho_{\rm d} \psi(r'_j) - \beta \Xi(r'_j)} \, e^{-i G_{\mathbf{m}_j} r'_j u} \, {\rm d}r'_j = \frac{4\pi}{V_{\rm B} G_{\mathbf{m}_j}} \int_0^{R_{\rm vir}} r'_j \, e^{-\beta b \rho_{\rm d} \psi(r'_j) - \beta \Xi(r'_j)} \times \\
&\times \sin\big(G_{\mathbf{m}_j} r'_j\big) \, {\rm d}r'_j = \frac{4\pi}{V_{\rm B}} \int_0^{R_{\rm vir}} \big(r'_j\big)^2 e^{-\beta b \rho_{\rm d} \psi(r'_j) - \beta \Xi(r'_j)} \, j_0\big(G_{\mathbf{m}_j} r'_j\big) \, {\rm d}r'_j
\end{aligned} \tag{L19}$$

Here $R_{\rm vir}$ and $V_{\rm vir}$ are the radius and the volume of the viral particle, and $b$ is the length of DNA segments.

As for $\frac{\delta}{\delta\Xi}$ functional derivative from Eq. (L16), it is clear that it equals to:

$$\frac{\delta \ln Z_{\psi\Xi}}{\delta\Xi(r)} + \frac{4\pi\beta^2 r^2}{v} \Xi(r) = 0 \tag{L20}$$

Where for $\frac{\delta \ln Z_{\psi\Xi}}{\delta\Xi(r)}$ derivative we have:

$$\frac{\delta \ln Z_{\psi\Xi}}{\delta\Xi(r)} \approx \frac{N_{\rm tot}}{\lambda_{\rm max}} \frac{\Pr^{\rm L}_{\lambda_{\rm max}} \mathbf{U} \times \frac{\delta}{\delta\Xi(r)} \mathbf{T}^{(6)}_{00} \times \Pr^{\rm R}_{\lambda_{\rm max}} \mathbf{Y}}{\Pr^{\rm L}_{\lambda_{\rm max}} \mathbf{U} \times \Pr^{\rm R}_{\lambda_{\rm max}} \mathbf{Y}} \tag{L21}$$

Now, from Eq. (L19) it follows that:

$$\frac{\delta\Phi_{00}(\mathbf{G}_{\mathbf{m}_j})}{\delta\psi(r)} = \begin{cases} -\dfrac{4\pi}{V_{\rm B}} \beta b \rho_{\rm d} r^2 \, e^{-\beta b \rho_{\rm d} \psi(r) - \beta \Xi(r)} j_0(r G_{\mathbf{m}_j}), & \text{if } r < R_{\rm vir} \\ 0 & \text{, otherwise} \end{cases} \tag{L22}$$

On the other hand, for $\frac{\delta\Phi_{00}}{\delta\Xi(r)}$ derivative we have:

$$\frac{\delta\Phi_{00}(\mathbf{G}_{\mathbf{m}_j})}{\delta\Xi(r)} = \begin{cases} -\dfrac{4\pi}{V_{\rm B}} \beta r^2 \, e^{-\beta b \rho_{\rm d} \psi(r) - \beta \Xi(r)} j_0(r G_{\mathbf{m}_j}), & \text{if } r < R_{\rm vir} \\ 0 & \text{, otherwise} \end{cases} \tag{L23}$$

Combining Eq. (L18) and Eq. (L21)-Eq. (L23), it becomes obvious that:

$$\frac{\delta \ln Z_{\psi\Xi}}{\delta\Xi(r)} = \frac{1}{b\rho_{\rm d}} \frac{\delta \ln Z_{\psi\Xi}}{\delta\psi(r)} \tag{L24}$$

Thus, from Eq. (L17), Eq. (L20) and Eq. (L24), it can be concluded that Eq. (L16) for the stationary phase fields,

$\psi_{\rm sp}$ and $\Xi_{\rm sp}$, is equivalent to the following system of equations:

$$\begin{cases} \dfrac{\delta \ln Z_{\psi\Xi}}{\delta\psi(r)} - 4\pi\beta r^2 \left[ \sum_j c_j q_j e^{-\beta q_j \psi(r)} + \varepsilon\Delta\psi(r) \right] = 0 \\ \dfrac{\delta \ln Z_{\psi\Xi}}{\delta\psi(r)} + \dfrac{4\pi\beta^2 b\rho_{\rm d} r^2}{v}\Xi(r) = 0 \end{cases} \tag{L25}$$

which can be further simplified by subtracting the first line of the equation from the second one:

$$\begin{cases} \dfrac{\delta \ln Z_{\psi\Xi}}{\delta\psi(r)} - 4\pi\beta r^2 \left[ \sum_j c_j q_j e^{-\beta q_j \psi(r)} + \varepsilon\Delta\psi(r) \right] = 0 \\ \Xi(r) = -\dfrac{v}{\beta b\rho_{\rm d}} \left[ \sum_j c_j q_j e^{-\beta q_j \psi(r)} + \varepsilon\Delta\psi(r) \right] \end{cases} \tag{L26}$$

From Eq. (L26) it can be seen that there exists a simple mathematical relation between the stationary phase fields, $\psi_{\rm sp}$ and $\Xi_{\rm sp}$, which makes it possible to easily obtain estimations of the electrostatic and volume-exclusion DNA interaction energies.

To this aim, it should be noted that the model calculations indicate that in the absence of the volume-exclusion effect, the stationary phase electrostatic potential, $\psi_{\rm sp}(r)$, is nearly uniform inside the viral capsid, see Figure 2(d). Addition of the volume-exclusion effect to the model does not change this trend as it even further helps DNA to uniformly spread throughout the inner space of the capsid. Thus, to the first order of approximation, we can safely assume that $\psi_{\rm sp}(r)$ and $\Xi_{\rm sp}(r)$ fields are nearly constant inside the viral particle and neglect $\varepsilon\Delta\psi(r)$ term in our estimations of the DNA-DNA electrostatic and volume-exclusion interaction energies.

Furthermore, following Appendix A, it is clear that $\sum_j$ sum in Eq. (L26) is dominated by monovalent ions, such as $Na^+$ and $Cl^-$, whose total concentration in the surrounding media under physiological conditions is equal to $c_{\rm ions} = 150$ mM for both positive and negative ions [8].

From the above notes and Eq. (L26), it is then not very hard to obtain the following approximate formula for the average value of the stationary phase volume-exclusion field, $\hat{\Xi}_{\rm sp}$:

$$\hat{\Xi}_{\rm sp} \approx -\frac{q_{\rm e} c_{\rm ions} v}{\beta b\rho_{\rm d}} \left[ e^{-\beta q_{\rm e}\hat{\psi}_{\rm sp}} - e^{\beta q_{\rm e}\hat{\psi}_{\rm sp}} \right] = \frac{2 q_{\rm e} c_{\rm ions} v}{\beta b\rho_{\rm d}} \sinh(\beta q_{\rm e}\hat{\psi}_{\rm sp}) \tag{L27}$$

Where $\hat{\psi}_{\rm sp}$ is the average stationary phase electrostatic potential inside the viral particle, $q_{\rm e}$ is the elementary charge and $\sinh(x)$ is the hyperbolic sine function.

By using Eq. (L27) and $v \approx b^2 R_{\rm d}$ formula for the excluded volume of DNA segments [42], where $R_{\rm d} = 1$ nm is the radius of DNA, one then can easily find the interaction energies, $E'_{\rm e}$ and $E'_{\rm v}$, of a single DNA segment of the length $b$ with the mean stationary phase fields, $\hat{\psi}_{\rm sp}$ and $\hat{\Xi}_{\rm sp}$, as: $E'_{\rm e} \approx b\rho_{\rm d}\hat{\psi}_{\rm sp}$ and $E'_{\rm v} \approx \hat{\Xi}_{\rm sp}$, see the exponential factor in Eq. (L19).

Plotting $E'_{\rm e}$ and $E'_{\rm v}$ energies as functions of the average $\hat{\psi}_{\rm sp}$ potential, it can be seen that in the physiologically relevant range of $-100$ mV $\leq \hat{\psi}_{\rm sp} \leq 0$ mV, DNA interaction with the surrounding electrostatic field created by electrically charged particles, including DNA itself, is by more than two orders of magnitude stronger than DNA-DNA interactions due to the volume-exclusion effect, see Figure S14. Only in the rightmost part of the graph corresponding to the physical conditions found in small viral particles the volume-exclusion effect starts to catch up with electrostatic DNA interactions due to distant DNA segments coming into direct physical contact with each other. Yet, even in the most extreme case of $\hat{\psi}_{\rm sp} = -100$ mV, the energy contribution of the volume-exclusion effect remains small in comparison to the one made by electrostatic forces, indicating that the latter play a dominant role in determining the DNA conformation.

*In vitro* experimental data support this theoretical observation by showing that the effective diameter of DNA is very sensitive to the ionic strength of the surrounding solution [56], pointing to the fact that DNA-DNA interactions are mainly governed by repulsive electrostatic forces rather than the volume-exclusion effect.

It should be noted that while in this Appendix section we have considered only the case of a viral particle, from Figure S14 it is clear that very similar results will also hold for chromosomal DNA packed inside the nucleus of a eukaryotic cell. Indeed, even if the excluded volume of nucleosomes and partial electrostatic screening of DNA by histone octamers (by $\sim 50\%$) are taken into account, DNA-DNA electrostatic repulsion forces still will be several times stronger than the volume-exclusion effect in the physiologically relevant range of $-10$ mV $\leq \hat{\psi}_{\rm sp} \leq 0$ mV. As a result, the contribution of the volume-exclusion effect into the DNA partition function can be omitted in the model calculations, as it was done in Appendix A.

Finally, it is worth mentioning that formulas such as Eq. (L1)-Eq. (L26) not only can be used to add the volume-exclusion effect as a higher-order correction to the model in future studies, but also to introduce HP1-dependent nucleosome-nucleosome cross-linking interactions responsible for phase-separation processes taking place in nuclei of living cells [57, 58]. This may provide further insights into molecular mechanisms responsible for chromatin organization in living cells, see *Discussion* section in the main text.

#### Appendix M: Nuclear transport

In the previous Appendix sections, it has been assumed that processes responsible for chromatin organization in nuclei of living cells can be rather well described by equilibrium thermodynamics. However, it is known that living cells usually operate quite far from thermodynamic equilibrium. For example, to overcome the energy barrier imposed by nuclear pore complexes (NPCs) on movement of large proteins ($> 50$ kDa) across the NE, eukaryotic cells utilize so-called nuclear transport system that relies on active physico-chemical processes that utilize the energy of GTP hydrolysis [59]. As a result, the question arises: to what extent such active processes may affect the main results presented in our study?

In this section, we address this question by considering nuclear transportation of histone-binding chaperones shuttling between the cell cytosol and nucleoplasm. To this aim, we utilize a minimal pseudo-first order kinetic model similar to that published in a recent study [60], which has been originally used by the authors to estimate the passive and active transportation rates of proteins through NPCs. This will make it possible to utilize the transportation rates measured in ref. [60] to evaluate corrections that need to be done to our model of DNA packaging in nuclei of living cells in order to accurately take into account the effect of the nuclear transportation system.

Specifically, following ref. [60], we will assume that there exist two types of processes by which histone-bound and unloaded chaperones, $\rm c_1$ and $\rm c_2$, can move between the cell cytosol and nucleoplasm: 1) via passive diffusion through NPCs or 2) by utilizing active nuclear transportation system mediated by importins and exportins that recognize nuclear localization (NLS) and nuclear export (NES) signals carried by cargo proteins, see schematic Figure S15(a). Indeed, it has been previously shown that histone-binding chaperones, such as Nap1, shuttle between the cell cytosol and nucleoplasm by utilizing both NLS and NES signals, making them targets for importin- and exportin-dependent nuclear transportation [19].

Experimental measurements performed in ref. [60] indicate that nuclear transportation processes can be rather well described by pseudo-first order kinetic rates, which in the general case may depend on the size of the transported protein and/or microenvironment conditions in which cells are grown [60]. By using this approach, it then straightforward to find the ratio between the cytosolic and nucleoplasmic concentrations of chaperones by imposing the steady-state condition on the nuclear import ($J_{\rm imp}$) and export ($J_{\rm exp}$) fluxes for each of the chaperone types: $J_{\rm imp} = J_{\rm exp}$. For example, for unloaded $c_1$ chaperone, we have:

$$J_{\rm imp} = (k^{\rm a}_{\rm imp} + k^{\rm p}_{\rm imp})\, c^{\rm cyto}_{\rm c_1 u} \quad \text{and} \quad J_{\rm exp} = (k^{\rm a}_{\rm exp} + k^{\rm p}_{\rm exp})\, c^{\rm nucl}_{\rm c_1 u} \quad \Longrightarrow \quad (k^{\rm a}_{\rm imp} + k^{\rm p}_{\rm imp})\, c^{\rm cyto}_{\rm c_1 u} = (k^{\rm a}_{\rm exp} + k^{\rm p}_{\rm exp})\, c^{\rm nucl}_{\rm c_1 u} \qquad \text{(M1)}$$

Where $k^{\rm a}_{\rm imp}$ and $k^{\rm a}_{\rm exp}$ are active import and export transportation rates, and $k^{\rm p}_{\rm imp}$ and $k^{\rm p}_{\rm exp}$ are passive import and export diffusion rates through NPCs, see Figure S15(a). $c^{\rm cyto}_{\rm c_1 u}$ and $c^{\rm nucl}_{\rm c_1 u}$ are cytosolic and nucleoplasmic concentrations of unloaded $\rm c_1$ chaperones, respectively. Very similar equations can be also written down for histone-bound chaperones,

$c_1$b and $c_2$b, as well as for unloaded $c_2$u chaperones.

From Eq. (M1), it can be seen that:

$$\frac{c_{c_1u}^{nucl}}{c_{c_1u}^{cyto}} = \frac{k_{imp}^{a} + k_{imp}^{p}}{k_{exp}^{a} + k_{exp}^{p}} \tag{M2}$$

To further simplify Eq. (M2), it is convenient to apply the theory of activated complex, from which it follows that:

$$\begin{cases} k_{imp}^{p} = k_{imp}^{p,0} \\ k_{imp}^{a} = k_{imp}^{a,0} \\ k_{exp}^{p} = k_{exp}^{p,0}\, e^{\beta q_{c_1u}\langle\psi\rangle} \\ k_{exp}^{a} = k_{exp}^{a,0}\, e^{\beta q_{c_1u}\langle\psi\rangle} \end{cases} \tag{M3}$$

Here $\langle\psi\rangle$ is the average nuclear electrostatic potential. $k_{imp}^{p,0}$, $k_{imp}^{a,0}$, $k_{exp}^{p,0}$ and $k_{exp}^{a,0}$ are transportation rates of a protein of the same size as $c_1$ chaperone, but with a net zero electrical charge. Indeed, in the presence of the nuclear electrostatic potential, the activation energies of the active and passive nuclear import remain unchanged. At the same time, the activation energies of the active and passive nuclear export gain $-q_{c_1u}\langle\psi\rangle$ offset due to the chaperone interaction with the nuclear electrostatic potential [Figure S15(b)], resulting in appearance of $e^{\beta q_{c_1u}\langle\psi\rangle}$ exponential factors in the right part of Eq. (M3).

Substituting Eq. (M3) into Eq. (M2), we get:

$$\frac{c_{c_1u}^{nucl}}{c_{c_1u}^{cyto}} = \frac{k_{imp}^{a,0} + k_{imp}^{p,0}}{k_{exp}^{a,0} + k_{exp}^{p,0}}\, e^{-\beta q_{c_1u}\langle\psi\rangle} = e^{\beta\eta - \beta q_{c_1u}\langle\psi\rangle} \tag{M4}$$

Where

$$\eta = k_B T \ln\left(\frac{k_{imp}^{a,0} + k_{imp}^{p,0}}{k_{exp}^{a,0} + k_{exp}^{p,0}}\right) \tag{M5}$$

Thus, in the presence of active nuclear transportation of histone-binding chaperones through NPCs, the ratio between the cytosolic and nucleoplasmic concentrations of chaperones deviates from that predicted by the Boltzmann law by $e^{\beta\eta}$ factor.

To estimate the value of the correction energy, $\eta$, it should be first noted that from the detailed balance condition it follows that $k_{exp}^{p,0} = k_{imp}^{p,0} = k_{imp}^{p}$. In addition, from Eq. (M3) it can be seen that regardless of the electrical charge of a transported protein, we have:

$$k_{exp}^{a,0} = \frac{k_{exp}^{a} k_{imp}^{p}}{k_{exp}^{p}} \tag{M6}$$

As a result, this makes it possible to estimate $\eta$ energy defined Eq. (M5) based on experimental measurements performed on any protein of a similar molecular weight. For example, it is possible to use experimental data shown in Figures 3B-C and S4J-K of ref. [60], from which it is not hard to find $k_{imp}^{p,0}$, $k_{imp}^{a,0}$, $k_{exp}^{p,0}$, $k_{exp}^{a,0}$ rates, and thus $\eta$ energy, for a protein of $27-67$ kDa size, covering practically the whole range of histone-binding chaperones and their complexes with histone dimers (for example, the molecular weight of Asf1 histone-binding chaperone is 32 kDa, Nap1 – 48 kDa). The final results of the calculations are shown in Table II.

As can be seen from the table, the absolute value of $\eta$ energy is $|\eta| < 0.6\, k_B T$ in the whole $27-67$ kDa range of protein sizes, suggesting that the nuclear transport has a rather negligible effect on localization of proteins carrying both NLS and NES signals, such as Nap1 histone-binding chaperones. As a result, based on Eq. (M4) it can be concluded that the distribution of histone-binding chaperones between the cell cytosol and nucleoplasm should very closely follow that predicted by the Boltzmann law, indicating that the equilibrium approach to description of chromatin organization in nuclei of living cells, which is adopted in our study, is well justified from a physical point of view. Since variation in the strength of NLS affinity to importins has almost the same effect on both import and export transportation rates of proteins (Figure S3E-F in ref. [60]), this conclusion remains valid regardless of the strength of NLS.

It should be noted that by making a few minor changes in the model, it is not very hard to take into account the deviation described by Eq. (M4) of the nucleocytoplasmic distribution of histone-binding chaperones from the Boltzmann law. Specifically, this can be done by using the following modified form of Eq. (A15) in model calculations:

$$Z^{\psi}_{\mathrm{c}_i\mathrm{u}} = \frac{Z^{\mathrm{in}}_{\mathrm{c}_i\mathrm{u}}}{\Lambda^3_{\mathrm{c}_i\mathrm{u}}} \int_{V^{\mathrm{osm}}_{\mathrm{cell}}} e^{\beta\eta_{\mathrm{c}_i\mathrm{u}}\chi_{\mathrm{nucl}}(\mathbf{r}) - \beta q_{\mathrm{c}_i\mathrm{u}}\psi(\mathbf{r})}\, d\mathbf{r} \quad \text{and} \quad Z^{\psi}_{\mathrm{c}_i\mathrm{b}} = \frac{Z^{\mathrm{in}}_{\mathrm{c}_i\mathrm{b}}}{\Lambda^3_{\mathrm{c}_i\mathrm{b}}} \int_{V^{\mathrm{osm}}_{\mathrm{cell}}} e^{\beta\eta_{\mathrm{c}_i\mathrm{b}}\chi_{\mathrm{nucl}}(\mathbf{r}) - \beta q_{\mathrm{c}_i\mathrm{b}}\psi(\mathbf{r})}\, d\mathbf{r} \tag{M7}$$

Where $Z^{\psi}_{\mathrm{c}_i\mathrm{u}}$ and $Z^{\psi}_{\mathrm{c}_i\mathrm{b}}$ are the partition functions of unloaded and histone-bound chaperones, $\mathrm{c}_i$ $(i = 1, 2)$, in the presence of the nuclear electrostatic potential, $\psi(\mathbf{r})$. $Z^{\mathrm{in}}_{\mathrm{c}_i\mathrm{u}}$ and $Z^{\mathrm{in}}_{\mathrm{c}_i\mathrm{b}}$ are the partition function parts corresponding to the inner degrees of freedom of the corresponding chaperones. $\Lambda_{\mathrm{c}_i\mathrm{u}}$, $\Lambda_{\mathrm{c}_i\mathrm{b}}$, $q_{\mathrm{c}_i\mathrm{u}}$ and $q_{\mathrm{c}_i\mathrm{b}}$ are the thermal de Broglie wavelengths and electrical charges of unloaded and histone-bound chaperones, $\mathrm{c}_i$ $(i = 1, 2)$, respectively. $\eta_{\mathrm{c}_i\mathrm{u}}$ and $\eta_{\mathrm{c}_i\mathrm{b}}$ are energy terms describing the effect of nuclear transport on nucleocytoplasmic distributions of histone-binding chaperones, which are defined by formulas similar to Eq. (M5). $\chi_{\mathrm{nucl}}(\mathbf{r})$ is the characteristic function of the cell nucleus: $\chi_{\mathrm{nucl}}(\mathbf{r}) = 1$ if the position vector $\mathbf{r}$ corresponds to a point located inside the cell nucleus, otherwise $\chi_{\mathrm{nucl}}(\mathbf{r}) = 0$. $V^{\mathrm{osm}}_{\mathrm{cell}}$ is the osmotically active volume of a cell. $\beta$ is the inverse of thermodynamic temperature: $\beta = 1/k_{\mathrm{B}}T$.

Thus, the presence of nuclear transport leads to $\eta_{\mathrm{c}_i\mathrm{u}}$ and $\eta_{\mathrm{c}_i\mathrm{b}}$ offsets of the chemical potentials of histone-binding chaperones localized in the cell nucleoplasm with respect to the chemical potentials of the same type of chaperones in the cell cytosol.

From Appendices A-B, it can be seen that application of the above changes to Eq. (A15) leads to the following form of Eq. (9) from the main text:

$$\mu^{\mathrm{tot}}_{\mathrm{pr}} \approx \mu^0_{\mathrm{pr}} - q_{\mathrm{oct}}\langle\psi\rangle + 2k_{\mathrm{B}}T\ln\theta_{\mathrm{c}_1} + 2k_{\mathrm{B}}T\ln\theta_{\mathrm{c}_2} + 2k_{\mathrm{B}}T\sum_{i=1,2}\ln\left(\frac{V^{\mathrm{osm}}_{\mathrm{cell}} + V^{\mathrm{pr}}_{\mathrm{nucl}}[e^{\beta\eta_{\mathrm{c}_i\mathrm{u}} - \beta q_{\mathrm{c}_i\mathrm{u}}\langle\psi\rangle} - 1]}{V^{\mathrm{osm}}_{\mathrm{cell}} + V^{\mathrm{pr}}_{\mathrm{nucl}}[e^{\beta\eta_{\mathrm{c}_i\mathrm{b}} - \beta q_{\mathrm{c}_i\mathrm{b}}\langle\psi\rangle} - 1]}\right) \tag{M8}$$

Here $\theta_{\mathrm{c}_i} = n^0_{\mathrm{c}_i\mathrm{b}}/n^0_{\mathrm{c}_i\mathrm{u}}$ $(i = 1, 2)$ are occupancy ratios of histone-binding chaperones, where $n^0_{\mathrm{c}_1\mathrm{b}}$, $n^0_{\mathrm{c}_2\mathrm{b}}$, $n^0_{\mathrm{c}_1\mathrm{u}}$ and $n^0_{\mathrm{c}_2\mathrm{u}}$ are the average total numbers of $\mathrm{c}_1$ and $\mathrm{c}_2$ chaperones in histone-bound and unloaded states, respectively, see Appendix A. $V^{\mathrm{pr}}_{\mathrm{nucl}}$ is the nucleus volume accessible to proteins, such as histone-binding chaperones, which in our study was set equal to the total nucleus volume, $V_{\mathrm{nucl}}$: $V^{\mathrm{pr}}_{\mathrm{nucl}} = V_{\mathrm{nucl}}$. Finally, $\mu^0_{\mathrm{pr}}$ is the standard Gibbs free energy of nucleosome formation in the presence of histone-binding chaperones under the standard experimental conditions: $\theta_{\mathrm{c}_1} = \theta_{\mathrm{c}_2} = 1$ and $\psi = 0$.

Comparing the above formula to Eq. (9), it can be seen that introduction of nuclear transport into the model leads only to a slight change in the last logarithmic term of Eq. (M8) – two new exponential factors, $e^{\beta\eta_{\mathrm{c}_i\mathrm{u}}}$ and $e^{\beta\eta_{\mathrm{c}_i\mathrm{b}}}$, appear in the numerator and denominator.

Using Eq. (M8), it then can be shown that $\eta_{\mathrm{c}_i\mathrm{u}}$ and $\eta_{\mathrm{c}_i\mathrm{b}}$ energies have a negligible effect on stability of nucleosomes, and thus the chromatin structure, as soon as $|\eta_{\mathrm{c}_i\mathrm{u}}|$ and $|\eta_{\mathrm{c}_i\mathrm{b}}|$ are less than 1 $k_{\mathrm{B}}T$ [compare Figure S16(b) to Figure 5 from the main text]. Since this criterion is satisfied under normal physiological conditions (see Table II), it can be concluded that the main results of our study remain valid even in the presence of active nuclear transport. Though, it should be noted that plots shown in Figures S16(a-c) indicate that under certain conditions living cells may utilize regulation of nucleocytoplasmic trafficking of histone-binding chaperones to induce changes in chromatin structure by increasing the absolute values of $\eta_{\mathrm{c}_i\mathrm{u}}$ and $\eta_{\mathrm{c}_i\mathrm{b}}$ energies above 1 $k_{\mathrm{B}}T$ level, which warrants future studies.

Finally, it should be noted that recent experimental studies show that proteins with a molecular weight $\leq 70$ kDa have access to only $\sim 60\% - 70\%$ of the cell nucleus volume under normal experimental conditions [61], – a number that can locally drop as low as $0\% - 10\%$ in nuclei of cells moving through narrow constrictions [62]. To investigate a potential role of such a protein exclusion effect by chromatin in regulation of the histone binding free energy to DNA, we varied $V^{\mathrm{pr}}_{\mathrm{nucl}}$ nucleus volume accessible to proteins in Eq. (M8) in $10\%V_{\mathrm{nucl}} - 70\%V_{\mathrm{nucl}}$ range, making the plots shown in Figures S16(d-f). By comparing the graphs displayed in Figure S16 to Figure 5 from the main text, it can be seen that: 1) exclusion of proteins by chromatin produces the same effect on the binding free energy of histone octamers to DNA as negative $\eta_{c_i u}$ and $\eta_{c_i b}$ energy terms, and 2) it has a negligible impact on the binding free energy of histone octamers to DNA in the case of $V^{\mathrm{pr}}_{\mathrm{nucl}} = 70\% \cdot V_{\mathrm{nucl}}$, which corresponds to normal experimental conditions [61]. The latter result justifies $V^{\mathrm{pr}}_{\mathrm{nucl}} = V_{\mathrm{nucl}}$ approximation, which has been used in our study.

TABLE I: Values of the model parameters used in calculations.

| **Parameter** | **Value** | **Description** |
|---|---|---|
| $b$ | 3.4 nm | Length of bare DNA segments. |
| $A$ | 50 nm | Bending persistence length of B-DNA [63, 64]. |
| $C$ | 95 nm | Twisting persistence length of B-DNA [65–67]. |
| $a, a_{\rm pr}$ | $a = a_{\rm pr} = \frac{A}{b} \approx 14.7$ | Dimensionless bending elasticities of bare DNA segments far away and next to the entry points of DNA into nucleosomes. |
| $c, c_{\rm pr}$ | $c = c_{\rm pr} = \frac{C}{b} \approx 27.9$ | Dimensionless twisting elasticities of bare DNA segments far away and next to the entry points of DNA into nucleosomes. |
| $T$ | 310 K | Environment temperature. |
| $N_{\rm bp}$ | 40 kbp (EMBL3 $\lambda$-phage) or 6.2 Gbp (human) | Length of chromosomal DNA [68, 69]. |
| $c_{\rm ions}$ | 150 mM | Cytosolic concentration of positive and negative ions [11–14]. |
| $q_{\rm c_1 b}, q_{\rm c_2 b}$ | $+9\, q_{\rm e}$ | Electrical charge of histone-bound chaperones estimated based on experimental data from ref. [10, 21, 22]. |
| $q_{\rm c_1 u}, q_{\rm c_2 u}$ | $-28.2\, q_{\rm e}$ | Electrical charge of unloaded chaperones estimated based on experimental data from ref. [10, 21, 22]. |
| $q_{\rm oct}$ | $+148.8\, q_{\rm e}$ | Electrical charge of histone octamers calculated based on the primary sequence of histones. |
| $\rho_{\rm d}$ | $-2q_{\rm e}$ per DNA base-pair | Linear electrical charge density of DNA. |
| $\varepsilon_r$ | 75 | Relative permittivity of water [70]. |
| $\theta_{\rm c_1}, \theta_{\rm c_2}$ | 0.17 | Occupancy ratios of histone-binding chaperones estimated based on experimentally measured occupancy fraction of chromosomal DNA by nucleosomes ($\sim 0.7 - 0.9$ [22, 71, 72]). |
| $\mu^0_{\rm pr}$ | 7 $k_{\rm B}T$ | Standard Gibbs free energy of nucleosome formation in the presence of histone-binding chaperones under the standard experimental conditions: $\theta_{\rm c_1} = \theta_{\rm c_2} = 1$ [32, 33]. |
| $r_{\rm pr}$ | 5.85 nm | The average distance between the entry and exit points of DNA in a nucleosome estimated based on crystallographic data from ref. [37, 38]. |
| $\mathbf{A}_{\rm in}$ | $(0, 1.92, -0.61)$ | Euler angles of the rotation matrix $\mathbf{A}_{\rm in}$ describing an equilibrium orientation of the DNA segment entering a nucleosome with respect to the nucleosome core estimated based on the crystallographic data from ref. [37, 38]. |
| $\mathbf{A}_{\rm out}$ | $(-0.61, 1.92, 0)$ | Euler angles of the rotation matrix $\mathbf{A}_{\rm out}$ describing an equilibrium orientation of the DNA segment exiting a nucleosome with respect to the nucleosome core estimated based on crystallographic data from ref. [37, 38]. |
| $K$ | 14 | Number of DNA segments bound to a histone octamer in each nucleosome, which corresponds to $\sim 145 - 147$ bp of nucleosomal DNA [37, 38]. |
| $n_{\rm max}$ | 18 | Number of terms in the Fourier-Bessel expansion series of the electrostatic potential, $\psi$. |
| $s_{\rm max}$ | 7-9 | The upper bound on the principal index of Wigner D-functions used in the expansion series of DNA transfer-functions. |
| $\Vert\mathbf{a}_1\Vert$, $\Vert\mathbf{a}_2\Vert$, $\Vert\mathbf{a}_3\Vert$ | $2.4\, R$ | Length of the primitive vectors $(\mathbf{a}_1, \mathbf{a}_2, \mathbf{a}_3)$ spanning a hexagonal Bravais lattice used in the model calculations, where $R$ is the radius of the cell nucleus or a viral particle. |
| $l_{\rm max}$ | 2500-14800 | The total number of reciprocal lattice nodes employed in computation of the DNA partition function. |

TABLE II: Correction energy term, $\eta$, estimated based on experimental measurements reported in ref. [60] for proteins of different molecular weights.

| **Correction energy term** | **Molecular weight of a protein** | | | |
|---|---|---|---|---|
| | 27 **kDa** | 41 **kDa** | 54 **kDa** | 67 **kDa** |
| $\eta$ | $-0.18\ k_{\mathrm{B}}T$ | $-0.58\ k_{\mathrm{B}}T$ | $-0.38\ k_{\mathrm{B}}T$ | $-0.02\ k_{\mathrm{B}}T$ |

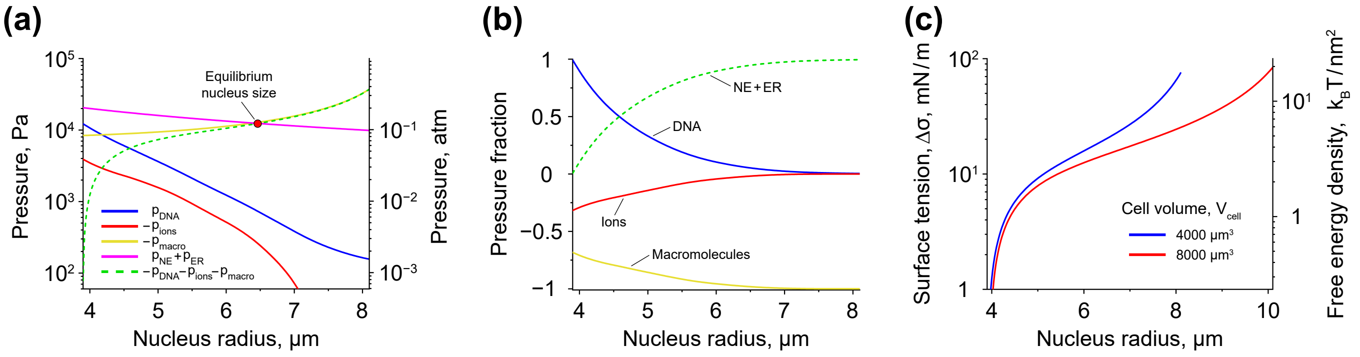


FIG. S1. **Cell nucleus size regulation.** **(a)** Pressures generated on the NE by DNA, ions, macromolecules, and the surface tensions of the NE and ER membrane in the case of the total cell volume of $V_{\rm cell} = 4000\ \mu{\rm m}^3$. The yellow curve representing the osmotic pressure created by macromolecules is plotted for the case of $n_{\rm macro} = 5 \cdot 10^9$ molecules; whereas, the magenta curve demonstrates the case of the NE and ER membrane surface tension difference of $\Delta\sigma = 20$ mN/m. In transfer-matrix calculations, the total length of DNA was set equal to 2.1 m, corresponding to the size of human genome of $\sim 6.2$ Gbp [69]. The length of bare DNA segments in the discretized polymer model was $b = 3.4$ nm. **(b)** Relative contributions of different cell components to the total positive / negative pressure acting on the NE in the case of the total cell volume of $V_{\rm cell} = 4000\ \mu{\rm m}^3$. **(c)** Difference between the surface tensions of the NE and ER membrane, $\Delta\sigma = \sigma_{\rm ER} - \sigma_{\rm NE}$, that has to be maintained by a living cell in order to retain the nucleus size, $R_{\rm nucl}$, at a specific value. Blue and red curves are calculated based on the green dashed curves shown in panel (a) and Figure 3(a), respectively, by using the following formula: $\Delta\sigma = -\frac{1}{4} R_{\rm nucl}(p_{\rm DNA} + p_{\rm ions} + p_{\rm macro})$, see Eq. (K9) and Eq. (6) in the main text. In the calculations, parameter $\zeta$ was put equal to $\zeta = 0.45$.

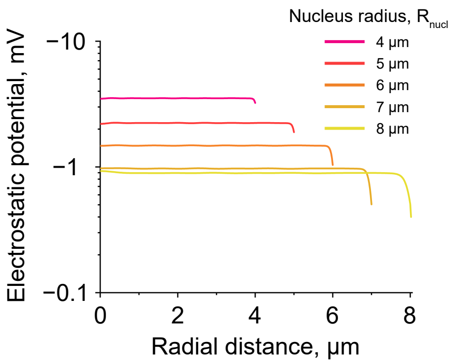


FIG. S2. **Nuclear electrostatic potential in nuclei of different sizes.** The plot shows results obtained for a cell with the total volume of $V_{\rm cell} = 4000\ \mu{\rm m}^3$.

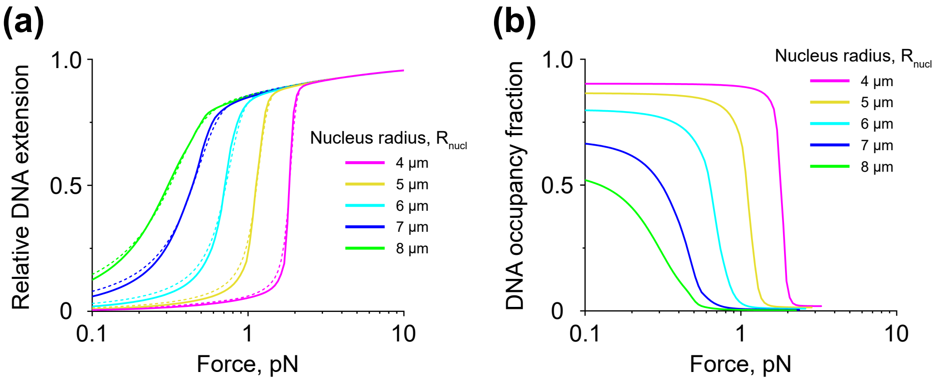


FIG. S3. **Correlation between the nucleus size and nucleosome stability.** **(a)** and **(b)** Force-extension and force-DNA occupancy fraction curves obtained by mechanical stretching of a small part of chromosomal DNA (2 $\mu$m in length) in nuclei of different sizes. Solid curves on panels (d) and (e) demonstrate results of transfer-matrix calculations performed on a cell with the total volume of $V_{\mathrm{cell}} = 4000\ \mu\mathrm{m}^3$. Dashed curves in panel (a) display results of the force-extension curves' fitting to the previously published model of a mechanically stretched DNA [3].

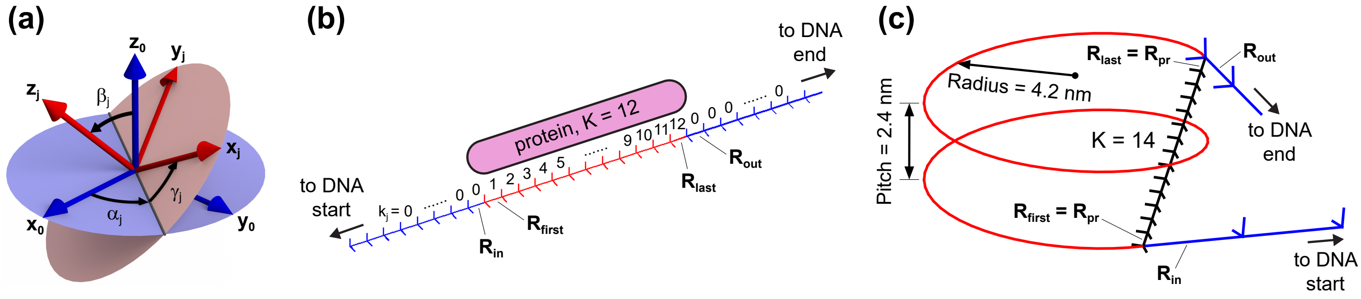


FIG. S4. **Model description of nucleoprotein complexes.** **(a)** In the discretized polymer model of DNA, 3D-orientation of each DNA segment shown in Figure 1(c) is represented by a corresponding Euler rotation matrix, $\mathbf{R}_j$, which is obtained as the composition of three successive revolutions of the coordinate frame $(\mathbf{x}_j, \mathbf{y}_j, \mathbf{z}_j)$ attached to each DNA segment through Euler angles $\alpha_j$, $\beta_j$ and $\gamma_j$ about the axes of the fixed global coordinate system, $(\mathbf{x}_0, \mathbf{y}_0, \mathbf{z}_0)$. Here $j$ is the index enumerating DNA segments: $j = 1, ..., N$. $N$ is the total number of segments in the polygonal chain representing DNA. **(b)** Schematic picture of a nucleoprotein complex in the case when a protein occupies $K = 12$ DNA segments upon binding to DNA. Bare DNA segments are displayed in blue color and protein-bound DNA segments engaged in formation of the nucleoprotein complex are shown in red. The numbers above the DNA segments indicate $k_j$ indexes of the corresponding DNA segments, which are used in the model to describe physical states of DNA segments and their location within nucleoprotein complexes. **(c)** Model representation of nucleosomes. For the sake of DNA partition function calculations, DNA segments constrained inside each nucleosome are represented by straight segments connecting the entry and exit points of DNA in the nucleosome, which are shown in black color in the figure. The red line indicates the real path of DNA through the nucleoprotein complex. In panels (b) and (c), $\mathbf{R}_{\rm in}$ and $\mathbf{R}_{\rm out}$ Euler rotation matrices denote the orientations of the DNA segments sitting next to the entry and exit points of the nucleoprotein complex; whereas, matrices $\mathbf{R}_{\rm first}$ and $\mathbf{R}_{\rm last}$ designate orientations of the first and the last DNA segments inside the complex.

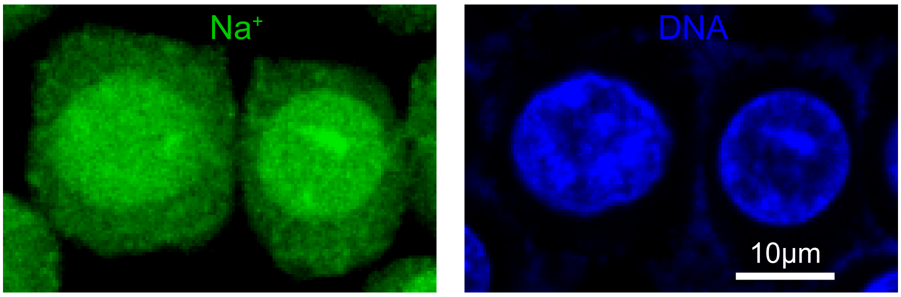


FIG. S5. **Confocal images of HEK-293 cells labelled with sodium indicator, ANG-2 (left panel), and DNA stain, Hoechst (right panel).** From the image it can be seen that the concentration of $Na^+$ ions is elevated in nuclei of living cells in comparison to the cell cytoplasm due to their interaction with the negative electrostatic potential of the cell nucleus and chromatin.

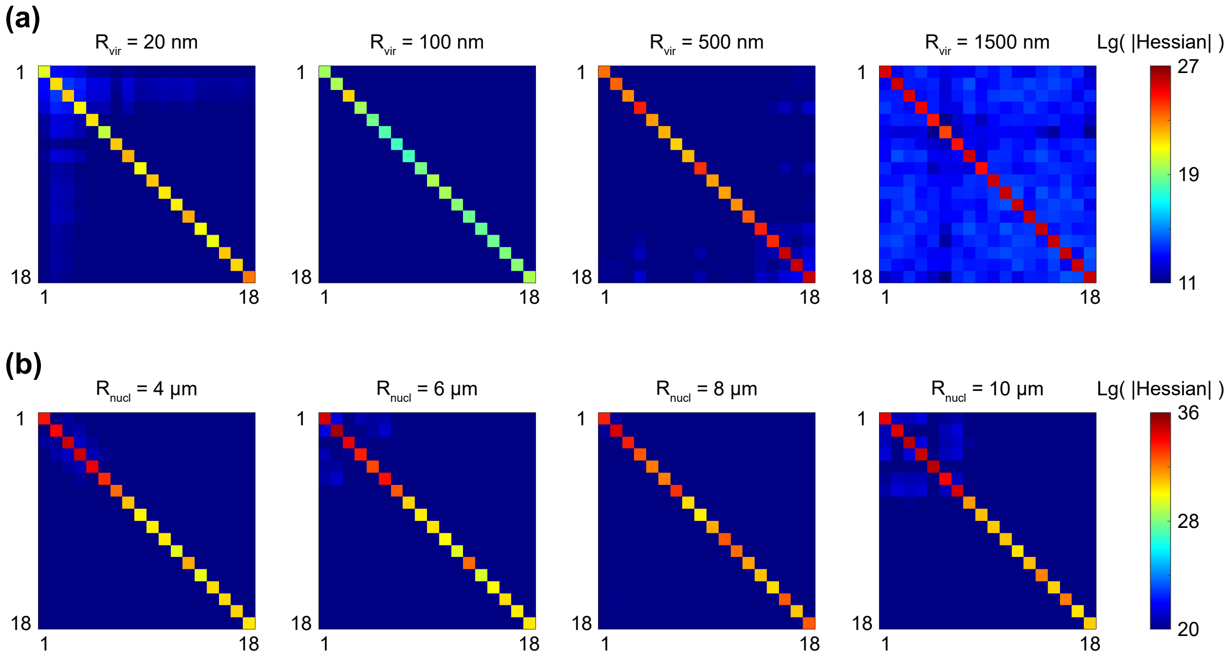


FIG. S6. **Hessian matrices.** The figure demonstrates the absolute values of the first 18 × 18 elements of $\left[\frac{\partial^2 \ln Z}{\partial\psi_n \partial\psi_m}\right]$ Hessian matrix plotted in the logarithmic scale for different sizes of **(a)** viral particles and **(b)** nuclei of living cells. From the figure it can be seen that all Hessian matrices have a nearly diagonal form. By taking into account that their diagonal elements are all negative, it can be concluded that these matrices are negative-definite, from which it immediately follows that the stationary phase potential, $\psi_{\rm sp}$, that solves Eq. (A37) corresponds to the maximum of the partition function or, which is the same, to the minimum of the free energy of the system. In the figure, $R_{\rm vir}$ and $R_{\rm nucl}$ indicate the radii of the viral capsid and the cell nucleus, accordingly.

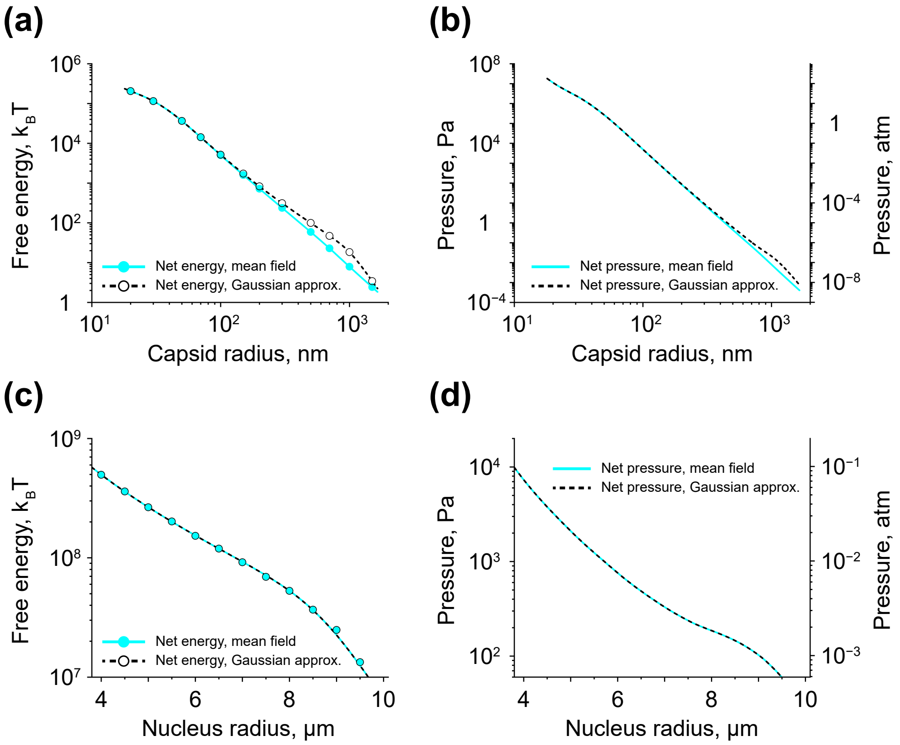


FIG. S7. **Mean field approach vs Gaussian approximation.** **(a, b)** The total free energy of a viral particle and the net pressure exerted by DNA and ions on the capsid wall as functions of the capsid radius, $R_{\rm vir}$. **(c, d)** The total free energy of DNA and ions and the net pressure developed by them on the NE as functions of the nucleus radius, $R_{\rm nucl}$. In the graphs, the data points indicate results of transfer-matrix calculations; whereas, the lines show smoothing spline interpolation. Solid cyan curves correspond to the case of the mean field approach; whereas, the dashed black curves represent the case of Gaussian approximation. As can be seen from the figure, field fluctuations near the stationary phase potential, $\psi_{\rm sp}$, have a negligible effect on the total free energy of the system as well as parameters describing its physical state, such as the pressure developed by DNA and ions, in the physiologically relevant range of 20 nm $\leq R_{\rm vir} \leq$ 100 nm and 4 $\mu$m $\leq R_{\rm nucl} \leq$ 10 $\mu$m.

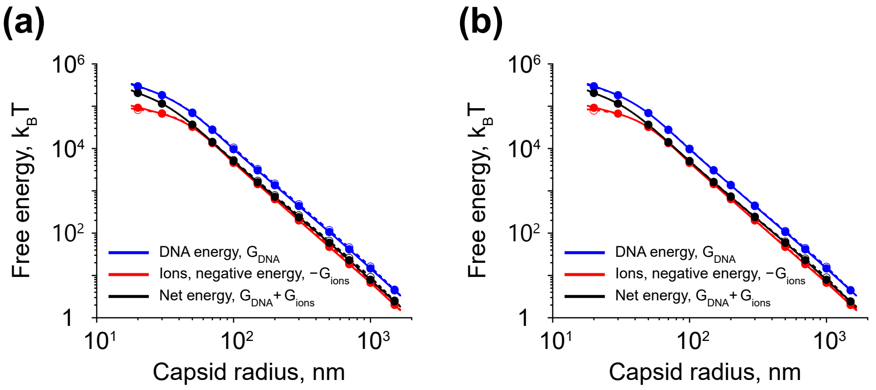


FIG. S8. **Robustness of the model predictions with respect to variation in the DNA segment size, $b$, and the total number of reciprocal lattice nodes, $l_{\mathbf{max}}$, used in the model computations. (a, b)** Contributions made by DNA (blue curve) and ions (red curve) into the total free energy of a viral particle (black curve) as functions of the capsid size. Data points shown in the graphs are the results of transfer-matrix calculations; whereas, the lines indicate smoothing spline interpolation. In both panels, solid curves correspond to the case of $b = 3.4$ nm and $l_{\max} = 5149$. As for the dashed curves, in panel (a) they demonstrate results obtained for $b = 1$ nm and $l_{\max} = 5149$ scenario; whereas, in panel (b) – for the case of $b = 3.4$ nm and $l_{\max} = 2535$. As can be seen from the figure, neither the DNA segment size, nor the total number of reciprocal lattice nodes have a strong effect on the graphs, suggesting robustness of the model to changes in the values of these parameters.

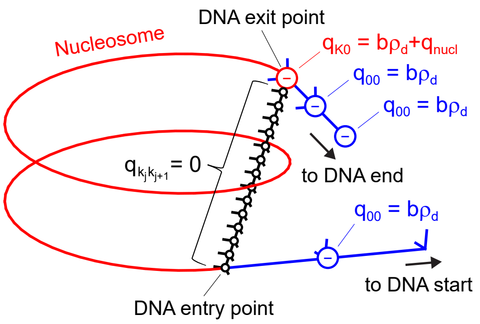


FIG. S9. **Distribution of electrical charges on DNA in the discretized polygonal chain model.** For the sake of faster calculations, the net electrical charge of each nucleosome ($q_{\mathrm{nucl}}$) as well as the bare DNA segment in front of it ($q_{\mathrm{bare}} = b\rho_{\mathrm{d}}$) was assigned to the joint located at the exit point of DNA from the nucleosome as schematically shown in the figure. Electrical charges of the rest of bare DNA segments ($q_{\mathrm{bare}} = b\rho_{\mathrm{d}}$) were assigned to the nearest downstream joints of the polygonal chain representing DNA. Furthermore, it has been assumed in the model that only the joints which are followed by a bare DNA segment contribute to the confinement energy term in Eq. (B1). As can be seen from the figure, these are exactly the same joints as ones possessing a non-zero electrical charge. In the figure, bare DNA segments are shown in blue color; whereas, DNA constrained inside the nucleosome is displayed in red. Subintervals of the polygonal chain representing the DNA segments bound to the nucleosome are shown in black color, see Appendix B and Figure S4(c) for more details.

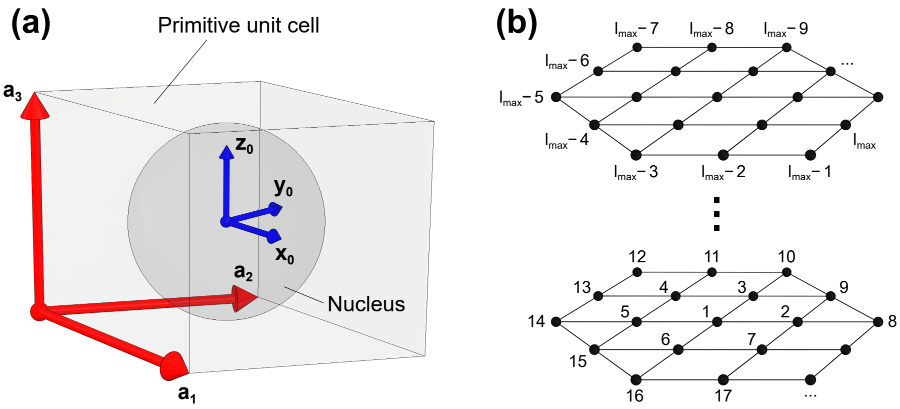


FIG. S10. **Primitive unit cell of the Bravais lattice and enumeration of the reciprocal lattice nodes.** **(a)** To estimate the DNA partition function, we utilized a hexagonal Bravais lattice spanned by primitive vectors, $(\mathbf{a}_1, \mathbf{a}_2, \mathbf{a}_3)$, which were long enough for a primitive unit cell of the Bravais lattice to cover the entire cell nucleus or a viral particle placed at the origin of the global coordinate system, $(\mathbf{x}_0, \mathbf{y}_0, \mathbf{z}_0)$, as schematically shown in the figure. **(b)** Schematic picture of one of the possible ways to enumerate the nodes of the reciprocal lattice, which was used in the model. $l_{\max}$ is the total number of nodes employed in the DNA partition function calculations, see Appendix D and Table I for more details.

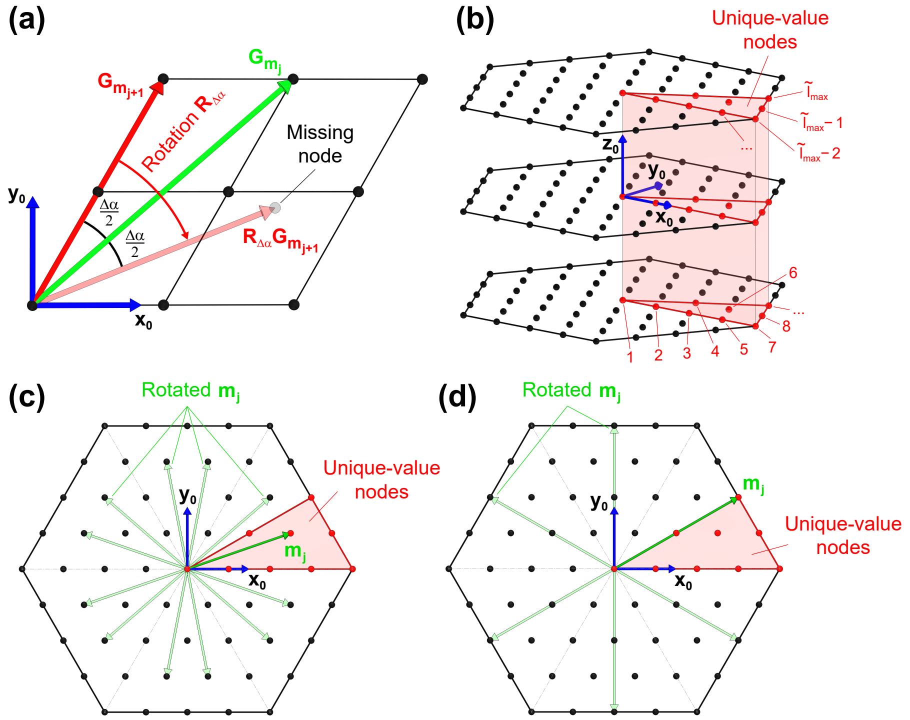


FIG. S11. **Unique-value nodes of the reciprocal lattice.** **(a)** In the general case, the wave-vector rotation method cannot be directly applied to Eq. (H12) due to the absence of necessary nodes in the reciprocal lattice. The panel demonstrates one of such cases. It can be seen from the figure that $\mathbf{G}_{\mathbf{m}_{j+1}}$ wave-vector cannot be rotated about $\mathbf{z}_0$-axis of the global coordinate system by $\Delta\alpha = 2(\alpha_{\mathbf{G}_{\mathbf{m}_j}} - \alpha_{\mathbf{G}_{\mathbf{m}_{j+1}}})$ angle as the reciprocal lattice does not have a corresponding node. **(b)** An example of unique-value nodes of the reciprocal lattice (shown in red color). These are the nodes that cannot be obtained from each other by rotation about $\mathbf{z}_0$-axis of the global coordinate system, while the rests of the reciprocal lattice can be generated from the unique-value nodes via such a rotation. The panel also shows one of the ways to enumerate the unique-value nodes, which was adapted in our study. $\widetilde{l}_{\max}$ is the total number of the unique-value nodes used in the DNA partition function calculations. **(c, d)** Two examples of possible rotations of a unique-value vector, $\mathbf{m}_j$, about $\mathbf{z}_0$-axis of the global coordinate system.

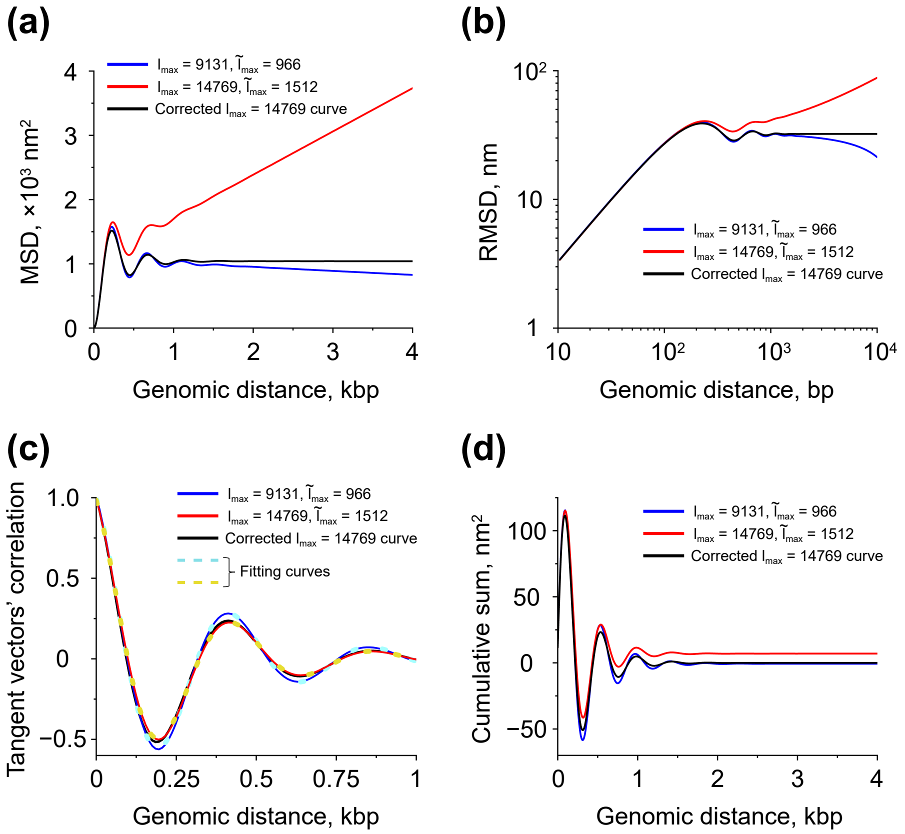


FIG. S12. **Tangent vector correlation function and the mean squared displacement (MSD) between two points on a viral DNA as a function of the genomic distance between them.** **(a, b)** MSD and RMSD curves obtained for different numbers of reciprocal lattice nodes, $l_{\rm max}$, by using Eq. (I11). **(c)** Tangent vector correlation function calculated for two different values of $l_{\rm max}$ with the help of Eq. (J2). Yellow and cyan dotted lines demonstrate fitting of the shown tangent vector correlation curves to the damped oscillation function defined by Eq. (J6). **(d)** Value of the tangent vector correlation sum in the right-hand side of Eq. (J5) as a function of the genomic distance between two points on DNA. In all panels, the black curve indicates results obtained by application of the renormalization procedure described by Eq. (J8) to the tangent vector correlation curve corresponding to $l_{\rm max} = 14769$ ($\widetilde{l}_{\rm max} = 1512$) case, which is displayed in red color in panel (c).

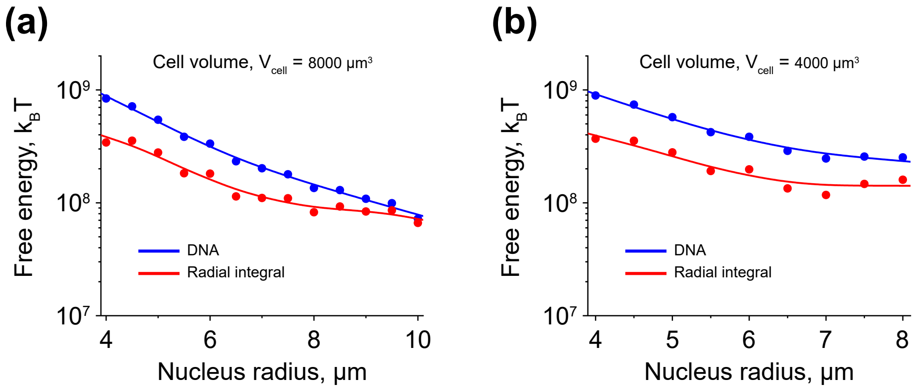


FIG. S13. **Smoothing spline interpolation of the free energy of DNA and the radial integral from Eq.** (K5)**.** Panels **(a)** and **(b)** show the results obtained for cells with the total volumes of $V_{\rm cell} = 8000\ \mu{\rm m}^3$ and $V_{\rm cell} = 4000\ \mu{\rm m}^3$, respectively. Solid lines indicate smoothing spline interpolation of the free energy of DNA, $G_{\rm DNA}$, (blue curves) and the radial integral from Eq. (K5), $c_{\rm ions} k_{\rm B} T \int_0^{R_{\rm nucl}} 8\pi r^2 \left[\cosh(\beta q_{\rm e} \psi_{\rm sp}(r)) - 1\right] {\rm d}r$, (red curves) based on fitting of the data points calculated at different values of the nucleus radius, $R_{\rm nucl}$. In the calculations, the total length of DNA was set equal to 2.1 m, corresponding to the size of human genome of $\sim 6.2$ Gbp [69].

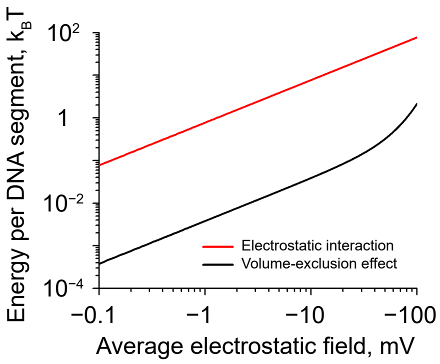


FIG. S14. **Energies of electrostatic and volume-exclusion DNA interactions per single DNA segment as functions of the strength of the average stationary phase electrostatic potential, $\hat{\psi}_{sp}$.** From the graph it can be seen that electrostatic forces dominate over the volume-exclusion effect in the whole range of $-100$ mV $\leq \hat{\psi}_{sp} \leq 0$ mV. The length of DNA segments used in the model calculations was $b = 3.4$ nm.

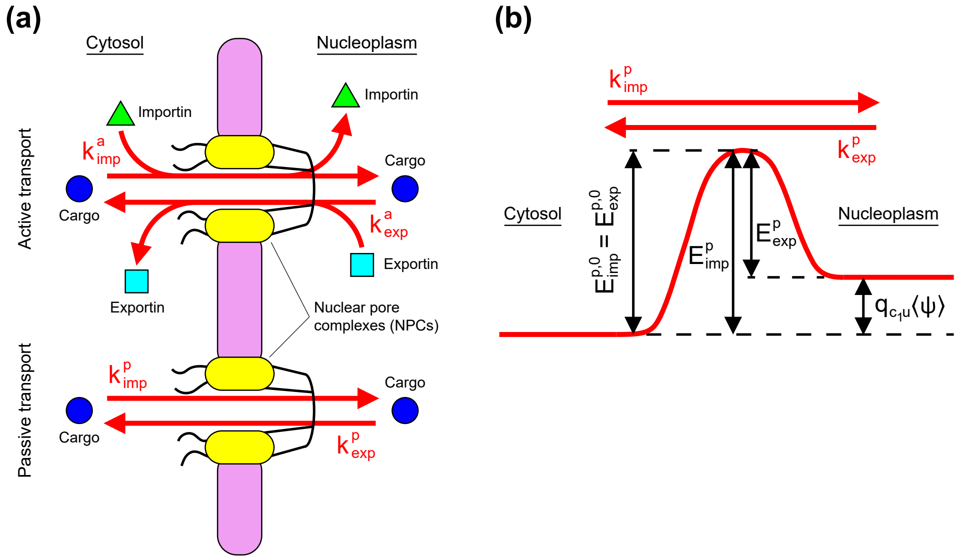


FIG. S15. **Pseudo-first order kinetic model of nuclear transport of proteins through nuclear pore complexes (NPCs).** **(a)** In the model, two types of processes by which proteins can move between the cell cytosol and nucleoplasm are considered: 1) via passive diffusion through NPCs or 2) by utilizing active nuclear transportation system mediated by importins and exportins that recognize nuclear localization (NLS) and nuclear export (NES) signals carried by proteins. Following ref. [60], both passive and active transportation of a cargo protein between the cell cytosol and the nucleoplasm are described in the model by pseudo-first order kinetic rates, $k^{\rm a}_{\rm imp}$, $k^{\rm a}_{\rm exp}$, $k^{\rm p}_{\rm imp}$ and $k^{\rm p}_{\rm exp}$, shown in the figure that can be determined based on photobleaching experiments. **(b)** Schematic picture of the activation energies, $E^{\rm p}_{\rm imp}$ and $E^{\rm p}_{\rm exp}$, of passive nuclear import and export of $\rm c_1u$ chaperones, respectively. From the figure it can be seen that due to the interaction of electrically charged chaperones (carrying a net charge of $q_{\rm c_1u}$) with the average nuclear electrostatic potential, $\langle\psi\rangle$, the activation energy of the passive nuclear export gains $-q_{\rm c_1u}\langle\psi\rangle$ offset with respect to the activation energy $E^{\rm p,0}_{\rm exp}$ of the passive nuclear export of proteins, which have a similar size, but carry a net zero electrical charge. At the same time, the activation energy of the passive nuclear import, $E^{\rm p}_{\rm imp}$, is independent from the electrical charge of transported proteins ($E^{\rm p}_{\rm imp} = E^{\rm p,0}_{\rm imp}$), as schematically shown in the figure. A very similar conclusion can be drawn for the activation energies of the active nuclear import and export of $\rm c_1u$ chaperones, as well as for the rest of chaperones, $\rm c_2u$, $\rm c_1b$ and $\rm c_2b$.

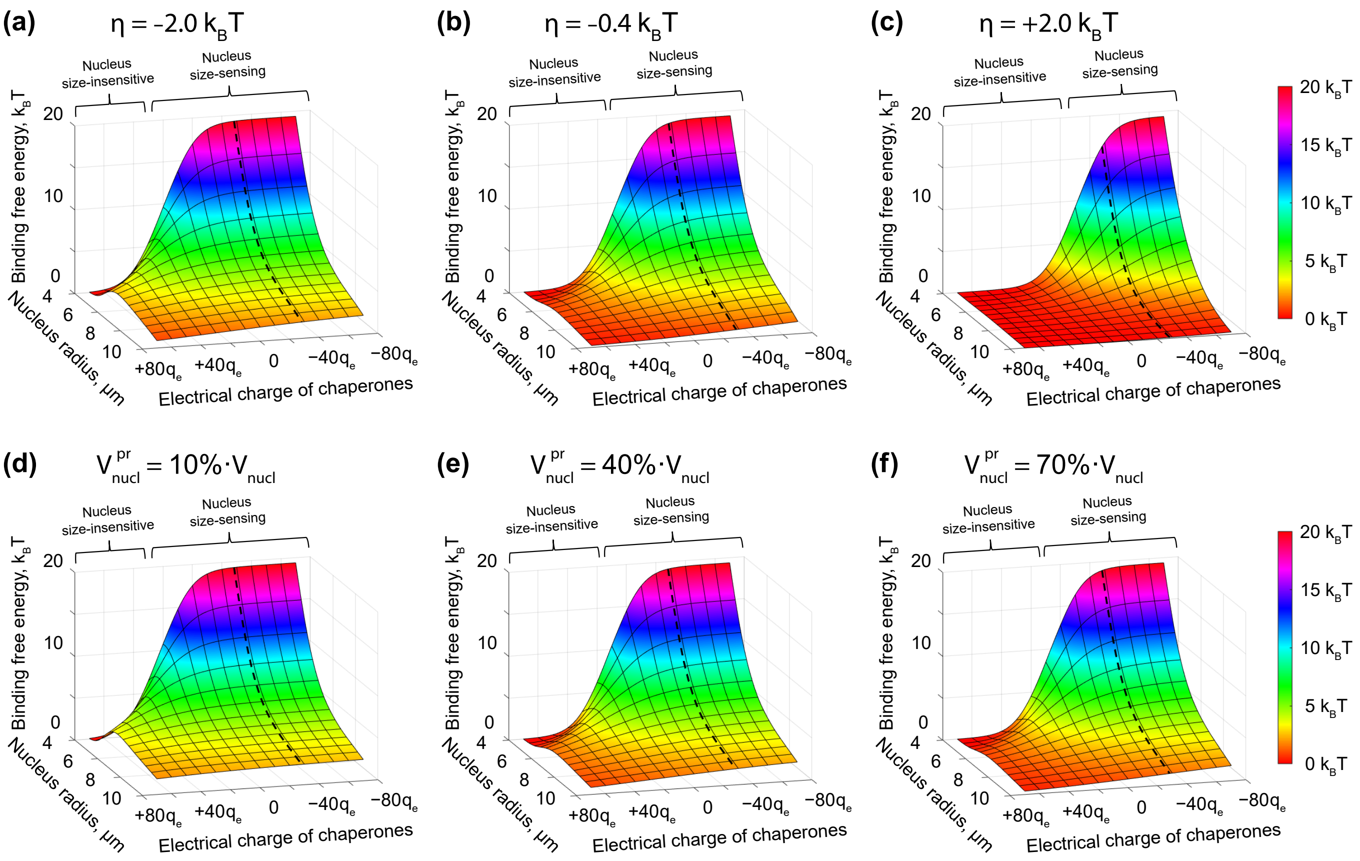


FIG. S16. **Roles of the nuclear transport (a-c) and protein exclusion effect by chromatin (d-f) in regulation of the binding free energy of histone octamers to DNA.** Plots demonstrate results of calculations based on Eq. (M8) performed for different electrical charges of histone-binding chaperones and cell nucleus sizes. For the sake of simplicity, it was assumed in the calculations that H2A·H2B-binding chaperones as well as H3·H4-binding chaperones, both have similar electrical charges: $q_{\mathrm{c_1 u}} = q_{\mathrm{c_2 u}}$. In panels (a-c), the nucleus volume accessible to histone-binding chaperones ($V^{\mathrm{pr}}_{\mathrm{nucl}}$) was put equal to the total volume of the cell nucleus ($V^{\mathrm{pr}}_{\mathrm{nucl}} = V_{\mathrm{nucl}}$); whereas, all $\eta_{\mathrm{c}_i \mathrm{u}}$ and $\eta_{\mathrm{c}_i \mathrm{b}}$ $(i = 1, 2)$ energy terms describing the effect of the nuclear transport onto the distribution of histone-binding chaperones between the cell cytosol and nucleoplasm were all set equal to the same parameter, $\eta$, which was varied from $-2.0\ k_{\mathrm{B}}T$ to $+2.0\ k_{\mathrm{B}}T$. The case of $\eta = -0.4\ k_{\mathrm{B}}T$ corresponds to the value estimated based on experimental measurements done on living cells under normal physiological conditions (see Appendix M and Table II). In panels (d-f), the nucleus volume accessible to histone-binding chaperones was ranged from $V^{\mathrm{pr}}_{\mathrm{nucl}} = 10\% \cdot V_{\mathrm{nucl}}$ to $V^{\mathrm{pr}}_{\mathrm{nucl}} = 70\% \cdot V_{\mathrm{nucl}}$; whereas, $\eta_{\mathrm{c}_i \mathrm{u}}$ and $\eta_{\mathrm{c}_i \mathrm{b}}$ $(i = 1, 2)$ energy terms were set equal to $0\ k_{\mathrm{B}}T$. Under normal physiological conditions, the nucleus volume accessible to proteins with a molecular weight $\leq 70$ kDa is equal to $\sim 60\% - 70\% \cdot V_{\mathrm{nucl}}$, see ref. [61]. Dashed curves shown in the graphs indicate model predictions based on estimation of the chaperone electrical charge ($q_{\mathrm{c_1 u}} = q_{\mathrm{c_2 u}} = -28.2\, q_{\mathrm{e}}$, where $q_{\mathrm{e}} = 1.6 \cdot 10^{-19}$ C is the elementary charge), which was obtained from experimental measurements of relative histone concentrations in the cytosol and nucleoplasm of HeLa cells [22], see Appendix A for details. By comparing the above graphs shown in panels (b) and (f) to Figure 5 from the main text, it can be seen that nuclear transport and exclusion of proteins by chromatin have a negligible effect on the binding free energy of histone octamers to DNA and, as a result, on chromatin structure under normal physiological conditions.

[1] A. K. Efremov, R. S. Winardhi, and J. Yan, Phys. Rev. E **94**, 032404 (2016).
[2] A. K. Efremov, R. S. Winardhi, and J. Yan, Polymers **9**, 74 (2017).
[3] A. K. Efremov and J. Yan, Nucleic Acids Res. **46**, 6504 (2018).
[4] S. F. Edwards, Proc. Phys. Soc. **85**, 613 (1965).
[5] H. Kleinert, *Path integrals in quantum mechanics, statistics, polymer physics, and financial markets.*, 5th ed. (World Scientific Publishing, Singapore, 2009).
[6] X. Qiu, D. C. Rau, V. A. Parsegian, L. T. Fang, C. M. Knobler, and W. M. Gelbart, Phys. Rev. Lett. **106**, 028102 (2011).
[7] D. W. Bauer, D. Li, J. Huffman, F. L. Homa, K. Wilson, J. C. Leavitt, S. R. Casjens, J. Baines, and A. Evilevitch, J. Virol. **89**, 9288 (2015).
[8] R. Milo and R. Phillips, *Cell biology by the numbers.* (Garland Science, New York, United States of America, 2016).
[9] T. J. Century, I. R. Fenichel, and S. B. Horowitz, J. Cell Sci. **7**, 5 (1970).
[10] H. Oberleithner, S. Wünsch, and S. Schneider, Proc. Natl. Acad. Sci. U.S.A. **89**, 241 (1992).
[11] M. V. Martinov, V. M. Vitvitsky, and F. I. Ataullakhanov, Biophys. Chem. **80**, 199 (1999).
[12] J. Keener and J. Sneyd, *Mathematical physiology I: cellular physiology.*, 2nd ed. (Springer, New York, United States of America, 2009).
[13] A. R. Kay, Front. Cell Dev. Biol. **5**, 41 (2017).
[14] A. Fazelkhah, K. Braasch, S. Afshar, E. Salimi, M. Butler, G. Bridges, and D. Thomson, Sci. Rep. **8**, 17818 (2018).
[15] E. H. Zhou, X. Trepat, C. Y. Park, G. Lenormand, M. N. Oliver, S. M. Mijailovich, C. Hardin, D. A. Weitz, J. P. Butler, and J. J. Fredberg, Proc. Natl. Acad. Sci. U.S.A. **106**, 10632 (2009).
[16] M. Guo, A. F. Pegoraro, A. Mao, E. H. Zhou, P. R. Arany, Y. Han, D. T. Burnette, M. H. Jensen, K. E. Kasza, J. R. Moore, F. C. Mackintosh, J. J. Fredberg, D. J. Mooney, J. Lippincott-Schwartz, and D. A. Weitz, Proc. Natl. Acad. Sci. U.S.A. **114**, E8618 (2017).
[17] L. De Koning, A. Corpet, J. E. Haber, and G. Almouzni, Nat. Struct. Mol. Biol. **14**, 997 (2007).
[18] C. M. Hammond, C. B. Strømme, H. Huang, D. J. Patel, and A. Groth, Nat. Rev. Mol. Cell Biol. **18**, 141 (2017).
[19] N. Mosammaparast, C. S. Ewart, and L. F. Pemberton, EMBO J. **21**, 6527 (2002).
[20] K. M. Keck and L. F. Pemberton, Biochim. Biophys. Acta **1819**, 277 (2012).
[21] M. Mazzanti, L. J. DeFelice, J. Cohen, and H. Malter, Nature **343**, 764 (1990).
[22] T. Weidemann, M. Wachsmuth, T. A. Knoch, G. Müller, W. Waldeck, and J. Langowski, J. Mol. Biol. **334**, 229 (2003).
[23] J. D. Pajerowski, K. N. Dahl, F. L. Zhong, P. J. Sammak, and D. E. Discher, Proc. Natl. Acad. Sci. U.S.A. **104**, 15619 (2007).
[24] H. Kimura and P. R. Cook, J. Cell Biol. **153**, 1341 (2001).
[25] P. Debye and E. Hückel, Phys. Z. **24**, 185 (1923).
[26] R. Milo, Bioessays **35**, 1050 (2013).
[27] J. C. Lagarias, J. A. Reeds, M. H. Wright, and P. E. Wright, SIAM J. Optim. **9**, 112 (1998).
[28] P. J. Mulligan, E. F. Koslover, and A. J. Spakowitz, J. Phys.: Condens. Matter **27**, 064109 (2015).
[29] A. Y. Grosberg, T. T. Nguyen, and B. I. Shklovskii, Rev. Mod. Phys. **74**, 329 (2002).
[30] D. Lohr, R. T. Kovacic, and K. E. Van Holde, Biochem. **16**, 463 (1977).
[31] A. Shimamura, D. Tremethick, and A. Worcel, Mol. Cell Biol. **8**, 4257 (1988).
[32] J. Mazurkiewicz, F. J. Kepert, and K. Rippe, J. Biol. Chem. **281**, 16462 (2006).
[33] K. Rippe, J. Mazurkiewicz, and N. Kepper, in *DNA interactions with polymers and surfactants.*, edited by R. Dias and B. Lindman (John Wiley & Sons, Hoboken, United States of America, 2008) Chap. 6.
[34] S. Tsonchev, R. D. Coalson, and A. Duncan, Phys. Rev. E **60**, 4257 (1999).
[35] S. Li, H. Orland, and R. Zandi, J. Phys. Condens. Matter **30**, 144002 (2018).
[36] I. M. Gelfand, R. A. Minlos, and Z. Y. Shapiro, *Representations of the rotation and Lorentz groups and their applications.* (Dover Publications, New York, United States of America, 2018).
[37] K. Luger, A. W. Mäder, R. K. Richmond, D. F. Sargent, and T. J. Richmond, Nature **389**, 251 (1997).
[38] J. M. Harp, B. L. Hanson, D. E. Timm, and G. J. Bunick, Acta Crystallogr. Sect. D - Biol. Crystallogr. **D56**, 1513 (2000).
[39] Y. Saad, *Numerical methods for large eigenvalue problems.* (SIAM, Philadelphia, United States of America, 2011).

[40] G. B. Arfken and H. J. Weber, *Mathematical methods for physicists.*, 6th ed. (Elsevier Academic Press, Burlington, United States of America, 2005).
[41] P. Castro-Villarreal and J. E. Ramírez, Phys. Rev. E **100**, 012503 (2019).
[42] M. Rubinstein and R. H. Colby, *Polymer physics.* (Oxford University Press, Oxford, United Kingdom, 2003).
[43] K. N. Dahl, S. M. Kahn, K. L. Wilson, and D. E. Discher, J. Cell Sci. **117**, 4779 (2004).
[44] J. D. Finan, K. J. Chalut, A. Wax, and F. Guilak, Ann. Biomed. Eng. **37**, 477 (2009).
[45] M. Webster, K. L. Witkin, and O. Cohen-Fix, J. Cell Sci. **122**, 1477 (2009).
[46] U. Seifert, Adv. Phys. **46**, 13 (1997).
[47] D. Boal, *Mechanics of the cell.*, 2nd ed. (Cambridge University Press, Cambridge, United Kingdom, 2012).
[48] R. Phillips, J. Kondev, J. Theriot, and H. G. Garcia, *Physical biology of the cell.*, 2nd ed. (Garland Science, New York, United States of America, 2013).
[49] U. Aebi, J. Cohn, L. Buble, and L. Gerace, Nature **323**, 560 (1986).
[50] M. W. Goldberg, I. Huttenlauch, C. J. Hutchison, and R. Stick, J. Cell Sci. **121**, 215 (2008).
[51] J. W. Newport, K. L. Wilson, and W. G. Dunphy, J. Cell Biol. **111**, 2247 (1990).
[52] L. Yang, T. Guan, and L. Gerace, J. Cell Biol. **139**, 1077 (1997).
[53] P. Jevtić, L. J. Edens, L. D. Vuković, and D. L. Levy, Curr. Opin. Cell Biol. **28**, 16 (2014).
[54] R. N. Mukherjee, P. Chen, and D. L. Levy, Nucleus **7**, 167 (2016).
[55] P.-G. de Gennes, *Scaling concepts in polymer physics.* (Cornell University Press, London, United Kingdom, 1979).
[56] V. V. Rybenkov, N. R. Cozzarelli, and A. V. Vologodskii, Proc. Natl. Acad. Sci. U.S.A. **90**, 5307 (1993).
[57] A. L. Nielsen, M. Oulad-Abdelghani, J. A. Ortiz, E. Remboutsika, P. Chambon, and R. Losson, Mol. Cell **7**, 729 (2001).
[58] S. Machida, Y. Takizawa, M. Ishimaru, Y. Sugita, S. Sekine, J.-i. Nakayama, M. Wolf, and H. Kurumizaka, Mol. Cell **69**, 385 (2018).
[59] L. J. Terry, E. B. Shows, and S. R. Wente, Science **318**, 1412 (2007).
[60] I. Andreu, I. Granero-Moya, N. R. Chahare, K. Clein, M. M. Jordán, A. E. M. Beedle, A. Elosegui-Artola, X. Trepat, B. Raveh, and P. Roca-Cusachs, bioRxiv (2021).
[61] A. Bancaud, S. Huet, N. Daigle, J. Mozziconacci, J. Beaudouin, and J. Ellenberg, EMBO J. **28**, 3785 (2009).
[62] J. Irianto, C. R. Pfeifer, R. R. Bennett, Y. Xia, I. L. Ivanovska, A. J. Liu, R. A. Greenberg, and D. E. Discher, Mol. Biol. Cell **27**, 4011 (2016).
[63] C. Bustamante, J. F. Marko, E. D. Siggia, and S. Smith, Science **265**, 1599 (1994).
[64] M. D. Wang, H. Yin, R. Landick, J. Gelles, and S. M. Block, Biophys. J. **72**, 1335 (1997).
[65] Z. Bryant, M. D. Stone, J. Gore, S. B. Smith, N. R. Cozzarelli, and C. Bustamante, Nature **424**, 338 (2003).
[66] S. Forth, C. Deufel, M. Y. Sheinin, B. Daniels, J. P. Sethna, and M. D. Wang, Phys. Rev. Lett. **100**, 148301 (2008).
[67] F. Mosconi, J. F. Allemand, D. Bensimon, and V. Croquette, Phys. Rev. Lett. **102**, 078301 (2009).
[68] A. Evilevitch, L. Lavelle, C. M. Knobler, E. Raspaud, and W. M. Gelbart, Proc. Natl. Acad. Sci. U.S.A. **100**, 9292 (2003).
[69] GRCh38.p13. Genome Reference Consortium. (2019).
[70] A. Catenaccio, Y. Daruich, and C. Magallanes, Chem. Phys. Lett. **367**, 669 (2003).
[71] J. L. Compton, M. Bellard, and P. Chambon, Proc. Natl. Acad. Sci. U.S.A. **73**, 4382 (1976).
[72] V. E. Tate and L. Philipson, Nucleic Acids Res. **6**, 2769 (1979).
